# The Complete Sequence and Comparative Analysis of Ape Sex Chromosomes

**DOI:** 10.1101/2023.11.30.569198

**Authors:** Kateryna D. Makova, Brandon D. Pickett, Robert S. Harris, Gabrielle A. Hartley, Monika Cechova, Karol Pal, Sergey Nurk, DongAhn Yoo, Qiuhui Li, Prajna Hebbar, Barbara C. McGrath, Francesca Antonacci, Margaux Aubel, Arjun Biddanda, Matthew Borchers, Erich Bornberg-Bauer, Gerard G. Bouffard, Shelise Y. Brooks, Lucia Carbone, Laura Carrel, Andrew Carroll, Pi-Chuan Chang, Chen-Shan Chin, Daniel E. Cook, Sarah J.C. Craig, Luciana de Gennaro, Mark Diekhans, Amalia Dutra, Gage H. Garcia, Patrick G.S. Grady, Richard E. Green, Diana Haddad, Pille Hallast, William T. Harvey, Glenn Hickey, David A. Hillis, Savannah J. Hoyt, Hyeonsoo Jeong, Kaivan Kamali, Sergei L. Kosakovsky Pond, Troy M. LaPolice, Charles Lee, Alexandra P. Lewis, Yong-Hwee E. Loh, Patrick Masterson, Kelly M. McGarvey, Rajiv C. McCoy, Paul Medvedev, Karen H. Miga, Katherine M. Munson, Evgenia Pak, Benedict Paten, Brendan J. Pinto, Tamara Potapova, Arang Rhie, Joana L. Rocha, Fedor Ryabov, Oliver A. Ryder, Samuel Sacco, Kishwar Shafin, Valery A. Shepelev, Viviane Slon, Steven J. Solar, Jessica M. Storer, Peter H. Sudmant, Sweetalana, Alex Sweeten, Michael G. Tassia, Françoise Thibaud-Nissen, Mario Ventura, Melissa A. Wilson, Alice C. Young, Huiqing Zeng, Xinru Zhang, Zachary A. Szpiech, Christian D. Huber, Jennifer L. Gerton, Soojin V. Yi, Michael C. Schatz, Ivan A. Alexandrov, Sergey Koren, Rachel J. O’Neill, Evan E. Eichler, Adam M. Phillippy

## Abstract

Apes possess two sex chromosomes—the male-specific Y and the X shared by males and females. The Y chromosome is crucial for male reproduction, with deletions linked to infertility^1^. The X chromosome carries genes vital for reproduction and cognition^2^. Variation in mating patterns and brain function among great apes suggests corresponding differences in their sex chromosomes. However, due to their highly repetitive nature and incomplete reference assemblies, ape sex chromosomes have been challenging to study. Here, using the methodology developed for the telomere-to-telomere (T2T) human genome, we produced gapless assemblies of the X and Y chromosomes for five great apes (chimpanzee, bonobo, gorilla, Bornean and Sumatran orangutans) and a lesser ape, the siamang gibbon. These assemblies allowed us to untangle the intricacies of ape sex chromosome evolution. We found that, compared to the Xs, the ape Ys vary greatly in size and have low alignability and high levels of structural rearrangements. This divergence on the Y arises from the accumulation of lineage-specific ampliconic regions, palindromes, transposable elements, and satellites. Our analysis of Y chromosome genes revealed expansions of multi-copy gene families and signatures of purifying selection. Thus, the Y exhibits dynamic evolution, while the X is more stable. Mapping short-read sequencing data to these assemblies revealed diversity and selection patterns on sex chromosomes of >100 great ape individuals. These reference assemblies are expected to inform human evolution and conservation genetics of nonhuman apes, all of which are endangered species.

## Introduction

Therian X and Y chromosomes are thought to have originated from a pair of autosomes ∼170 million years ago (MYA)^3^. The X chromosome, typically present in two copies in females and one copy in males, has mostly retained the gene content and order from the original autosomal pair^4^. The Y chromosome, typically present in one copy in males, has acquired the sex-determining gene *SRY* and other male-specific genes and mutations, which were fixed by inversions preventing recombination between the Y and the X over most of their lengths^5,6^. Lacking recombination, the Y has contracted in size and accumulated deleterious mutations and repetitive elements, leading to differences in size and gene content between the Y and the X. The recent human ‘T2T’ (gapless and complete) assembly revealed an X chromosome of ∼154 Mb with 796 protein-coding genes^7^, and a Y chromosome of ∼63 Mb with 107 protein-coding genes^8^. In addition to the *pseudoautosomal region*s (PARs), where the Y still recombines with the X, and *ancestral regions*, which originated from the original autosomal pair, the human Y has long *ampliconic regions* with extensive intrachromosomal homology.

Ampliconic regions harbor *palindromes*—long inverted repeats undergoing gene conversion, which counteracts the accumulation of deleterious mutations^9^. Similar to the human Y, the human X possesses PARs^7^, ancestral regions, and several palindromes^10^.

Whereas human sex chromosomes have been recently completely sequenced^7,8^, the full characterization of sex chromosomes in our closest relatives—non-human apes—has remained enigmatic despite their importance in informing human disease and evolution, and species conservation. In general, therian sex chromosomes have been understudied. Due to the haploid nature and high repetitive content of the Y, most previous studies assembled female genomes, omitting the Y altogether^11^. The ape Y chromosomes were sometimes sequenced with targeted methods^6,12,13^ or via shotgun sequencing of male genomes^14,15^, yet such assemblies were usually fragmented, collapsed, and incomplete. The ape X chromosomes were deciphered to a greater level of contiguity (e.g.,^16–18^), but their assemblies, particularly for long satellite arrays, remained unfinished, preventing their complete characterization.

Earlier cytogenetic studies demonstrated lineage-specific amplifications and rearrangements leading to large size variation among great ape Y chromosomes (e.g.,^19^). The initial assemblies of the human and chimpanzee Ys revealed remarkable differences in structure and gene content^6,12^ despite short divergence time, and an acceleration of substitution rates and gene loss on the Y was observed in the common ancestor of bonobo and chimpanzee^15^. The Y chromosome of the common ancestor of great apes had likely already possessed ampliconic sequences and multi-copy gene families^15^, and all ape sex chromosomes share the same evolutionary strata^14^, while experiencing lineage-specific amplification and loss of ampliconic genes^14,15^. This progress notwithstanding, the lack of complete ape sex chromosome assemblies prevented detailed inquiries into the evolution of ampliconic regions, palindromes, segmental duplications, structural variants, satellites, transposable elements (TEs), and gene copy number. Utilizing the experimental and computational methods developed for the T2T assembly of the human genome^8,20^, we deciphered the complete sequences of sex chromosomes from six ape species and studied their structure and evolution.

## Results

### Ape sex chromosome assemblies

To perform a comparative analysis of great ape sex chromosomes, we built genome assemblies for most extant great ape species—chimpanzee, bonobo, western lowland gorilla (later called ‘gorilla’), Bornean orangutan (later called ‘B. orangutan’), and Sumatran orangutan (later called ‘S. orangutan’). We also assembled the genome of an outgroup—the siamang, representing one of four genera of gibbons (lesser apes). The assemblies included two pairs of closely related species: B. and S. orangutans (*Pongo*), which diverged from each other ∼1 MYA, and chimpanzee and bonobo (*Pan*), which diverged from each other ∼2.5 MYA (Table S1). The human lineage diverged from the *Pan*, gorilla, *Pongo*, and gibbon lineages approximately 7, 9, 17, and 20 MYA, respectively (Fig. 1A, Table S1). The studied species differ in their dispersal and mating patterns (Table S2), potentially affecting sex chromosome structure and evolution. We isolated high-molecular-weight DNA from male cell lines for these species (Table S3, Note S1, Note S2) and used it for high-coverage Pacific Biosciences (PacBio) HiFi, Ultra-Long Oxford Nanopore Technologies (UL-ONT), and Hi-C sequencing (Methods). The sequencing depth among samples ranged from 54-109× for HiFi, 28-73× for UL-ONT, and 30-78× for Hi-C (Table S4). We had access to parental DNA for the studied bonobo and gorilla individuals (Table S5) and sequenced it to 51-71× depth with Illumina short-read technology (Table S4).

**Figure 1.**
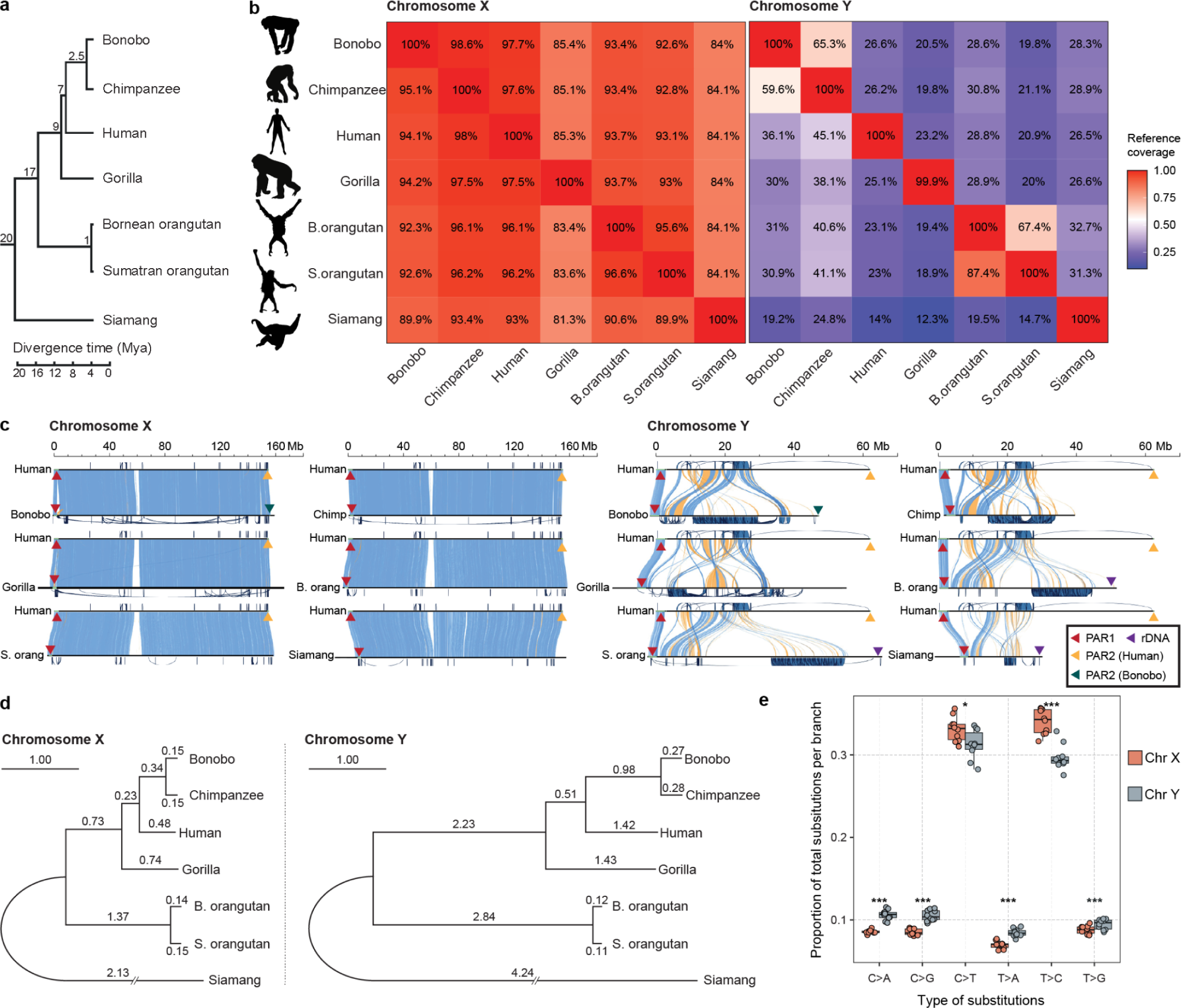
Chromosome alignability and divergence. **(a)** The phylogenetic tree of the studied species (see text and Table S1 for references of divergence times). **(b)** Pairwise alignment coverage of chromosomes X and Y (% of reference, as shown on the x-axis, covered by the query, as shown on the y-axis). **(c)** Alignment of ape sex chromosomes against the human T2T assembly^8,20^. Blue and yellow bands indicate the direct or inverted alignments, respectively, between the chromosomes. Pseudoautosomal regions (PARs) and ribosomal DNA arrays (rDNA) are indicated by triangles (not to scale). Intrachromosomal segmental duplications (SDs) are drawn outside the axes. The scale bars are aligned to the human chromosome. **(d)** Phylogenetic trees of nucleotide sequences on the X and Y chromosomes using Progressive Cactus^78^. Branch lengths (substitutions per 100 sites) were estimated from multi-species alignment blocks including all seven species. **(e)** Substitution spectrum differences between chromosomes X and Y, i.e. a comparison of the proportions of six single-base nucleotide substitution types among total nucleotide substitutions per branch between X and Y (excluding PARs). The distribution of the proportion of each substitution type across phylogenetic branches is shown. The significance of differences was evaluated with a two-sided *t*-test and marked with * for *p*<0.05 and *** for *p*<0.0005. ‘B. orang’ and ‘B. orangutan’ stand for Bornean orangutan, and ‘S. orang’ and ‘S. orangutan’ stand for Sumatran orangutan.

Genome assemblies were generated with Verkko^21^ using the HiFi and UL-ONT data, with haplotypes phased using either parental *k*-mers or Hi-C evidence (Methods). The sex chromosomes were clearly distinguishable from the autosomes in the assembly graphs, with several X and Y chromosomes assembled completely with telomeres on each end (Fig. S2). The remaining sex chromosomes were finished via manual curation and validated, resulting in version 1.1 of the assemblies (Table S6; Methods).

Altogether, we generated T2T assemblies for siamang and B. orangutan X and Y, for which prior assemblies were unavailable, and for bonobo, chimpanzee, gorilla, and S. orangutan X and Y, for which lower-quality assemblies were available^12,15–18^ (Fig. 2). Compared with the previous assemblies, newly generated sequences accounted for 24–45% and 2.6-16% of the total chromosome length on the Y and on the X, respectively (8.6–30 Mb and 3.9–28 Mb of sequence, respectively, Table S7). The sequences gained in these assemblies had a high frequency of motifs able to form non-canonical (non-B) DNA structures (Fig. 2; *p*<2.2×10^-16^ for logistic regressions in each species with previous assemblies, Table S8), which are known to be problematic sequencing targets^22^. Combining sequencing technologies, as done herein, remedies sequencing limitations in such regions^22^.

**Figure 2.**
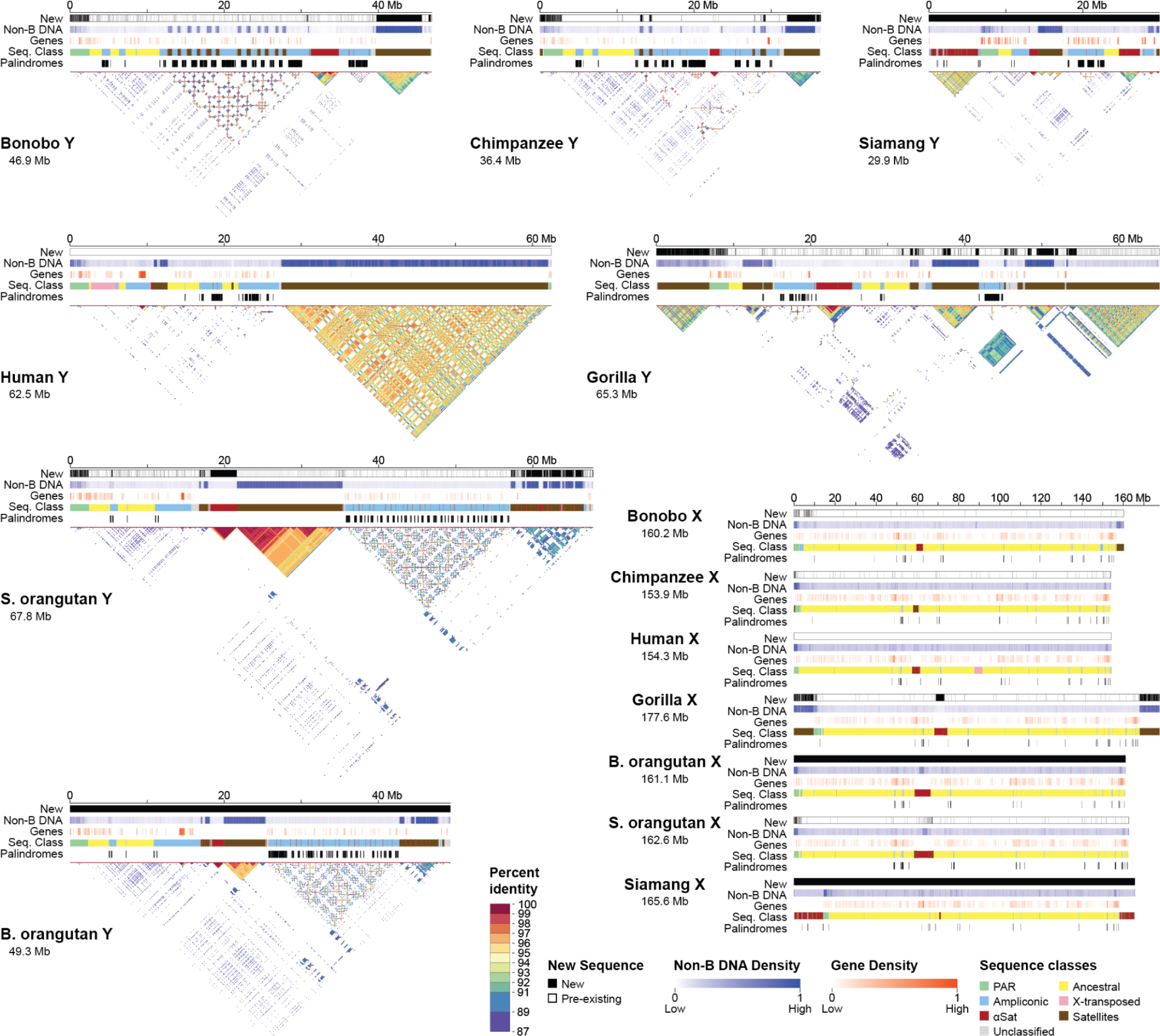
Sequences gained, non-B DNA, genes, sequence classes, palindromes, and intrachromosomal similarity in the assemblies. Tracks for newly generated sequence (in black) relative to previous assemblies (see text), non-B DNA density, gene density (up to 11 genes per 100 kb window), sequence classes (see color legend), and palindromes (in black) are shown. The X and Y are portrayed on different scales. No previous references existed for the Bornean orangutan or siamang, hence the solid black bars for the new sequence tracks. No new sequence was added to the existing T2T human reference in this study and thus the new sequence tracks are empty (white). The gene density tracks are normalized across all species and chromosomes; the non-B DNA density tracks are calibrated independently for each chromosome; in both cases, darker shades indicate higher density. Self-similarity dot plots using a modified version of Stained Glass^124^ are shown for the Y chromosomes (see percent identity legend). ‘B. orangutan’ and ‘S. orangutan’ stand for Bornean and Sumatran orangutan, respectively. In these plots, satellite arrays are visible as blocks of color, directed segmental duplications as horizontal lines, and inverted/palindromic repeats as vertical lines.

The variation in length was larger among the Y than among the X chromosomes across the studied ape species (including human X and Y^7,8^, Fig. 2). Ape Y chromosomes ranged from 30 Mb in siamang to 68 Mb in S. orangutan and differed by as much as 19 Mb between the two orangutans and 11 Mb between bonobo and chimpanzee. The X chromosomes ranged from 154 Mb in chimpanzee and human to 178 Mb in gorilla and differed by only 1.5 Mb between the two orangutans and 6.3 Mb between bonobo and chimpanzee.

### High interspecific variation on the Y

Across all pairwise species comparisons, the percentage of sequence aligned was lower for ape Y than ape X chromosomes (Fig. 1B, Fig. 1C). Only 14–27% of the human Y was covered by alignments to the other ape Y chromosomes, whereas as much as 93–98% of the human X was covered by alignments to the other ape X chromosomes (Fig. 1C). The same pattern was observed for closely related species, with only 60–87% of the Y, but >95% of the X, aligned between them (Fig. 1C).

By analyzing sequence similarity between the X and the Y of the same species, we identified PARs (Fig. 1C; Table S9; Methods), which undergo recombination and thus differ only at the haplotype level between the two sex chromosomes^6^. All species possessed a homologous 2.2–2.5-Mb PAR1, but independently acquired PAR2 sequences were identified in human and bonobo. The PAR2 is ∼330 kb long in human^8^ and ∼95 kb in bonobo (this study) yet they are not homologous (Note S3). The subsequent analyses were conducted excluding PARs unless noted otherwise.

In the sequences experiencing interspecies variation, 83–86% of base pairs on the X and 99% of bases on the Y were affected by large-scale structural variants (SVs; Fig. 1C), while the remaining base pairs were affected by single-nucleotide variants (SNVs; Fig. S3, Fig. S4; Table S10; Methods). Inversions were abundant on the Y (Table S10), consistent with its palindromic architecture. Inversions and insertions were ∼8-fold and 3-fold longer on the Y than the X, respectively (average sizes of 12.1 Mb vs. 1.5 Mb, and 38.2 kb vs. 11.9 kb, respectively; *p*<2.2×10^-16^, Wilcoxon ranked-sum tests). The number of SVs positively correlated with the lengths of phylogenetic branches (Fig. S5, Table S11), with a higher slope for the Y (15.8 SVs/Mb/MY) than for the X (6.1 SVs/Mb/MY), indicating a more rapid accumulation of SVs on the former than on the latter. To identify SVs with potential functional significance in the human lineage, we studied overlaps with genes for 334 and 1,711 human-specific SVs on the Y and the X, respectively (Additional File 1; Table S12). On the Y, we detected an 80-bp deletion disrupting the first exon of *DAZ4*, and an insertion of the previously reported 3.6-Mb X-transposed region (XTR, a human-specific duplication from the X to the Y^6^) including 13 genes. Outside of gene copy-number changes, human-specific inversions affected 11 genes on the Y, and human-specific insertions and deletions affected 23 genes on the X. Thus, SVs represent one of the dominant types of genetic variation on the X and particularly on the Y, and might have functional consequences.

The phylogenetic analysis of multi-species alignments (Methods) for the X, and separately for the Y, revealed the expected species topology (Fig. 1A) but detected higher substitution rates on the Y than on the X for all the branches (Fig. 1D), consistent with male mutation bias^23,24^. For instance, the human-chimpanzee divergence was 2.68% on the Y and 0.97% on the X. For the Y, we detected an 11% acceleration of substitution rates in the *Pan* lineage and a 9.2% slowdown in the *Pongo* lineage, as compared to substitution rates in the human lineage (*p*<10^-5^, relative rate tests; Table S13). For the X, the substitution rates were more similar in magnitude among the branches (Table S13). These results indicate a stronger male mutation bias for the *Pan,* and a weaker bias for the *Pongo,* lineages as compared to that for the human lineage. Strong male mutation bias in the *Pan* lineage is consistent with increased sperm production due to sperm competition (Table S2).

Comparing nucleotide substitution spectra between the two sex chromosomes, we found C>A, C>G, T>A, and T>G substitutions to be significantly more abundant on the Y than on the X, and C>T and T>C substitutions to be more abundant on the X than on the Y (Fig. 1E). These findings are broadly consistent with sex-specific signatures of *de novo* mutations from other studies; C>A, C>G and T>G were shown to be enriched in paternal *de novo* mutations, whereas C>T mutations—in maternal *de novo* mutations^25^. C>G might be related to meiotic double-strand breaks in the male germline^26^.

### Ampliconic regions and palindromes

To study the evolution of sex chromosomes outside of PARs, we separated the assemblies into ancestral, ampliconic, and satellite regions (Methods; Fig. 2; Table S14; Additional File 2). The ancestral regions (also called ‘X-degenerate’ on the Y^6^), which are the remnants of the autosomal past, ranged from 138–147 Mb among species on the X, but were much shorter, from 3.6–7.5 Mb, on the Y, consistent with sequence loss due to the lack of recombination on the latter. We did not find XTRs^6^ on the Y chromosomes of nonhuman apes (Note S4).

Ampliconic regions, defined as long (>90-kb) multi-copy sequences with >50% identity between copies (Methods), ranged from 3.8–6.9 Mb on the X, but were longer on the Y, where they ranged from 9.7–28 Mb and substantially contributed to its length variation among species (Fig. 2, Table S14). These regions were shorter (by 2.5-25 Mb) in previous Y chromosome assemblies^12,15^ than in our T2T assemblies, suggesting their collapse in the former. Ampliconic regions on the X were shared among species to a large degree (Ext. Data Fig. 1A); for instance, we could detect their homology among the African great apes. In contrast, we could detect homology between Y ampliconic regions only in pairs of closely related species—bonobo and chimpanzee, and the two orangutans (Ext. Data Fig. 1B)—yet these regions still differed in organization (Fig. S6), suggesting extremely rapid evolution.

Within ampliconic regions, we located abundant palindromes, defined as >8-kb inverted repeats of sequences with ≥98% identity (i.e. arms) frequently separated by a spacer (Fig. 2, Fig. 3A; Additional File 3; Methods).

**Figure 3.**
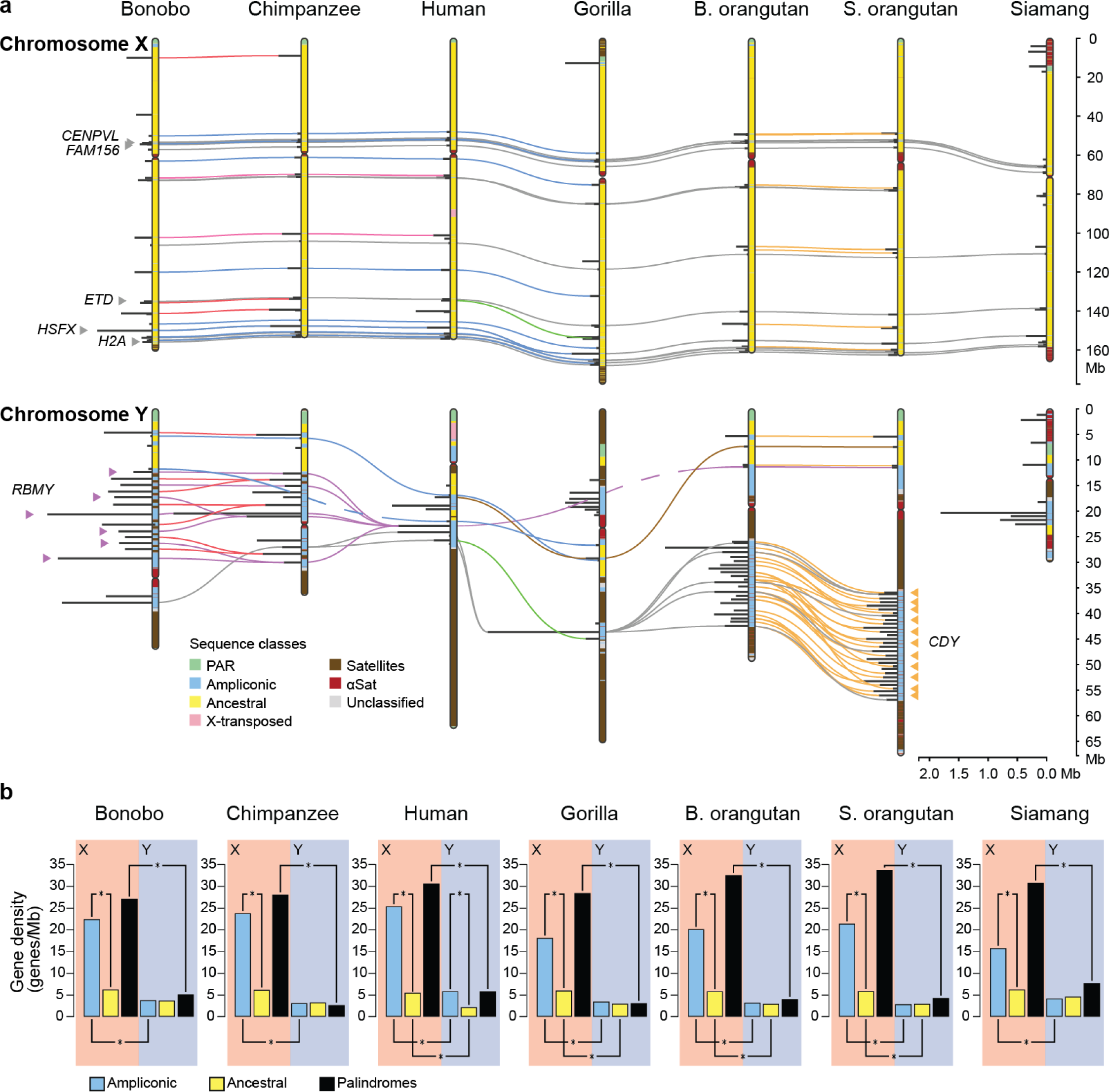
Conservation of palindromes and gene density in different sequences classes. **(a)** All palindromes are shown as horizontal lines perpendicular to the chromosomes (painted with sequence classes), and shared palindromes are connected by colored lines. Several gene families (*CDY* and *RBMY*) that expanded in lineage-specific palindromes are indicated. See Table S41 for the original data. On the X chromosome, several gene families present in palindromes shared among all species are highlighted (*CENPVL1*, *FAM156*, *ETD*, *HSFX*, *H2A*). **(b)** Gene density (in genes per Megabase) for different sequence classes on the X and the Y chromosomes. Only significant comparisons are shown. The significance of differences in gene densities ws computed using goodness of fit (chi-squared) test with Bonferroni correction for multiple tests. See Table S38 for the original data and p-values. The interactive version of the plot can be found at https://observablehq.com/d/6e3e88a3e017ec21.

Palindromes on the Y were on average two to three times longer (Fig. 3A, Fig. S7A, significant *p*-values for one-sided Wilcoxon rank sum tests in most cases, Table S15), and had significantly higher coverage (*p*=2.12×10^-3^ for two-sided Wilcoxon rank sum test, Table S15), than on the X for all species, supporting their role in rescuing deleterious mutations through intrachromosomal recombination and gene conversion on the Y^5,9^. Consistent with gene conversion, we found higher GC content in palindrome arms than spacers on X and Y (*p*=3.08×10^-2^ and *p*=1.04×10^-2^, respectively, for two sample one-sided *t*-tests, Fig. S7B). Palindromes on the X displayed a high degree of conservation among species (Fig. 3A; Table S16); 21 and 10 palindromes were shared among African great apes and among all great apes, respectively. Palindromes on the Y displayed a lower degree of conservation (Fig. 3A; Table S16); two and no palindromes were shared among African great apes and among all great apes, respectively, and numerous palindromes were species-specific or shared by closely related species only.

Segmental duplications (SDs, Methods), defined as >1-kb multi-copy sequences with >90% identity (Methods), constituted 22.8–55.9% of the length of non-human ape Y chromosomes, compared to only 4.0–7.2% of the X chromosomes (Fig. 1C, Table S17). The *Pan* and *Pongo* lineages each showed an almost two times higher percentage of their Y chromosomes occupied by SDs compared to the other ape lineages (average 48.7% vs. 26.6%, *p*=0.057 Mann-Whitney U test). We found little evidence of lineage-specific SDs on the X, but nevertheless observed a gain of up to 2.2 Mb of interchromosomal SDs in the T2T vs. previous X assemblies^16–18^. SDs largely overlapped ampliconic regions and palindromes (Note S5).

### Repetitive element composition and methylation

Our comprehensive annotations (Methods) revealed that 71–85% and 62–66% of Y and X chromosome length, respectively, consisted of repetitive elements (Fig. 4A, Table S18)—TEs, satellites, and simple/low-complexity regions—compared to only 53% of the human T2T autosomal length^27^. On the Y, the repetitive element content (Table S18, Table S19) and distributions (Ext. Data Fig. 2) varied greatly among species, substantially contributing to its length variation. On this chromosome, repeats were predominantly composed of satellites and simple/low complexity regions (Fig. 4A). The repetitive element content was significantly higher in the ancestral than ampliconic regions (∼65.5% vs. ∼46.9%; *p*<0.001, Mann-Whitney U test; Fig. S8, Table S20), reflecting the absence of recombination in the former and frequent intrachromosomal recombination in the latter^28^. The X had a more consistent repetitive element content among species (Fig. 4A, Table S18), composed mainly of retroelements and enriched for L1s^29^ (Table S19). The distributions of repeats along the X were similar among species (Ext. Data Fig. 2), with notable exceptions in the expansion of alpha satellites at the non-centromeric regions in siamang^30,31^, of the SAR/HSat1A satellite in non-human African apes, and of subtelomeric arrays of the pCht/StSat satellite in gorilla^32^ (Fig. 4B, Ext. Data Fig. 2). The repetitive element content of X ancestral regions was significantly lower than that for Y ancestral regions (∼59.3% vs. ∼65.5%; *p*<0.001, Mann-Whitney U test; Fig. S8, Table S20) and significantly higher than that for Y ampliconic regions (∼46.9%; *p*<0.001, Mann-Whitney U test), consistent with different recombination rates among these regions. PARs maintained a consistent repeat content and distribution across the apes analyzed (Ext. Data Fig. 2, Fig. S8, Table S20).

**Figure 4.**
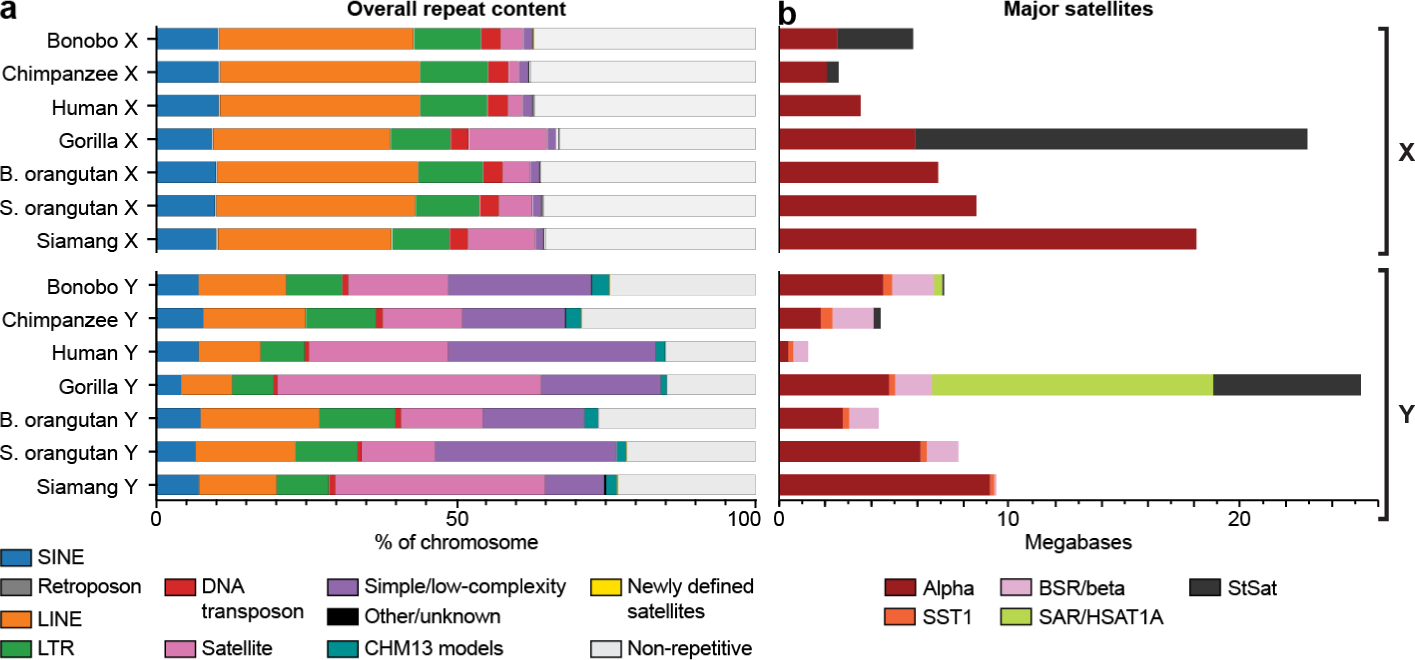
Repeats on ape sex chromosomes. **(a)** Overall repeat annotations across each of the ape sex chromosomes are depicted as a percentage of total nucleotides. Each repeat class is defined by color, with gray representing non-repetitive DNA. Previously uncharacterized human repeats derived from the CHM13 genome analyses are demarcated in teal, adding 0.02% to 2.84% of annotations in each of the non-human ape sex chromosomes. Newly defined satellites (Methods) are depicted in light orange. **(b)** The number of bases on each X and Y chromosome comprising canonical satellites are shown, with each satellite represented by a different color according to the included key. Of note, StSat/pCht and SAR/HSat1A satellites have undergone expansion on gorilla, bonobo, and chimpanzee X and Y chromosomes. Alpha satellites, present in all species, form large subterminal expansions in siamang gibbon (Ext. Data Fig. 2).

We identified 13 previously uncharacterized composite repeats (Fig. S9, Table S21, Table S22), two variants of *DXZ4* repeats, and 33 previously unknown satellites (Fig. S10, Table S23). The latter accounted for an average of 317 kb and 61 kb on each X and Y chromosome, respectively. Variable TE types and satellite arrays, including previously unknown satellites, expanded in a lineage-specific manner (Fig. 4A, Fig. 4B, Fig. S11, Ext. Data Fig. 2, Table S24, Table S25) either via intrinsic TE mobility or other mechanisms. For example, the bonobo-specific satellite Ariel flanked PAR2 in a 318-unit array on the X and a 134-unit array on the Y (Note S3). Lineage-specific expansions on the Y contributed more to interspecies variation than those on the X, but had similar patterns for both sex chromosomes between closely related species (Note S6).

Our T2T assemblies have for the first time allowed us to explore the distribution of motifs able to form non-B DNA structures—A-phased repeats, direct repeats, G-quadruplexes (G4s), inverted repeats, mirror repeats, short tandem repeats (STRs), and Z-DNA^33^—which have been implicated in numerous cellular processes, including replication and transcription^34^. Such motifs (Methods) covered from 6.3–8.7% of the X and from 10–24% of the Y (Methods; Table S26). Each non-B DNA motif type usually occupied a similar fraction (Table S26) and was located in similar regions of the Xs among species, with direct repeats frequently located at the subtelomeric regions and inverted repeats at the centromeric regions (Fig. S12). In contrast, the Ys exhibited a wide range of variation in content and location of different non-B DNA types (Table S26, Fig. S12). Non-B DNA was frequently enriched at satellites (Fig. S13, Table S27), suggesting functional roles. For instance, the LSAU satellite^35^ exhibited an overrepresentation of G4s, where they might function as mediators of epigenetic modifications^36^ consistent with variable methylation levels at this satellite among apes^37^. Across species, we also observed an enrichment of inverted repeats at alpha satellites, consistent with the suggested role of non-B DNA in centromere formation^38^.

Given the strong impact of DNA methylation on repetitive elements and genome composition, we analyzed 5mC DNA methylation (hereafter ‘methylation’) patterns across ape sex chromosomes using long-read data mapped to these T2T assemblies. Previous studies suggested that, in females, the inactive X may have lower global methylation than the active X^39,40^, which is transcriptionally more active and less heterochromatic. We thus hypothesized that, in males, the Y, given its relative transcriptional inactivity^41^ and high heterochromatin content, may have lower global methylation than the active X. In line with this expectation, the Y (excluding PARs) exhibited lower methylation levels than the X in long-range windows (Table S28; Ext. Data Fig. 3A).

DNA methylation was higher for PAR1 than the rest of the X chromosome in all species (Wilcoxon rank-sum test, *p*-values in Table S28; Ext. Data Fig. 3A), which may be due to differences in recombination levels, as methylation is known to be elevated in regions with high recombination rates^42^. Methylation differences between each PAR2 and the rest of the X were not significant (Fig. S14A), likely due to their recent emergence. Methylation levels were significantly higher in ampliconic regions, which undergo intrachromosomal recombination, than ancestral regions in chimpanzee, human, and B. orangutan X chromosomes (Table S28, Ext. Data Fig. 3), but were not significantly different between these two regions on the X of other species and were lower in ampliconic than ancestral regions on the Y (Ext. Data Fig. 3). Thus, the relationship between methylation and recombination might be different for intra-vs. interchromosomal recombination. Most groups of repetitive elements followed the general pattern of highest methylation in PAR1, intermediate in non-PAR X, and lowest in non-PAR Y (Ext. Data Fig. 3B, Table S28). The same pattern was observed in satellites (with the exception of human, which showed non-significant trends), despite their recent and frequent lineage-specific expansions. These patterns suggest rapid evolution of methylation on ape sex chromosomes.

### Evolution of centromere and rDNA arrays

We next examined the evolution of centromeres on X (cenX) and Y (cenY) chromosomes. Previous studies indicated that primate centromere sequences underwent repeated remodeling cycles, in which new variants of 171-bp alpha satellite (AS) repeat monomers emerged and expanded within progenitor arrays, while vestigial layers of old displaced centromeres in the flanks degraded and shrank (Fig. 5A)^43,44^. Indeed, each major primate lineage has active centromeres corresponding to a different AS suprachromosomal family (SF) group. Accordingly, cenXs in African apes are composed of ‘younger’ SF1–3 (Fig. 5B), while the ‘older’ SF5 and yet older SF4 form active centromeres in *Pongo* and siamang lineages, respectively. Further, active arrays on cenX were flanked by older SF vestigial layers in all primates studied (e.g., by SF5, SF4, and SF6–11 in African apes; Fig. 5B)^45,46^. In contrast to cenX, whose chromosomal position has been stable throughout primate evolution, the chromosomal position of cenY is variable, and lacks older flanking layers (Fig. 5B).

**Figure 5.**
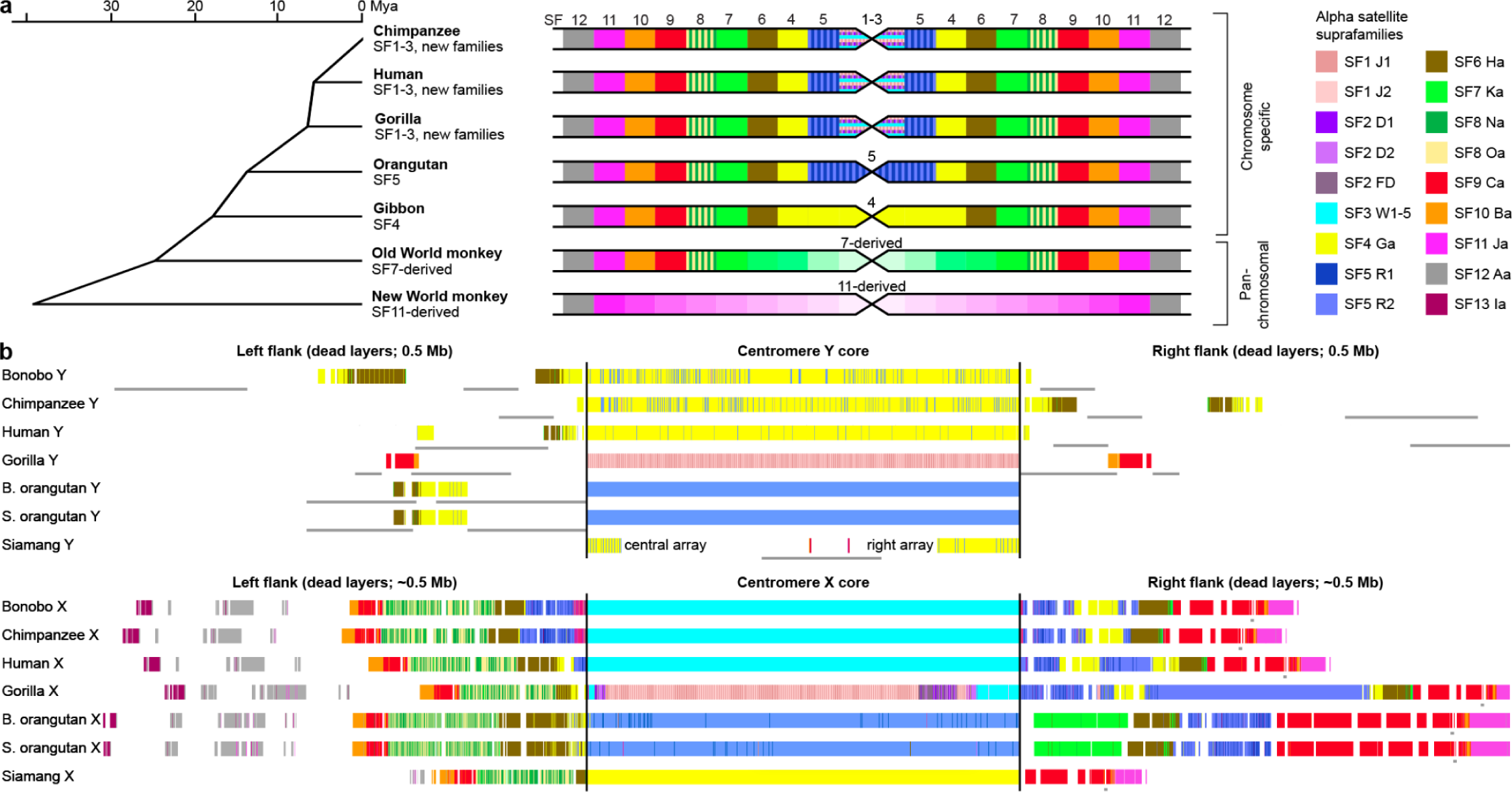
Centromeres on ape sex chromosomes. **(a)** The left panel shows the primate phylogenetic tree with active alpha satellite (AS) suprafamilies (SFs) specified. Chromosome-specific organization indicates that the active centromere in each chromosome has a different higher order repeat (HOR). In pan-chromosomal organization, all centromeres have similar repeats. The right panel shows the generalized centromeres for each branch (not to scale) with SF composition of the active core indicated in the middle and of the dead flanking layers on the sides. Each branch has one or several SFs fewer than in African apes but may have a number of branch-specific layers not shared with the human lineage (shown by hues of the same color). The African ape centromere cores are shown as horizontal bars of SF1-SF3 to represent that each chromosome usually has only one SF, and the SF differs with each chromosome. **(b)** The UCSC Genome Browser tracks showing the SF composition of centromere cores (not to scale) and the flanks for cenY and cenX. CenX is always surrounded by stable vestigial layers, which represent the remnants of dead ancestral centromeres, while cenY has a ‘naked’ centromere devoid of standard monomeric layers. Thin gray lines under the tracks show overlaps with the segmental duplication tracks, which are abundant among cenY flanks. In gorilla cenX, SF3 (cyan) was replaced by SF2 (purple) and then by SF1 (pink). The colors for all AS SFs are listed in the included key. See details in Note S7.

CenY is defined by an older SF4 in human and *Pan* lineages^8,47^, rather than the younger SF1-3 typical of cenX and other African ape centromeres. This ‘lagging’ pattern was not observed in other ape cenYs, which aligned with expectations (Fig. 5B). For example, cenY in gorilla is defined by SF1, and contains motifs important for the binding of centromere protein B (CENP-B boxes; Fig. S15A,E), which are typical of the younger SF1-3.

CENP-B boxes are absent in the SF4 arrays in human and *Pan* cenY. CENP-B is a key component of the inner kinetochore and the absence of CENP-B boxes can impact centromere fidelity and function^48^.

Ape centromeres consist of higher-order repeats (HORs), where subsets of ordered AS monomers are arranged as a larger repeating unit with high sequence similarity between copies (Table S29, Table S30, Note S7, Methods). HORs on cenX and cenY are lineage-specific in apes, with the exception of the shared cenX HOR in human and *Pan*. In closely related species (e.g. chimpanzee and bonobo, or two orangutans) we observed the same HORs; however, their arrays differed in length, structural variant composition (StVs), and centromere dip regions (CDR, or the signature methylation pattern that marks the kinetochore location^46,49^) (Ext. Data Fig. 4, Fig. S15B–C). Further classification of HORs revealed species-specific HOR haplotypes (HORhaps^45,46^) with subtle signatures of array remodeling, comparable to the turnover of SFs (Ext. Data Fig. 4, Fig. S15D, Note S7). Finally, SF4 AS arrays were identified in the siamang in both centromeres and subtelomeric regions^30^. In contrast to the highly similar subtelomeric arrays (Fig. S15F), the non-telomeric arrays in siamang were distinct, or chromosome-specific, similar to the other apes^30,44,50^.

Ribosomal DNA (rDNA) arrays were found on the Y chromosomes of siamang, S. orangutan, and B. orangutan^51,52^, but not on any X chromosomes. Individual UL-ONT reads confirmed the presence of three copies for S. orangutan and one copy for B. orangutan, but were not long enough to span the siamang array. Instead, FISH was used to estimate the size of the siamang array at 16 copies and confirm the absence of rDNA signal on all other sex chromosomes (Ext. Data Fig. 5A-B, Fig. S16, Table S31, Methods). Evidence of active 45S transcription was found for both the siamang and S. orangutan arrays, while the single B. orangutan unit appeared silent (Ext. Data Fig. 5C-E). Beyond the genomes assembled here, we also found rDNA on the Y chromosomes of white-cheeked and black crested gibbons (Note S8).

#### Protein-coding genes

Our gene annotations (Table S32; Methods) indicated the presence of a high percentage of BUSCO genes on the X chromosomes (Table S33), and of most previously known Y chromosome genes (Fig. 6). We manually curated Y chromosome genes (Methods) and validated the copy number of several Y multi-copy gene families with droplet digital PCR (ddPCR; Table S34, Table S35). As a rule, genes were single-copy in ancestral regions and multi-copy in ampliconic regions (Table S36, Table S37). On the X, gene density was ∼2.5–5-fold higher in the ampliconic than ancestral regions (16–25 vs. 5.3–6.1 genes/Mb, respectively; Fig. 3B; Table S38) and was higher still in palindromes (20-32 genes/Mb; Fig. 3B). Palindromes shared among species contained many housekeeping gene families (e.g., *CENPVL*, *H2A*, and *FAM156*; Table S37, Table S38). Gene density was uniformly lower on the Y than on the X (Fig. 3B), with a low density in both ancestral (2.0–4.5 genes/Mb) and ampliconic (2.8–5.7 genes/Mb) regions.

**Figure 6.**
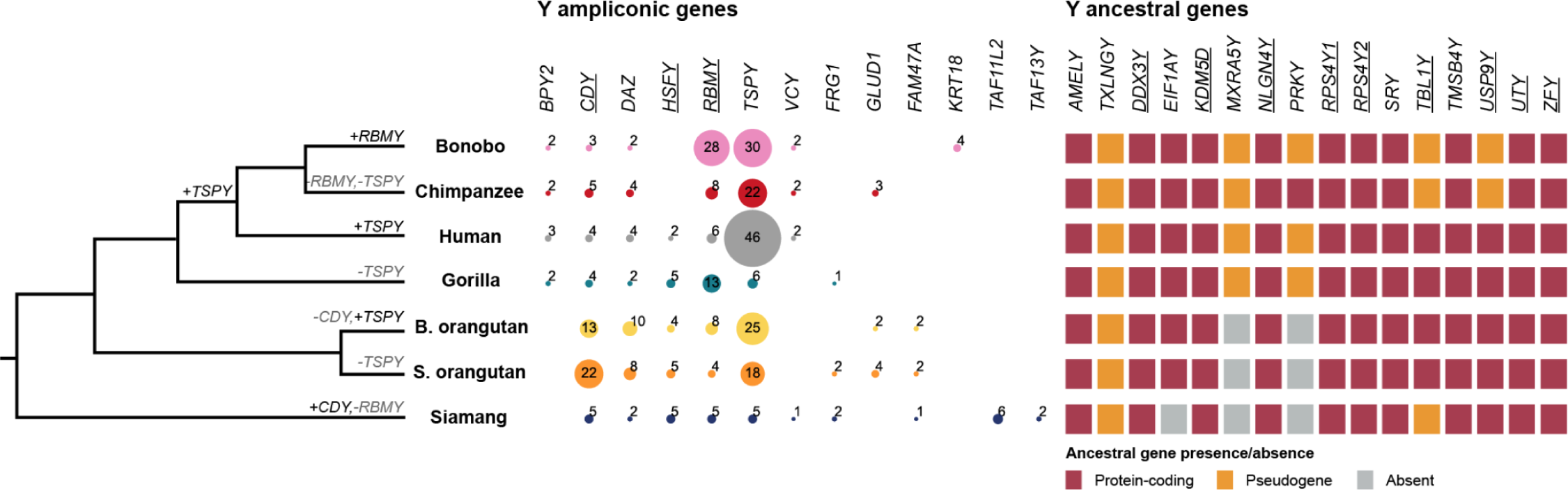
Gene evolution on the Y chromosome. Significant gains and losses in ampliconic gene copy number (Note S10) are shown on the phylogenetic tree. Copy number (noted by number and shown by circle size) or absence of ampliconic genes, and presence, pseudogenization, or absence (i.e., deletion) of ancestral (i.e. X-degenerate) genes, on the Y chromosome. Genes showing signatures of purifying selection (Methods) are underlined. *PRY* and *XKRY* were found to be a pseudogene in all species studied and thus are not shown. The *RBMY* gene family harbored two distinct gene variants, each present in multiple copies, within both orangutan species (Fig. S19).

The ancestral (or ‘X-degenerate’) gene content on the Y was generally well conserved (Fig. 6, Note S9), with the exception of three genes (*TXLNGY*, *MXRA5Y*, and *PRKY*) that were pseudogenized or lost in all or nearly all studied apes (Table S39). Ten ancestral genes were present in all studied apes, and nine out of 13 ancestral genes analyzed exhibited signatures of purifying selection (*p*≤0.05, LRTs; Table S40), suggesting their functional importance. Notably, all four ancestral genes found to be retained in eutherian mammals in another study^4^ were present in apes, and three of them (*DDX3Y*, *UTY*, and *ZFY* but not *SRY*) exhibited a signature of purifying selection, i.e., the nonsynonymous-to-synonymous rate ratio, d_N_/d_S_, below 1 (*p*≤0.05, likelihood ratio tests, LRTs; Table S40).

Among multi-copy genes on the Y and on the X, we detected ampliconic gene families, defined as families with at least two copies having ≥97% sequence identity at the protein level in at least one species (Table S36, Table S37). Many of them were located in palindromes. The proportion of ampliconic among multi-copy gene families was lower on the X than the Y chromosome (29/123 vs. 14/20, *p*=2.68×10^-6^, chi-squared test) highlighting a lower percentage of the X length covered by ampliconic sequences and palindromes (Table S14).

Nevertheless, we still found several copious ampliconic gene families on the X—*GAGE*, *MAGE*, and *SPANX*, the products of which are exclusively or predominantly expressed in testis (Table S37).

Among Y ampliconic gene families, nine were described previously^6,14^ (*BPY2*, *CDY*, *DAZ*, *HSFY*, *RBMY*, *TSPY*, *VCY*, *FRG1*, and *GLUD1*), with the majority functioning in spermatogenesis^6^, and four (*FAM47AY*, *KRT18Y*, *TAF13Y*, and *TAF11L2Y*) are described here for the first time (Fig. 6, Table S36). We discovered that some ampliconic gene copies were present at several palindromes and/or located outside of palindromes (Ext. Data Fig. 6, Table S41). We found episodes of significant lineage-specific expansions and contractions in the previously described ampliconic gene families (Fig. 6, Note S10). For example, *RMBY* expanded in bonobo, *CDY* in S. orangutan, and *TSPY* in human. These results based on one individual per species are largely consistent with prior ddPCR results obtained for multiple individuals per species^41^. *TSPY*—the only ampiconic gene family predominantly located in tandem arrays outside of palindromes (in all species but bonobo and siamang, Table S41)—had a high copy number in all ape species but gorilla and siamang (Fig. 6). A phylogenetic analysis identified mainly species- and genus-specific clades (Ext. Data Fig. 7) with short branches for individual *TSPY* protein-coding copies, suggesting sequence homogenization due to recombination between palindrome arms and/or direct repeats^53^. The newly described families had more limited species distribution and were usually less copious than the previously described families (Fig. 6). We found no evidence of positive selection acting on Y ampliconic gene families (Table S40). A significant signal of purifying selection was detected for only three (*CDY*, *HSFY,* and *RBMY*) out of seven gene families analyzed (*p*≤0.05, LRTs; Table S40). Congruous with an observation for human and macaque^5^, apes had a lower group-mean d_N_/d_S_ for Y ancestral than ampliconic genes (0.38 vs. 0.69, joint model fit, LRT *p*-value<10^-10^), suggesting stronger purifying selection acting on the former genes.

The characteristic DNA methylation levels near the transcription start sites of protein-coding genes (Fig. S14B-C) and their relationship with gene expression (Fig. S14D) implies the importance of promoter hypomethylation in the regulation of gene expression^54^ on both sex chromosomes. Previous studies indicated that *de novo* genes, defined as species- or genus-specific genes arising from noncoding sequences, play a role in fertility and frequently have testis-specific expression^55^. Thus, the Y chromosome is a promising candidate for *de novo* gene formation. Using the new T2T assemblies, we were able to trace the emergence of two candidate *de novo* genes specific to ape Y chromosomes—one in bonobo and one in siamang (Note S11).

#### Intraspecific ape diversity and selection

Our T2T assemblies enabled the first X and Y chromosome-wide analyses of great ape intraspecific diversity. Aligning short sequencing reads from 129 individuals across 11 subspecies (Table S42A) to T2T and previous assemblies (Methods), we detected a higher proportion of reads mapping and a lower mismatch rate to the T2T assemblies in most cases (Ext. Data Fig. 8A, Fig. S17A, Table S42). We further found that the variants identified relative to the T2T assemblies contained fewer SNVs and small indel homozygous variants (Fig. S17B, Table S42), which can arise from structural errors in the reference genome^56^, and largely restored the expected site frequency spectrum (Ext. Data Fig. 8B). However, eastern lowland and mountain gorillas still contained a substantial number of homozygous variants (Fig. S16C), highlighting the need for additional species- and subspecies-specific references. Within the chimpanzee Y, the T2T assembly identified more variants due to the increased length and more uniform read distribution (Ext. Data Fig. 8C). Finally, within a segment of the ampliconic region on gorilla Y, we found a 33-fold reduction in variants (Ext. Data Fig. 8D), likely due to a collapse of this segment in the previous assembly.

Leveraging the more accurate and complete variant calls, we next studied the nucleotide diversity of the different species. Across the X chromosome, the diversity of S. orangutans was higher than that of B. orangutans (*p*<0.001, Mann-Whitney U test, Ext. Data Fig. 8E), in agreement with prior work^57^. In the *Pan* lineage, central chimpanzees retained the highest diversity (*p*-values≤0.01, Mann-Whitney U tests), followed by eastern chimpanzees. Nigeria-Cameroon and western chimpanzees retained a relatively low diversity, potentially signaling historical population bottlenecks^58^. The western lowland gorillas retained a higher diversity than the eastern lowland and mountain gorillas (*p*-values<0.002, Mann-Whitney U tests), both of which have undergone a prolonged population decline^59^. In most subspecies studied, the Y exhibited a substantially lower diversity than the X (*p*-values≤0.01, Mann-Whitney U test, Ext. Data Fig. 8E), as was reported in humans^60^. Among the great apes, bonobos displayed the highest diversity on the Y.

Of particular interest was putative selection on the Y, which can evolve rapidly due to different levels of sperm competition among ape species^6^ (Table S2). We analyzed combined chimpanzee and gorilla samples for nucleotide diversity and Tajima’s D and derived expected values from neutral simulations (Note S12). In gorillas, the observed Y/X diversity ratio was considerably lower than in simulations. In chimpanzees, this ratio aligned with neutrality only at very low male effective population size (N_m_ < 0.25 N_f_). Because male effective population size is high in chimpanzees^61^, this suggests selection reduced diversity on the Y in both species, consistent with reports for humans^60^. Tajima’s D results suggested purifying selection drives this reduction in diversity on the Y in both species (Note S12). Additionally, we identified 45 genes in gorilla and 81 genes in chimpanzee overlapping with candidate regions of selection (Note S12). Finally, incorporating diversity information, we found no evidence of positive selection on Y ancestral genes in chimpanzee and gorilla (Note S13).

## Discussion

Our complete assemblies have revealed the evolution of great ape sex chromosomes in unprecedented detail. In contrast to the X, the Y has undergone rapid evolution in all ape species. It accumulated repetitive elements and experienced elevated rates of nucleotide substitutions, intrachromosomal rearrangements, and segmental duplications, likely due to a loss of recombination over most of its length. It also has reduced global levels of DNA methylation, linked to the low expression levels of many of its genes^41^. Because of this degradation, the Y was suggested to be on its way towards extinction in mammals^2^. Our study suggests that it is still present in apes in part because it harbors several protein-coding genes evolving under purifying selection, similar to observations for rhesus macaque^62^. Future studies should investigate Y non-coding genes and regulatory elements, which may be essential for males and further contribute to selective pressure.

Palindromes are thought to be critical for counterbalancing the degradation of the Y by enabling intrachromosomal recombination and gene conversion^10^. Thus, we expected Y palindromes to be conserved, but instead found many of them to be lineage-specific. Rapid acquisition of new Y palindromes might be due to random genetic drift, which is expected to be strong on the Y due to its small effective population size^63^, and/or due to species-specific selection. Our analysis of Y ampliconic genes, which are primarily located in palindromes and play a role in spermatogenesis, did not provide evidence of species-specific selection. Instead we found a higher ratio of nonsynonymous-to-synonymous mutations for ampliconic vs. single-copy genes. This pattern is consistent with either relaxation of functional constraints or a higher rate of fixation of beneficial mutations due to gene conversion in ampliconic genes^5^—possibilities that should be distinguished by future analyses. Notably, copies of some Y ampliconic genes were present at multiple locations on the Y, and not just within a single palindrome or tandem repeat, providing an additional mechanism safeguarding genes on the non-recombining Y chromosome. The X chromosome also undergoes less recombination than the autosomes as, outside of PARs, it does not recombine in males. We found that it has utilized some of the same strategies to preserve its genetic content, including maintaining palindromes in all apes studied and having ampliconic gene copies at multiple locations.

In addition to gene amplifications, a variety of lineage-specific satellite expansions were observed in the apes, with some specific to the Y (e.g. SAR/HSAT1A accumulation in gorilla Y) and some shared between X and Y (e.g. alpha satellite expansions in siamang). These observations prompt a question about the functionality of these satellites, including the ones enriched in non-B DNA, since such structures may serve as binding sites for protein regulators^34^ and may be involved in defining centromeres^38^. Satellites on the *Drosophila* sex chromosomes were shown to contribute to regulation of gene expression of autosomal genes^64^ and to reproductive isolation among species^65^; similar phenomena should be investigated in apes. Future work is needed to clarify the potential role of satellites in recombination. In some of the species studied here, subtelomeric satellites distal to the PAR were found to be shared between X and Y. If recombination occurs within these satellites, our current PAR annotation will need to be expanded to include them. Additionally, the putative PAR2 sequence discovered in bonobo is flanked by an Ariel repeat that may serve as a cis-acting factor for increased double-strand break formation, as was found for a mo-2 minisatellite in mouse^66^. However, the bonobo PAR2 sequence was also found at the ends of several autosomes (Note S3) and thus might act as a general facilitator of recombination or represent a subtelomeric duplication^67^. The presence of active rDNA arrays on the Y of some ape species also hints at ectopic recombination between the Y and the short arms of the rDNA-bearing acrocentric chromosomes^8,68^.

Mapping intraspecific short-read data from multiple non-human ape individuals revealed intriguing patterns of diversity and highlighted the critical need for collecting additional samples. Further intraspecific studies, comparing the complete sex chromosomes of multiple individuals per species (as was recently done for the human Y^69^) and subspecies are required to reveal the full landscape of ape sex chromosome evolution. Such studies will be useful for investigating sex-specific dispersal and will greatly inform conservation efforts in non-human ape species, all of which are endangered. In humans, both sex chromosomes are important for reproduction^1,2^, genes on the X are also critical for cognition^2^, abnormal X chromosome gene dosage underlies female bias in autoimmune disorders^70^, and X-linked mutations are responsible for 10% of Mendelian disorders^71^, even though the X constitutes only ∼5% of the genome^7^. Thus, we expect these T2T assemblies to be pivotal for understanding disease-causing mutations and human-specific traits.

## Methods

### Sequencing and assemblies

#### Sequencing

We built a collection of male fibroblast and lymphoblastoid cell lines for these species (Table S3, Note S1, Note S2), each karyotyped (Fig. S1) to confirm absence of large-scale chromosomal rearrangements, and isolated high-molecular weight DNA from them. Whole-genome DNA sequencing (WGS) was performed using three different sequencing technologies. To obtain long and accurate reads, Pacific Biosciences (PacBio) HiFi sequencing was performed on a Sequel II with a depth of >60×. To obtain ultralong (>100-kb) reads, ONT sequencing was performed on a PromethION to achieve ≥100 Gb (≥29× depth). To assist with assemblies, paired-end short-read sequencing was performed on Hi-C (Dovetail Omni-C from Cantata Bio) libraries sequenced on Illumina NovaSeq 6000, targeting 400 M pairs of 150-bp reads (≥30× depth) per sample. For bonobo and gorilla parents, we generated paired-end short reads on an Illumina NovaSeq 6000 to achieve ≥518 M pairs of 151 bp reads (≥51× depth) for each sample. Full-length transcriptome sequencing was performed on testes tissue from specimens other than the T2T genome targets (Table S43) using PacBio Iso-Seq on up to three SMRT (8M) cells using a Sequel II.

#### Assemblies

The complete, haplotype-resolved assemblies of chromosomes X and Y were generated using a combination of Verkko^21^ and expert manual curation. Haplotype-specific nodes in the Verkko graphs were labeled using parental-specific *k*-mers when trios were available (bonobo and gorilla) or Hi-C binned assemblies in the absence of trios (chimpanzee, orangutans, and siamang). Haplotype-consistent contigs and scaffolds were automatically extracted from the labeled Verkko graph, with unresolved gap sizes estimated directly from the graph structure (see Rautiainen *et al.*^21^ for more details).

During curation, the primary component(s) of chromosomes X and Y were identified based on the graph topology as visualized in Bandage^72^ and using MashMap^73^ alignments of the assembly to the CHM13 human reference^20^. Several X and Y chromosomes were automatically completed by Verkko and required no manual intervention; for the remainder, manual interventions were employed (Table S6). Using available information such as parent-specific *k*-mer counts, depth of coverage, and node lengths, some artifactual edges could be removed and simple non-linear structures resolved. For more complex cases, ONT reads aligned through the graph were used to generate multiple candidate resolutions, which were individually validated to select the one with the best mapping support. Disconnected nodes due to HiFi coverage gaps were joined and gap-filled using localized, ONT-based Flye^74^ assemblies. The resulting gapless, telomere-to-telomere (T2T) assemblies were oriented based on MashMap alignments to the existing reference genomes of the same or related species (Table S7); in v1.1 of the assemblies, all chromosomes were oriented to start with PAR1.

To validate the T2T assemblies of chromosomes X and Y, we aligned all available read data (Table S4) to the assemblies to measure agreement between the assemblies and raw sequencing data. Specific alignment methods differed for the various data types (Supplemental Methods), but the general principles from McCartney *et al.*^75^ were followed. Validation of the assemblies was done in multiple ways to assess assembly completeness and correctness. Coverage analysis, erroneous *k*-mers, and haplotype-specific *k*-mers (for the two trios) were manually inspected using IGV^76^, and assembly QV was calculated using Merqury^77^. Completeness of each chromosome was confirmed by the identification of telomeric arrays on each end and uniform coverage of long read mappings, with an absence of clipped reads or other observable mapping artifacts.

### Alignments

#### Multi-species whole-chromosome alignments

To estimate the substitution rates on the X and Y chromosomes, we used CACTUS^78^ to generate seven-species multiple alignments, first for the X sequences, and separately for the Y sequences. Sequences were softmasked using repeat annotations (see below). We provided CACTUS with a guide tree, (((((bonobo,chimp),human),gorilla),(sorang,borang)),gibbon), but did not provide branch lengths.

#### Pairwise alignments

To compute the percentage of sequences aligned and to study structural variants and segmental duplications, the pairwise alignment of the human chromosome X and Y was performed against each of chromosome X and Y of the six ape species using minimap2.24^79^. To support other analyses, lastz^80^ was used to compute pairwise alignments of X and Y chromosomes for each species.

### Nucleotide substitution analysis

#### Nucleotide substitution frequency analysis

Substitution rates were estimated (separately for the X and the Y chromosomes) for CACTUS alignment blocks containing all seven species with the REV model implemented in PHYLOFIT^81^.

#### Nucleotide substitution spectrum analysis

Substitution spectrum analysis was conducted using 13-way CACTUS^78^ alignments, which, in addition to the seven studied species, include six ancestral species sequences reconstructed by CACTUS^78^. Triple-nucleotide sequences with 5’ base identical among 13 sequences and 3’ base identical among 13 sequences were used for downstream substitution spectrum analysis. For each branch, 96 types of triple-nucleotide substitutions were grouped into six types based on the middle base substitutions (C>A, C>G, C>T, T>A, T>C and T>G). To compare the distribution of substitution types between chromosome X and chromosome Y and PAR1, we applied *t*-test to the proportions of each substitution type per branch, using Bonferroni correction for multiple testing.

### Segmental duplications and structural variants

#### Segmental duplications (SDs)

The SD content in humans and non-human primates was identified using SEDEF (v1.1)^82^ based on the analysis of genome assemblies soft-masked with TRF v.4.0.9^83^, RepeatMasker^84^, and Windowmasker (v2.2.22)^85^. The SD calls were additionally filtered to keep those with sequence identity >90%, length >1 kb, and satellite content <70%. Lineage-specific SDs were defined by comparing the putative homologous SD loci, defined as containing 10 kb syntenic sequence flanking the SD. The lineage-specific SDs of each species were identified based on non-orthologous locations in the genomes.

#### Structural variants

Structural variants were identified against the human reference genome, CHM13v2.0, via minimap (v2.24) pairwise alignment of ape chromosomes against the human chromosome X and Y^79,86^; 50-bp–300-kb sized SVs with PAV^87^. Larger events were identified and visually inspected using the Saffire SV variant calling pipeline (https://github.com/wharvey31/saffire_sv). The human-specific structural variants were identified by intersecting the variant loci of six ape species; deletions in the six ape species relative to human reference chromosome as putative human-specific insertions, and insertions as putative human-specific deletions. The phylogenetic branch of origin of each SV was predicted using maximum parsimony. As a limitation of this analysis, the SVs for branches including ancestors of the reference species, i.e. human ancestors (i.e. human-chimpanzee-bonobo, human-chimpanzee-bonobo-gorilla, and human-chimpanzee-bonobo-gorilla-orangutan common ancestors) were not computed.

### Palindromes and ampliconic regions

#### Palindrome detection and grouping

We developed *palindrover* in order to screen the X and Y chromosomes for palindromes with ≥98% sequence identity, length ≥8 kb, and spacer ≤500 kb, only keeping candidates with <80% of repetitive content. After aligning the arms with lastz^80^ (alignments with identity <85%, gaps >5%, <500 matched bases, or covering less than 40% of either arm, were discarded), we identified orthologous palindromes and grouped paralogous palindromes on the same chromosome.

#### Overview of the workflow for sequence class annotations

We annotated sequence classes following^6^, with modifications. First, PARs and satellite repeat tracks were created (by aligning X and Y chromosomes for PARs, and by merging adjacent (within 1 kb) RepeatMasker^84^ annotation spanning >0.25 Mb). Next, ampliconic regions were identified as a union of palindromes and regions with high intrachromosomal similarity (i.e. similar to other locations within non-PAR, here identified as consecutive 5-kb windows mapping with ≥50% identity to the repeat-masked chromosomes using blastn from BLAST+ v.2.5.0^88,89^, excluding self-alignments, and spanning >90 kb). The remaining subregions on the Y were annotated as ancestral or ampliconic if overlapping respective genes. Subregions nested within two matching classes were annotated as such.

### Satellite and repeat analysis

#### Satellite and repeat annotations

We produced comprehensive repeat annotations for both X and Y chromosomes across the ape lineage by integrating a combination of known repeats and models identified in human CHM13^27,90^ and T2T-Y^8^, and *de novo* repeat curation (Table S18). To identify canonical and novel repeats on chromosomes X and Y, we utilized the previously described pipeline^27^, with modifications to include both the Dfam 3.6^91^ and Repbase (v20181026)^92^ libraries for each species during RepeatMasker^93^ annotation. A subsequent RepeatMasker run was completed to include repeat models first identified in the analysis of T2T-CHM13 (Table S44), and the resulting annotations were merged. To identify and curate previously undefined satellites, we utilized additional TRF^83^ and ULTRA^94^ screening of annotation gaps >5 kb in length. To identify potential redundancy, satellite consensus sequences generated from gaps identified in each species were used as a RepeatMasker library to search for overlap in the other five analyzed primate species.

Consensus sequences were considered redundant if there was a significant annotation overlap in the RepeatMasker output. Subsequently, final repeat annotations were produced by combining newly defined satellites and 17 variants of pCht/StSat derived from Cechova et al.^95^ and merging resulting annotations. Newly defined satellites that could not be searched using RepeatMasker^93^ due to complex variation were annotated using TRF^83^ and manually added. Tandem composite repeats were identified using self-alignment dotplots and subsequently curated using BLAT^96^ to identify unit lengths and polished using a strategy defined in^97^. Composite repeats were compiled in a distinct repeat annotation track from canonical repeat annotations.

Lineage-specific insertions/expansions were characterized by identifying unaligned regions from CACTUS alignments of the seven primate X and Y chromosomes with halAlignExtract^98^. Unaligned regions were filtered by length and for tandem repeats using TRF^83^ and ULTRA^94^. RepeatMasker^93^ was used to identify the content of the lineage-specific insertions/expansions using the approach described above.

#### Non-B DNA annotations

G-quadruplex motifs were annotated with Quadron^99^, and other types of non-B DNA motifs—with gfa (https://github.com/abcsFrederick/non-B_gfa). To compute non-B DNA density, we used bedtools ‘coverage’ command to count the number of overlaps between each 100 kb window and non-B DNA motifs. We used glm function implemented in R to perform simple and multiple logistic regression to evaluate the relationship between non-B DNA density and sequences gained by the new assemblies. The non-B DNA enrichment analysis for satellites is described in Supplemental Methods.

#### Centromere analysis

To analyze centromeres, we annotated alpha-satellites (AS) and built several tracks at the UCSC Genome Browser (https://genome.ucsc.edu/s/fedorrik/primatesX and https://genome.ucsc.edu/s/fedorrik/primatesY): (1) Suprachromosomal Family (SF) tracks using human-based annotation tools^46^ and utilizing score/length thresholds of 0.7, 0.3, and no threshold; (2) AS-strand track; (3) Higher Order Repeat (HOR) track using species-specific tools specifically designed for this project (https://github.com/fedorrik/apeXY_hmm) and methods described in ^46^; (4) Structural Variation (StV, i.e. altered monomer order) tracks in HORs; (5) CENP-B sites visualized by running a short match search with the sequence YTTCGTTGGAARCGGGA. Other methods are described in Supplemental Methods and Note S7.

### Gene annotations and analysis

#### Gene annotations at the NCBI

The *de novo* gene annotations of the six primate assemblies were performed by the NCBI Eukaryotic Genome Annotation Pipeline as previously described for other genomes^100,101^, between March 20 and May 31, 2023. The annotation of protein-coding and long non-coding genes was derived from the alignments of primate transcripts and proteins queried from GenBank and RefSeq, and same-species (but usually not the same-individual) RNA-seq reads and PacBio Iso-Seq queried from the Sequence Read Archive to the WindowMasker^85^ masked genome. cDNAs were aligned to the genomes using Splign^102^, and proteins were aligned using ProSplign. The RNA-seq reads (Additional File 4), ranging from 673 million (*Pongo pygmaeus*) to 7.3 billion (*Pan troglodytes*) were aligned to the assembly using STAR^103^, while the Iso-Seq reads (ranging from none for *Symphalangus syndactylus* to 27 million for *Gorilla gorilla*) were aligned using minimap2^79^. Short non-coding RNAs, rRNAs, and tRNAs were derived from RFAM^104^ models searched with Infernal cmsearch^105^ and tRNAscan-SE^106^, respectively.

#### Gene annotations at the UCSC

Genome annotation was performed using the Comparative Annotation Toolkit (CAT)^107^. First, whole-genome alignments between the primate (gorilla, chimpanzee, bonobo, Sumatran orangutan, Bornean orangutan, and siamang) and human GRCh38, and T2T-CHM13v2 genomes were generated using CACTUS^78^, as described above. CAT then used the whole-genome alignments to project the UCSC GENCODEv35 CAT/Liftoff v2 (https://cgl.gi.ucsc.edu/data/T2T-primates-chrXY/chm13.draft_v2.0.gene_annotation.gff3) annotation set from CHM13v2 to the primates. In addition, CAT was given Iso-Seq FLNC data to provide extrinsic hints to the Augustus PB (PacBio) module of CAT, which performs *ab initio* prediction of coding isoforms. CAT was also run with the Augustus Comparative Gene Prediction (CGP) module, which leverages whole-genome alignments to predict coding loci across many genomes simultaneously (i.e. gene prediction). CAT then combined these *ab initio* prediction sets with the human gene projections to produce the final gene sets and UCSC assembly hubs used in this project.

#### Curation and analysis of ancestral (X-degenerate) genes

For the Y chromosome, we collected annotations from the NCBI Eukaryotic Genome Annotation Pipeline (RefSeq), CAT, and Liftoff. We extracted ancestral gene annotations from each and mapped them onto the Y chromosome sequence for each in Geneious^108^. We identified that every gene was present and manually curated an annotation set with the most complete exonic complement across annotations. We extracted all CDS regions for each gene and aligned them. For the X chromosome, we extracted ancestral gene copies from the RefSeq annotations using gffread^109^ and aligned them. All alignments were examined and curated by eye, and missing genes and exons were confirmed using BLAST^89^. All present genes were aligned to their orthologs and their gametologs, where we identified genes with significant deviations (truncations of 20% or greater) relative to known (functional) Y copies in other ape species, or their X chromosome counterpart, as pseudogenes (Table S39). These alignments were also used to identify gene conversion events using GeneConv^110^ and to detect selection (see below).

#### Detection of multi-copy and ampliconic gene families

We used blastp for all protein sequences of all protein-coding genes (as annotated by NCBI) against a blast database built from these sequences, separately for the X and the Y chromosome. To infer homology we used a cutoff of 50% sequence identity of at least 35% of protein lengths^111^. We then clustered genes into multi-copy families using a simplified single linkage approach (if genes A and B shared similarity and so did genes B and C, we created a group of genes A, B, and C). To overcome the shortcomings of this method we removed gene clusters where no genes within one species shared high enough similarity.

For each multi-copy gene family we collected the counts of occurrences of gene copies, the sequence classes assigned to the regions where these copies occur, and all pairwise identities of gene copies within one species (Table S36, Table S37). Among multi-copy gene families we then delineated ampliconic families as those that had ≥97% sequence identity between at least two copies in a family in at least one species, which we chose because it was a natural breakpoint in the pairwise sequence identity distribution for Y multi-copy genes (Fig. S20). This method identified all previously known Y ampliconic gene families (*BPY2*, *CDY*, *DAZ*, *HSFY*, *RBMY*, *TSPY*, *VCY*, *FRG1*, and *GLUD1*), as well as four new ones (*FAM47A*, *KRT18*, *13Y*, and *TAF11L2*).

#### Curation of ampliconic genes

We first collected annotations from the NCBI annotation pipeline, CAT, and Liftoff. To these annotations, we added mappings from human and species-specific gene sequences onto the latest assemblies and included the full Iso-Seq transcripts. To combine these annotations, we first performed an interval analysis to find all annotated, mapped, or predicted copies, with one or more sources of evidence and then manually curated the final set of protein-coding and pseudogene copies for each of these genes (Table S45).

#### ddPCR ampliconic gene copy number validations

Copy numbers were determined with ddPCR using the protocols described in ^13^ and ^41^. The sequences of the primers for bonobo, chimpanzee, gorilla, Bornean and Sumatran orangutan were from ^41^. The primers for siamang were designed using Geneious Prime software^108^ and are available in Table S34. ddPCR conditions are described in Table S35.

#### *TSPY* gene analysis

The UCSC table browser was used to retrieve and export the TSPY sequences. For every genome, the appropriate gene annotation dataset was selected with the specific regions defined using the locations of the curated *TSPY* copies. The sequence was retrieved in the 5’ UTR, CDS exons, 3’ UTR, and intron regions and the generated fasta files were then used for alignment with MAFFT v7.520^112^.

Maximum-likelihood phylogenies were inferred using IQTree (v2.0.3)^113^ with the best-fit substitution model estimated by ModelFinder^114^ (best-fit model according to BIC: TVM+F+G4). Node support values were estimated using 10,000 ultrafast bootstrap replicates^115^ with hill-climbing nearest neighbor interchange (`-bnni` flag) to avoid severe model violations. Nodes with <95% ultrafast bootstrap support were collapsed as polytomies.

#### Estimating rDNA copy number and activity by FISH and immuno-FISH

Chromosome spreads were prepared and labeled as described previously^116^. To estimate rDNA copy number and activity from FISH and ImmunoFISH images, individual rDNA arrays were segmented, the background-subtracted integrated intensity was measured for every array, and the fraction of the total signal of all arrays in a chromosome spread was calculated for each array. Similarly, the fraction of the total UBF fluorescence intensity, indicative of RNA PolI transcription^117^, was used to estimate the transcriptional activity of the chrY rDNA arrays. The total rDNA copy number in a genome was estimated from Illumina sequencing data based on *k*-mer counts. Full details are available in Supplemental Methods.

#### Gene-level selection using interspecific fixed differences

To detect selection from interspecific comparison of gene sequences, we started with alignments of ancestral or ampliconic genes, using one consensus sequence per species for ampliconic gene families that were present in at least four species. For these alignments, we inferred ML phylogeny with raxml-ng (GTR+G+I, default settings otherwise), and looked for evidence of gene-level episodic diversifying selection (EDS) using BUSTED with site-to-site synonymous rate variation and a flexible random effects branch-site variation for d_N_/d ^118,119^. Because all alignments were relatively short, we also fitted the standard MG94 + GTR model where d_N_/d_S_ ratios were constant across sites and were either shared by all branches (global model) or estimated separately for each branch (local model). We tested for d_N_/d_S_ ≠ 1 using a likelihood ratio test (global model). To investigate branch-level variability in d_N_/d_S_, we used a version of the local model where all branches except one shared the same d_N_/d_S_ ratio and the focal branch had its own d_N_/d_S_ ratio; *p*-values from branch-level d_N_/d_S_ tests were corrected using the Holm-Bonferroni procedure. Finally, to compare mean in global d_N_/d_S_ between ampliconic and ancestral genes, we performed a joint MG94 + GTR model fit to all genes, with the null model that d_N_/d_S_ is the same for all genes, and the alternative model that d_N_/d_S_ are the same within group (ampliconic or ancestral), but different between groups. All analyses were run using ^120^.

### Methylation analysis

#### CpG methylation calling

To generate CpG methylation calls, Meryl^77^ was used to count *k*-mers and compute the 0.02% most frequent 15-mers in each ape draft diploid assembly. ONT and PacBio reads were mapped to the corresponding draft diploid assemblies with Winnowmap2^121^ and filtered to remove secondary and unmapped reads. Modbam2bed (https://github.com/epi2me-labs/modbam2bed) was used to summarize modified base calls and generate a CpG methylation track viewable in IGV^122^.

#### Methylation analysis

Using the processed long-read DNA methylation data to analyze large sequence classes (PAR1, Ampliconic regions, ancestral regions), we split these regions into 100-kb bins and calculated mean methylation levels of all CpGs within each bin. For smaller sequence classes, such as specific repetitive elements, we generated mean methylation levels from individual elements themselves. For human data, we added another filtering step to remove regions where two long-read sequencing platforms yielded highly divergent results (mostly Yq12 region); non-human methylation data were concordant between the two sequencing platforms (Fig. S18) and thus were used in their entirety. Promoters were defined as regions 1 kb upstream of the transcription start site.

#### Diversity analysis

We collected short-read sequencing data from 129 individuals across 11 distinct great ape subspecies (Table S42A) and aligned the reads to previous (using the previous reference of S. orangutan reference for B. orangutan data) and T2T sex chromosome assemblies. We next performed variant calling with GATK Haplotype Caller^123^, conducted joint genotyping with GenotypeGVCFs^123^, and removed low-confident variants. To further enhance the accuracy and completeness of variant detection, we adopted the masking strategy proposed by the T2T-CHM13v2.0 human chrY study^8^, in which PARs and/or Y chromosome were masked in a sex-specific manner. After generating karyotype-specific references for XX and XY samples, we realigned the reads of each sample to the updated references and called variants. The new variant set was validated reconstructing the Y chromosome phylogeny and estimating the time-to-most-recent common ancestor on it (Note S14). Using the complete variant call sets, we quantified the nucleotide diversity of each subspecies with VCFtools. For chromosome X, we assessed the diversity in PAR and ancestral regions. For chromosome Y, we computed the nucleotide diversity in ancestral regions.

## Data Availability

The raw sequencing data generated in this study have been deposited in the Sequence Read Archive under BioProjects PRJNA602326, PRJNA902025, PRJNA976699-PRJNA976702, and PRJNA986878-PRJNA986879. The genome assemblies and NCBI annotations are available from GenBank/RefSeq (see Table S46 for accession numbers). The CAT/Liftoff annotations are available in a UCSC Genome Browser Hub: https://cgl.gi.ucsc.edu/data/T2T-primates-chrXY/. The reference genomes, alignments and variant calls are also available within the NHGRI AnVIL: https://anvil.terra.bio/#workspaces/anvil-dash-research/AnVIL_Ape_T2T_chrXY. The alignments generated for this project are available at: https://www.bx.psu.edu/makova_lab/data/APE_XY_T2T/ and https://public.gi.ucsc.edu/~hickey/hubs/hub-8-t2t-apes-2023v1/8-t2t-apes-2023v1.hal (with the following additional information: https://public.gi.ucsc.edu/~hickey/hubs/hub-8-t2t-apes-2023v1/8-t2t-apes-2023v1.README.md). Additional Data Files include human-specific structural variant coordinates (File 1), sequence class coordinates (File 2), palindrome coordinates (File 3), and RNA-Seq and IsoSeq datasets used for gene annotations (File 4). Primary data related to the cytogenetic evaluation of the rDNA is deposited in the Stowers Institute Original Data Repository under accession LIBPB-2447: https://www.stowers.org/research/publications/libpb-2447.

## Code Availability

The source code created to generate the results presented in this paper is publicly available on GitHub (https://github.com/makovalab-psu/T2T_primate_XY) and provided at Zenodo at DOI 10.5281/zenodo.10680008. All external scripts and programs are also linked through this GitHub repository.

## Acknowledgements

We would like to acknowledge Rebeca Campos-Sanchez, Francesca Chiaromonte, Tamara Goldfarb, Bernard de Massy, Terence D. Murphy, Morgan Park, Shashikant Pujar, Cynthia Steiner, Dylan J Taylor, Marta Tomaszkiewicz, and Allison Watwood for their assistance and/or advice. We are grateful to Bernadette Weissensteiner and Kate Anthony who assisted with primate cell culture, to PSU Genomics Core Facility, PSU Sartorius Cell Culture Facility, and PSU College of Medicine Genome Sciences Core Facility for their technical assistance, and to San Diego Zoological Society Frozen Zoo and Tissue and DNA collection, Coriell Institute, Smithsonian Institute, University of Texas MD Anderson Cancer Center, and Tulsa Zoo for providing samples and/or cell lines used in this study. This work utilized the computational resources of the NIH HPC Biowulf cluster (https://hpc.nih.gov), and of the Computational Biology Core and sequencing at the Center for Genome Innovation, both in the Institute for Systems Genomics at the University of Connecticut. This work was supported, in part, by the Intramural Research Program of the National Human Genome Research Institute, National Institutes of Health (NIH, B.D.P, S.N., G.G.B., S.Y.B., A.D., E.P., A.R., S.J.S., A.S., A.C.Y., S.K., A.M.P.), by the National Center for Biotechnology Information of the National Library of Medicine, NIH (F.T.-N., D.H., P.M., K.M.M.), by the NIH awards R35GM151945 (to K.D.M.), HG002385 and HG010169 (to E.E.E.), R01GM146462 (to P.M.), R01CA266339 (to J.L.G.), R01GM123312 (to R.O.), R35GM146926 (to Z.A.S), R35GM146886 (to C.D.H.), R01HG011641 (to S.V.Y.), U01CA253481 and U24HG010263 (to M.C.S.), R35GM124827 (to M.A.W.), R01HG011274 (to K.H.M.), and HG007497 (to C.L. and E.E.E.), by the National Science Foundation awards 2138585 and 1931531 (to P.M.), EF-2204761 (to S.V.Y.), and by the Center for Integration in Science of the Ministry of Aliyah, Israel (IAA). K.H.M. is a Searle Scholar, E.E.E. is an investigator of the Howard Hughes Medical Institute. T.M.L. was supported by the NIH T32 GM102057 Computation, Bioinformatics, and Statistics (CBIOS) Training Program Grant at Penn State Uniuversity.

## Author Contributions

V.A.S. is currently retired. B.D.P. performed computational validations, NCBI submissions, chimpanzee subspecies identification, biosample registration, figure generation, and overall project and consortium coordination. R.S.H. generated alignments, identified pseudoautosomal boundaries and palindromes, including their sharing, and performed substitution analysis. M.C. classified assemblies into sequence classes and identified ampliconic regions, and performed palindrome analysis. G.A.H., P.G.S.G., J.M.S., R.J.O. and S.J.H. performed repetitive element annotation, manual curation, analyses and dfam submissions, J.M.S. performed lineage-specific repeat analyses, and G.A.H. generated tracks for figures. K.P. performed gene density analyses, visualized palindrome sharing, identified multi-copy and ampliconic gene families. S.N. and S.K. performed sequence assemblies. G.H. and B.P. generated multi-species alignments. A.S. generated dotplots. S.J.S. performed rDNA array copy number estimation, base calling, and alignment, and generated methylation tracks. D.A.Y., W.T.H., and H.J. performed segmental duplication and structural variation analyses. Q.L., A.B., M.C.S., R.C.M.C., M.G.T., C.D.H., T.M.L.P., S., Z.A.S., P.Ha., C.L., and S.L.K.P. performed diversity and selection analyses. K.P., P.He., F.T.N., D.H., P.M., M.A.W., B.J.P., M.G.T., and M.D. performed gene annotations and analyses. K.K. performed non-B DNA analysis. X.Z. performed substitution spectrum analysis and assisted in figure preparation. D.E.C., K.S., P.C.C., and A.C. performed DeepConsensus calling. M.A. and E.B. performed de novo gene analysis. C.S.C. analyzed palindrome structure in orangutans. P.H.D. and J.L.R. provided HiFi data for bonobo. I.A.A., F.R., V.A.S., V.S., and K.H.M. performed centromere analysis. S.V.Y., D.A.H., and Y.H.E.L. performed methylation analysis. T.P., M.B., and J.L.G. performed rDNA analysis. A.D. and E.P. generated karyotypes. G.A.H., Lu.C. and R.J.O. confirmed the siamang karyotype. L.D.G. and M.V. performed karyotype confirmation and FISH analysis on rDNA. H.Z. performed ddPCR and maintained cell culture. A.C.Y., S.Y.B., and G.G.B. generated UL-ONT and Illumina sequences. S.S. and R.E.G. generated HiC libraries. K.M.M., A.P.L., and G.H.G. generated HiFi and IsoSeq PacBio sequences. A.R., P.M., and S.J.C.C. participated in project discussions, S.J.C.C. also collected gene ontology and mating system information, and A.R. performed methylation comparison between two sequencing platforms. La.C., Lu.C., and O.A.R. provided samples. Lu.C. also provided karyotype confirmation. B.C.M.G. coordinated project resources, maintained cell culture, performed ddPCR, and RNA extractions. K.D.M., E.E.E., and A.M.P. provided project leadership and coordination, and are co-leading the Primate T2T consortium. K.D.M. wrote the manuscript with contributions from the other authors.

## Additional Information

Supplementary Information is available for this paper: Supplementary Figures, Supplementary Tables, Supplementary Methods, Supplementary Notes, and Additional Data Files.

---

**Extended Data Figure 1.**
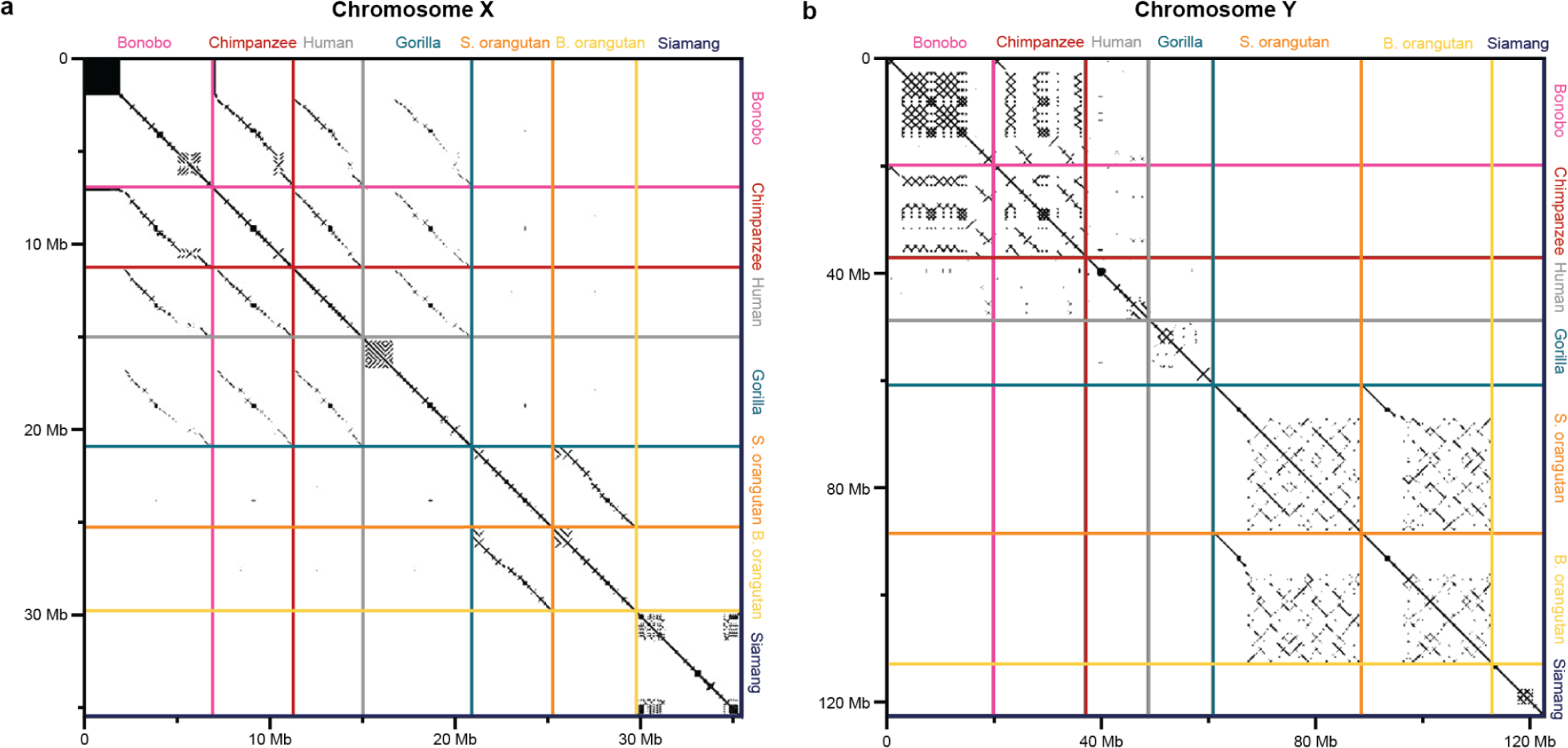
Conservation of ampliconic regions across species. A comparison of ampliconic regions on the (**A**) X chromosomes and (**B)** Y chromosomes between species with similarities highlighted using a dot plot analysis. Ampliconic regions were extracted and concatenated independently for each species and visualized with gepard^125^ using a window size of 100.

**Extended Data Figure 2.**
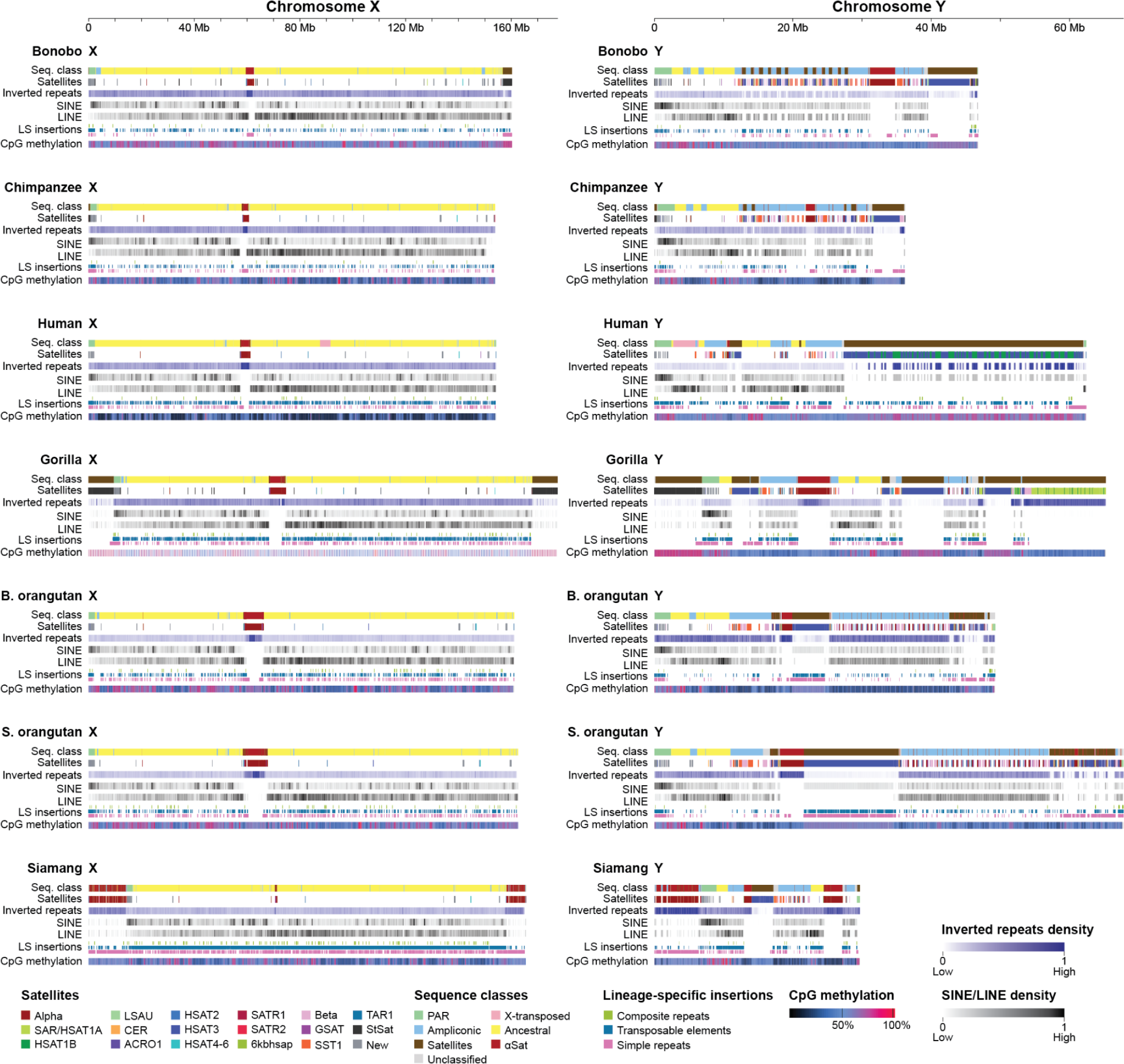
Repeats and satellites on the X and Y chromosomes. Repeats and satellites shown with sequence class annotations and CpG methylation for chromosomes X and Y. The scales are different between chromosomes X and Y. The tracks for each species are: (1) sequence class annotation, (2) satellites, (3) inverted repeats, (4) SINEs, (5) LINEs, (6) lineage-specific (LS) insertions of composite repeats (green), transposable elements (blue), and satellites, simple repeats, and low-complexity repeats (pink), and (7) CpG methylation. The inverted repeat, SINE, and LINE tracks are plotted in blocks with darker colors representing a higher density (density values are calibrated independently for each chromosome/species). CpG methylation is also displayed on a gradient between dark blue (low methylation) and magenta (high methylation) based on the percentage of supporting aligned ONT reads. The remaining tracks (sequence class, satellites, and LS insertions) are displayed as presence/absence (color/no color). The class and satellite tracks are discrete, whereas the LS insertions are plotted as mini tracks to avoid overplotting where >1 label applies.

**Extended Data Figure 3.**
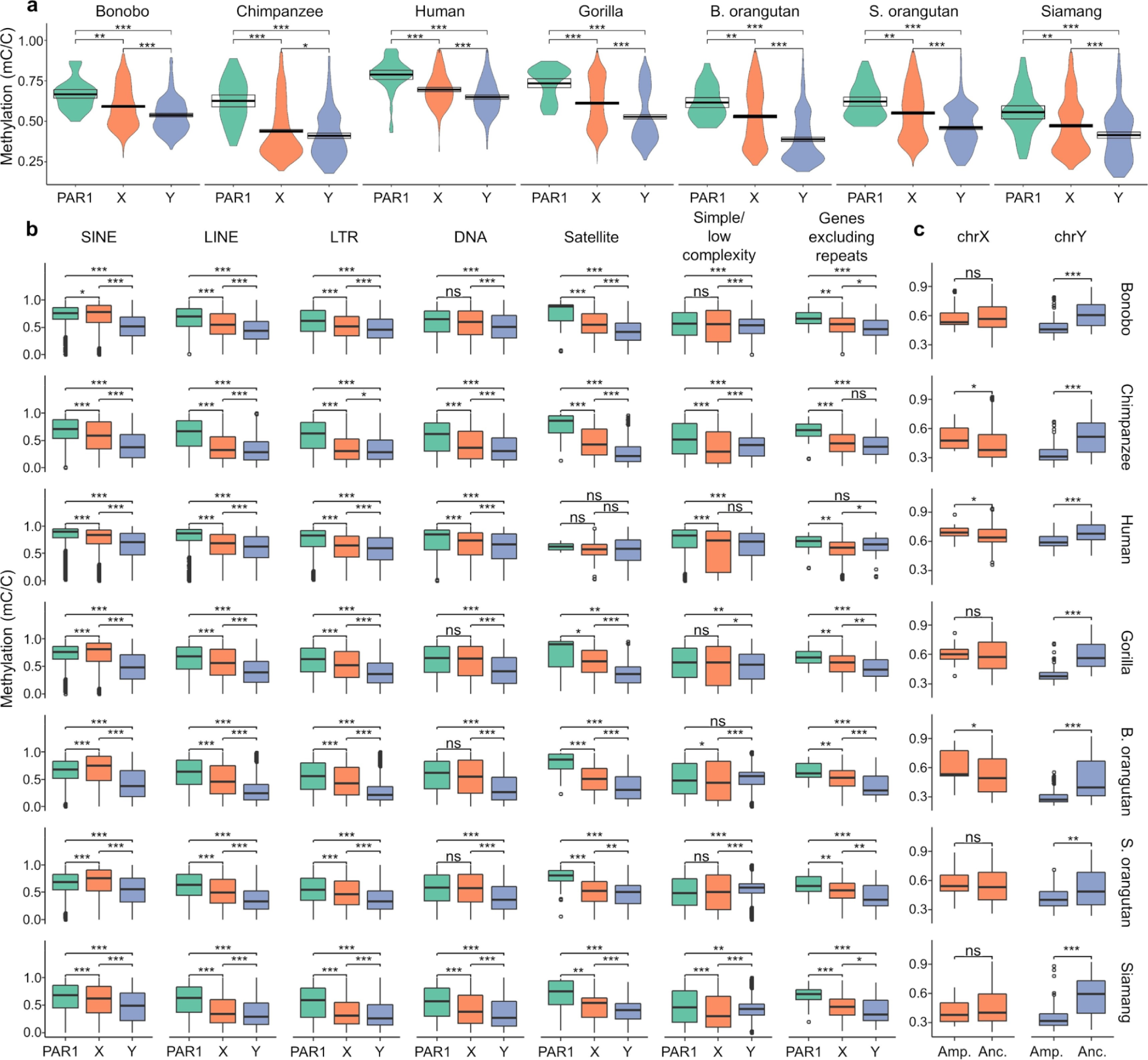
Methylation patterns. **(a)** DNA methylation levels in Pseudoautosomal region 1 (PAR1; teal), non-PAR chromosome X (orange), and non-PAR chromosome Y (periwinkle). *p*-values were determined using Wilcoxon rank-sum tests. * *p*<0.05; ** *p*<10^-3^; *** *p*<10^-6^. (**b**) Differences in DNA methylation levels between different repeat categories as well as protein-coding genes (after excluding repetitive sequences). (**c**) Differences in methylation levels between ampliconic and ancestral regions in the X and the Y chromosomes (in 100 kb bins).

**Extended Data Figure 4.**
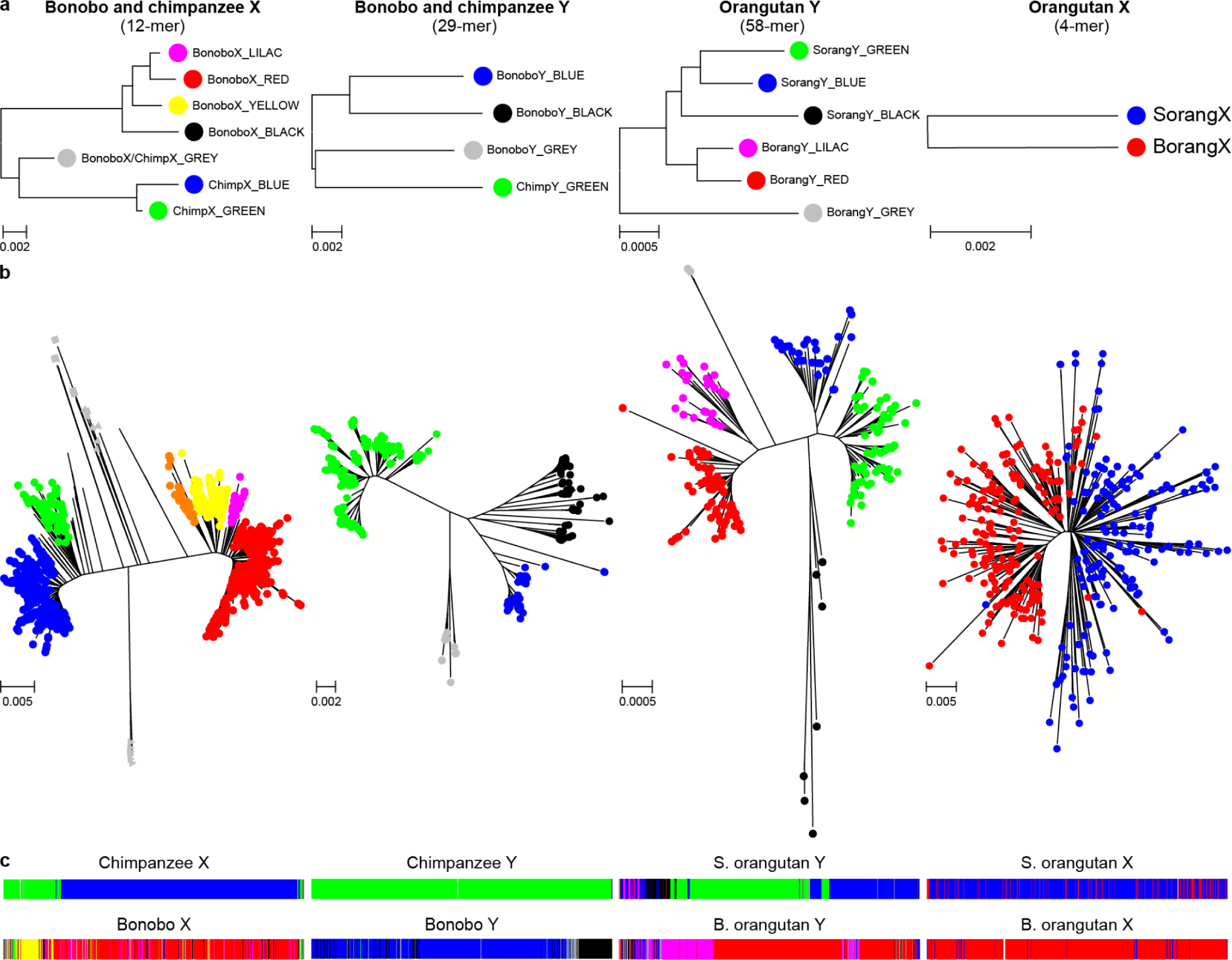
Alpha satellite higher order repeat (HOR) haplotypes are species-specific in *Pongo* and *Pan* (except for the few distal HOR copies) **(a)** Consensus HOR haplotype (HORhap) phylogenetic trees, **(b)** HOR trees, and **(c)** HORhap UCSC Genome Browser annotation tracks for active alpha satellite arrays of chromosomes X and Y in two *Pan* and two *Pongo* species (see Methods in Note S7) are shown. Each colored branch in a HOR tree represents a HORhap. All branches in HOR trees are species-specific, except for the GREY cluster in *Pan* cenX tree, where mixing of chimpanzee (square markers) and bonobo (triangle markers) HORs were observed (Note S7). Each branch was extracted to obtain HORhap consensus sequence and HMM further used in HMMER-based HORhap classification tool^46^ to produce HORhap annotations. The larger branches with shorter twigs correspond to the younger large active HORhap arrays; the smaller branches with longer twigs correspond to the older and smaller side arrays. The thinnest and longest branches make up the oldest and smallest peripheral arrays which often cannot be seen in the track panels. The *Pongo* X tree has a ‘star-like’ shape and does not have obvious HORhaps; HORs colored by species indicate almost no mixing between species and species-specific consensus sequences show three consistent differences (Fig. S15D). Thus, we concluded that the species did not share the same HORhaps, but no significant divides could be seen in the tree due to the short HOR length (a 4-mer), as detailed in Note S7. The age of the HORhaps is also confirmed by consensus trees where the oldest GREY twigs branch out closer to the root and are nearly equidistant to the active HORhap branches of respective species. Hence they likely resemble the HORs that existed in the common ancestor of both species. Thus, all but the oldest HORhaps are species-specific and indicate considerable evolution that occurred after the species diverged.

**Extended Data Figure 5.**
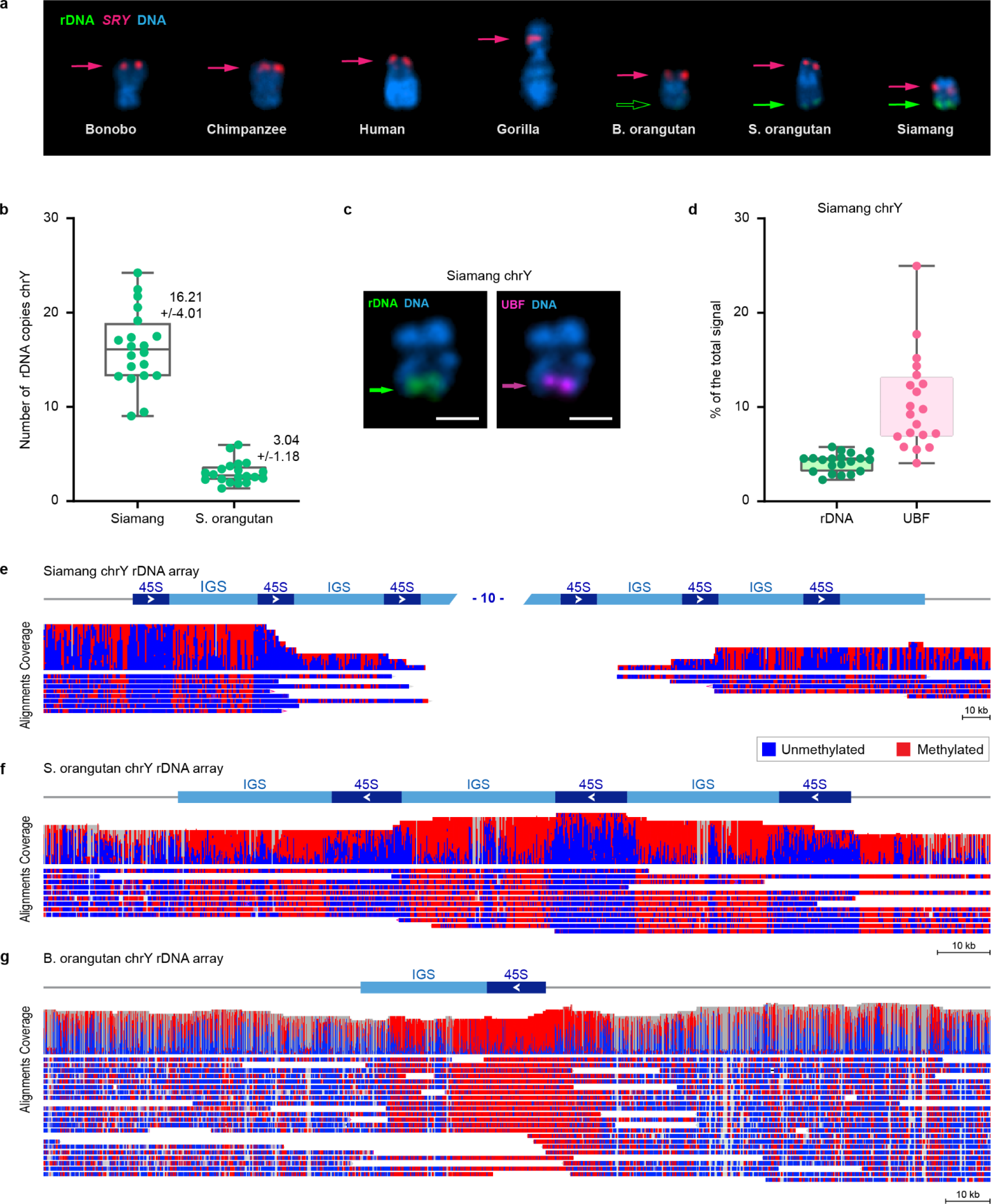
Estimation of rDNA copy number and activity on chromosome Y arrays. (**a**) Gallery view of Y chromosomes from species in this study. Chromosomes were FISH-labeled with rDNA-(BAC RP11-450E20, green) and *SRY*-containing (BAC RP11-400O10, red) BAC probes and counter-stained with DAPI. Siamang and both orangutans’ Y chromosomes have rDNA signal on the distal ends of the q-arms. (**b**) Siamang and Sumatran orangutan chrY rDNA copy number was quantified from the fraction of the total fluorescent intensity of rDNA signals on all chromosomes (from chromosome spreads as in panel **a**) and the Illumina sequencing estimate of the total copy number of rDNA repeats in the genome (339 copies in siamang, 814 in Sumatran orangutan). The mean and standard deviations from 20 chromosome spreads are shown near each box plot. The rounded average of rDNA arrays on chrY were 16 copies for siamang and 3 copies for Sumatran orangutan. (**c**) A representative image of siamang chrY labeled by immuno-FISH with rDNA probe (green) and the antibody against rDNA transcription factor UBF (magenta). The chrY rDNA array is positive for the UBF signal. (**d**) Quantification of siamang chrY rDNA and UBF expressed as the fraction of the total fluorescent intensity of all rDNA-containing chromosomes in a chromosome spread. The mean and standard deviations from 20 chromosome spreads are shown near each box plot. ChrY rDNA arrays contain on average ∼10% of the total chromosomal UBF signal. Siamang **(e)** and Sumatran **(f)** and Bornean **(g)** orangutan read-level plots showing ONT methylation patterns at the chrY rDNA locus and surrounding regions. The coverage track shows the depth of sequencing coverage across the rDNA array, and the methylation track displays the methylation status of individual cytosines. Hypomethylation of the 45S units is evidence of active transcription in siamang and S. orangutan, but not B. orangutan. Only reads >100 kb that are anchored in unique sequence outside the rDNA array and (except for Bornean orangutan) span at least two 45S units are shown.

**Extended Data Figure 6.**
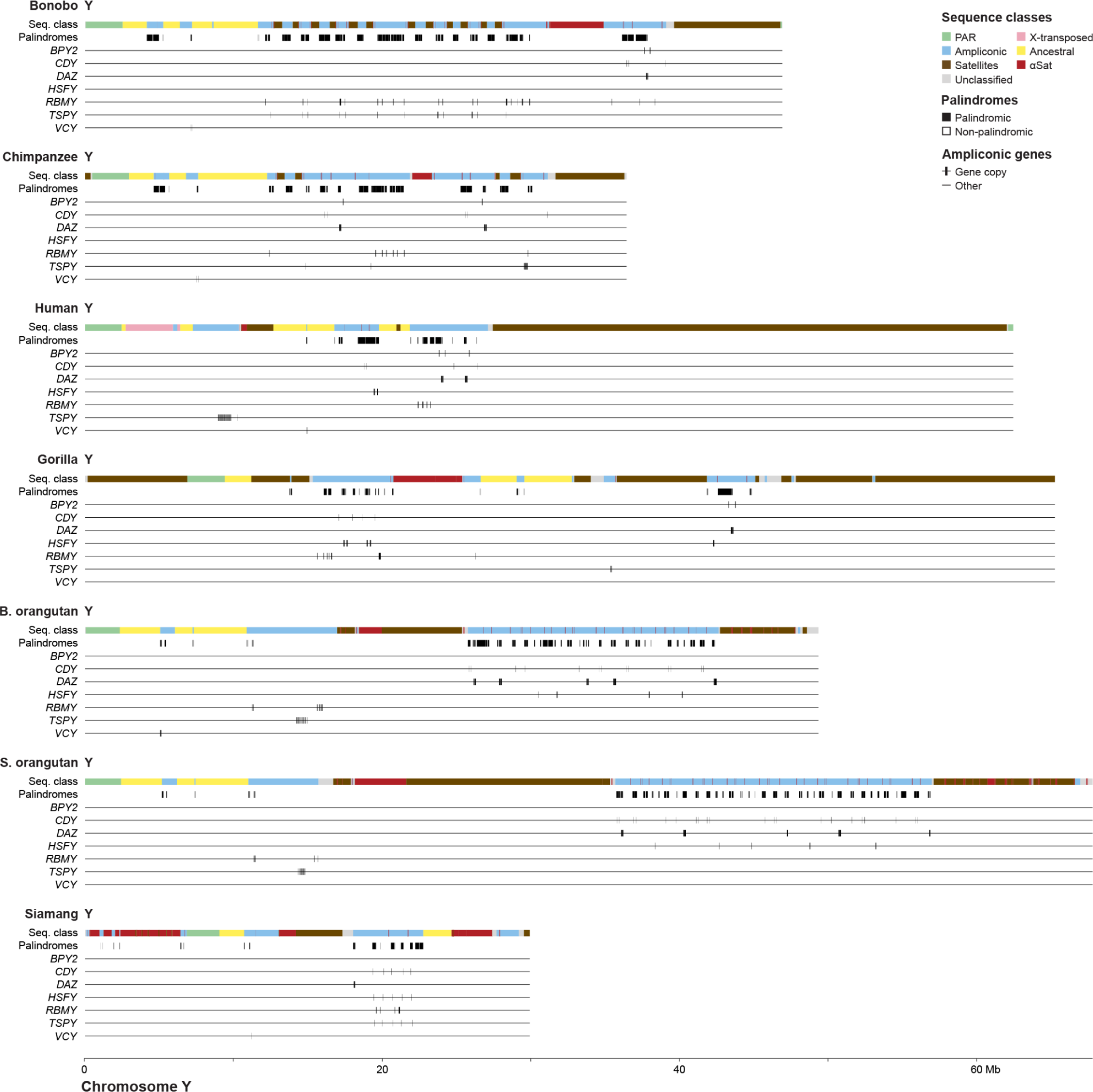
Positions of ampliconic gene families on the Y chromosome. Locations of protein-coding ampliconic genes, grouped by family, are shown with sequence class annotations and palindrome locations on each Y chromosome. The tracks for each species are: (1) sequence class annotation, (2) palindromes, and (3-9) ampliconic gene families: *BPY2*, *CDY*, *DAZ*, *HSFY*, *RBMY*, *TSPY*, and *VCY*. The sequence class track has a discrete class annotation for every base. All other tracks are displayed as presence/absence (color/no color) with the ampliconic gene family tracks containing a horizontal midline to help the eye with the sparse display. All Y chromosomes are plotted on the same scale.

**Extended Data Figure 7.**
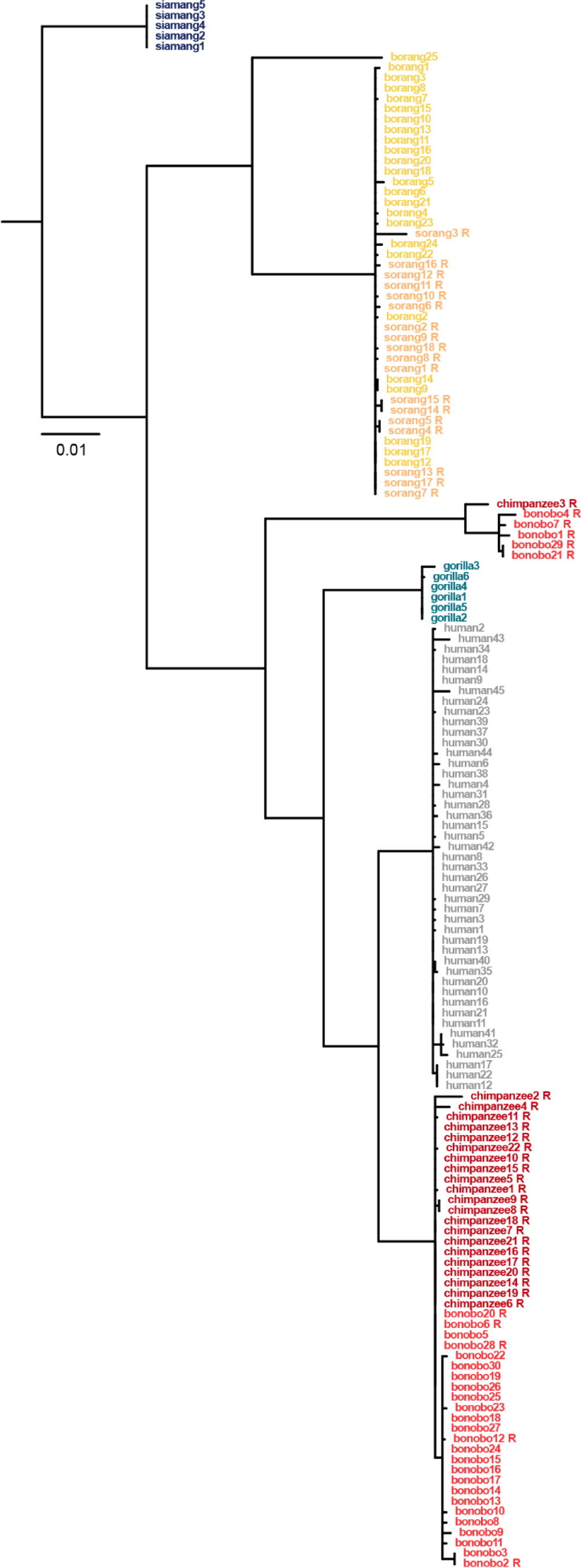
Phylogenetic analysis of the *TSPY* gene family. Phylogenetic analysis (see Methods) of the protein-coding copies of the *TSPY* gene family in great apes, using siamang as an outgroup, uncovered mostly genus-specific clustering suggesting homogenization among copies. Gene copies (numbered for each species) were extracted from the manually curated set (Table S45) and included 5’ and 3’ UTRs, CDS exons, and introns. These sequences were aligned and used to infer a maximum likelihood phylogeny (see Methods for details) with 10,000 ultrafast bootstrap replicates. Nodes with <95% bootstrap support were collapsed. ‘R’ indicated a reverse orientation as compared with assembly sequences.

**Extended Data Figure 8.**
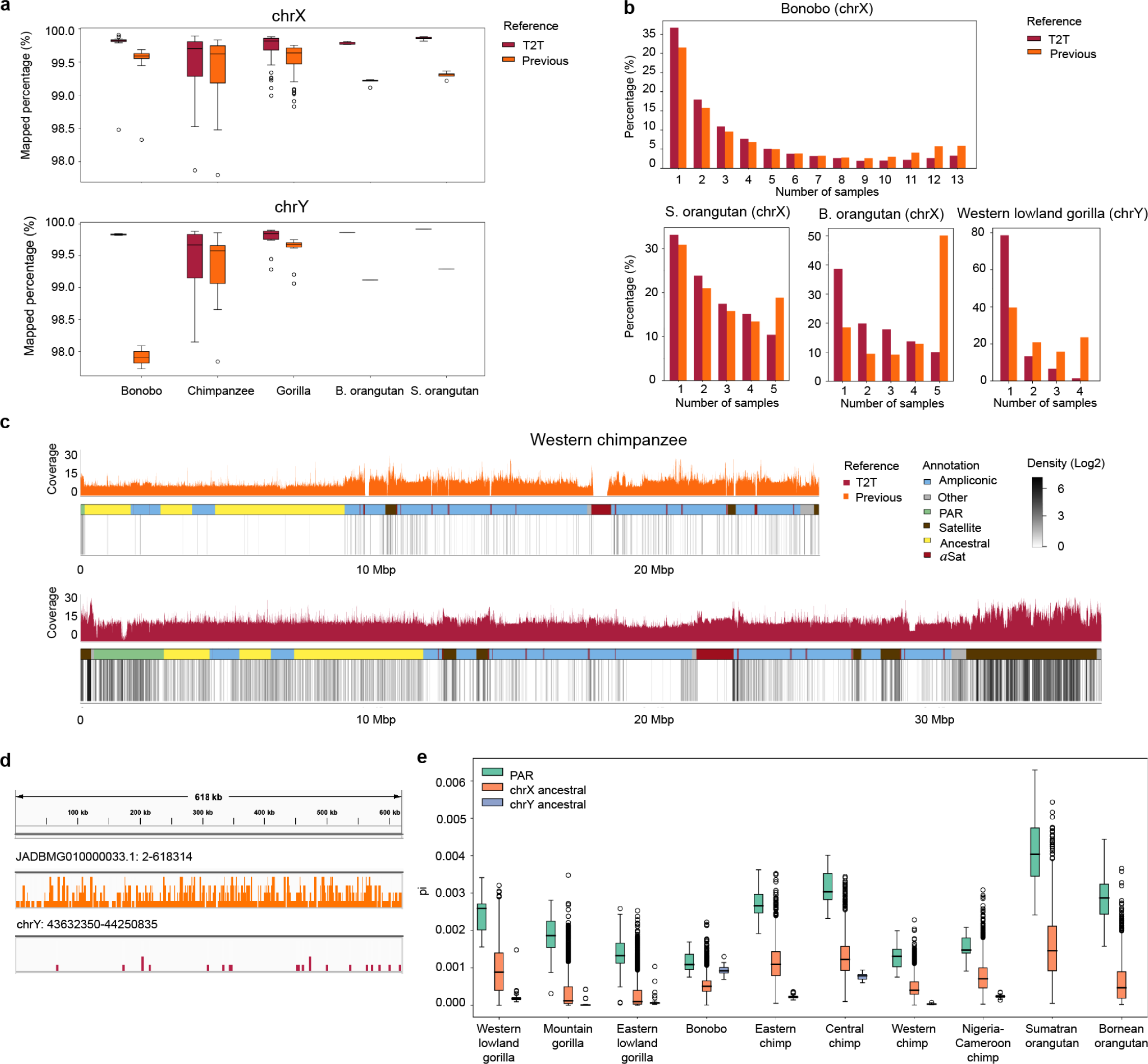
T2T assemblies facilitate short-read mapping and enable the analysis of genetic diversity in great apes. **(a)** The percentage of short reads mapped to T2T vs. previous sex chromosome assemblies (using the previous reference assembly of Sumatran orangutan for Bornean orangutan data). (**b**) Allele frequencies (y-axis) of variants called from reads mapped to T2T vs. previous assemblies. (**c**) Coverage and variant density (in log2 values of densities per 10 kb) distribution across previous (shown in the reverse orientation) and T2T assemblies for western chimpanzee. Peak variant densities were observed at 5.9 for previous chrY and at 7.6 for T2T chrY. (**d**) Distributions of variant allele frequencies on JADBMG010000033.1 (positions 2 to 618,314, upper), a contig from a previous chrY assembly, and T2T chrY (positions 43,632,350 to 44,250,835, bottom), for western lowland gorilla, visualized using IGV. (**e**) Nucleotide diversity (pi)^126^ in pseudoautosomal regions (PARs), ancestral regions of chromosome X, and ancestral regions of chromosome Y. ‘Chimp’ stands for chimpanzee. See Table S42.

