## Supplementary Information for "The Complete Sequence and Comparative Analysis of Ape Sex Chromosomes"

Supplemental Information includes 20 Supplemental Figures (this file), Supplemental Methods (this file), 14 Supplemental Notes (this file), and 47 Supplemental Tables (a separate Excel file).

### Supplemental Figures

#### Figure S1. Karyotypes of cell lines used in the study

(**A**) bonobo, (**B**) chimpanzee, (**C**) gorilla, (**D**) B. orangutan, (**E**) S. orangutan, (**F**) siamang. For great apes, the homologous human chromosomes are labeled by Roman numerals, with chromosome ten underlined. Great apes karyotypes have been assembled following ISCN[1] and the Atlas of Mammalian Chromosomes[2] as references.

**A**

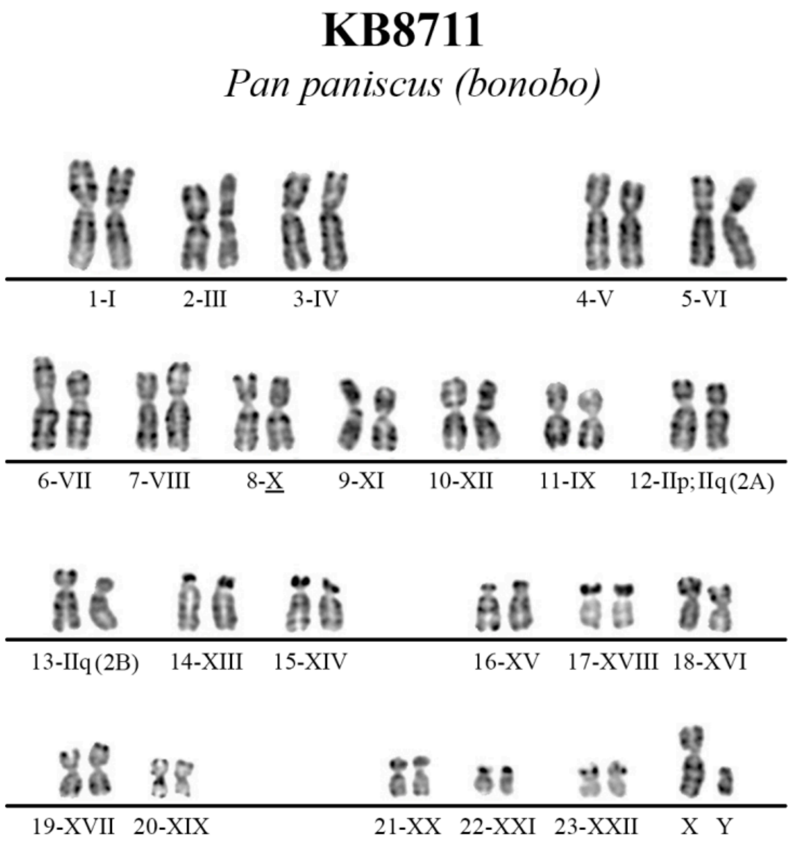

B

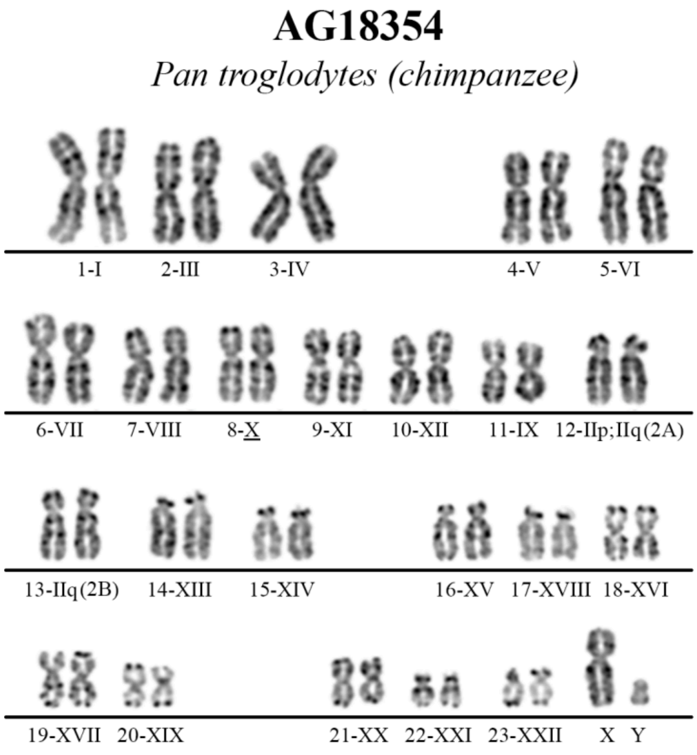
AG18354
Pan troglodytes (chimpanzee)
1-I
2-III
3-IV
4-V
5-VI
6-VII
7-VIII
8-X
9-XI
10-XII
11-IX
12-IIp;IIq(2A)
13-IIq(2B)
14-XIII
15-XIV
16-XV
17-XVIII
18-XVI
19-XVII
20-XIX
21-XX
22-XXI
23-XXII
X
Y

C

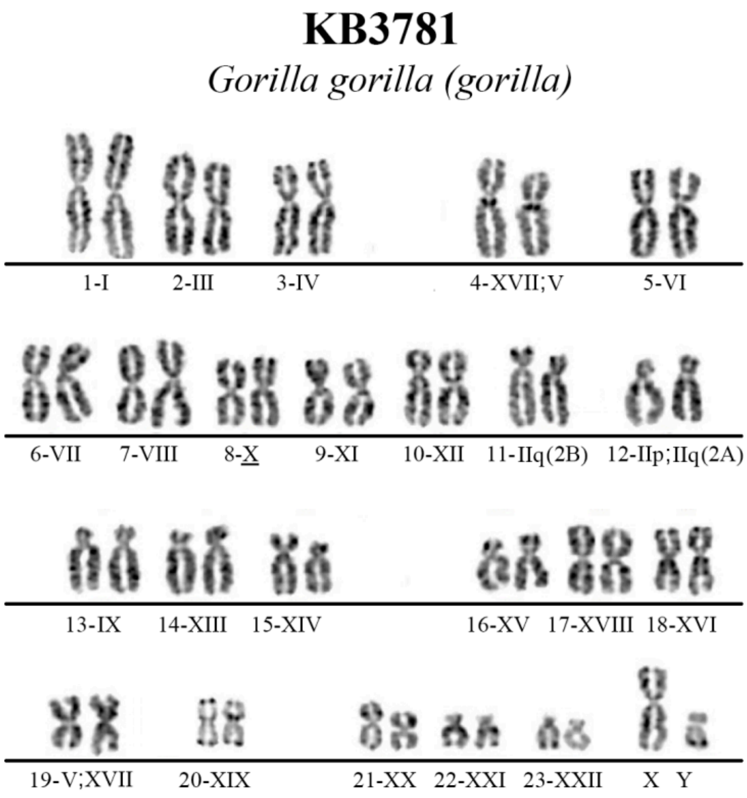
KB3781
Gorilla gorilla (gorilla)
1-I
2-III
3-IV
4-XVII;V
5-VI
6-VII
7-VIII
8-X
9-XI
10-XII
11-IIq(2B)
12-IIp;IIq(2A)
13-IX
14-XIII
15-XIV
16-XV
17-XVIII
18-XVI
19-V;XVII
20-XIX
21-XX
22-XXI
23-XXII
X
Y

D

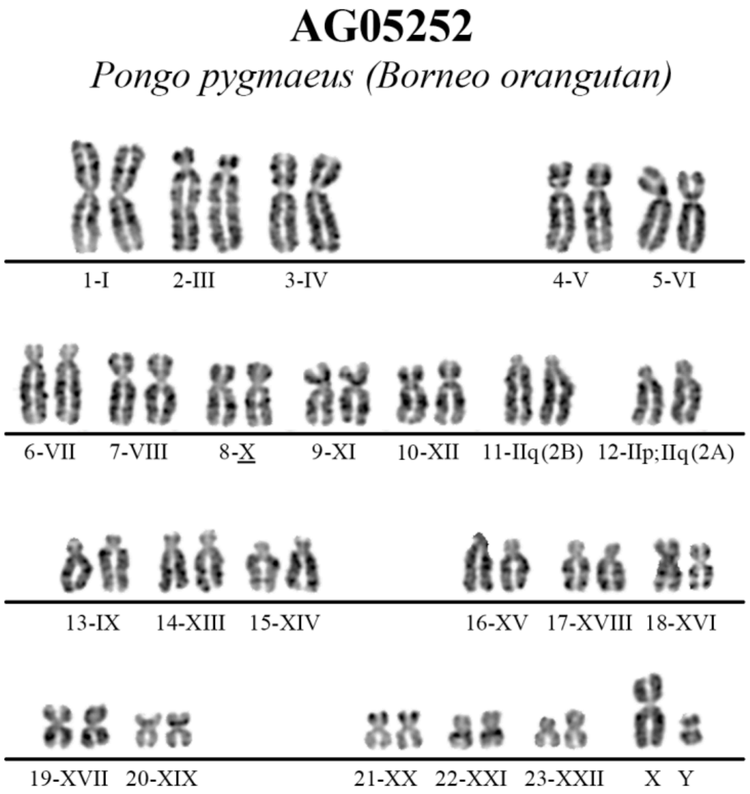
AG05252
Pongo pygmaeus (Borneo orangutan)
1-I 2-III 3-IV 4-V 5-VI
6-VII 7-VIII 8-X 9-XI 10-XII 11-IIq(2B) 12-IIp;IIq(2A)
13-IX 14-XIII 15-XIV 16-XV 17-XVIII 18-XVI
19-XVII 20-XIX 21-XX 22-XXI 23-XXII X Y

E

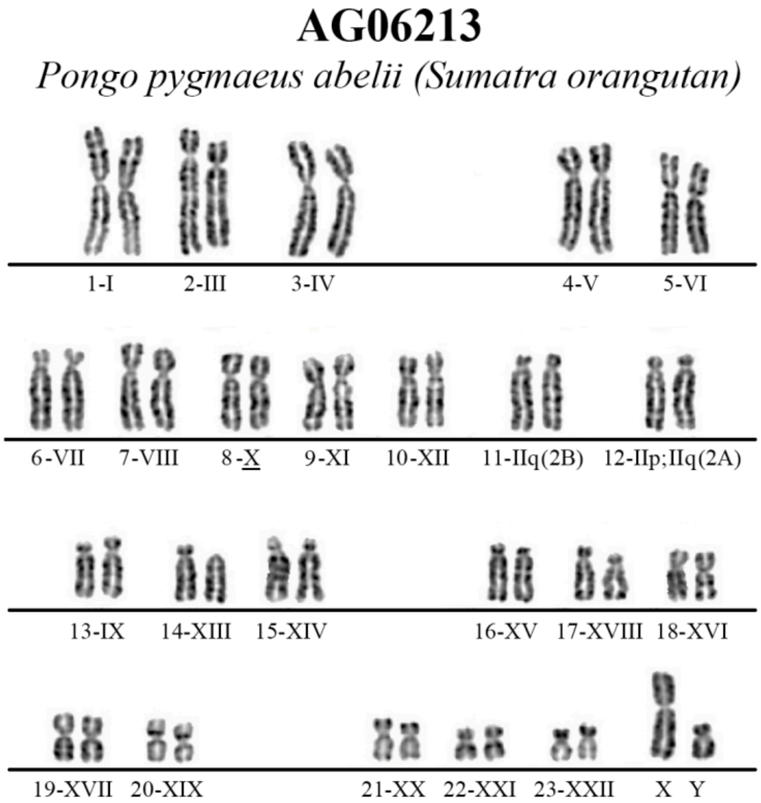
AG06213
Pongo pygmaeus abelii (Sumatra orangutan)
1-I 2-III 3-IV 4-V 5-VI
6-VII 7-VIII 8-X 9-XI 10-XII 11-IIq(2B) 12-IIp;IIq(2A)
13-IX 14-XIII 15-XIV 16-XV 17-XVIII 18-XVI
19-XVII 20-XIX 21-XX 22-XXI 23-XXII X Y

**F** Jambi (*Symphalangus syndactylus*)

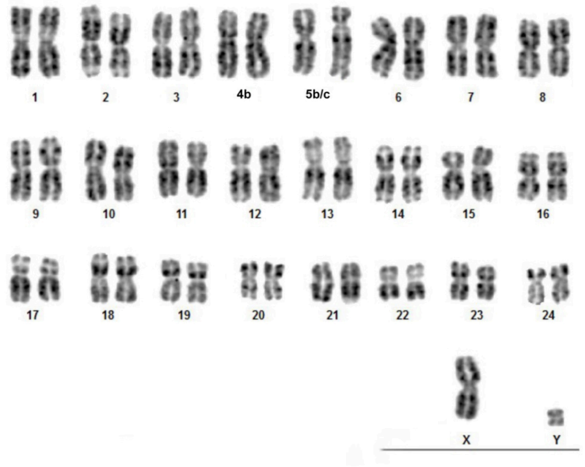

### Figure S2. Assembly graphs prior to manual curation

The HiFi-based graphs from Verkko[3] are in homopolymer-compressed space and have gone through automated resolution using ONT reads. The manual resolution to get T2T X and Y chromosomes came after this step. The visualizations were generated in Bandage[4]. (**A**) The entire assembly (i.e. all chromosomes) with chromosomes X and Y colored respectively red and blue, with all other sequences in gray. (**B**) Subset of the graphs from panel A showing only chromosomes X and Y, with red (X) and blue (Y) colors being assigned by trio (bonobo and gorilla) or Hi-C (chimpanzee, orangutans, and siamang) phasing. Gray nodes are still associated with X or Y but were not assigned a haplotype by the phasing algorithm, though the assignment was inferred from the graph structure during manual curation.

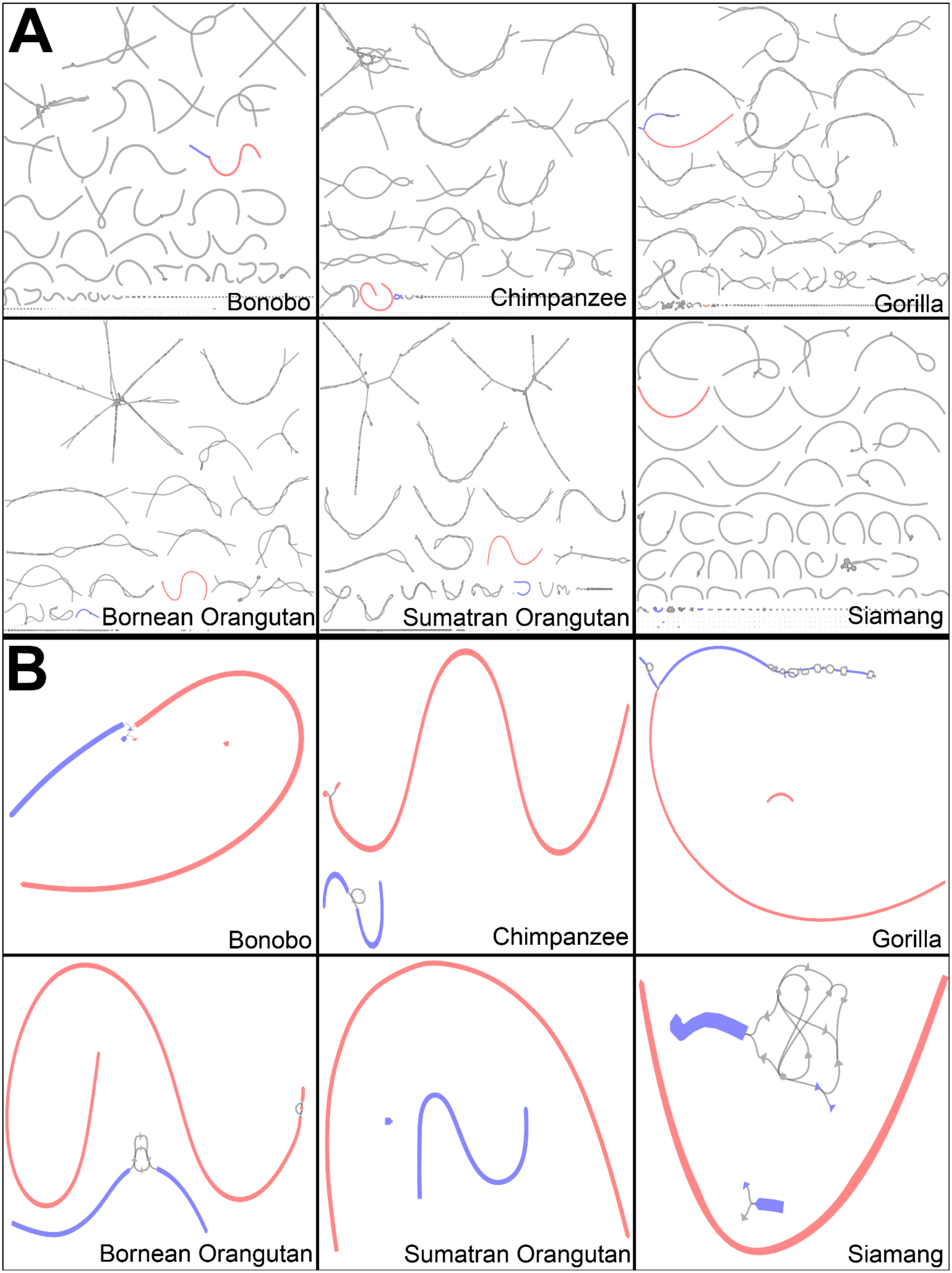
A
Bonobo
Chimpanzee
Gorilla
Bornean Orangutan
Sumatran Orangutan
Siamang
B
Bonobo
Chimpanzee
Gorilla
Bornean Orangutan
Sumatran Orangutan
Siamang

### Figure S3. Ape Y chromosome structural variants

Human Y chromosome (**A**) pseudoautosomal region (PAR) and (**B**) non-PAR aligned to bonobo, chimpanzee, gorilla, B. orangutan, S. orangutan, and gibbon Y chromosomes. The aligned regions are represented by green bars while the parts that are not aligned are indicated by orange bars. The number of bases affected by different classes of variants in (**C**) PAR and (**D**) non-PAR of human chromosome Y, in comparisons with nonhuman apes.

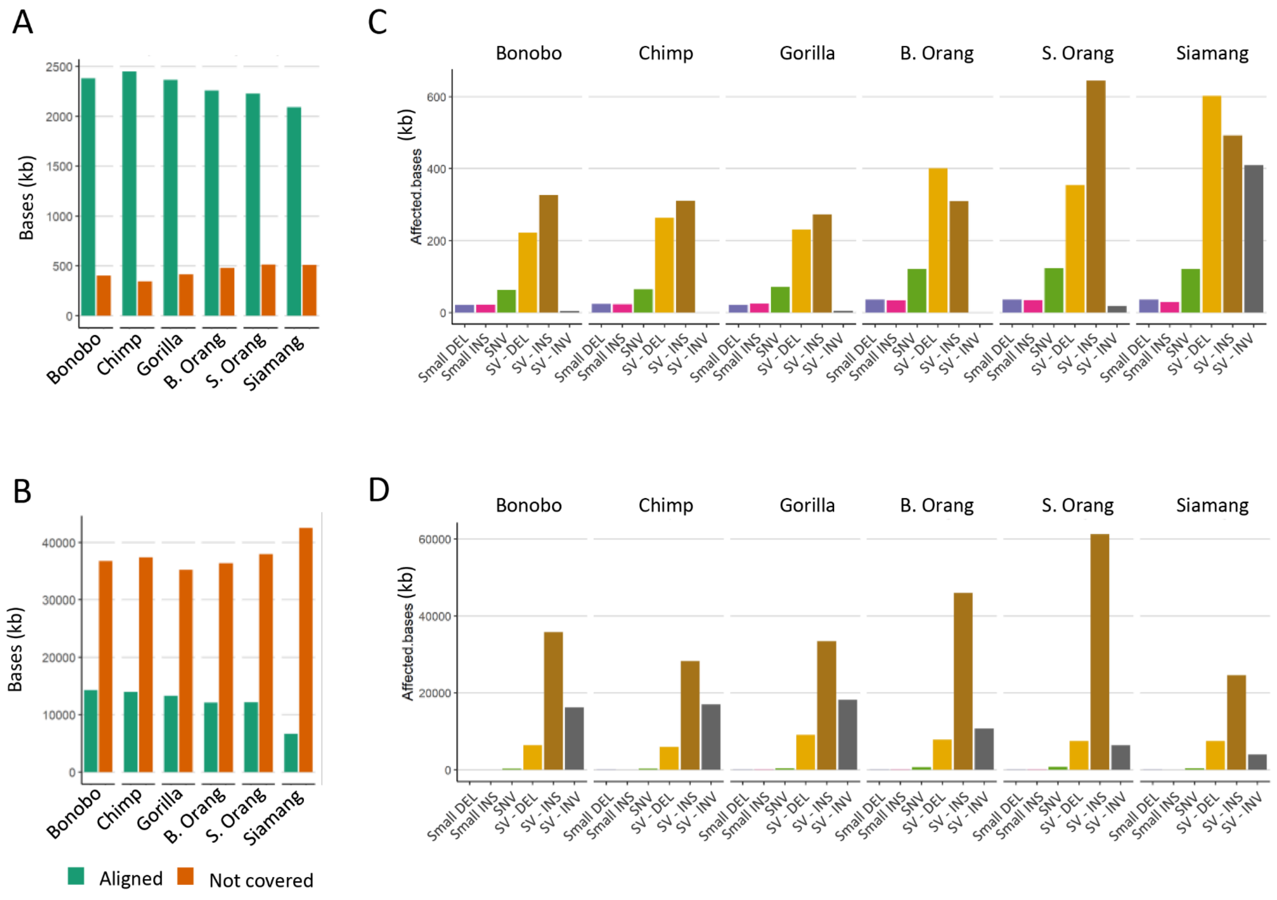

### Figure S4. Ape X chromosome structural variants

Human X chromosome (**A**) pseudoautosomal region (PAR) and (**B**) non-PAR aligned to bonobo, chimpanzee, gorilla, B. orangutan, S. orangutan, and gibbon X chromosomes. The aligned regions are represented by green bars while the parts that are not aligned are indicated by orange bars. The number of bases affected by different classes of variants in (**C**) non-PAR and (**D**) non-PAR of human chromosome X, in comparisons with nonhuman apes.

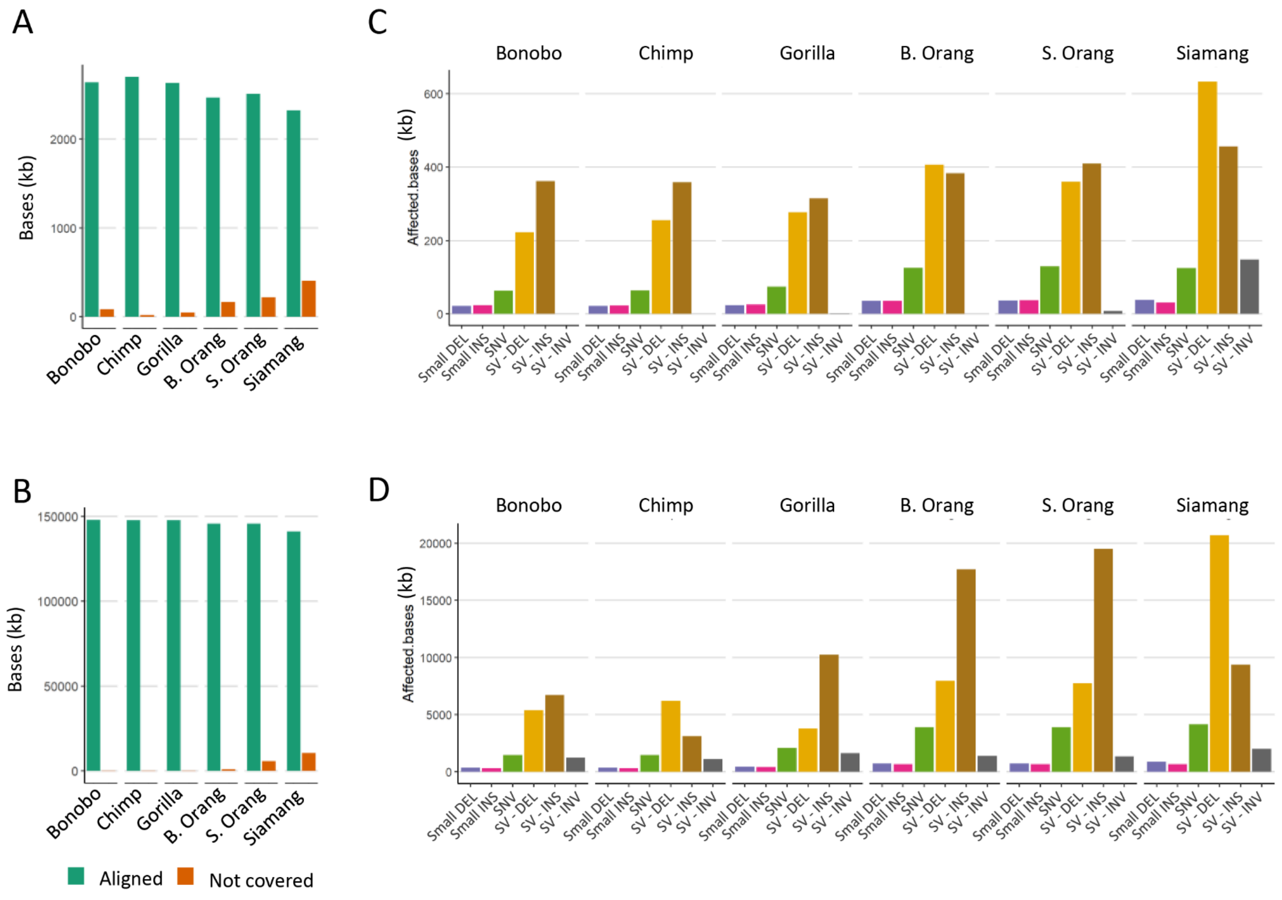

### Figure S5. Correlation of the number of structural variants (SVs) with ape phylogeny

(**A**) The number of putative structural variants originating from different phylogenetic branches and, in the parentheses, the number of structural variants overlapping with exons of human and non-human primate genes. The variants along the ancestral branches including the reference species, human, were not computed (see Supplemental Methods for details). (**B**) Correlation between divergence time in MY vs. sum of the number of structural variants/Mb. The blue points and dotted line represent the Y chromosomes (slope of 15.8 SVs/Mb/MY) and the orange points and the orange dotted line represent data points for the X chromosomes (slope of 6.1 SVs/Mb/MY). (**C**) Correlation between divergence time in MY vs. sum of the number of structural variants in PARs. The blue points and dotted line represent the Y chromosomes and the orange points and line represent data for the X chromosomes.

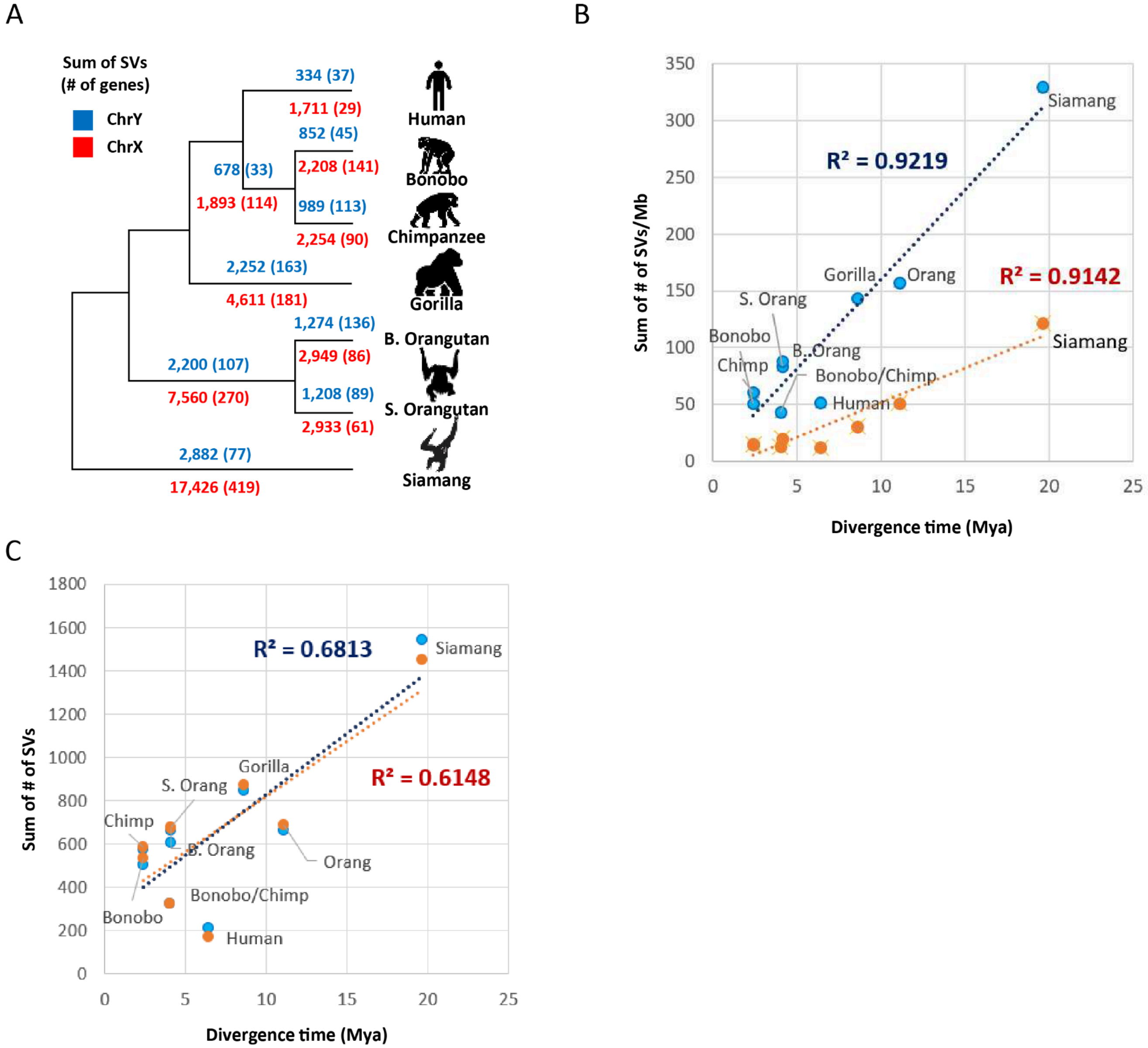

### Figure S6. The organization of long ampliconic regions on Bornean and Sumatran orangutan Y chromosomes

(**A**) Dotplot analysis of the studied ampliconic regions against themselves in each orangutan species separately. (**B**) The ampliconic regions were decomposed with the PanGenome Research ToolKit (PGR-TK based on graph decomposition[5]).

**A**

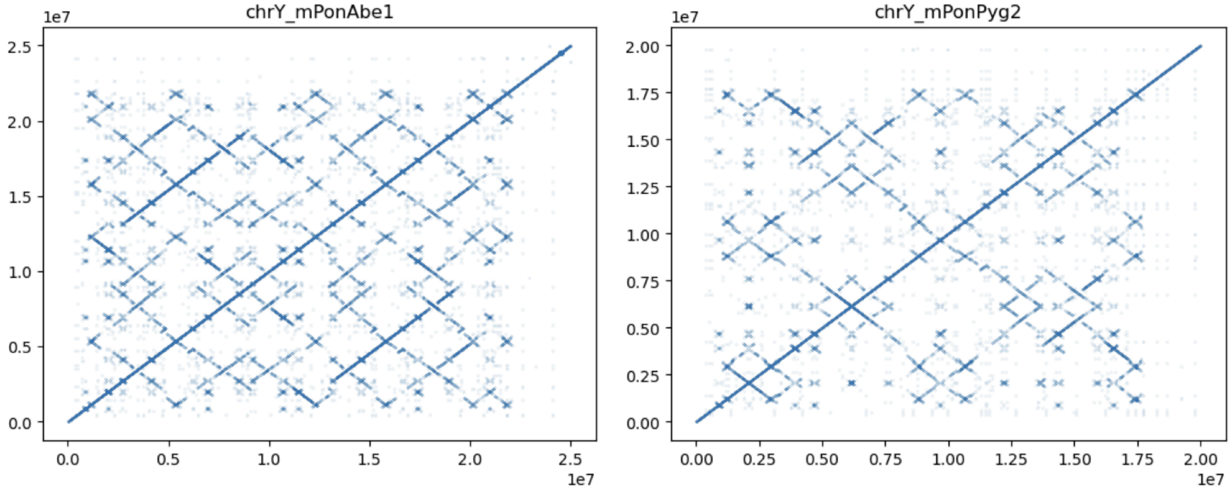

**B**

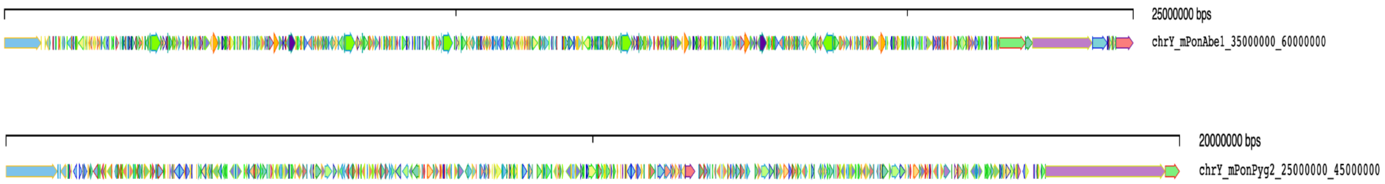

### Figure S7. Palindrome analyses

We analyzed GC content of palindrome arms and a spacer, their respective lengths, and the identity of corresponding arms in each palindrome (using stretcher from the EMBOSS Software Suite[6]). We found shorter spacers and increased arm lengths associated with the higher sequence identity between the arms (Table S15). By comparing GC content between arms and spacers jointly for all species, we found a median difference of 0.01, or 1%, for both X and Y chromosomes ($p$=0.0308 and 0.01037, respectively, for two sample one-sided $t$-test). Increased GC content in arms compared to their corresponding spacers is consistent with gene conversion acting on these palindromes in sex chromosomes. Compared with those on the Y, palindromes on the X have higher overall GC content (both in arms and in spacers, $p=2.2\times10^{-16}$, two-sided two-sample $t$-test, Fig. S7CD), providing more donor sites for GC-biased gene conversion. Additonally, we found a negative correlation between arms' GC content and spacer length, consistent with GC-biased gene conversion, on the X, but not the Y, although this effect was weak and not present in all the species (Fig. S7KL, Table S15). Except for siamang, shorter spacers were associated with higher sequence identity between the arms (Fig. S7G, Fig. S7H; significant Pearson's correlations in most cases, Table S15), consistent with more efficient gene conversion for sequences located closer to each other[7]. An increase in arms' length was associated with increased sequence identity between them (Fig. S7D, Fig. S7F; significant Pearson's correlations in 50% of cases tested, Table S15), suggesting that longer palindromes undergo gene conversion more frequently, and consistent with long gene conversion tracts in palindromes[8].

Palindromes with spacers of size 0 were excluded from the analysis (n=3). For the calculation of the GC content of the arms and length, we used program geecee from the EMBOSS Software Suite and only the values from the first arm were used (capitalizing on the high sequence identity). Please note that human was not included in the analysis, but has been studied previously[9–12]. The points in the scatterplots might be overlapping, and such cases will be represented by darker shade of colors.

(**A**) Comparison of palindrome arm lengths between X and Y chromosomes. (**B**) Difference between the GC content of arms versus their corresponding spacers for X and Y chromosomes. (**C**) GC content of palindrome spacers for X and Y chromosomes. (**D**) GC content of palindrome arms for X and Y chromosomes. (**E**) The scatterplot of GC content of arms and GC content of their corresponding spacers for the chromosome X. (**F**) The scatterplot of GC content of arms and GC content of their corresponding spacers for the chromosome Y. (**G**) The scatterplot of the spacer length and sequence identity of the palindrome arms for the chromosome X. (**H**) The scatterplot of the spacer length and sequence identity of the palindrome arms for the chromosome Y. (**I**) The scatterplot of the arm length and sequence identity of the palindrome arms for the chromosome X. (**J**) The scatterplot of the arm length and sequence identity of the palindrome arms for the chromosome Y. (**K**) The scatterplot of the spacer length and GC content of the palindrome arms for the chromosome X. (**L**) The scatterplot of the spacer length and GC content of the palindrome arms for the chromosome Y. (**M**) The scatterplot of the arm length and GC content of the palindrome arms for the chromosome X. (**N**) The scatterplot of the arm length and GC content of the palindrome arms for the chromosome Y.

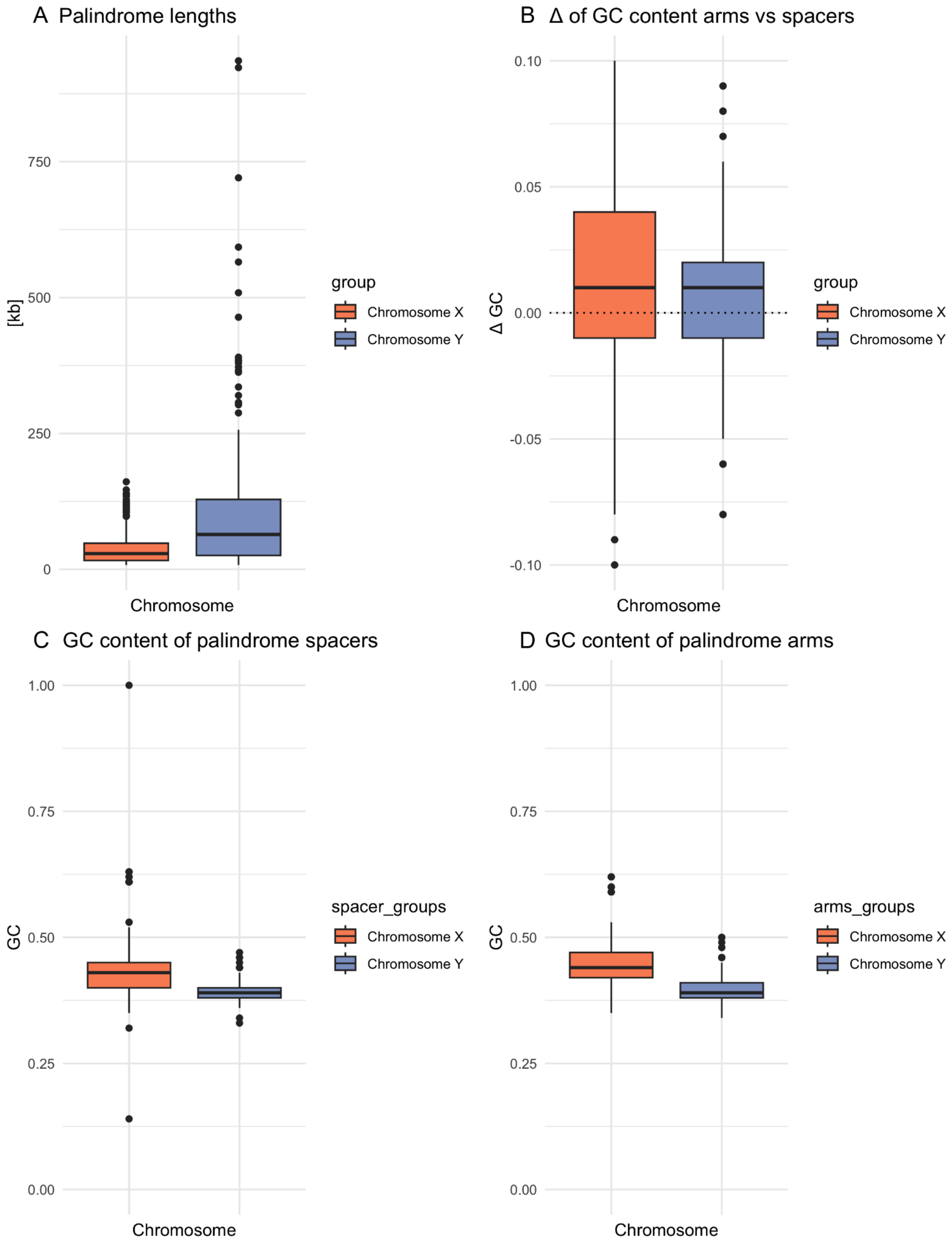
A Palindrome lengths
[kb]
750
500
250
0
Chromosome
group
Chromosome X
Chromosome Y
B Δ of GC content arms vs spacers
Δ GC
0.10
0.05
0.00
-0.05
-0.10
Chromosome
group
Chromosome X
Chromosome Y
C GC content of palindrome spacers
GC
1.00
0.75
0.50
0.25
0.00
Chromosome
spacer_groups
Chromosome X
Chromosome Y
D GC content of palindrome arms
GC
1.00
0.75
0.50
0.25
0.00
Chromosome
arms_groups
Chromosome X
Chromosome Y

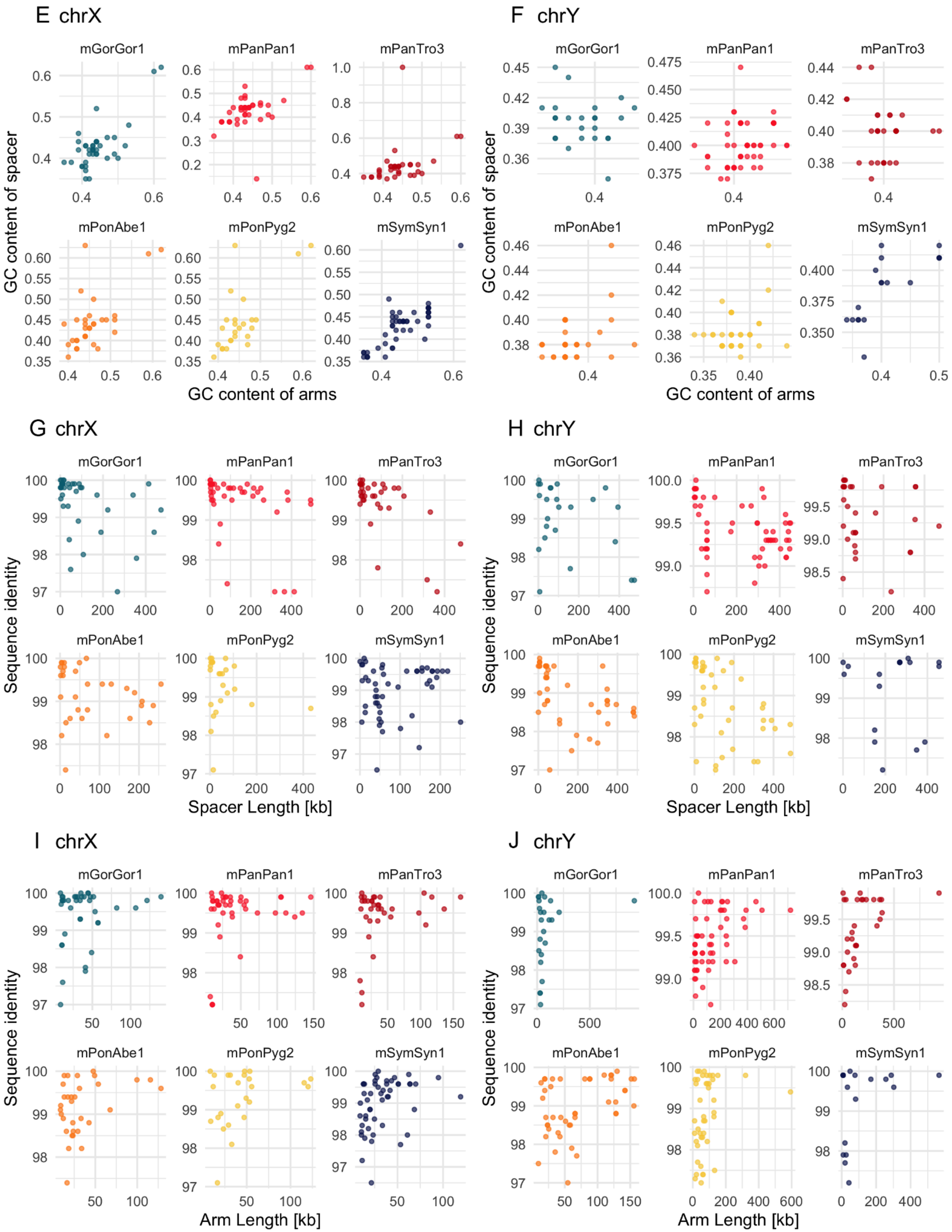
E chrX
F chrY
mGorGor1
mPanPan1
mPanTro3
mPonAbe1
mPonPyg2
mSymSyn1
GC content of spacer
GC content of arms
G chrX
H chrY
Sequence identity
Spacer Length [kb]
I chrX
J chrY
Arm Length [kb]

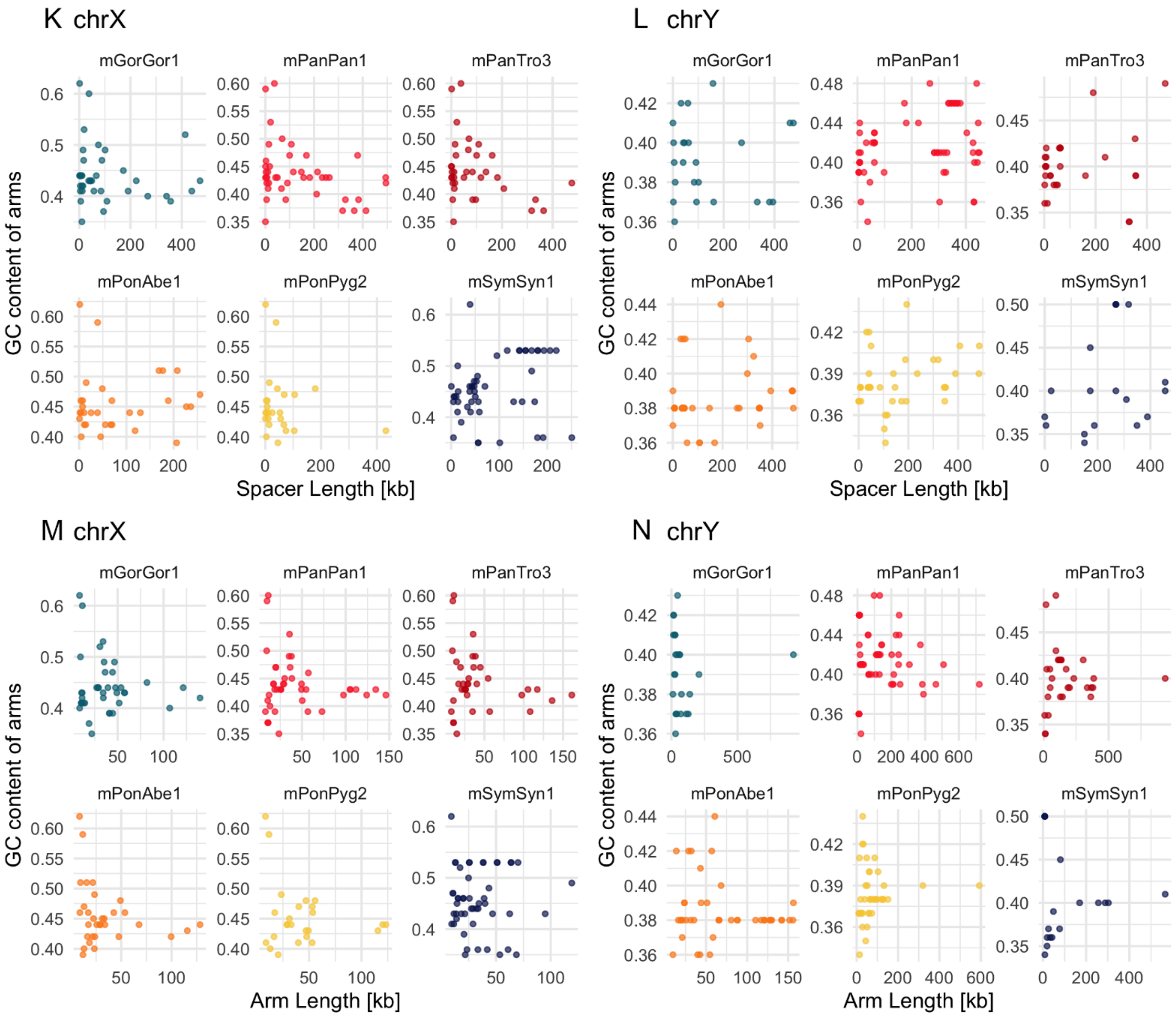
K chrX
L chrY
M chrX
N chrY
mGorGor1
mPanPan1
mPanTro3
mPonAbe1
mPonPyg2
mSymSyn1
GC content of arms
Spacer Length [kb]
Arm Length [kb]

### Figure S8. Repeat content by sequence class on the X and the Y chromosomes

Proportions of repeats and non-repetitive DNA are shown across ampliconic regions **(A)**, PARs **(B)**, ancestral (X-ancestral or Y X-degenerate regions) **(C)**, and satellite regions **(D)**. Repeat classes correspond to the colors depicted in the key below. Data depicted refers to Table S20.

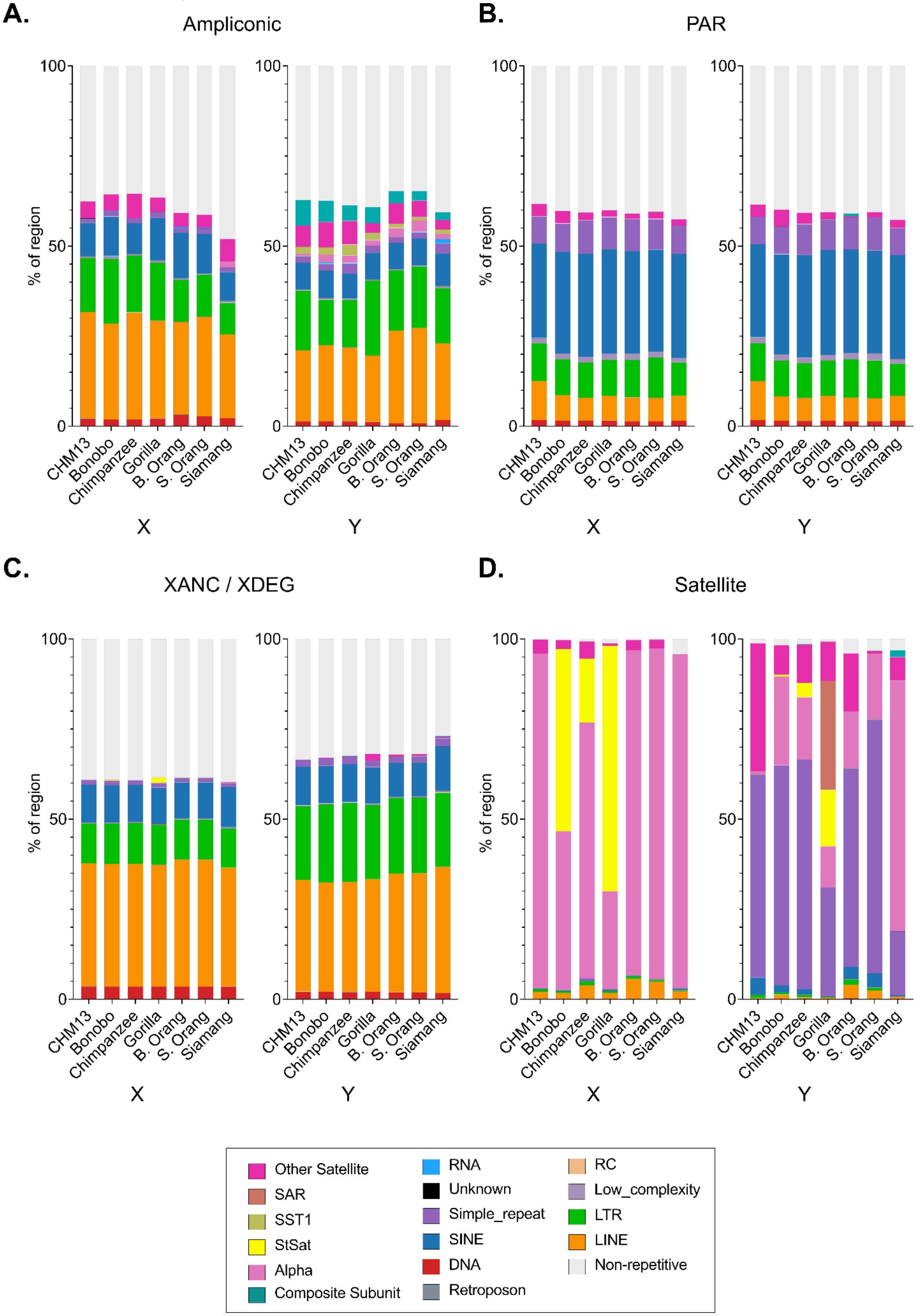

### Figure S9. Locations of new composite repeats across the ape chromosomes

Ideogram of X and Y chromosomes indicating locations of annotated composite repeats as singletons (black), duplicates (purple), and arrays (pink); corresponding to Table S22. Red indicates centromeres.

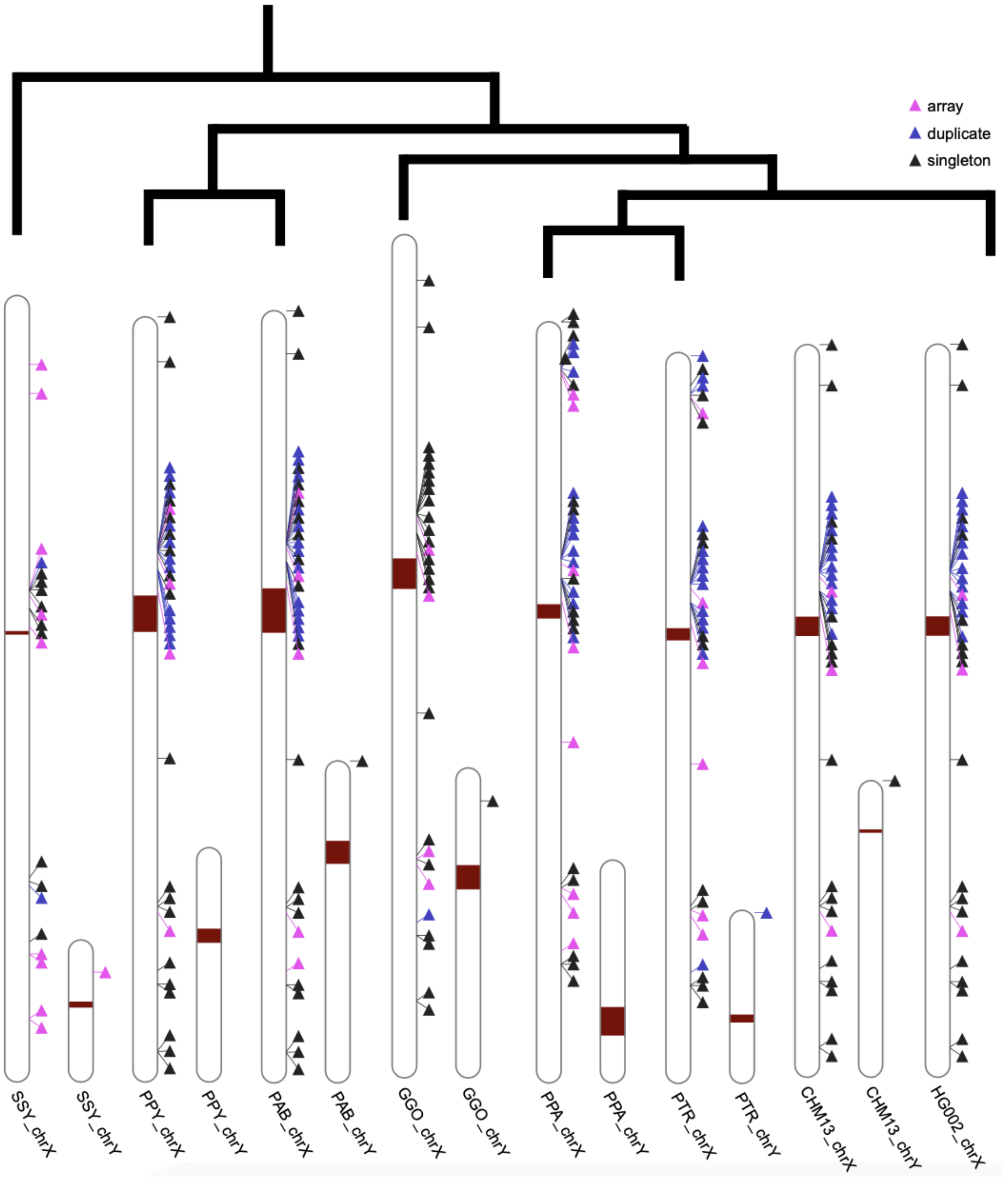

#### Figure S10. New satellites

The number of new satellite monomers on each ape X and Y chromosome are shown via categorical heat map for each of the 25 repeat models able to be surveyed across all species (Table S23). Copy numbers of the repeats listed are grouped by monomer copy number; satellites undetected in an ape genome are denoted as black boxes. Both the HG002 (diploid) and CHM13 (haploid) human samples are included. Of the 25 satellite models surveyed across all apes, only 15 were identified in all apes (collapsing variants of DXZ4 into one satellite), suggesting lineage-specific evolution of many of the newly discovered satellites.

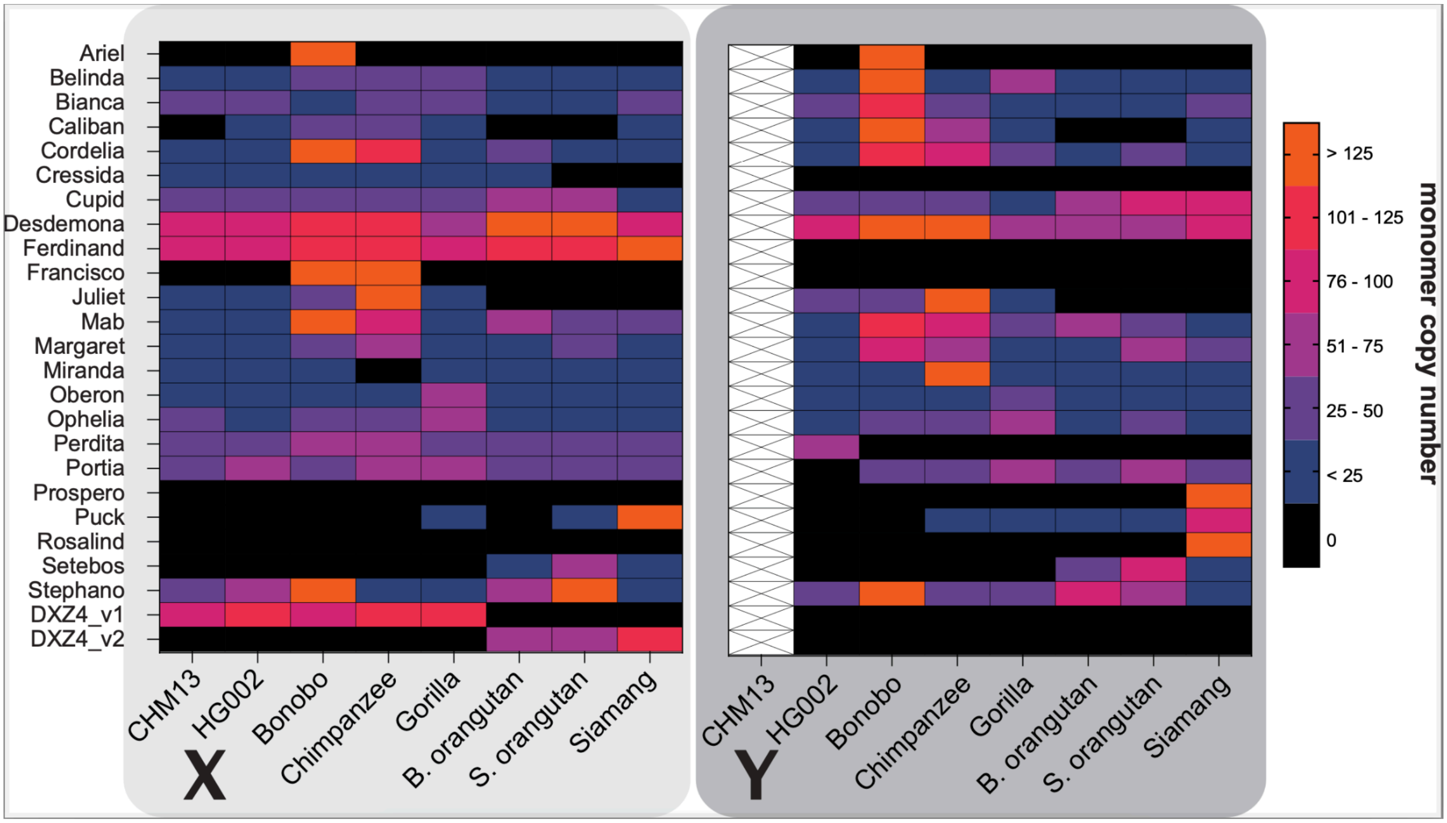

### Figure S11. Lineage-specific repeats on ape sex chromosomes

The number of bases comprising lineage-specific repeat expansions across each of the ape sex chromosomes.

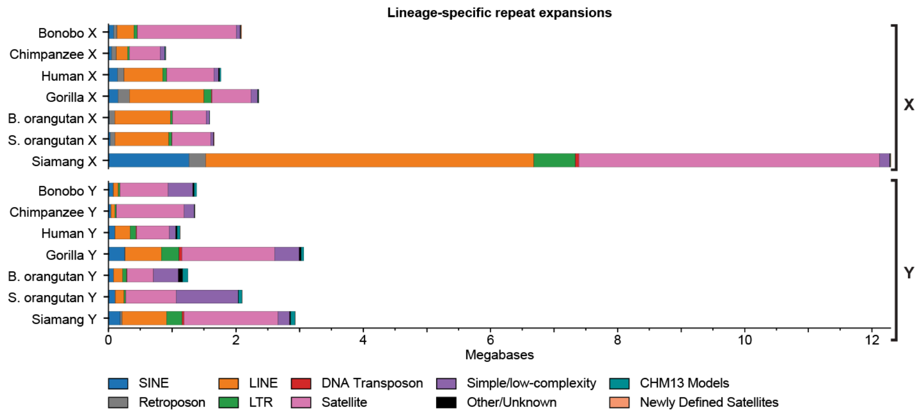

### Figure S12. Non-B DNA annotations

To color the barplot, for each species and each chromosome, we used the aligned intervals files that specify which section of the T2T assembly maps to a previous assembly (shown in white). Immediately below each barplot, for each species, each chromosome, and each non-B DNA type, we used non-B DNA density to color the barplot. For each 100-kp window, the absence of non-B DNA is shown in white, the highest density of non-B DNA is shown in black, and other non-B DNA density is shown in various shades of gray. The density was normalized separately for each chromosome. (**A**) A-phased repeats (APR), (**B**) direct repeats (DR), (**C**) G-quadruplexes (GQ), (**D**) mirror repeats (MR), (**E**) short tandem repeats (STR), (**F**) Z-DNA (Z).

**A**

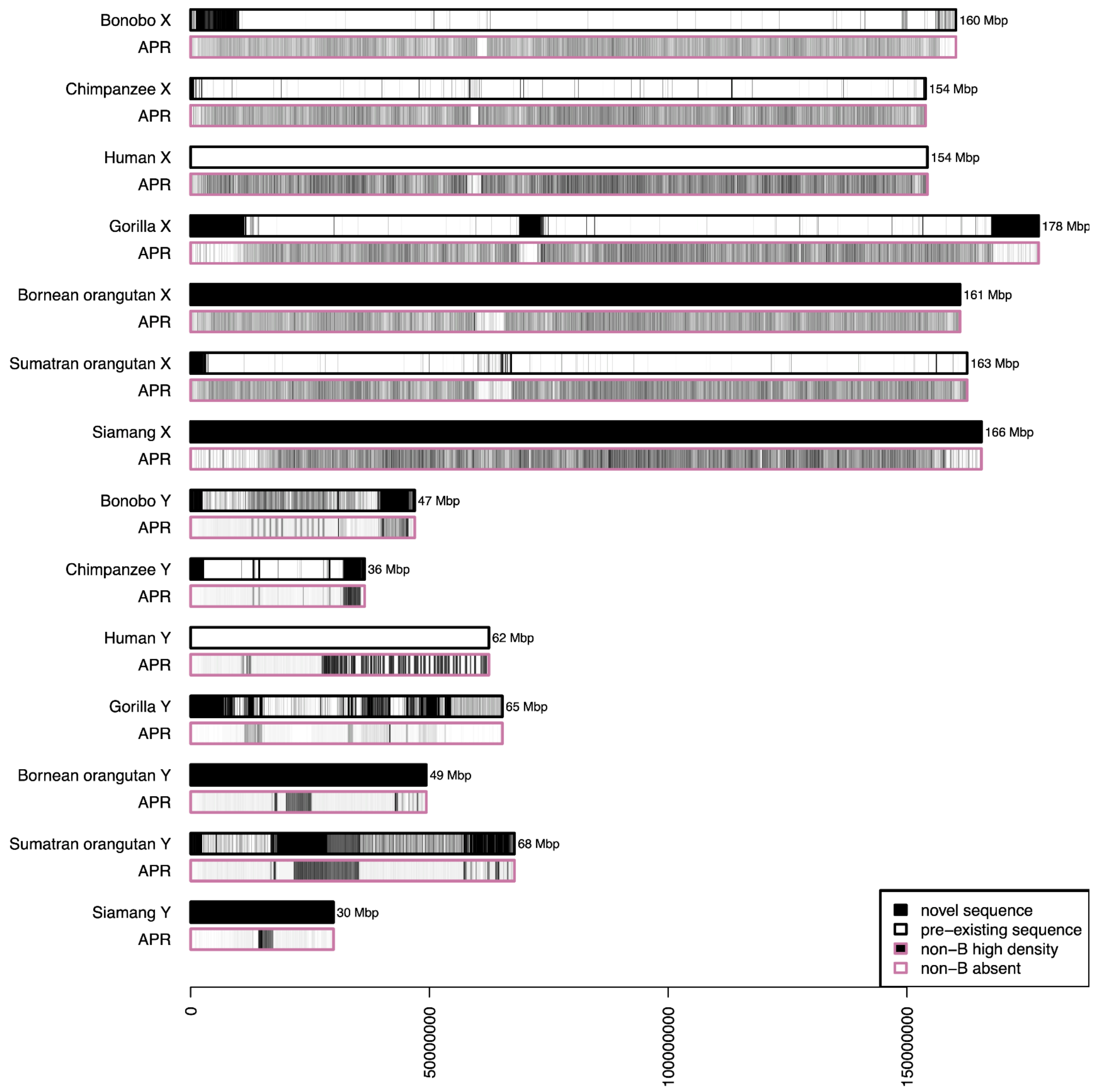

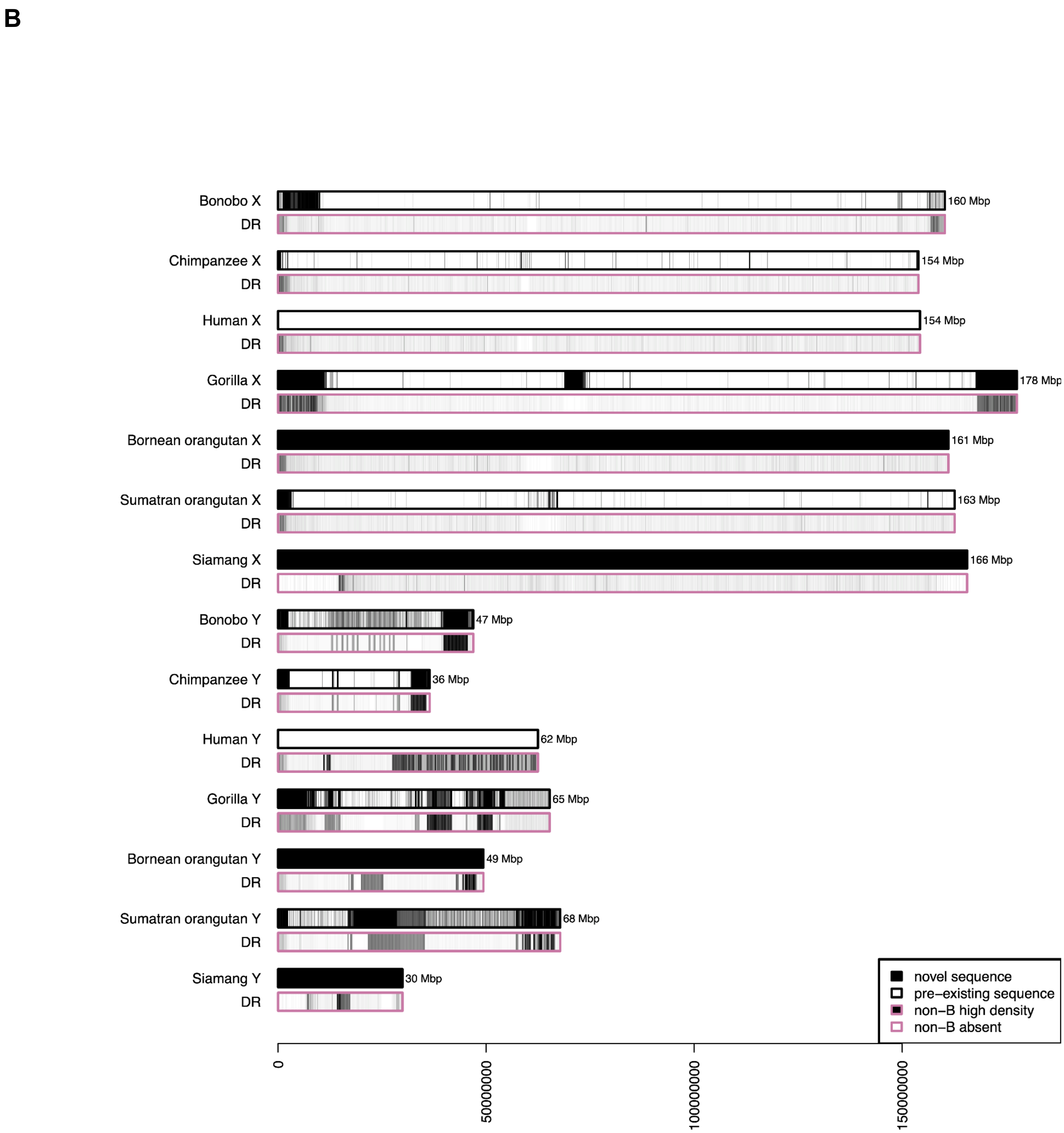
B
Bonobo X
DR
160 Mbp
Chimpanzee X
DR
154 Mbp
Human X
DR
154 Mbp
Gorilla X
DR
178 Mbp
Bornean orangutan X
DR
161 Mbp
Sumatran orangutan X
DR
163 Mbp
Siamang X
DR
166 Mbp
Bonobo Y
DR
47 Mbp
Chimpanzee Y
DR
36 Mbp
Human Y
DR
62 Mbp
Gorilla Y
DR
65 Mbp
Bornean orangutan Y
DR
49 Mbp
Sumatran orangutan Y
DR
68 Mbp
Siamang Y
DR
30 Mbp
novel sequence
pre–existing sequence
non–B high density
non–B absent
0
50000000
100000000
150000000

C

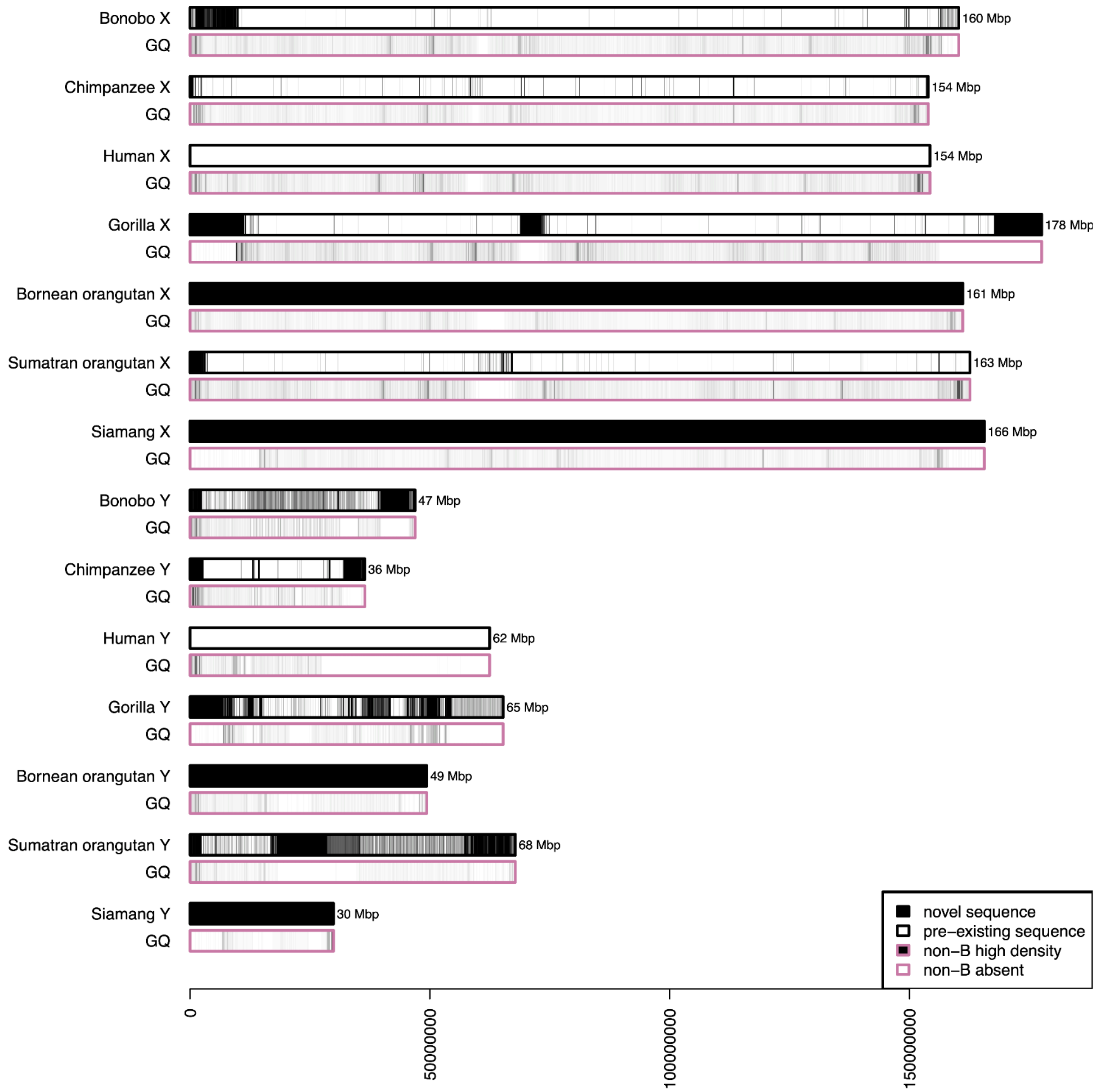

Bonobo X
GQ
160 Mbp
Chimpanzee X
GQ
154 Mbp
Human X
GQ
154 Mbp
Gorilla X
GQ
178 Mbp
Bornean orangutan X
GQ
161 Mbp
Sumatran orangutan X
GQ
163 Mbp
Siamang X
GQ
166 Mbp
Bonobo Y
GQ
47 Mbp
Chimpanzee Y
GQ
36 Mbp
Human Y
GQ
62 Mbp
Gorilla Y
GQ
65 Mbp
Bornean orangutan Y
GQ
49 Mbp
Sumatran orangutan Y
GQ
68 Mbp
Siamang Y
GQ
30 Mbp
novel sequence
pre–existing sequence
non–B high density
non–B absent
0
50000000
100000000
150000000

D

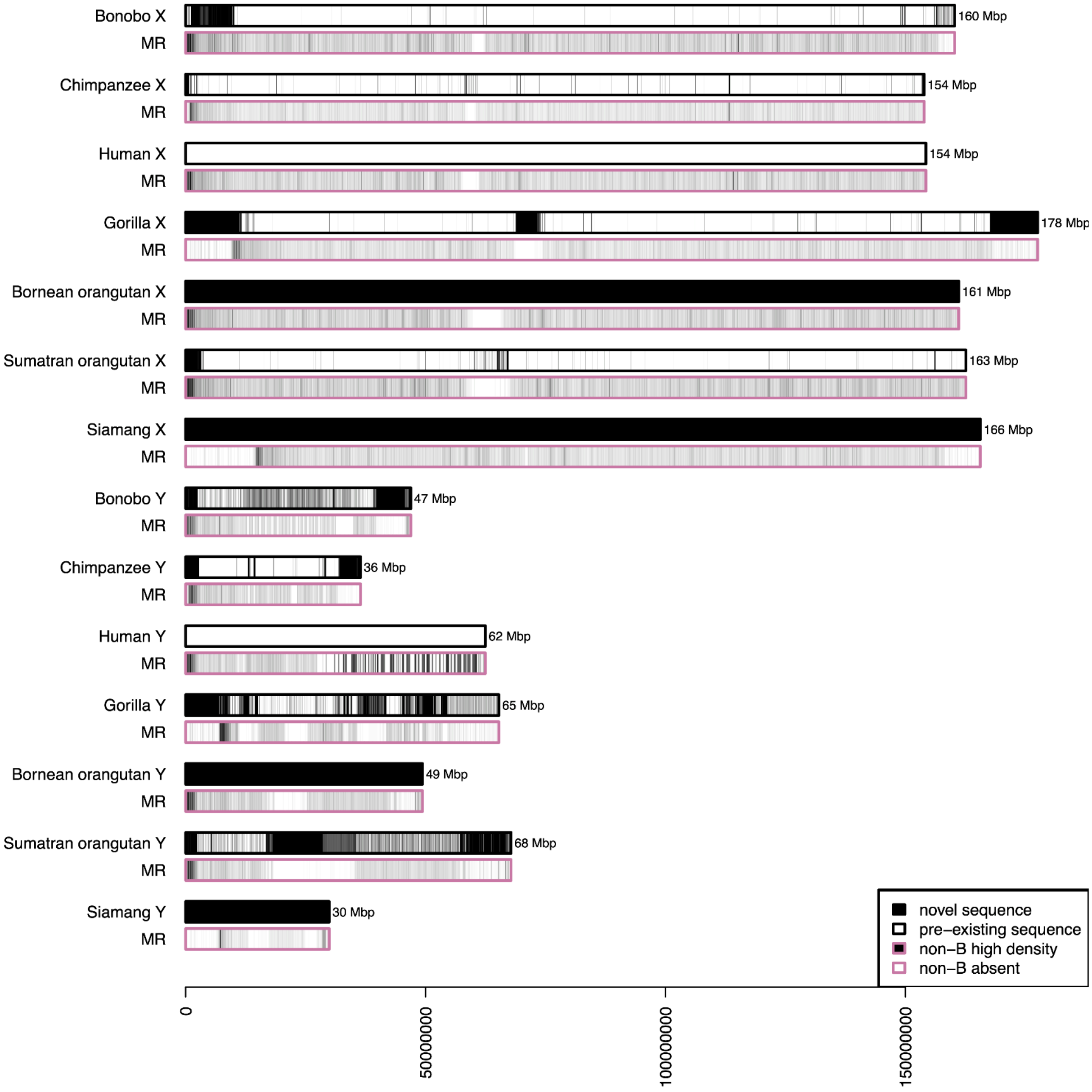
Bonobo X
MR
160 Mbp
Chimpanzee X
MR
154 Mbp
Human X
MR
154 Mbp
Gorilla X
MR
178 Mbp
Bornean orangutan X
MR
161 Mbp
Sumatran orangutan X
MR
163 Mbp
Siamang X
MR
166 Mbp
Bonobo Y
MR
47 Mbp
Chimpanzee Y
MR
36 Mbp
Human Y
MR
62 Mbp
Gorilla Y
MR
65 Mbp
Bornean orangutan Y
MR
49 Mbp
Sumatran orangutan Y
MR
68 Mbp
Siamang Y
MR
30 Mbp
novel sequence
pre−existing sequence
non−B high density
non−B absent
0
50000000
100000000
150000000

**E**

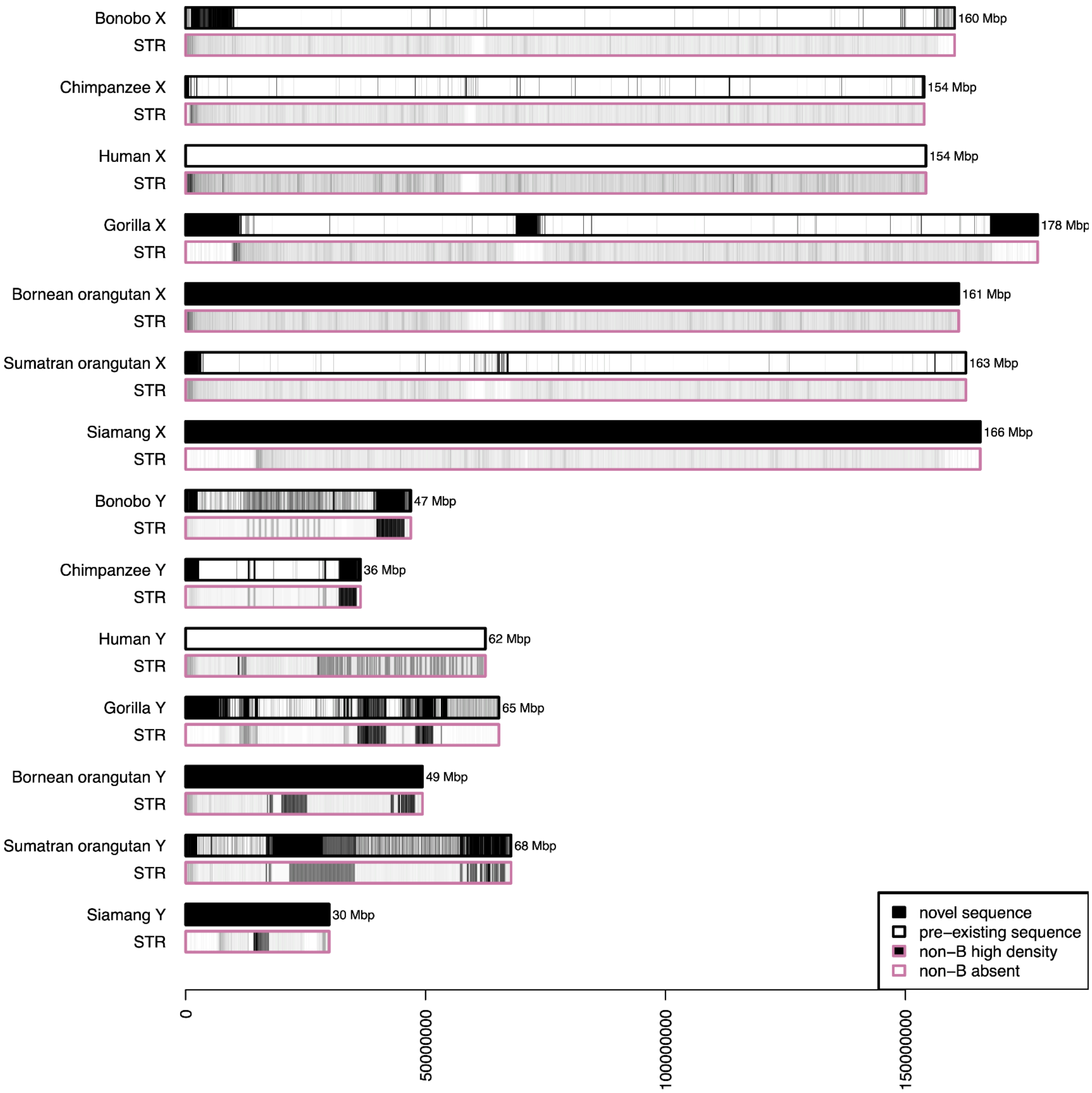

Bonobo X
STR
160 Mbp
Chimpanzee X
STR
154 Mbp
Human X
STR
154 Mbp
Gorilla X
STR
178 Mbp
Bornean orangutan X
STR
161 Mbp
Sumatran orangutan X
STR
163 Mbp
Siamang X
STR
166 Mbp
Bonobo Y
STR
47 Mbp
Chimpanzee Y
STR
36 Mbp
Human Y
STR
62 Mbp
Gorilla Y
STR
65 Mbp
Bornean orangutan Y
STR
49 Mbp
Sumatran orangutan Y
STR
68 Mbp
Siamang Y
STR
30 Mbp
novel sequence
pre–existing sequence
non–B high density
non–B absent
0
50000000
100000000
150000000

**F**

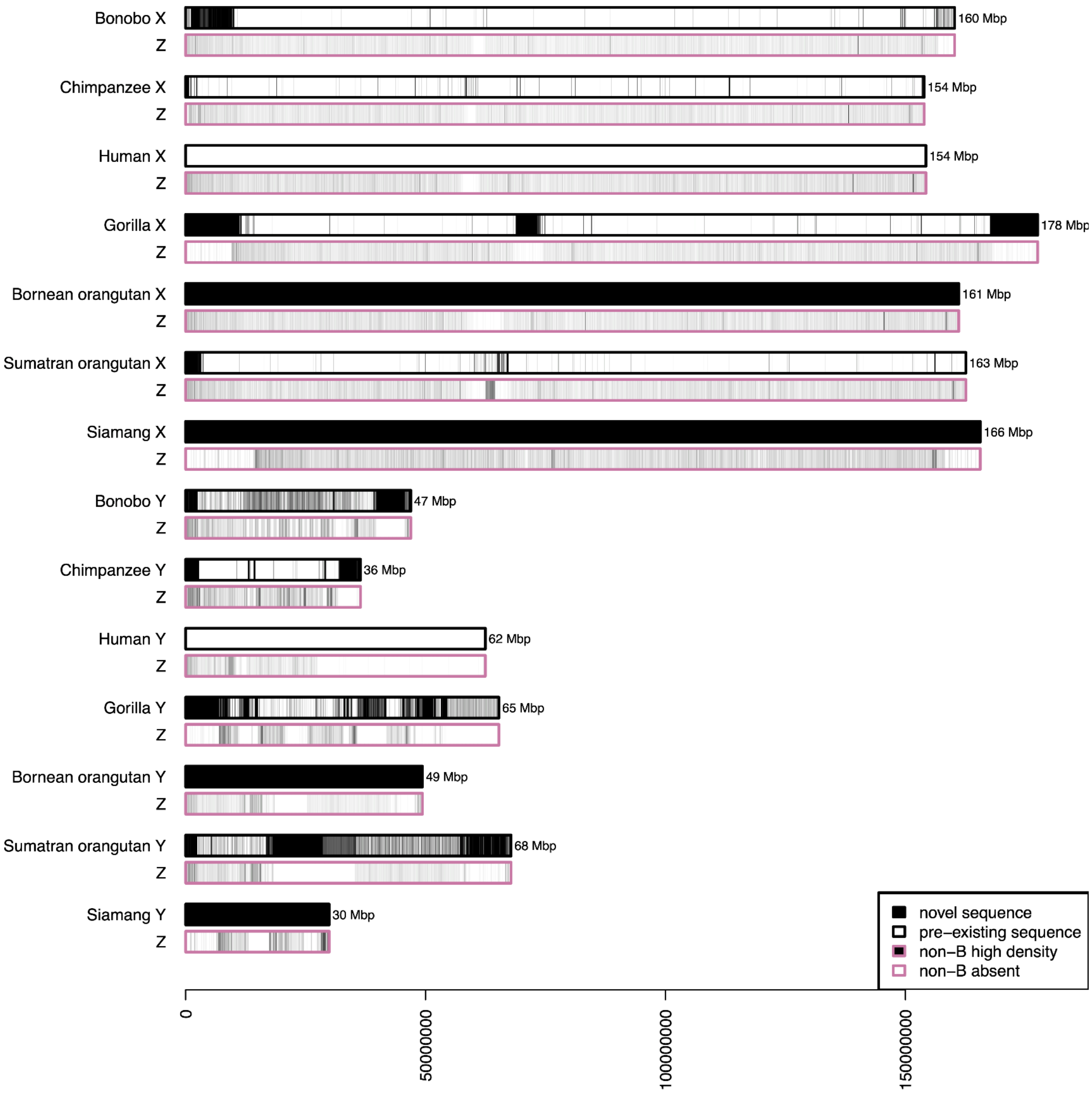

Bonobo X
Z
160 Mbp
Chimpanzee X
Z
154 Mbp
Human X
Z
154 Mbp
Gorilla X
Z
178 Mbp
Bornean orangutan X
Z
161 Mbp
Sumatran orangutan X
Z
163 Mbp
Siamang X
Z
166 Mbp
Bonobo Y
Z
47 Mbp
Chimpanzee Y
Z
36 Mbp
Human Y
Z
62 Mbp
Gorilla Y
Z
65 Mbp
Bornean orangutan Y
Z
49 Mbp
Sumatran orangutan Y
Z
68 Mbp
Siamang Y
Z
30 Mbp
novel sequence
pre−existing sequence
non−B high density
non−B absent
0
50000000
100000000
150000000

Figure S13. The fold enrichment of transposable elements and satellites in particular types of non-B DNA, as compared to non-B DNA average frequency on the X and Y

(**A**) bonobo and chimpanzee, (**B**) human and gorilla, (**C**) B. orangutan and S. orangutan, and (**D**) siamang.

**A**

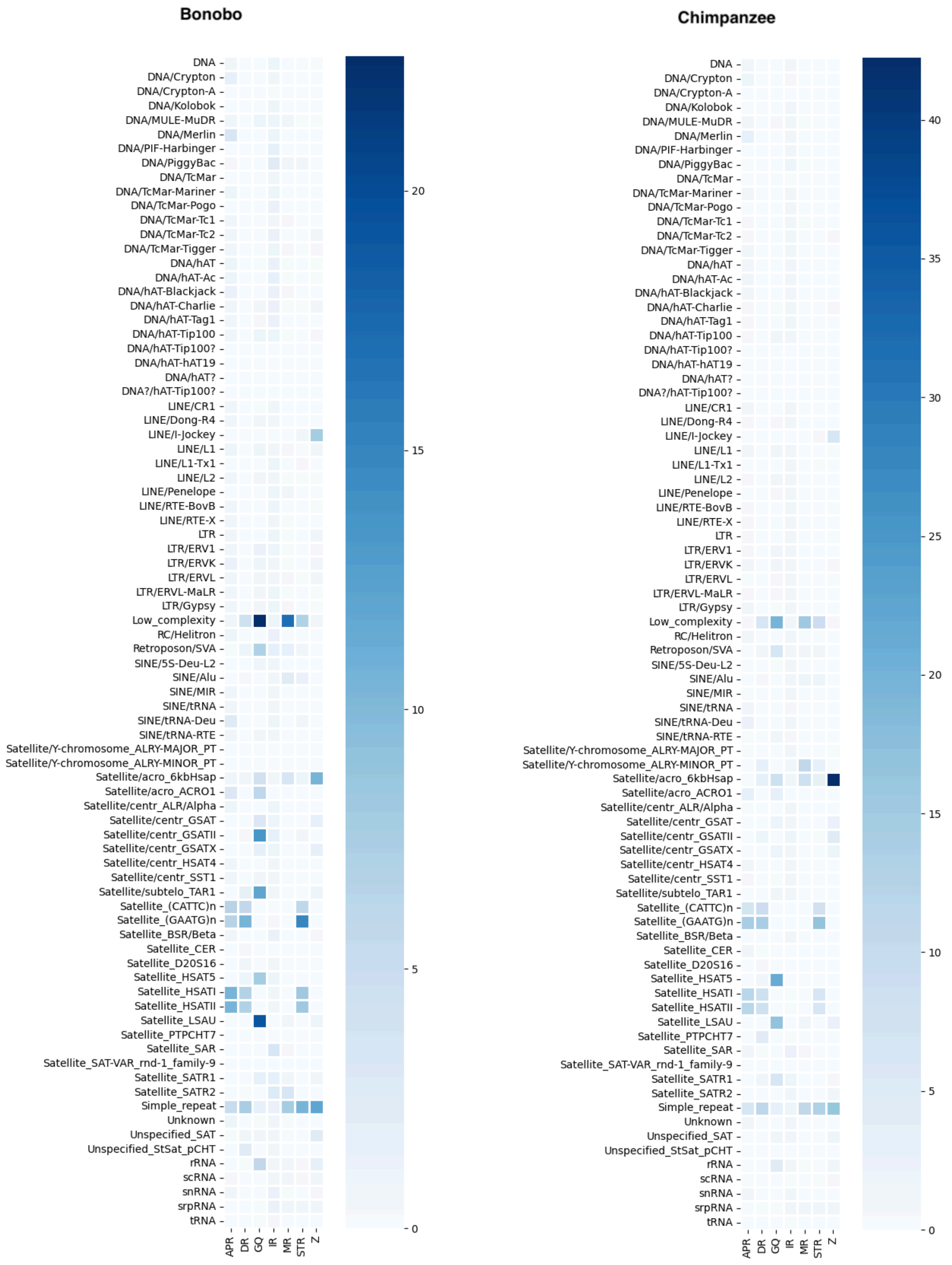

**B**

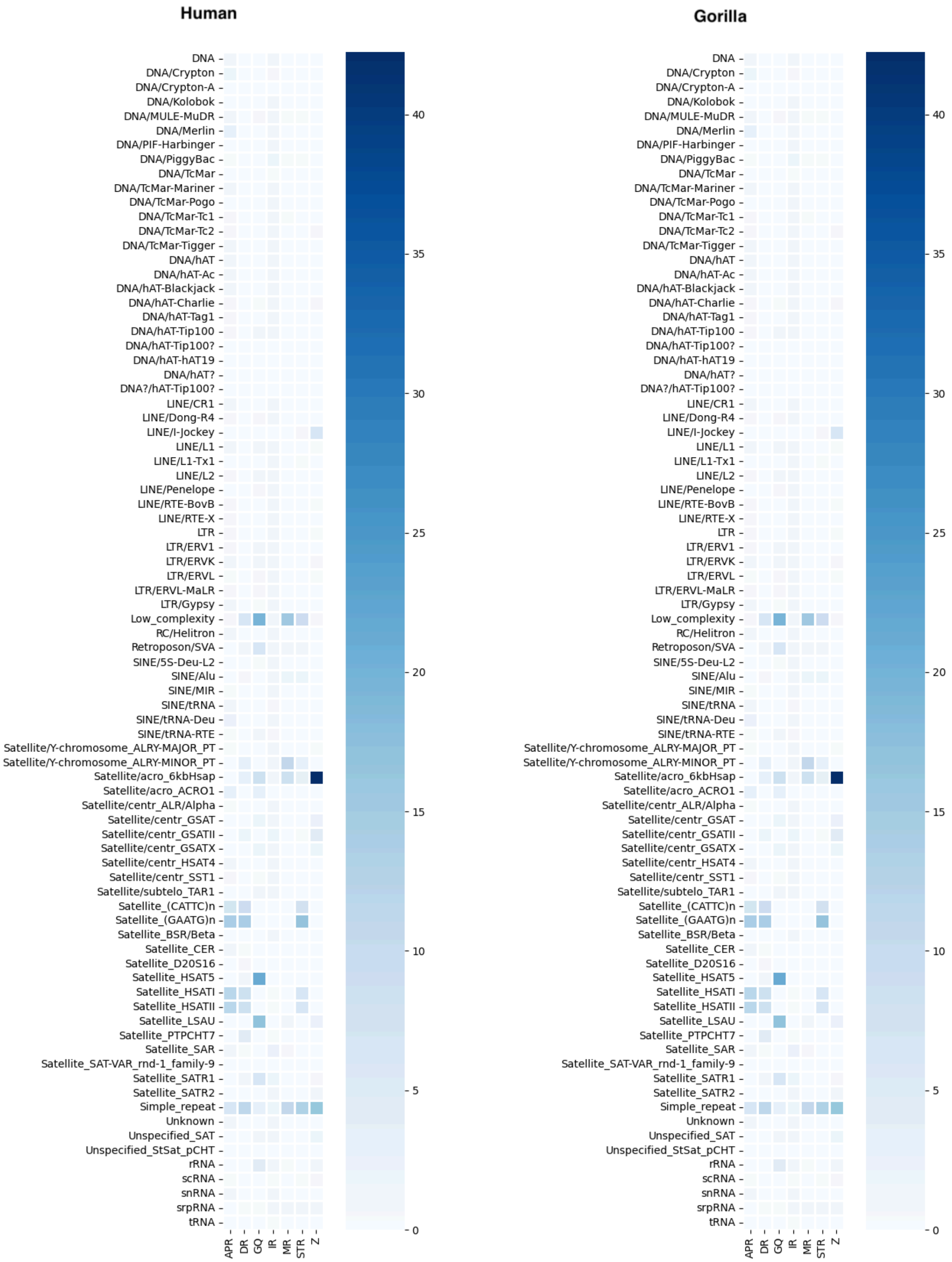
Human
Gorilla
DNA
DNA/Crypton
DNA/Crypton-A
DNA/Kolobok
DNA/MULE-MuDR
DNA/Merlin
DNA/PIF-Harbinger
DNA/PiggyBac
DNA/TcMar
DNA/TcMar-Mariner
DNA/TcMar-Pogo
DNA/TcMar-Tc1
DNA/TcMar-Tc2
DNA/TcMar-Tigger
DNA/hAT
DNA/hAT-Ac
DNA/hAT-Blackjack
DNA/hAT-Charlie
DNA/hAT-Tag1
DNA/hAT-Tip100
DNA/hAT-Tip100?
DNA/hAT-hAT19
DNA/hAT?
DNA?/hAT-Tip100?
LINE/CR1
LINE/Dong-R4
LINE/I-Jockey
LINE/L1
LINE/L1-Tx1
LINE/L2
LINE/Penelope
LINE/RTE-BovB
LINE/RTE-X
LTR
LTR/ERV1
LTR/ERVK
LTR/ERVL
LTR/ERVL-MaLR
LTR/Gypsy
Low_complexity
RC/Helitron
Retroposon/SVA
SINE/5S-Deu-L2
SINE/Alu
SINE/MIR
SINE/tRNA
SINE/tRNA-Deu
SINE/tRNA-RTE
Satellite/Y-chromosome_ALRY-MAJOR_PT
Satellite/Y-chromosome_ALRY-MINOR_PT
Satellite/acro_6kbHsap
Satellite/acro_ACRO1
Satellite/centr_ALR/Alpha
Satellite/centr_GSAT
Satellite/centr_GSATII
Satellite/centr_GSATX
Satellite/centr_HSAT4
Satellite/centr_SST1
Satellite/subtelo_TAR1
Satellite_(CATTC)n
Satellite_(GAATG)n
Satellite_BSR/Beta
Satellite_CER
Satellite_D20S16
Satellite_HSAT5
Satellite_HSATI
Satellite_HSATII
Satellite_LSAU
Satellite_PTPCHT7
Satellite_SAR
Satellite_SAT-VAR_rnd-1_family-9
Satellite_SATR1
Satellite_SATR2
Simple_repeat
Unknown
Unspecified_SAT
Unspecified_StSat_pCHT
rRNA
scRNA
snRNA
srpRNA
tRNA
APR
DR
GQ
IR
MR
STR
Z
40
35
30
25
20
15
10
5
0

C

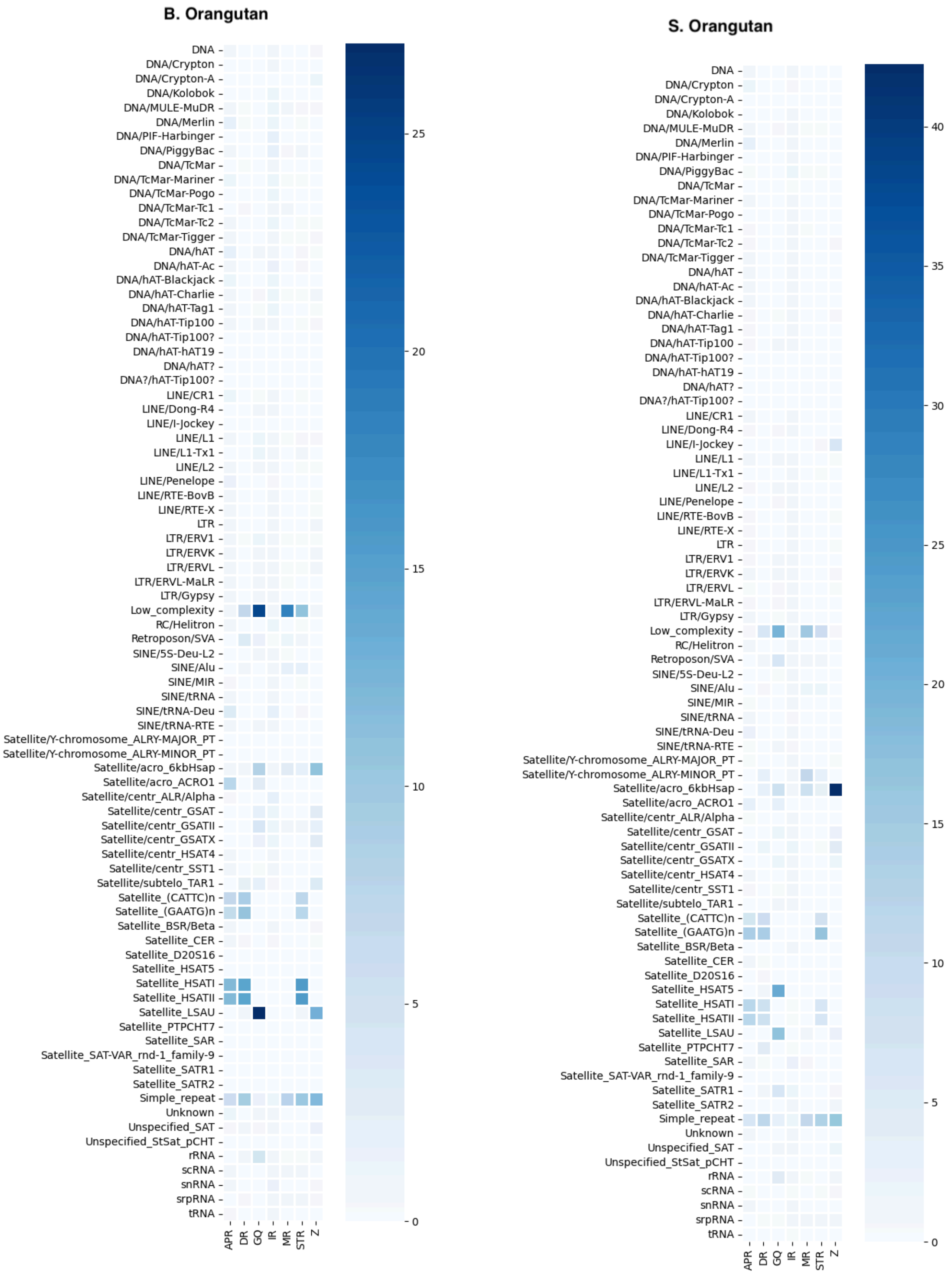

B. Orangutan
DNA
DNA/Crypton
DNA/Crypton-A
DNA/Kolobok
DNA/MULE-MuDR
DNA/Merlin
DNA/PIF-Harbinger
DNA/PiggyBac
DNA/TcMar
DNA/TcMar-Mariner
DNA/TcMar-Pogo
DNA/TcMar-Tc1
DNA/TcMar-Tc2
DNA/TcMar-Tigger
DNA/hAT
DNA/hAT-Ac
DNA/hAT-Blackjack
DNA/hAT-Charlie
DNA/hAT-Tag1
DNA/hAT-Tip100
DNA/hAT-Tip100?
DNA/hAT-hAT19
DNA/hAT?
DNA?/hAT-Tip100?
LINE/CR1
LINE/Dong-R4
LINE/I-Jockey
LINE/L1
LINE/L1-Tx1
LINE/L2
LINE/Penelope
LINE/RTE-BovB
LINE/RTE-X
LTR
LTR/ERV1
LTR/ERVK
LTR/ERVL
LTR/ERVL-MaLR
LTR/Gypsy
Low_complexity
RC/Helitron
Retroposon/SVA
SINE/5S-Deu-L2
SINE/Alu
SINE/MIR
SINE/tRNA
SINE/tRNA-Deu
SINE/tRNA-RTE
Satellite/Y-chromosome_ALRY-MAJOR_PT
Satellite/Y-chromosome_ALRY-MINOR_PT
Satellite/acro_6kbHsap
Satellite/acro_ACRO1
Satellite/centr_ALR/Alpha
Satellite/centr_GSAT
Satellite/centr_GSATII
Satellite/centr_GSATX
Satellite/centr_HSAT4
Satellite/centr_SST1
Satellite/subtelo_TAR1
Satellite_(CATTC)n
Satellite_(GAATG)n
Satellite_BSR/Beta
Satellite_CER
Satellite_D20S16
Satellite_HSAT5
Satellite_HSATI
Satellite_HSATII
Satellite_LSAU
Satellite_PTPCHT7
Satellite_SAR
Satellite_SAT-VAR_rnd-1_family-9
Satellite_SATR1
Satellite_SATR2
Simple_repeat
Unknown
Unspecified_SAT
Unspecified_StSat_pCHT
rRNA
scRNA
snRNA
srpRNA
tRNA
APR
DR
GQ
IR
MR
STR
Z
25
20
15
10
5
0
S. Orangutan
DNA
DNA/Crypton
DNA/Crypton-A
DNA/Kolobok
DNA/MULE-MuDR
DNA/Merlin
DNA/PIF-Harbinger
DNA/PiggyBac
DNA/TcMar
DNA/TcMar-Mariner
DNA/TcMar-Pogo
DNA/TcMar-Tc1
DNA/TcMar-Tc2
DNA/TcMar-Tigger
DNA/hAT
DNA/hAT-Ac
DNA/hAT-Blackjack
DNA/hAT-Charlie
DNA/hAT-Tag1
DNA/hAT-Tip100
DNA/hAT-Tip100?
DNA/hAT-hAT19
DNA/hAT?
DNA?/hAT-Tip100?
LINE/CR1
LINE/Dong-R4
LINE/I-Jockey
LINE/L1
LINE/L1-Tx1
LINE/L2
LINE/Penelope
LINE/RTE-BovB
LINE/RTE-X
LTR
LTR/ERV1
LTR/ERVK
LTR/ERVL
LTR/ERVL-MaLR
LTR/Gypsy
Low_complexity
RC/Helitron
Retroposon/SVA
SINE/5S-Deu-L2
SINE/Alu
SINE/MIR
SINE/tRNA
SINE/tRNA-Deu
SINE/tRNA-RTE
Satellite/Y-chromosome_ALRY-MAJOR_PT
Satellite/Y-chromosome_ALRY-MINOR_PT
Satellite/acro_6kbHsap
Satellite/acro_ACRO1
Satellite/centr_ALR/Alpha
Satellite/centr_GSAT
Satellite/centr_GSATII
Satellite/centr_GSATX
Satellite/centr_HSAT4
Satellite/centr_SST1
Satellite/subtelo_TAR1
Satellite_(CATTC)n
Satellite_(GAATG)n
Satellite_BSR/Beta
Satellite_CER
Satellite_D20S16
Satellite_HSAT5
Satellite_HSATI
Satellite_HSATII
Satellite_LSAU
Satellite_PTPCHT7
Satellite_SAR
Satellite_SAT-VAR_rnd-1_family-9
Satellite_SATR1
Satellite_SATR2
Simple_repeat
Unknown
Unspecified_SAT
Unspecified_StSat_pCHT
rRNA
scRNA
snRNA
srpRNA
tRNA
APR
DR
GQ
IR
MR
STR
Z
40
35
30
25
20
15
10
5
0

**D**

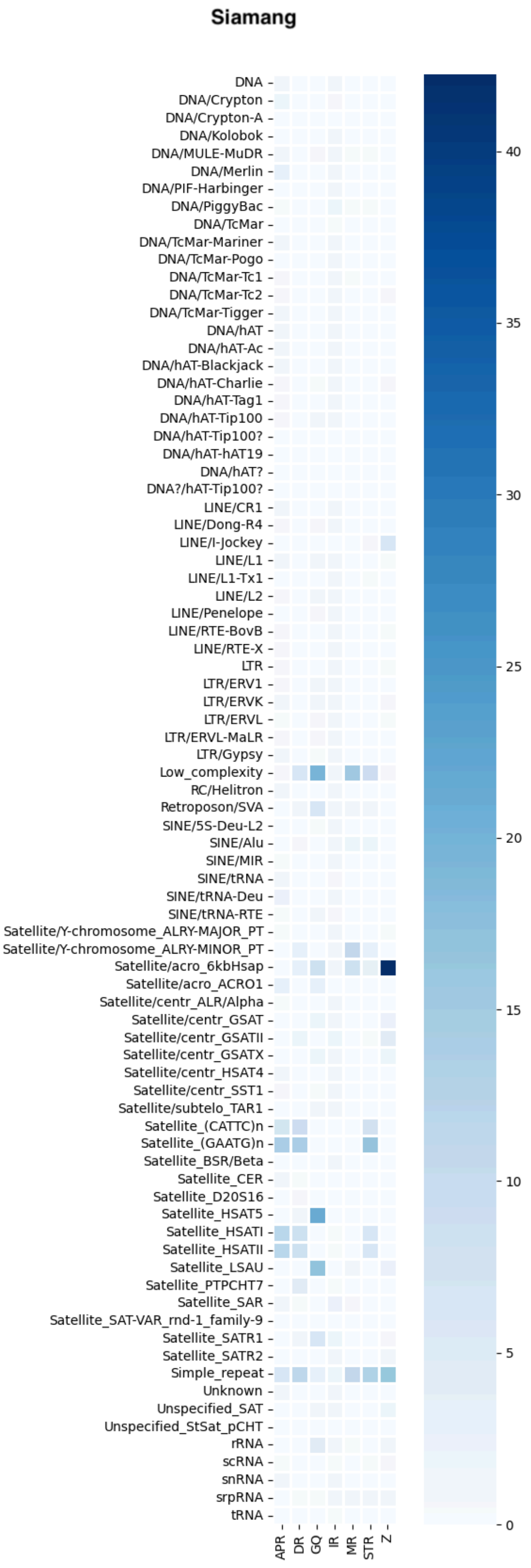

Siamang
DNA
DNA/Crypton
DNA/Crypton-A
DNA/Kolobok
DNA/MULE-MuDR
DNA/Merlin
DNA/PIF-Harbinger
DNA/PiggyBac
DNA/TcMar
DNA/TcMar-Mariner
DNA/TcMar-Pogo
DNA/TcMar-Tc1
DNA/TcMar-Tc2
DNA/TcMar-Tigger
DNA/hAT
DNA/hAT-Ac
DNA/hAT-Blackjack
DNA/hAT-Charlie
DNA/hAT-Tag1
DNA/hAT-Tip100
DNA/hAT-Tip100?
DNA/hAT-hAT19
DNA/hAT?
DNA?/hAT-Tip100?
LINE/CR1
LINE/Dong-R4
LINE/I-Jockey
LINE/L1
LINE/L1-Tx1
LINE/L2
LINE/Penelope
LINE/RTE-BovB
LINE/RTE-X
LTR
LTR/ERV1
LTR/ERVK
LTR/ERVL
LTR/ERVL-MaLR
LTR/Gypsy
Low_complexity
RC/Helitron
Retroposon/SVA
SINE/5S-Deu-L2
SINE/Alu
SINE/MIR
SINE/tRNA
SINE/tRNA-Deu
SINE/tRNA-RTE
Satellite/Y-chromosome_ALRY-MAJOR_PT
Satellite/Y-chromosome_ALRY-MINOR_PT
Satellite/acro_6kbHsap
Satellite/acro_ACRO1
Satellite/centr_ALR/Alpha
Satellite/centr_GSAT
Satellite/centr_GSATII
Satellite/centr_GSATX
Satellite/centr_HSAT4
Satellite/centr_SST1
Satellite/subtelo_TAR1
Satellite_(CATTC)n
Satellite_(GAATG)n
Satellite_BSR/Beta
Satellite_CER
Satellite_D20S16
Satellite_HSAT5
Satellite_HSATI
Satellite_HSATII
Satellite_LSAU
Satellite_PTPCHT7
Satellite_SAR
Satellite_SAT-VAR_rnd-1_family-9
Satellite_SATR1
Satellite_SATR2
Simple_repeat
Unknown
Unspecified_SAT
Unspecified_StSat_pCHT
rRNA
scRNA
snRNA
srpRNA
tRNA
APR
DR
GQ
IR
MR
STR
Z
40
35
30
25
20
15
10
5
0

#### Figure S14. Methylation patterns

(**A**) Differences in DNA methylation levels in long-range (100-kb) windows including PAR2. PAR2 is incorporated for comparison in bonobo and human. Boxplot represents the mean value and mean standard error from bootstrapping. P-values are determined using the Wilcoxon rank-sum tests (***$p<10^{-9}$; **$p< 10^{-6}$; *$p< 10^{-3}$). Only significant pairwise comparisons are shown. (**B-C**) DNA methylation across transcription units, shown from -3 kb of the transcription start site (TSS) to +3 kb of the transcription end site (TES) of protein-coding genes from (B) the X and (C) the Y chromosomes. (**D**) Spearman's correlations between promoter methylation and gene expression ranks. While X genes (N = 591) tended to show significant negative correlations, Y (N = 19) or PAR (N = 15) genes exhibited lower and often not significant correlations.

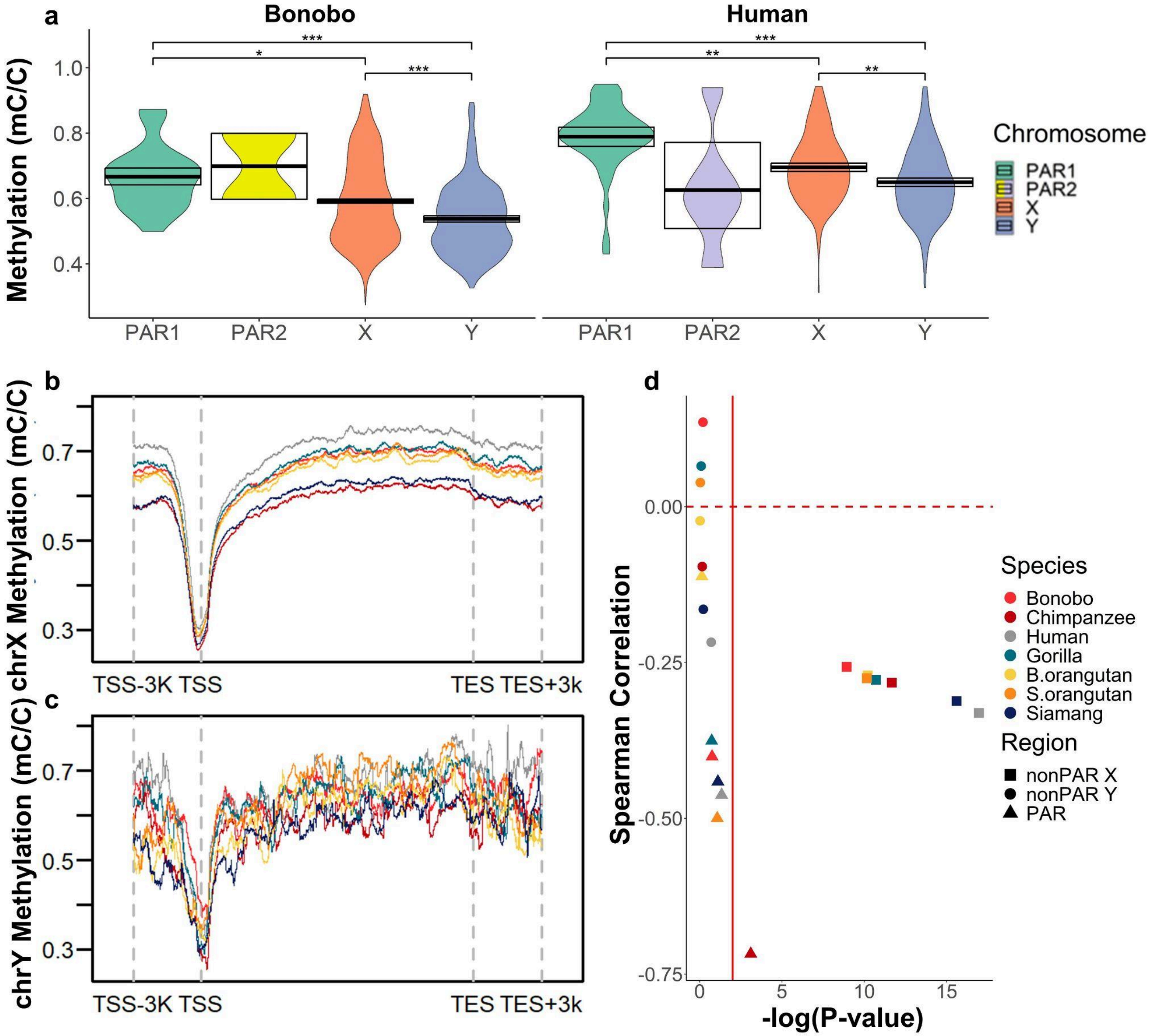

### Figure S15. Alpha satellite (AS) organization in primate X and Y centromeres

**(A)** AS suprafamily (SF) content of the whole Y chromosomes (upper panel) and X chromosomes (lower panel) of indicated species. In all cases but the siamang, the largest AS array is presumably centromeric. The smaller arrays indicate AS-containing SDs. The black horizontal bars show the short match track indicating the predicted CENP-B sites which have high density in cenXs except siamang and in gorilla cenY, but not in the other cenYs. Closely related species have the same SFs in the centromere and somewhat similar, but not identical, patterns of AS-containing SDs in the arms. Note that cenX in chimpanzee is inverted relative to human and bonobo (see details in Note S7). Siamang stands alone in having large sub-telomeric AS arrays, all of which are indicated, and two large non-telomeric arrays in cenY, either of which could serve as a centromere. Contrary to expectations shown Fig. 4B, human [13](HGOO2), chimpanzee and bonobo cenYs are formed by SF4, but not by the new families (lagging cenY). Also the much more variable position of cenY in a chromosome relative to cenX is observed. **(B)** and **(C)** Additional analysis of the orangutan species cenX. HOR-tracks, StV-tracks, SF-tracks, strand-tracks and methylation tracks covering the entire centromere are shown for each assembly (see details in Note S7). The patterns of monomeric layers at the ends of SF-tracks and the patterns of inversions at the ends in the strand-tracks are stable between species, and the general architecture of the centromere core is the same, but minor differences in domain organization are seen between species. Dips in methylation intensity (CDRs) at the right side of the orange arrays in both species indicate the kinetochore position, and thus active arrays. The StV tracks show that the HORs in the orange array are the mixture of two StVs, the full-length 4-mer HOR (monomers 1-4, shown in black) and the 2-mer deleted variant (monomers 3-4; shown in red). *Pongo abelii* has approximately equal proportions of both variants, and *Pongo pygmaeus* has a much smaller number of dimers, and these differences hold through the entire length of the arrays. **(C)** Quantification of the differences in tetramer and dimer content described in (**B**). Altogether, the evidence in panels b and c, and in Extended Data Fig. 4 indicates that all HOR domains in orangutan centromeres were already present before the split of the two species, but one or both active arrays have undergone near-complete remodeling after the split which led to species-specific HORhap and StV patterns. **(D)** Differences between the HOR consensus sequences in two closely related species; differences between *Pan* species are much larger than between *Pongo* species, in line with the relative differences in their divergence times. The differences between X and Y chromosomes apparently go against the genome-wide trend (Y evolves faster than X), however, we do not believe it to be a significant trend due to the sampling specifics in the centromeres (see Note S7). **(E)** Inversion analysis using SF- and AS strand-tracks demonstrating centromere inversion in gorilla cenX relative to other African apes and the changes in the centromere identity in the gorilla branch. The upper left panel shows the chimpanzee centromere that represents the ancestral architecture, which is still shared by humans, chimpanzee, and bonobo, and was presumably shared by their common ancestor with gorilla, before the gorilla branch-specific inversion occurred. The active SF3 AS array appears in cyan color in the SF track and in reverse orientation in the strand track (red). This is the arrangement typical of human, chimpanzee and bonobo cenX. In the bottom left panel, a similar view of gorilla cenX is shown;cyan represents the split dead centromere displaced to the flanks in the SF-track, and the new centromere core appears in direct orientation in the strand track (blue). The right two panels show the close-ups of the breakpoints, which are indicated by a red/blue color switch in the strand track overlapped by cyan in the HOR track. One can also observe the cenX identity changes with cyan (SF3) on both flanks, purple (SF2) next to it proximally, and pink (SF1) in the center. Hence the centromere has been inverted either before or after the identity changes, and both inversion breakpoints are in the SF3 arrays. **(F)** Unusual organization of AS in siamang chromosomes X and Y. SF-tracks show that the majority of AS in these chromosomes is SF4 (yellow). Huge SF4 subtelomeric AS arrays are present on both arms in the X and on the short arm in the Y. Some parts of the subtelomeric arrays are inverted. Relationships between HORs in various domains are noted.

a

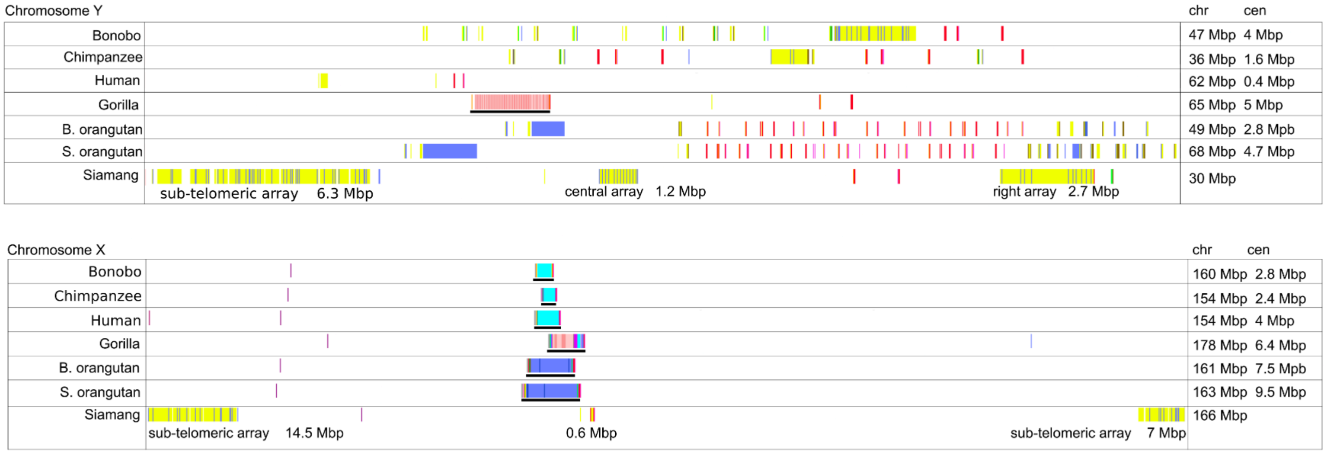

b

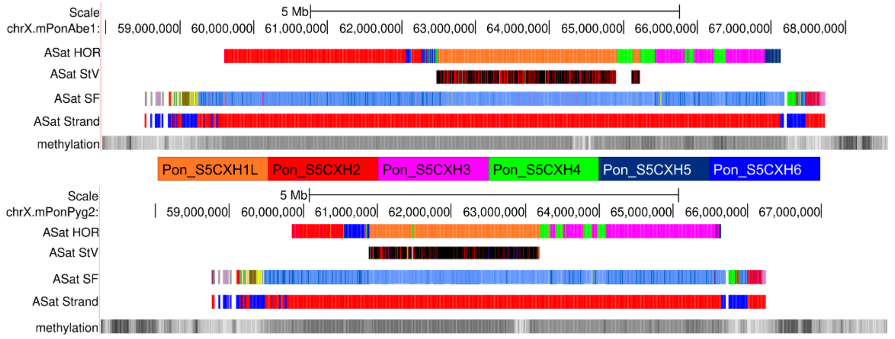

c

| stv (live orange array) | mPonPyg2 | mPonAbe1 |
|---|---|---|
| Pon_S5CXH1L.1-4 | 2915 | 2498 |
| Pon_S5CXH1L.3-4 | 896 | 2314 |
| Pon_S5CXH1L.1-2 | 30 | 56 |

d

| | HOR length (mons) | differences per HOR | differences per mon |
|---|---|---|---|
| Pan X | 12 | 60 | 5 |
| Pan Y | 29 | 110 | 3.8 |
| Pongo X | 4 | 3 | 0.75 |
| Pongo Y | 58 | 23 | 0.4 |

e

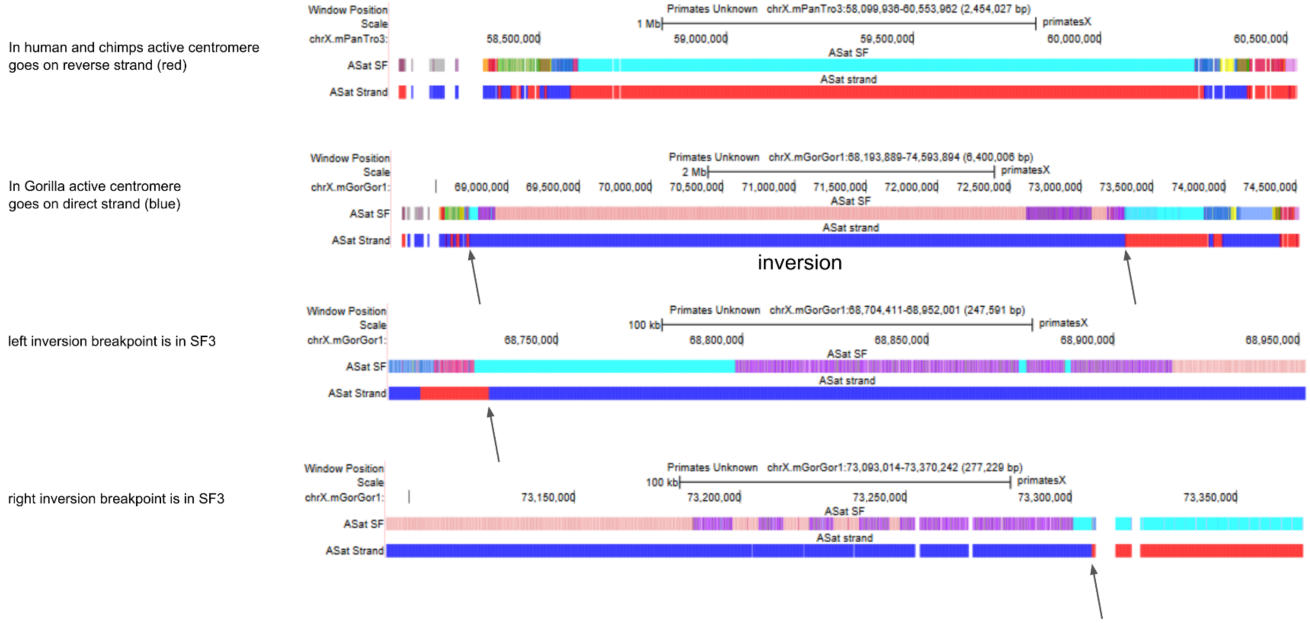

f

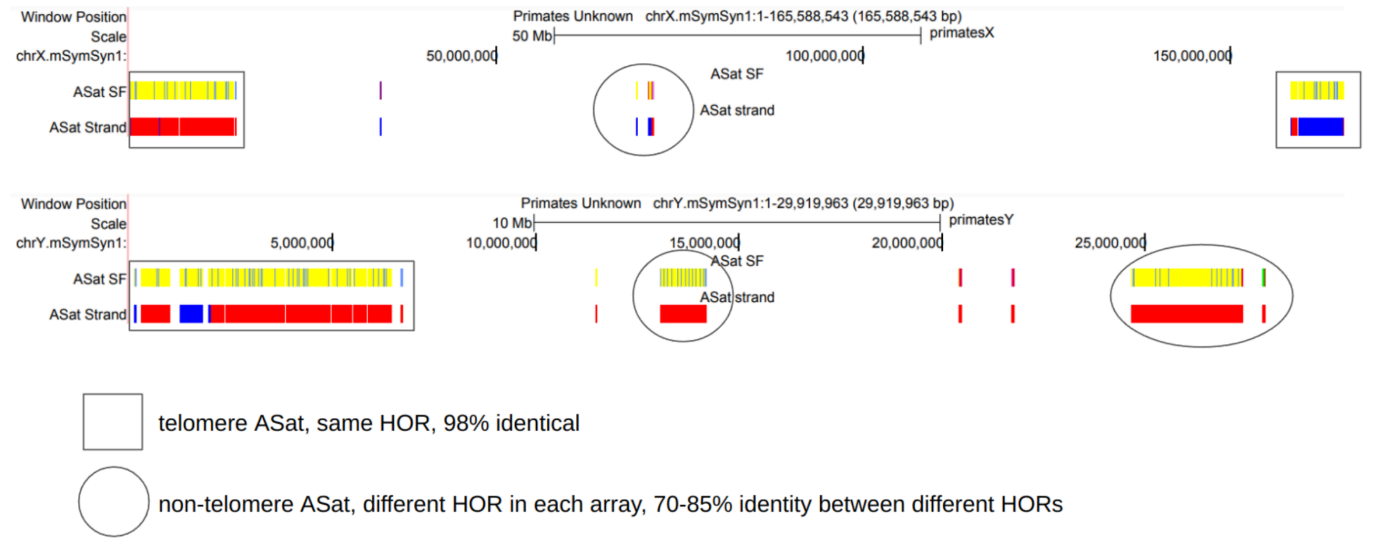
Window Position
Scale
chrX.mSymSyn1:
Primates Unknown chrX.mSymSyn1:1-165,588,543 (165,588,543 bp)
50 Mb
primatesX
50,000,000
100,000,000
150,000,000
ASat SF
ASat Strand
ASat SF
ASat strand
Window Position
Scale
chrY.mSymSyn1:
Primates Unknown chrY.mSymSyn1:1-29,919,963 (29,919,963 bp)
10 Mb
primatesY
5,000,000
10,000,000
15,000,000
20,000,000
25,000,000
ASat SF
ASat Strand
ASat SF
ASat strand
telomere ASat, same HOR, 98% identical
non-telomere ASat, different HOR in each array, 70-85% identity between different HORs

### Figure S16. Ribosomal DNA arrays

**(A)** Karyogram of all rDNA-containing chromosomes in S. orangutan. Chromosomes were labeled by FISH with probes for rDNA (BAC RP11-450E20, green) and *SRY* (BAC RP11-400O10, red). DNA was counter-stained with DAPI. Most of the somatic acrocentric chromosomes (9 pairs) contain rDNA arrays of various sizes on the p-arms, while the small chrY rDNA array is located on the distal q-arm. **(B)** Quantification of rDNA copy number on chromosome Y in Sumatran orangutan. Chromosome spreads were labeled by FISH with probes for rDNA and *SRY* as in panel A. The rDNA copy number on chromosome Y was calculated from its fraction of the total fluorescent intensity of the rDNA signals on all chromosomes and the Illumina sequencing estimate of the total copy number of rDNA repeats in the genome. The box plot shows means with standard deviations of all somatic rDNA-containing chromosomes and chromosome Y from 20 chromosome spreads. **(C)** Quantification of rDNA copy numbers on all rDNA-containing chromosomes in siamang. Chromosome spreads were labeled as in panel A. Since in this species rDNA is only present on one pair of somatic chromosomes and chromosome Y, all rDNA-containing chromosomes were discerned, and the copy number of rDNA repeats per chromosome was calculated as in panel B. Somatic non-Y rDNA chromosome pair had distinct large (L) and small (S) rDNA arrays, allowing haplotype separation by the array size. **(D)** Karyogram of all rDNA-containing chromosomes in siamang labeled by immuno-FISH with rDNA probe (green) and the antibody against rDNA transcription factor UBF (magenta). All rDNA arrays on both copies of somatic rDNA-containing chromosomes and on chrY were positive for the UBF signal. **(E)** Quantification of siamang rDNA and UBF expressed as the fraction of the total fluorescent intensity of all rDNA-containing chromosomes in a chromosome spread. The box plot shows means with standard deviations of rDNA and UBF fractions of the total signal present on chrY and both copies of somatic rDNA-containing chromosomes from 20 spreads.

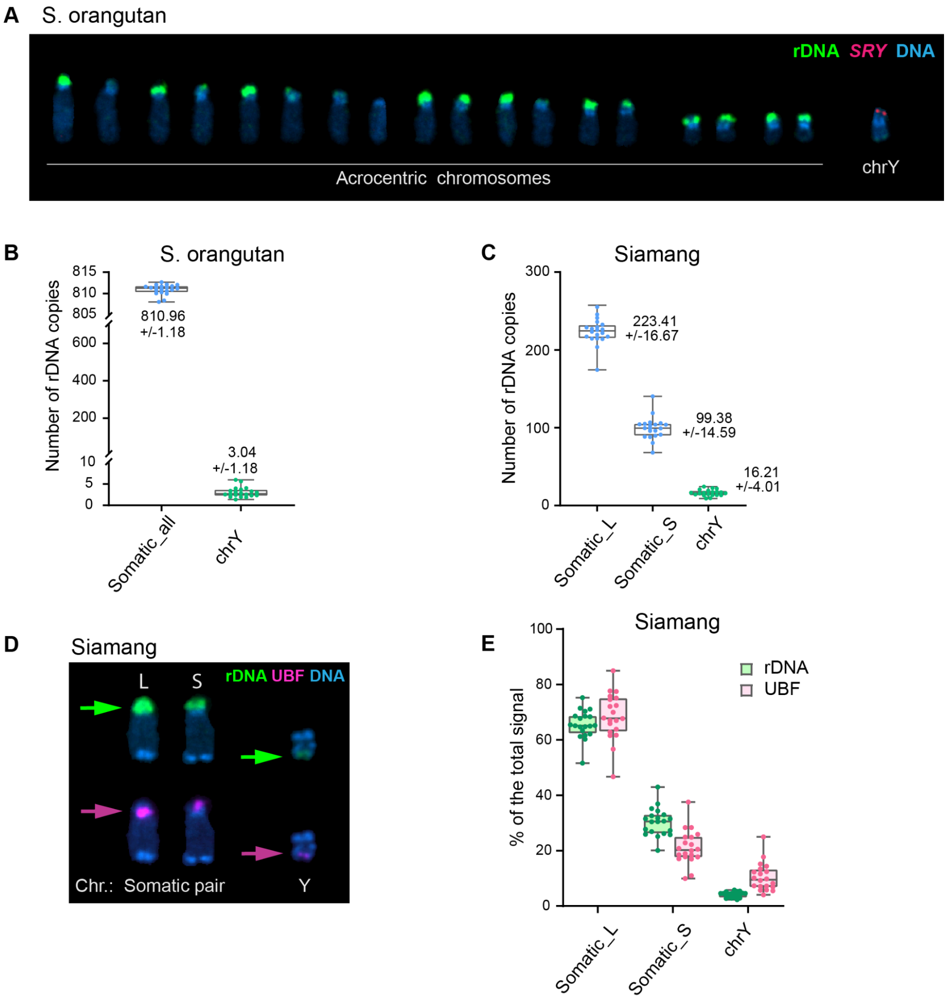
A S. orangutan
rDNA SRY DNA
Acrocentric chromosomes
chrY
B S. orangutan
Number of rDNA copies
815
810
805
600
400
200
10
5
0
810.96
+/-1.18
3.04
+/-1.18
Somatic_all
chrY
C Siamang
Number of rDNA copies
300
200
100
0
223.41
+/-16.67
99.38
+/-14.59
16.21
+/-4.01
Somatic_L
Somatic_S
chrY
D Siamang
L
S
rDNA UBF DNA
Chr.: Somatic pair
Y
E Siamang
% of the total signal
100
80
60
40
20
0
rDNA
UBF
Somatic_L
Somatic_S
chrY

### Figure S17. Diversity analyses

(**A**) The genome-wide mismatch rate using T2T and previous assemblies. In most cases, we observed lower mismatch rates between reads and references for the T2T vs. previous assemblies, demonstrating the superiority of the T2T assemblies as new reference genomes. The gorilla and chimpanzee X chromosomes exhibited similar mismatch rates between the old and the new assemblies, suggesting the high quality of the previous assemblies. The chimpanzee Y chromosome had a higher mismatch rate in the T2T than previous assembly, likely driven by the addition of repetitive regions and multi-mapping of some reads to them. (**B**) Variant counts for T2T, T2T masking-PARs (Pseudoautosomal regions), and previous assemblies. In some cases (e.g. bonobo X, Bornean orangutan X, Sumatran orangutan X, bonobo Y, and gorilla Y), we called fewer variants with the T2T as reference compared to the previous assembly, likely due to the reduced mismatch rate of the former assemblies and the use of species-specific references for Bornean orangutan. In other cases (e.g. bonobo Y and chimpanzee Y), we called more variants, which we attribute to the increased length and resolution of the new assemblies. (**C**) Allele frequency histogram of variants across shared subspecies samples relative to the T2T and previous genome assemblies. The x-axis indicates the number of subspecies samples, and the y-axis represents the percentage of variants observed within these samples.

**A**

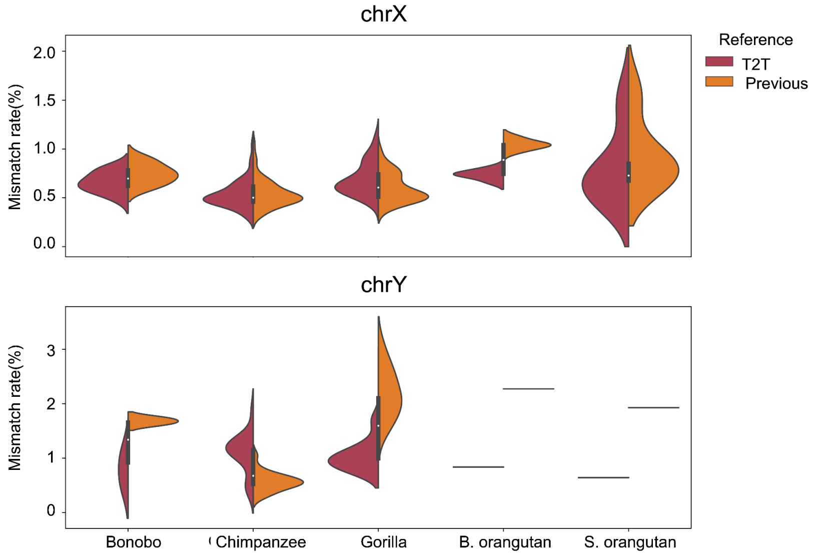

**B**

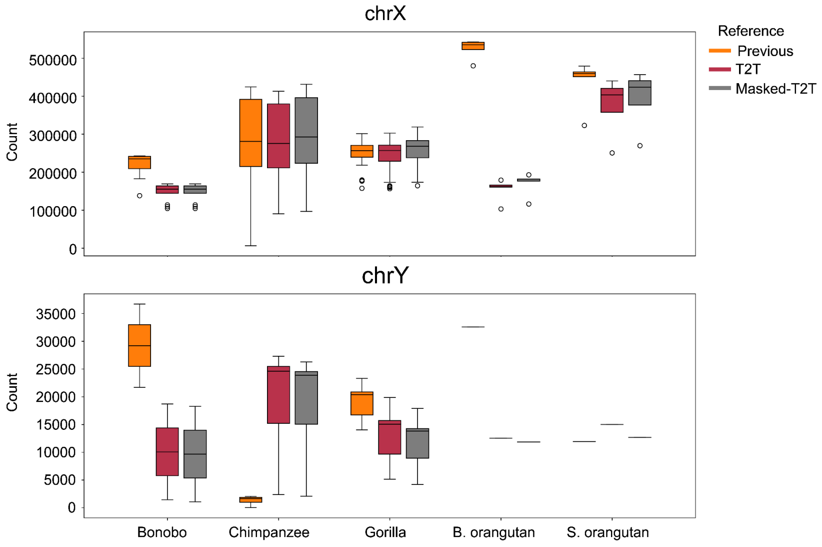

**C**

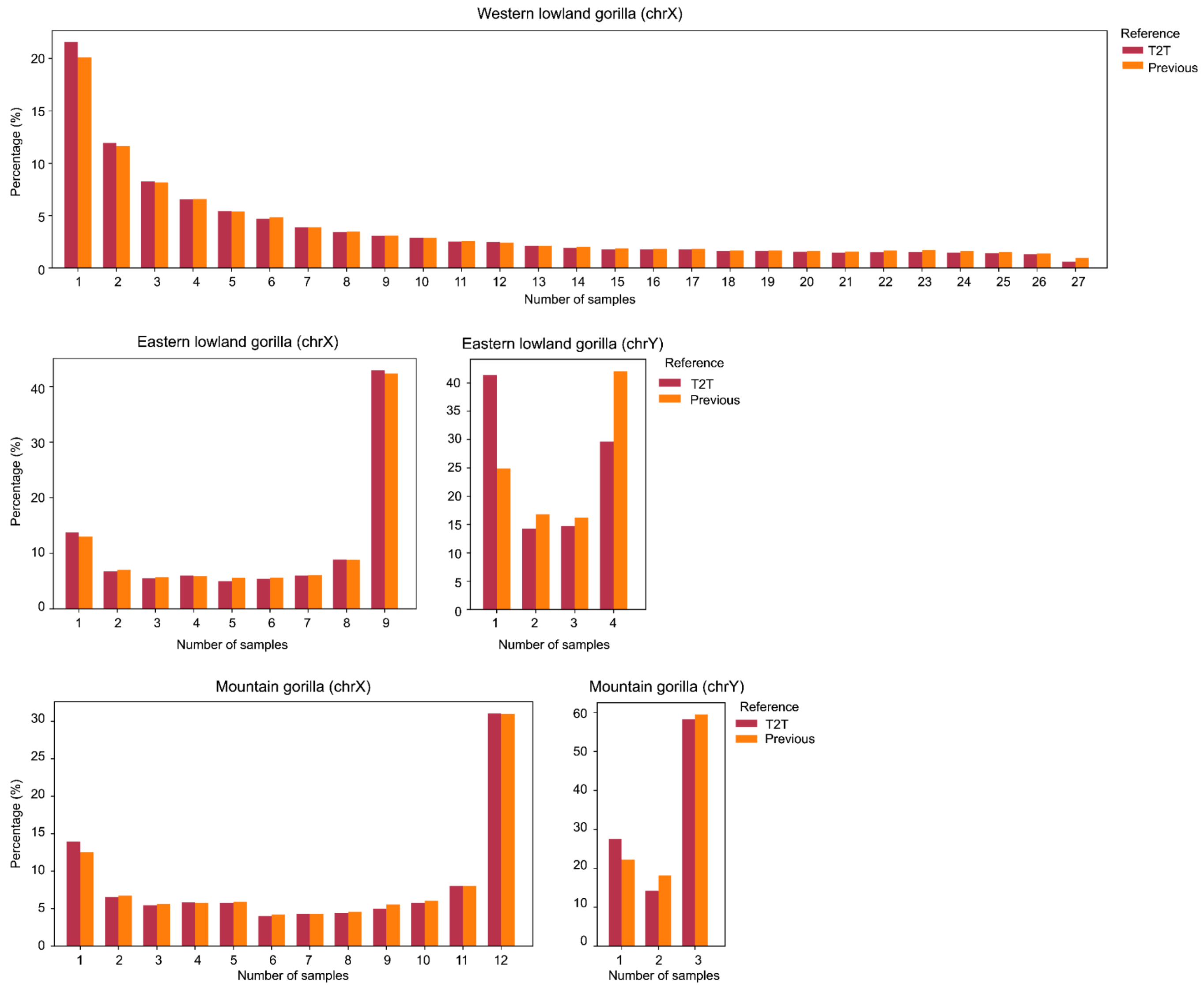
Western lowland gorilla (chrX)
Percentage (%)
Number of samples
Reference
T2T
Previous
Eastern lowland gorilla (chrX)
Eastern lowland gorilla (chrY)
Mountain gorilla (chrX)
Mountain gorilla (chrY)

### Figure S18. Methylation levels analyzed with ONT and PacBio HiFi long reads for the Y chromosomes agree across satellites

(**A**) bonobo; (**B**) chimpanzee; (**C**) gorilla; (**D**) B. orangutan; (**E**) S. orangutan; (**F**) siamang. The first two rows show HiFi and ONT coverage. The next two rows show the % of 5mC over C methylation for each HiFi and ONT read. Unlike seen in the human Yq12 heterochromatic region [13], where methylation and coverage disagreed across the HSat1B and HSat3 satellites, the methylation pattern between the two platforms concordantly agrees across different satellite repeat structures regardless of the non-human primate species in this study, at least for the sex chromosomes. Two-base microsatellite repeat composition pattern (e.g. AT = % bases composed with stretches of consecutive A and T bases in every 128 bp non-overlapping window) is shown as AT, GC, GA and TC tracks. Sequence class annotation is shown at the bottom, with color codes used as in Fig. 2.

**A**

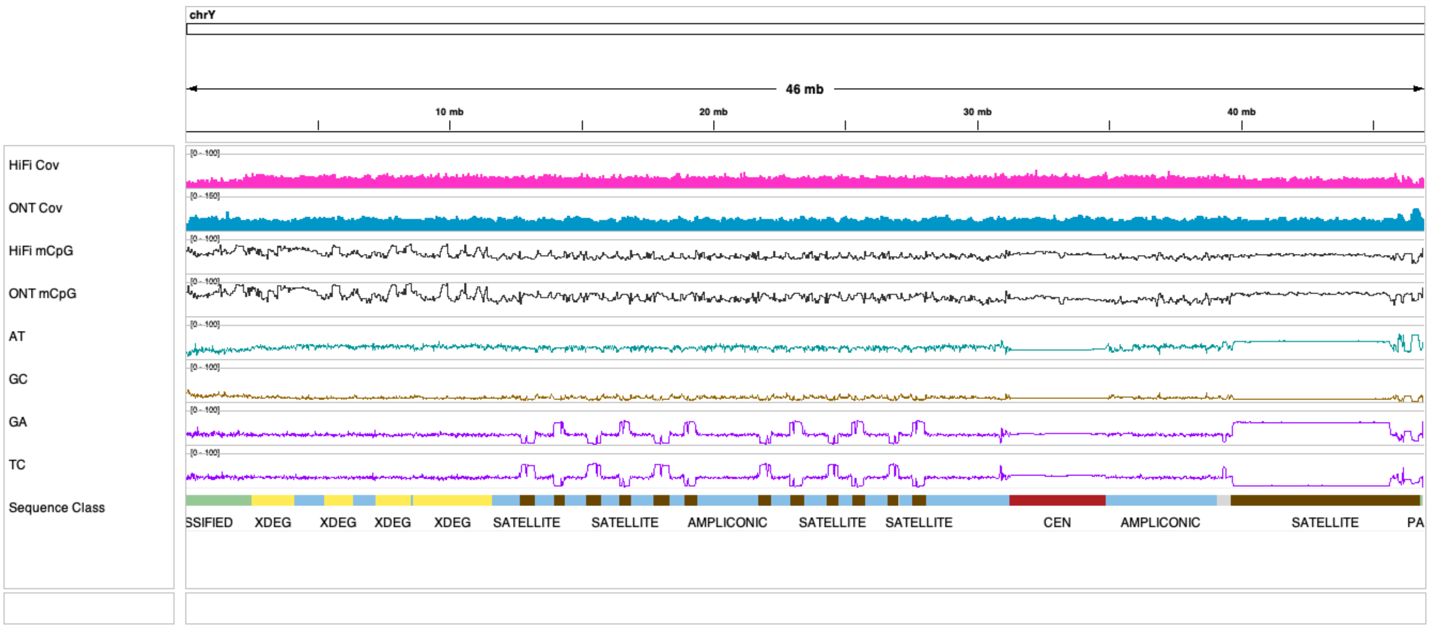

**B**

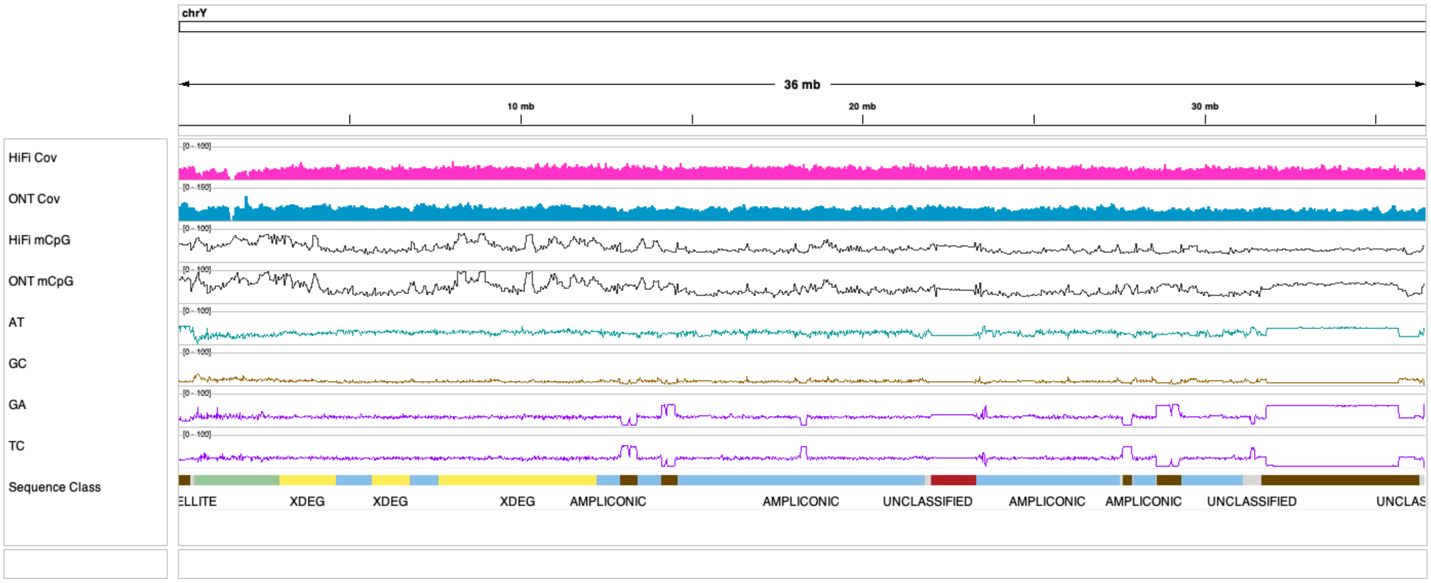

C

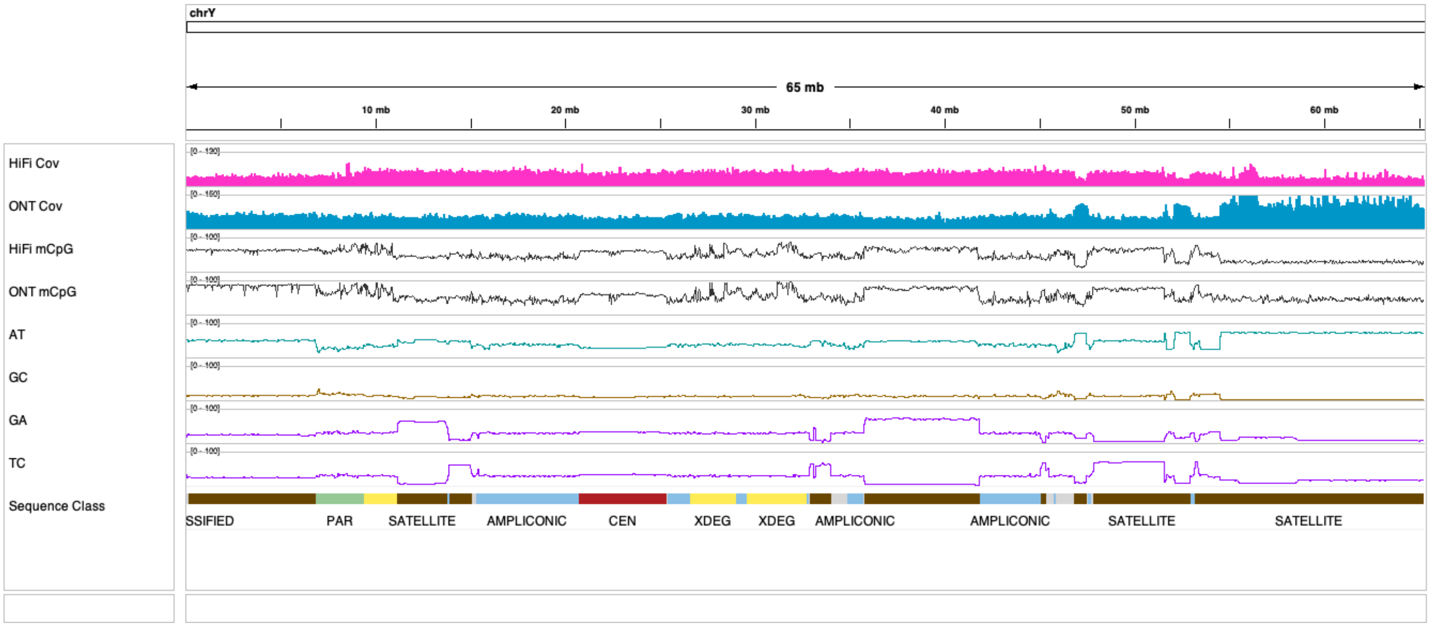
chrY
65 mb
10 mb
20 mb
30 mb
40 mb
50 mb
60 mb
HiFi Cov
ONT Cov
HiFi mCpG
ONT mCpG
AT
GC
GA
TC
Sequence Class
SSIFIED
PAR
SATELLITE
AMPLICONIC
CEN
XDEG
XDEG
AMPLICONIC
AMPLICONIC
SATELLITE
SATELLITE

D

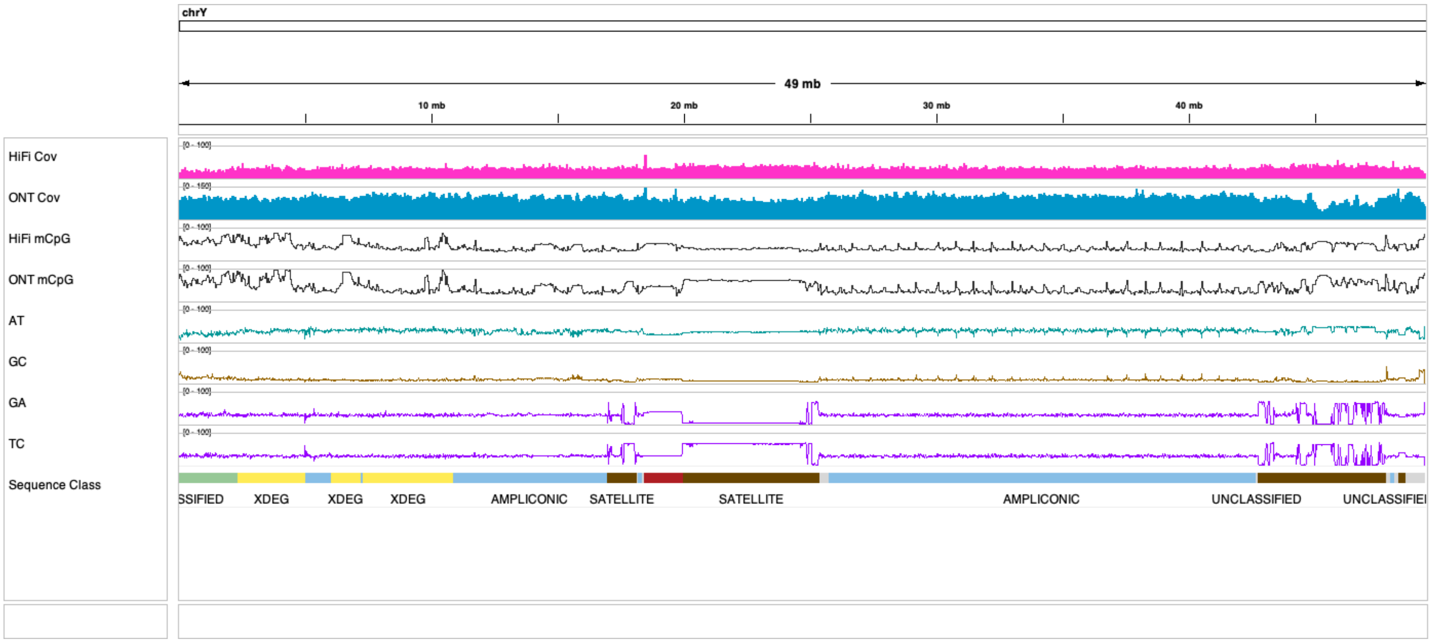
chrY
49 mb
10 mb
20 mb
30 mb
40 mb
HiFi Cov
ONT Cov
HiFi mCpG
ONT mCpG
AT
GC
GA
TC
Sequence Class
SSIFIED
XDEG
XDEG
XDEG
AMPLICONIC
SATELLITE
SATELLITE
AMPLICONIC
UNCLASSIFIED
UNCLASSIFIEI

**E**

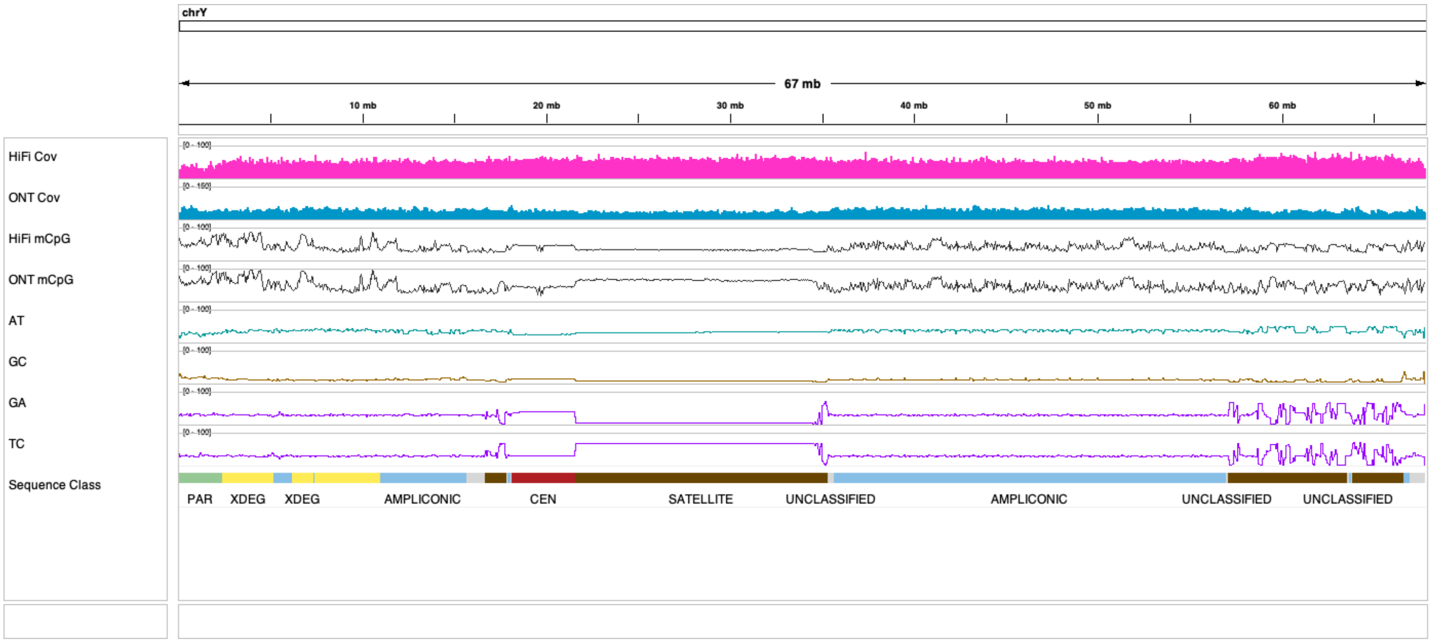
chrY
67 mb
10 mb
20 mb
30 mb
40 mb
50 mb
60 mb
HiFi Cov
ONT Cov
HiFi mCpG
ONT mCpG
AT
GC
GA
TC
Sequence Class
PAR
XDEG
XDEG
AMPLICONIC
CEN
SATELLITE
UNCLASSIFIED
AMPLICONIC
UNCLASSIFIED
UNCLASSIFIED

**F**

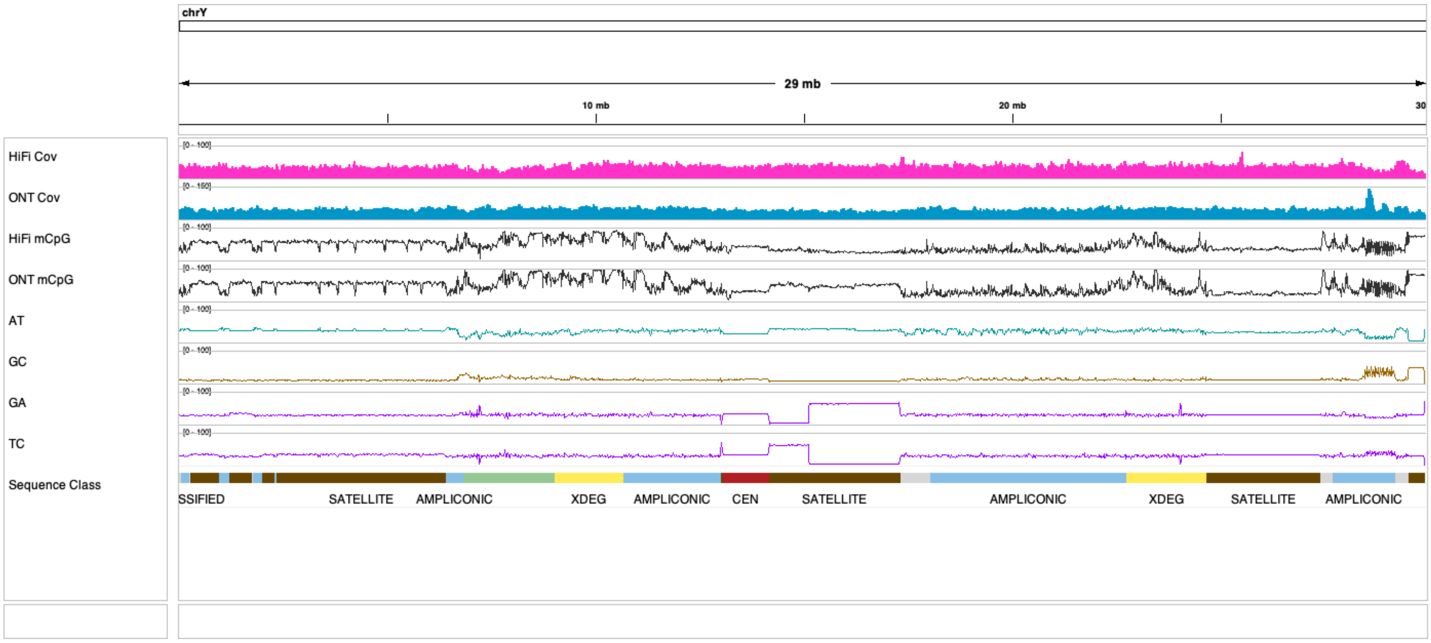
chrY
29 mb
10 mb
20 mb
30
HiFi Cov
ONT Cov
HiFi mCpG
ONT mCpG
AT
GC
GA
TC
Sequence Class
SSIFIED
SATELLITE
AMPLICONIC
XDEG
AMPLICONIC
CEN
SATELLITE
AMPLICONIC
XDEG
SATELLITE
AMPLICONIC

### Figure S19. *RBMY* copies in orangutan

(**A**) Similarity matrices of protein-coding copies (full genomic sequence) of the *RBMY* gene in the *Pongo* genus show two distinct groups in both the Sumatran and the Bornean orangutan. Identifiers of copies highlighted in bold are located on the same palindrome (one copy on each arm) while the others are repeated outside of the palindrome in tandem. Identifiers are based on NCBI annotations. (**B**) Overview of the alignments of individual copies. The copies located in palindromes are shown in the first two rows. Multisequence alignment done in (and screenshot captured from) the Geneious Prime program[14] using the Clustal Omega algorithm.

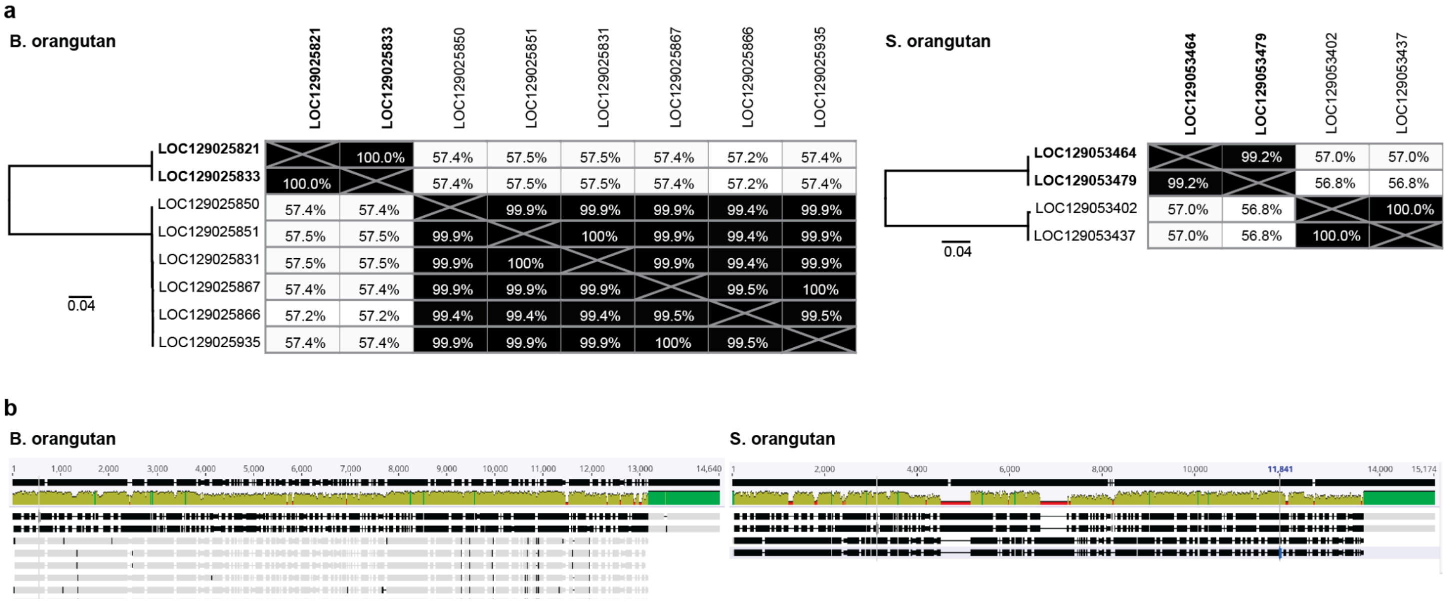

#### Figure S20. Distribution of pairwise identities for multi-copy genes on the Y chromosome within clusters of homology

The distribution of pairwise identities shows a natural breakpoint at 97% (black dashed line) which we chose as a cutoff for the identification of ampliconic gene families.

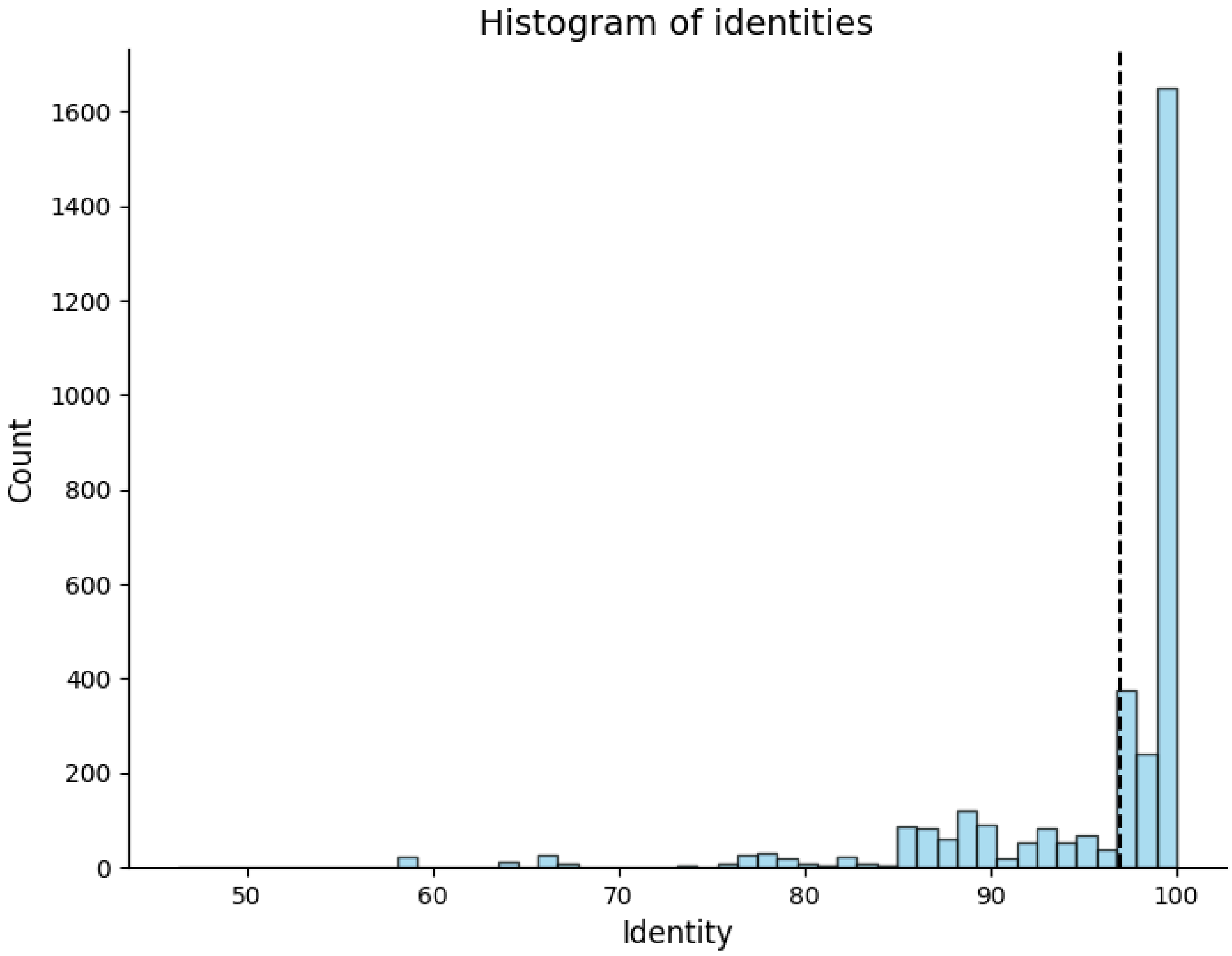

### Supplemental Methods

#### Cell lines

**Cell line propagation.** The cell lines used in the study are listed in Table S3. All cell lines were from male animals kept in captivity. The cell lines were propagated under the following conditions:

- Bonobo, Bornean orangutan, and gorilla fibroblasts: Alpha MEM with L-glutamine (Gibco) supplemented with 10% fetal bovine serum (Gemini Bio) and 1× Pen/Strep (Corning).
- Sumatran orangutan fibroblasts: Alpha MEM with L-glutamine supplemented with 15% fetal bovine serum, 1× Nonessential amino acids (Gibco) and 1× Pen/Strep (Corning).
- Siamang and chimpanzee lymphocytes: RPMI 1640 with phenol red and without L-glutamine (Gibco) supplemented with 10% fetal bovine serum, 1× Sodium Pyruvate (Gibco), 1× Nonessential amino acids, 1× Antibiotic/Antimycotic (Millipore-Sigma) , and freshly added 1× L-glutamine (Gibco). Medium was refrigerated for <2 months to maintain the appropriate L-glutamine concentration.

All cell lines were incubated at 37ºC with 5% $CO_2$ in a humidified chamber in standard tissue culture flasks with vented caps. Adherent fibroblast lines were maintained horizontally to produce cell monolayers. Non-adherent lymphocytes were incubated semi-upright to encourage formation of lymphocyte aggregates, which promote proliferation.

**Pellet generation.** To generate pellets, cells grown under standard culturing conditions were counted by hemocytometer and viability was determined by Trypan Blue exclusion (Gibco). For adherent cells, monolayers were trypsinized (0.25% Trypsin, 1mM EDTA; Gibco) and collected in an equal volume of growth medium containing FBS to inhibit further trypsinization. Lymphocytes were counted directly from the growth medium without trypsinization. The number of cells collected for each pellet was based on the approximate input required to generate genomic DNA for specific library preparations and analyses. To obtain a pellet, the desired number of cells were centrifuged for 8 minutes at room temperature at 1000g. After removing the supernatant by aspiration, pellets were resuspended in 1 ml of Dulbecco Phosphate Buffered Saline (DPBS; Corning), transferred to sterile screw cap 1.5-ml tubes and centrifuged again under the same conditions. Supernatants were removed by aspiration and the pellets were snap-frozen in liquid nitrogen and stored at -80ºC.

#### Karyotyping

Metaphase slide preparations were made from cultured fibroblast or lymphoblastoid cells lines after mitotic arrest with Colcemid (0.015 µg/mL, 16 to 18 hours) (GIBCO, Gaithersburg, MD), hypotonic treatment (0.075 mol/L KCl, 20 minutes, 37°C), and fixation with methanol–acetic acid (3:1). Slides were prepared by standard air-drying technique as described previously [15]. DAPI banding techniques were performed to identify structural and numerical chromosome aberrations [15]. Metaphases were analyzed with a fluorescent Microscope Zeiss M2 using Applied Spectral Imaging software INC, Carlsbad, CA.

#### Sequencing

**PacBio HiFi sequencing at the University of Washington.** High-molecular-weight DNA was isolated at PSU 5 mln. cells for each cell line using the Monarch HMW DNA Extraction Kit for Blood and Cells (New England Biolabs). PacBio HiFi data were collected from each sample as described in [16]. DNA quantity was assessed at receipt at the University of Washington PacBio Sequencing Services facility and at each subsequent step using the Qubit High Sensitivity DNA kit (ThermoFisher) read on a DS-11 FX instrument (DeNovix) and DNA fragment length distributions evaluated on a FEMTO Pulse capillary electrophoresis instrument (Agilent). Specifically, DNA was sheared using Megaruptor 3 (Diagenode) using settings to target 20-kb mode insert length. SMRTbell libraries were generated with the Express Template Prep Kit v2 (PacBio) (all samples) or SMRTbell Prep Kit v3 (PacBio) (for Jim_KB3781_GGO only) according to manufacturer's protocols. Size selection was performed with PippinHT (Sage Science) using a 15- or 17-kb high-pass protocol.

All HiFi libraries were sequenced at the University of Washington on a Sequel II instrument using 30-hour movie times and 2-hour pre-extension. Data sets were generated with P2/C2 chemistry (AG05252_PPY, AG06213_PAB, Jim_KB3781_GGO, PR00251_PPA), P2.2/C2 chemistry (AG05252_PPY, AG06213_PAB, AG18354_PTR, Jambi_SSY, Jim_KB3781_GGO, PR00251_PPA), or P3.2/C2 chemistry (Jim_KB3781_GGO). HiFi consensus (from subreads) was performed with pbccs v6+ (v6.0.0-6.5.0) for all SMRTcells, and with DeepConsensus v0.3rc0[17] for all but three (of 11) gorilla SMRTcells, which were sequenced later.

**Ultra-Long ONT sequencing at the NISC Sequencing Center (NIH).** Frozen cell pellets (containing 50 mln. cells each) generated at PSU were thawed and resuspended in 40 µl PBS for every 6 million cells. High molecular weight DNA was extracted using the protocol 'Nanobind UHMW DNA Extraction–Cultured, Cells Protocol' with Nanobind CBB Big DNA Kit (Pacific Biosciences). The DNA size was assessed on a pulse field gel. Quantitation and purity were determined using NanoDrop and Qubit (Thermo Fisher). Libraries were made using Ultra-Long DNA Sequencing Kit (SQK-ULK001,Oxford Nanopore) along with Nanobind UL Library Prep Kit with UHMW DNA Aux Kit (Pacific Biosciences). One extraction prep was used per library. FRA (fragmenting reagent) was added at a ratio of approximately 1 µl FRA per 14 ug DNA.

Each library was run for 72 hr on a PromethION flow cell version 9.4.1 using 3 loadings per flow cell. All samples were sequenced across 5 flow cells except for the Bornean orangutan, which was sequenced across 8 additional flow cells to achieve at least 100 GB in reads with length >100 kb. All flowcells were basecalled using Guppy v5+ on instrument, and the subset that was basecalled with <v6 was subsequently re-basecalled with v6+ until every flowcell had been basecalled with a Guppy version ranging from 6.0.0 to 6.1.1. These reads were used in the assembly. Some models did not enable methylation calling, and thus all flowcells were later basecalled again to obtain modified base information (see Supplemental Methods – CpG methylation calling).

**Hi-C library preparation at the UCSC.** Frozen pellets of 5 million cells per sample were stored at -80°C degrees at PSU prior to shipment to UC Santa Cruz on dry ice. We generated Omni-C libraries from primate cell lines using the Dovetail Omni-C kit (Cantata Bio) and followed the manufacturer's protocol with a few modifications. Briefly, we counted and aliquoted 1 million cells, which were then fixed using formaldehyde and DSG. We performed a nuclease digestion on the fixed cell aliquots using DNAse I and adjusted the concentration of nuclease until a suitable distribution of DNA fragments for proximity ligation was obtained. We then performed proximity ligation as described by the manufacturer, with end-repaired chromatin undergoing the ligation of a biotinylated bridge oligo prior to a final intra-aggregate ligation of bridge containing ends. We purified the proximity-ligated DNA product, which was used as input into an NEB Ultra II (New England Biolabs) library preparation. The resulting libraries underwent a streptavidin capture to enrich for biotin containing proximity-ligated products. We split the capture product into two replicates prior to the final index PCR in order to maximize the final complexity of the library.

**Hi-C library sequencing at PSU.** Fragment sizes of 150-1400 to 1600 bp, as determined by Fragment Analyzer, were included in the sequencing runs. Qubit was used for quantification of each sample. All dual indexed libraries were pooled and sequenced on an Illumina NovaSeq 6000 instrument using a single NovaSeq S2 flow cell; 400 million read pairs at 2×150 bp/sample by the Genome Sciences Core at Penn State Hershey. For demultiplexing, a bcl2fastq version 2.20.0 was used.

**Illumina sequencing at the NISC Sequencing Center (NIH).** PCR-free libraries were generated from 1 µg genomic DNA using a Covaris LE220-plus to shear the DNA and the TruSeq® DNA PCR-Free HT Sample Preparation Kit (Illumina) for library generation. The median insert sizes were ~400 bp. Libraries were tagged with unique dual index DNA barcodes to allow pooling of libraries and minimize the impact of barcode hopping. Libraries were pooled for sequencing on the NovaSeq 6000 (Illumina) using v.1.5 chemistry to obtain at least 570 million read pairs of 2x150-bp per individual library.

**IsoSeq sequencing at the University of Washington.** RNA isolated from testes tissues from the Makova Lab collection (Table S46) was prepared for PacBio Iso-Seq full-length transcriptome sequencing using the protocol 'Preparing Iso-Seq libraries using SMRTbell prep kit 3.0' (PacBio).

PTR_8720-2, PPA_5013: Samples were processed in two replicates each using barcoded adapter ligation for sample identification. After cDNA generation and amplification using the NEBNext Single Cell/Low Input cDNA Synthesis kit (New England Biolabs), samples were size-fractionated using a 0.86×/1.2× volume ratio of SMRTbell Cleanup Beads. The larger fraction was pooled with sufficient shorter fraction to bring sample mass up to 160 ng per replicate. After repooling, barcoded adapters (Table S46) from the Barcoded Overhang Adapter kit 8A/8B were added using the SMRTbell Prep Kit v3 (PacBio). All four replicates were pooled together for sequencing on three SMRT Cell 8Ms on a PacBio Sequel II instrument using chemistry P2.1/C2.0 or P2.2/C2.0. Samples were circular consensus-analyzed and demultiplexed in SMRT Link v11.0 and filtered for estimated quality scores ≥Q10. All replicate files per sample were processed together through the IsoSeq3 pipeline in SMRT Link.

OR6737_GGO, Ppyab_1991-51_PAB, Ppypy_3405_PPY: Samples were barcoded (Table S46) in the cDNA PCR step per the alternative option in the protocol. After reverse transcription and cleanup, the second strand synthesis and cDNA amplification step with barcoded primers (IDT) added sample barcodes to both ends of each molecule. Samples were size-fractionated and rebalanced as described above and prepared for sequencing using the SMRTbell Prep Kit v3 (PacBio). Libraries were sequenced on 1 or 2 SMRT Cell 8M per sample on a Sequel II instrument using chemistry P3.1/C2. Samples were circular consensus-analyzed in SMRT Link v11.0 and filtered for estimated quality scores ≥Q10. Each sample was processed for barcode removal through Iso-Seq analysis using the IsoSeq3 pipeline in SMRT Link.

#### Generating assemblies and computational validations

**Verkko assembly.** Verkko[18] v1.1 was run with the following parameters: `--slurm -d out_dir --graphaligner /path/to/GraphAligner --mbg /path/to/MBG --hifi-coverage ${X} --hifi /path/to/hifi/*.fq.gz --nano /path/to/nano/*.fq.gz`, where `${X}` differs per sample (example value: 30), set to approximately half of the HiFi coverage in Table S4. Note that the HiFi reads called with DeepConsensus were used, which meant 3/11 SMRTcells for Gorilla were not included. The following parameters were added for the two trio samples (gorilla & bonobo) to specify the Meryl databases for haplotype-specific, homopolymer-compressed kmers (k=30): `--hap-kmers maternal.compressed.k30.meryl paternal.compressed.k30.meryl trio`. For the four non-trio samples (chimpanzee, orangutans, and siamang), the Rukki (path walking) step was run manually with pseudo-haplotype-specific markers from the Hi-C data (see the Supplemental Methods' section 'Hi-C phasing'). The gorilla and bonobo haplotype-specific, homopolymer-compressed *k*-mers were generated with Meryl v1.3[19] and Merqury v1.3[19] using the following commands:

```
meryl count compress k=30 output mat.compressed.k30.meryl mat.fq.gz

meryl count compress k=30 output pat.compressed.k30.meryl pat.fq.gz

hapmers.sh mat.compressed.k30.meryl pat.compressed.k30.meryl
```

**Assembly and alignment visualization.** The assemblies were visualized and tangles were inspected using Bandage[4] v0.8.1 or BandageNG (https://github.com/asl/BandageNG) v2022.09. Chromosomes X and Y are often easy to identify because they are largely homozygous (i.e., long stretches without bubbles) and one end of each is joined at the PAR. Identification is especially striking when the graph is colored using trio markers. Alignments were visualized with IGV[18] v2.15.4.

**Assumed genome sizes.** Genome sizes were calculated from C-values taken from the Animal Genome Size Database (Gregory, T. R. Animal Genome Size Database. https://www.genomesize.com 2023) as found on Genomes on a Tree (GoaT; https://goat.genomehubs.org, [20]). Every picogram of DNA was assumed to be 978 Mb [21]. When no values were present for a given species, ancestral values were used. Median values were used when multiple values were available.

**Assembly evaluation with Merqury.** QV, k-mer completeness, and k-mer spectra plots were calculated with Merqury[19] v1.3. The following command was used for the trios: `$MERQURY/merqury.sh read-db.meryl hap1.meryl hap2.meryl hap1.fasta hap2.fasta out_prefix`. For those without parental data, the hap1.meryl and hap2.meryl positional parameters were omitted. Since no WGS Illumina data was generated as part of this study for the bonobo sample (mPanPan1), previously-generated WGS Illumina data (SRR11032812[22]) was used to create the “read-db.meryl”.

**Hi-C phasing.** Hi-C phasing has since been implemented in Verkko/Rukki, but it was not available at the time. Instead, the HiFi and HiC data was assembled with Hifiasm[23] v0.16.1-r375. Hifiasm was run with the following parameters: `hifiasm -o asm -t 48 --h1 /path/to/hic/*R1_001.fastq.gz --h2 /path/to/hic/*R2_002.fastq.gz /path/to/hifi/*.fastq.gz`. Markers were extracted by using Merqury[19] v1.3 with the commands:

```
cat asm.hic.hap1.p_ctg.gfa |awk '{if (match($1, "^S")) { print ">"$2; print $3}}' |fold -c > asm.hic.hap1.p_ctg.fasta

meryl k=21 memory=30 count compress asm.hic.hap1.p_ctg.fasta asm.hic.hap1.p_ctg.k21.meryl

cat asm.hic.hap2.p_ctg.gfa |awk '{if (match($1, "^S")) { print ">"$2; print $3}}' |fold -c > asm.hic.hap2.p_ctg.fasta

meryl k=21 memory=30 count compress asm.hic.hap2.p_ctg.fasta asm.hic.hap2.p_ctg.k21.meryl

$MERQURY/trio/hapmers.sh asm.hic.hap1.p_ctg.k21.meryl asm.hic.hap2.p_ctg.k21.meryl -no-filt
```

**MashMap alignments for chromosome orientation and chromosome naming.** MashMap[24] v2.0 was run with the following parameters: `-f map -k 16 --pi 95 -s 1000000`. Alignments were filtered to ignore alignments where the query (contig/scaffold) was covered by <50%. When present, the existing references were used to help orient the autosomal contigs. The Bornean orangutan assembly utilized the Sumatran orangutan reference, and the siamang assembly utilized the T2TCHM13v2.0 reference. Chromosome-level contigs and scaffolds were named, in addition to being oriented, based on these alignments; thus chromosome naming reflects the human-centric naming of the other great ape references. The next version of the genome assembly, including T2T autosomes, is expected to follow a different chromosome naming convention based on cytological standards (e.g., based on length of the chromosome) instead of on homology.

**Tangle resolution and patching with ONT data.** In some cases, the output contigs from Verkko had an overlap that was not merged automatically. In these cases, we confirmed the overlap size by examining the GraphAligner[25] output mapping the ONT reads to the graph. The contigs were then merged using the join_ctgs.py script from Nurk et al.[26]. In cases where separate contigs were output due to a complex tangle, the ONT alignments were consulted to select the correct path. In cases where no ONT read unambiguously spanned the assembly gap (e.g., due to a complex tangle or coverage dropout in the HiFi reads), a local assembly of relevant ONT reads was performed. Relevant ONT reads were identified based on overlap with the appropriate contig ends. The assembly was performed with Flye[27] v2.9-b1768 using the following parameters: `flye --asm-coverage 40 --nano-raw ont.subset.fq.gz`. The resulting assembled contig was inserted into the gap after trimming off the portion that overlapped the existing contigs surrounding the gap using the published pipeline[26].

**Alignments for assembly validation.** For short reads, alignments were performed with BWA-MEM2[28] v2.2.1. The index was built with this command: `bwa-mem2 index -p asm asm.fa.gz`. The alignments were performed with this command: `bwa-mem2 bwa -o output.sam asm reads1.fq.gz reads2.fq.gz`. All non-primary alignments were ignored. For long reads, Winnowmap2[29] v2.03 was used with the following parameters: `-x map-pb --MD -a -Y -L --eqx --cs -I 8G -W asm.repetitive.k15.tsv -o output.sam asm.fa.gz reads.fq.gz`. If ONT reads were used instead of PacBio HiFi reads, the `-x` parameter was changed to map-ont. The repetitive kmers file provided to -W was generated with Meryl[30] v1.03: `meryl count k=15 output asm.k15.meryl asm.fa.gz` and `meryl print greater-than distinct=0.9998 asm.k15.meryl > asm.repetitive.k15.tsv`. All non-primary alignments were ignored. For the assemblies with parental Illumina data, the alignments were further filtered using marker-assisted filtering using the scripts in https://github.com/arangrhie/T2T-Polish/tree/master/marker_assisted, as described in McCartney et al.[31].

#### Sequences gained

Each new assembly's X or Y was mapped to the corresponding X or Y from an older assembly (Table S7), using Winnowmap[32]. As per winnowmap recommendations, high-frequency 19-mers were identified first, using Meryl[19] with the following parameters:

```
meryl count k=19 output oldAssembly.meryldb oldAssembly.fa

meryl print greater-than distinct=0.9998 oldAssembly.meryldb >
oldAssembly.repeats

winnowmap -x asm20 -c --eqx -t 4 -W oldAssembly.repeats oldAssembly.fa
newAssembly.fa
```

For Table S7 any overlaps in the mapped intervals were collapsed. The length of the resulting intervals was summed, subtracted from the chromosome length, then compared to that length.

#### Non-B DNA annotations

To find non-B DNA forming motifs, we ran gfa (https://github.com/abcsFrederick/non-B_gfa). gfa creates six output files for the following non-B DNA types: A-phased repeats, short tandem repeats, direct repeats, mirror repeats, inverted repeats, and Z-DNA. We filtered gfa output files based on spacer length for inverted, direct, and mirror repeats (kept rows where spacer length ≤15). We generated G4 annotations using Quadron[33].

For each species, and separately for chromosomes X and Y, we created adjacent 100-kb windows across the chromosome using the 'bedtools makewindows' command[34]. For each species, and for chromosomes X and Y, we used 'bedtools coverage' command to count the number of overlaps between each 100-kb window and the non-B DNA specified in the concatenated gfa output file. Finally, for each 100-kb window, we counted the bases that have non-zero count, to avoid double counting the overlapping non-B DNA bases. This generates the non-B DNA density file that we used in the non-B DNA density plots and summary tables.

We ran simple and multiple Logistic Regression (LR) to evaluate the relationship between non-B DNA density and sequence novelty. A sequence in T2T assembly is considered novel if at least part of the sequence does not map to a previous, non-T2T assembly. For simple LR, for each species and for each non-B DNA type, the independent variable is the non-B DNA density for each 100-kb window in chromosomes X and Y and the dependent variable is a binary value: ‘0’ if the 100-kb window is not novel, and ‘1’ otherwise. In addition to seven non-B DNA densities, we also ran a simple LR for all non-B DNA density files combined. For multiple LR, for each species, the independent variables are the non-B DNA density of seven non-B DNA types; the dependent variable is determined identical to simple LR. We used R language’s glm (Generalized Linear Model) function to perform simple and multiple LR.

For repeat enrichment analysis, we used repeat annotations as described in Supplemental Methods’ section ‘Repeat and satellite annotations’. We ran ‘bedtools merge’ on the repeat bed files to combine overlapping intervals and avoid double counting them. We then ran 'bedtools intersect' to find the overlaps between the merged repeats and each of 7 non-B DNA annotations. For each species, and for each repeat type, we summed up the lengths of the intervals in the merged repeat files. For each species, for each repeat type, and for each non-B DNA type, we summed up the lengths of intersections between the merged repeat types and the non-B DNA types; this generated a table composed of 63 rows (number of repeat types) and 7 columns (number of non-B DNA types). We then divided the values in each row of this table by the corresponding value in the list to normalize the value to be between 0 and 1. Finally, we divided table cells by their corresponding average (for the X and Y chromosomes) non-B DNA density (of a particular type). This has allowed us to detect enrichment, i.e. cases when density in a certain cell was higher than the average non-B DNA density across the sex chromosomes.

#### Alignments

**Pairwise alignments with minimap.** To compute the percentage of sequences aligned and to study structural variants and segmental duplications, the pairwise alignment of the human chromosome X and Y was performed on the chromosome X and Y of the six ape species using minimap2.24[35] with options: `-m 10 -A 1 -B 2 -O2,12 -n2 -g 100 -cx asm20 -r 200,100000 --eqx -Y -s 1000`.

**Pairwise alignments with lastZ**. To support other analyses, lastz[36] was used to compute pairwise alignments of X and Y chromosomes. Five groups of alignments were computed—intra-species Y vs Y, intra-species X vs X, intra-species X vs Y, inter-species Y vs Y, and inter-species X vs X. The same steps and parameters were used for all groups, as follows. The assembly's X and Y chromosomes were softmasked according to the repeat annotations (Supplemental Methods' section 'Repeat and satellite annotations'). Alignment scoring parameters were set to match those indicated as appropriate for primates at http://genomewiki.ucsc.edu/index.php/Hg19_conservation_lastz_parameters. In detail, `--notransition` was used with this scoring matrix (equivalent to human-chimp.v2 at the page linked above):

```
gap_open_penalty   =   600 # O

gap_extend_penalty =   150 # E

hsp_threshold      =  3000 # K

gapped_threshold   =  4500 # L

x_drop             =   900 # X

y_drop             = 15000 # Y

     A    C    G    T

A   90 -330 -236 -356

C -330  100 -318 -236

G -236 -318  100 -330

T -356 -236 -330   90
```

**Multi-species alignments with CACTUS.** The 7-way sequence alignments were constructed from T2T chrX and separately chrY genomes of seven primates (bonobo, chimpanzee, human, gorilla, Bornean orangutan, Sumatran orangutan, and siamang) by CACTUS[37]. Pseudoautosomal regions were removed from both sex chromosome alignments. CACTUS also reconstructed six ancestral nodes (Anc_ALL, Anc_OGHBC, Anc_GHBC, Anc_O, Anc_HBC, Anc_BC).

#### Phylogenetic analysis

X and Y chromosome alignments separately generated using CACTUS[37] were converted to Multiple Alignment Format (MAF) using `hal2maf` (version 2.2)[38] with T2T-CHM13 set as the reference assembly, pseudoautosomal regions removed, and the following parameters: `--noAncestors --onlyOrthologs`. A custom `BioPython` script was used to extract 1-to-1 orthology blocks and convert the alignment format to FASTA, where each extracted alignment block contained a single sequence per species. X and Y maximum-likelihood phylogenies were inferred using IQTree (version 2.0.3)[39] with the best-fit substitution model estimated by `ModelFinder`[40] and node support estimated using 10,000 ultrafast bootstrap[41] replicates.

#### Substitution frequency analysis

To provide conservative estimates of substitution rates, we removed duplicates from the multi-species CACTUS alignments using MAFDUPLICATEFILTER from the MAFTOOLS suite[42]. This step replaces multiple sequences from the same species by the sequence closest to the consensus of an alignment block.To obtain conservative estimates, we only retained alignment blocks where all seven species were present. This step was performed using maf_filter_to_species_set.

We ran PHYLOFIT[43] on our filtered alignment using REV, with the following settings, separately for X and Y:

```
phyloFit --seed=35707 --EM --subst-mod REV --nrates 4 --tree
"(((((bonobo,chimp),human),gorilla),(sorang,borang)),gibbon)”
7species.multifasta --out-root 7species.phyloFit
```

We assessed the significance of the difference in branch lengths as follows. We counted the number of gap-free columns in the CACTUS alignments of chromosome Y (or X). We then estimated the number of substitutions along each lineage by multiplying the sum of the branch lengths along each lineage by the number of alignment columns. These were then used as $n_1$ and $n_2$, with $n=n_1+n_2$ in the following test described in Moorjani et al. (2016)[44] to compute an estimate of the chi-squared statistic.

$$\chi^2 \approx 2\left[n_1 log\left(\frac{n_1}{0.5n}\right) + \left(n - n_1\right) log\left(\frac{n-n_1}{0.5n}\right)\right]$$

#### X and Y substitution spectrum analysis

We extracted all the no-gap triple-nucleotide sequences from the 13-way (i.e. including reconstructed ancestors) CACTUS sequence alignments of chrX and chrY. The 12 branches are denoted as ALL_OGHBC (branch between Anc_ALL and Anc_OGHBC), ALL_gibbon (branch between Anc_ALL and gibbon), OGHBC_GHBC (branch between Anc_OGHBC and Anc_GHBC), OGHBC_O (branch between Anc_OGHBC and Anc_O), GHBC_HBC (branch between Anc_GHBC and Anc_HBC), GHBC_gorilla (branch between Anc_GHBC and gorilla), HBC_BC (branch between Anc_HBC and Anc_BC), HBC_human (branch between Anc_HBC and human), BC_bonobo (branch between Anc_BC and bonobo), BC_chimp (branch between Anc_BC and chimp), O_sorang (branch between Anc_O and Sumatran orangutan) and O_borang (branch between Anc_O and Bornean orangutan). Since there were no outgroups used to reconstruct the anc_all node, we are less confident in this reconstructed node, and thus the ALL_OGHBC and ALL_gibbon branches were removed from downstream analysis.

To remove PARs in each sex chromosome, alignment blocks overlapping with PAR annotations in any of the seven species were excluded. Triple-nucleotide sequences with 5’ base identical among 13 sequences and 3’ base identical among 13 sequences were used for downstream substitution spectrum analysis. For each branch, 192 types of triple-nucleotide substitutions were counted. Then by merging G with C as well as A with T, the counts data were consolidated into 96 types of triple-nucleotide substitutions. 96 types of triple-nucleotide substitutions were grouped into six types based on the middle base substitutions (C>A, C>G, C>T, T>A, T>C and T>G). To compare the distribution of the six substitution types between chrX and chrY, we applied a *t*-test to the proportions of each substitution type per branch in each group (chrX vs. chrY). Bonferroni correction was applied.

#### Gene annotations

**At the NCBI.** Almost all coding genes (96.7% for *S. syndactylus* to 98.08% for *P. paniscus*) were fully supported by alignments over more than 95% of their length. The completeness of the gene sets was estimated to be 93.92% (*S. syndactylus*) to 99.13% (G. gorilla) by BUSCO[45] version 4.0.2 run in -protein’ mode using the primates_odb10 marker set. See Table S32 for annotation statistics.

**At the UCSC.** The cactus(v2.6.0)[37]

(https://github.com/ComparativeGenomicsToolkit/cactus/releases/tag/v2.6.0) command for generating the alignments between the primate and human assemblies for the following input: `(((GCF_029281585.1:0.00993,((GCF_028858775.1:0.00272,GCF_029289425.1:0.00269):0.00415,(hs1:0.00025,hg38:0.00025):0.00619):0.00046):0.00509,(GCF_028885655.1:0.000945412,GCF_028885625.1:0.000915022):0.01864):0.00107106,GCF_028878055.1:0.00990798);`

For more detailed information see: http://public.gi.ucsc.edu/~hickey/hubs/hub-8-t2t-apes-2023v1/8-t2t-apes-2023v1.README.md RNA-Seq reads were aligned using minimap2 [35] using the following command: `minimap2 -a -x sr --sam-hit-only --secondary=no --eqx -t 4 mmdb/0.mmi rnaseq_data/0_0.fasta` Iso-Seq reads were aligned using minimap2[35] using the following command: `minimap2 -ax splice:hq -uf --sam-hit-only --secondary=no --eqx -t 4 mmdb/0.mmi isoseq_data/0_0.fasta` CAT [46] (commit: 5889b03380d92455b909c1ca0535fd590abbbe54) (https://github.com/ComparativeGenomicsToolkit/Comparative-Annotation-Toolkit) was run using the following command: `luigi --module cat RunCat --hal=8-t2t-apes-2023v1.hal --ref-genome=hs1 -- workers=10 --config=t2t.augustus.wholeGenome.chm13.config --work-dir chm13_2023v1/cat_work --out-dir chm13_2023v1/cat_output --local-scheduler --binary-mode local --augustus --augustus-pb --augustus-cgp --maxCores 45 --assembly-hub >& log_chm13_2023v1_CAT.txt` using the CHM13v2 annotation from UCSC GENCODEv35 CAT/Liftoff v2 as input along with this config file:

```
[ANNOTATION]
hs1 = chm13v2.gff3
[BAM]
GCF_029281585.1 = rnaseq/gorilla_gorilla/gorillaGorilla.final.wholeGenome.bam
GCF_028858775.1 = rnaseq/pan_troglodytes/panTroglodytes.final.wholeGenome.bam
GCF_029289425.1 = rnaseq/pan_paniscus/panPaniscus.final.wholeGenome.bam
GCF_028885655.1 = rnaseq/pongo_abelii/pongoAbelii.final.wholeGenome.bam
GCF_028885625.1 = rnaseq/pongo_pygmaeus/pongoPygmaeus.final.wholeGenome.bam
GCF_028878055.1 =
rnaseq/symphalangus_syndactylus/symphalangusSyndactylus.final.wholeGenome.bam
[ISO_SEQ_BAM]
GCF_029281585.1 = isoseqs/gorilla_gorilla/gorillaGorilla.final.wholeGenome.bam
GCF_028858775.1 = isoseqs/pan_troglodytes/panTroglodytes.final.wholeGenome.bam
GCF_029289425.1 = isoseqs/pan_paniscus/panPaniscus.final.wholeGenome.bam
GCF_028885655.1 = isoseqs/pongo_abelii/pongoAbelii.final.wholeGenome.bam
GCF_028885625.1 = isoseqs/pongo_pygmaeus/pongoPygmaeus.final.wholeGenome.bam
```

The liftoff[47] annotations were created using the following example command using version 1.6.3 (https://github.com/agshumate/Liftoff): `liftoff primate_x.fasta hs1.fa -sc 0.95 -g chm13v2.gff3 -polish` using the following reference annotations (1) UCSC GENCODEv35 CAT/Liftoff v2 and (2) NCBI RefSeqv110 To create the CAT/Liftoff annotation, we complemented the CAT result with missed GENCODE genes and putative additional paralogs from the Liftoff annotation using only predictions that did not overlap any CAT annotations.

#### Repeat and satellite annotations

**Identification and annotation of known repeat loci.** To identify canonical and novel repeats on chromosomes X and Y, we utilized the pipeline previously described in Hoyt et. al. 2022 [48]. Briefly, a RepeatMasker (v4.1.2-p1) run (RM1) was completed on each chromosome with a combined library of Dfam 3.6 and Repbase (v20181026) repeat sequences using sensitive settings (`-s`), the RepeatMasker compatible NCBI BLAST search engine RMBlast (`-e ncbi`), and the species tag of each respective ape (`-species` [one of: chimpanzee, bonobo, gorilla gorilla, bornean orangutan, sumatran orangutan, or gibbon]). Identified repetitive loci were used to generate hardmasked X and Y chromosomes, and a subsequent RepeatMasker run (RM2) was completed on each with a custom file containing 68 repeat models first identified in the analysis of T2T-CHM13 using the `-lib` flag.

The resulting repeat annotations from the two successive rounds of RepeatMasker were compiled into a single annotation by filtering out any repeat loci obtained from RM2 (T2T-CHM13 derived loci) with a Smith-Waterman score of 250 or less, eliminating repeat loci derived from RM1 that overlap with those remaining from RM2, and concatenating the filtered loci, hereafter referred to as 'RM-Merge-1'. RM-Merge-1 repeat loci were used to hardmask the sex chromosomes prior to repeat modeling to identify novel repeat structures.

**Satellite repeat modeling and curation.** Satellites and tandem repeats were identified in the above RepeatMasker analysis using both sequence similarity to known satellite sequences in existing repeat databases and the incorporated Tandem Repeats Finder (TRF) (trf409.linux64) screening. We used additional TRF screening (v4.09) and ULTRA (v1.0) with a periodicity of 1000 (`-p 1000`) on each ape sex chromosome to identify tandemly repeated sequences missed in the RM analysis. To curate satellite annotations, we identified ≥5-kb gaps in genomic features using bedtools (v2.29.0) by subtracting RM-Merge-1 and gene loci from the chromosome sequences and filtering based on length. Tandem repeats found in the resulting annotation gaps were manually curated using TRF and ULTRA outputs to identify consensus monomer sequences. Nomenclature for previously unknown satellites followed Hoyt et al, 2022[48] (in this case, moons/satellites of Neptune and Uranus).

**Final repeat annotation and compilation.** In order to produce a final set of repeat annotations including newly identified repeat models for chromosomes X and Y, a final RepeatMasker run (RM3) was completed using satellites curated above. Additionally, 17 variants of pCht/StSat, derived from Cechova et al.[49], were added to the database. Of the satellites identified, 23 were searched via RepeatMasker using the `-lib` flag. 19 loci were composed of consensi that were too AT-rich, variable, or lacked enough complexity for RepeatMasker to accurately annotate; monomers of 18 loci were annotated manually by independent TRF validation, and 1 was annotated regionally as AT-rich. The repeat compilation pipeline described in Hoyt et al., 2022 [48] was performed a second time to compile RM-Merge-1 and RM3 output resulting in RM-Merge-2. The RepeatMasker utility script buildSummary.pl (available at https://github.com/rmhubley/RepeatMasker) was used to summarize overall repeat annotations.

A note of caution—we discovered that prior taxonomic labeling of repeat library entries for repeats once considered lineage-specific (e.g. PtERV with a current taxonomic label in the repeat library of *Pan troglodytes*, therefore missed in searches of the gorilla and bonobo genomes) can lead to their exclusion from repeat annotations in a comparative framework due to incorrect lineage specification within the database.

**Satellite and composite repeat identification.** To identify larger tandemly arrayed structures, chromosomes X and Y were split into 200-kb windows and self alignment plots were generated using Minidot (v2016, available at https://github.com/thackl/minidot). The presence of large (>20-kb) regions with tandemly repeated sequence patterns, as described in Hoyt et al., 2022[48] as a tandemly repeating unit consisting of two or more repeat subunits, were visually identified and curated using IGV.

Each composite unit consensus sequence was generated by first extracting an approximate single composite unit within a predicted composite locus. Using this sequence, we used BLA [50] to identify and collect the fasta sequence of all unit-length and near-unit length insertions. The composite was then polished using a previously published strategy and associated scripts[51]. Briefly, the insertions were aligned to the approximate consensus sequence via the alignAndCallConsensus.pl script (available at https://github.com/Dfam-consortium/RepeatModeler). The script was run until the local alignments stabilized and the overall score of the total alignment could no longer be improved. The command line argument used for each composite was: `$ alignAndCallConsensus.pl -c <consensus.fa> -e <insertions.fa> -ma 14 -re 10 -html`, where `-ma 14` indicates a matrix based on 14% sequence divergence (the lowest % divergence available as part of the script), `-re 10` indicates the maximum number (in this case, 10) of alignment iterations to attempt in order to produce a stable alignment, and `-html` indicates that a user-friendly

HTML multiple sequence alignment (MSA) should be generated as part of the output. Each alignment was visually assessed, and minor adjustments were made according to the recommendations based on [51]. An example of a minor adjustment would be to trim the consensus sequence if minimal (less than three) sequences extended to the edges of the sequence.

Custom BLAST databases were generated from each assembly in order to search for individual composite units using their consensi, thereby determining the composite unit copy number across primate X and Y chromosomes. Due to the variability seen in the composite units between species, the BLASTN search required at least a 40% match to the composite consensus to be considered. All instances were manually checked for appropriate composite unit structure and categorized as ‘full length’ if their length exceeded 75% of the consensus length or as ‘fragmented’ if their length was less than that. These results are reported in Tables S21 and S22. Composite tracks were generated to highlight all composite unit loci (full length and fragmented). Each species-specific track includes additional information about the composite unit, including composite name, length of the unit, percent divergence from the consensus sequence, percent of the consensus sequence length, and its status as either ‘full length’ or ‘fragmented’.

**Identification of lineage-specific insertions and repetitive elements.** Lineage-specific (LS) insertions were ascertained by utilizing the hierarchical alignment (HAL) file produced as a result of the alignment of the seven primate X and Y chromosomes by CACTUS, and the associated halTools package. For each species, the halAlignExtract (with `-complement` option) output was used to determine unaligned regions in the alignment, based upon the parent genome in the HAL file. A subsequent analysis merged fragments separated by ≤20 bp, utilized a combination of TRF and ULTRA to remove insertions consisting of mostly simple and low complexity repeats, and removed insertions <70 bp and >15,000 bp. To identify the content of the LS insertions, a RepeatMasker analysis using a sequential approach was used. The first round of RepeatMasker consisted of identifying canonical repeats. The second round was used to identify the remaining insertions with a custom library based on new repeats found as part of this study.

**Visualization.** For generation of repeat density heatmaps across primate and human X and Y chromosomes (Ext. Data Fig. 2), BEDTools [52] coverage was used to calculate base-pair coverage across 100-kb windows per repeat class or category followed by subsequent visualization with Rideogram v0.2.2 [53].

#### Centromeric satellite analysis

To perform the analysis of centromeres, we created various annotations and built custom annotation tracks in UCSC Human Genome Browser as follows:

1. Suprachromosomal Family (SF)-tracks, done as described in Altemose et al.[54], show SF-classes of monomers for all AS arrays using score/length ratio threshold 0.7, which is good for viewing alpha satellite (AS) in the arms, as it filters out unreliable very short and low-score hits. Coverage in the centromeres may not be continuous in siamang where many full-length monomers give low scores with the human-based HMM profiles.
2. SF-tracks with reduced threshold 0.3 and with no threshold were built to improve coverage in siamang at the expense of seeing more noise in the arms.
3. AS-strand track, same as SF-track, but colored to show only the orientation of AS. Blue color indicates AS on direct strand and red on reverse strand.
4. HOR-track which shows what identified high order repeat (HOR) of a given species each monomer belongs to. These tracks were built using species-specific HMMER-based tools especially developed for this paper. The HORs identified are listed in Table S29, the HMMs built to identify each monomer of the HORs are available on https://github.com/fedorrik/apeXY_hmm. Methods of HOR evaluation and tool construction were described previously[54].
5. StV-tracks showing structural variation (altered monomer order) in HORs were built for all active and some inactive AS arrays in all centromeres. Note that StV annotations are different from SV annotations used in this project. The former only involves alpha satellites and shows only the unusual presence or absence of whole alpha-satellite monomers in the context of our HOR annotation. Both smaller (submonomeric) and larger-than-HOR indels are not covered. The summary statistics for these tracks are provided in Table S30. The script for building the StV-track is available on https://github.com/fedorrik/stv.

6. Functional CENP-B sites were visualized by running a short match search with the sequence YTTCGTTGGAARCGGGA.
7. For centromere analysis, we also used repeat annotation (see Supplemental Methods' section 'Repeat and satellite annotations'), methylation (see Supplemental Methods' section 'CpG methylation') and segmental duplication (see Supplemental Methods' section 'Segmental duplications') tracks built for this paper.

SF-tracks were built and color-coded as described[54] using human-based annotation tools, because SF classification covers all alpha satellites in all primates, and the SF-tools based on human HMMs work in all primates if the detection threshold is adjusted appropriately. To the contrary, HOR annotation tools have to be species-specific or at least genus-specific, as different apes have different HOR sets and human-based HOR classification tools would not work properly on non-human centromeres. However, the closely related species, such as chimpanzee and bonobo, and Sumatran and Bornean orangutans possessed the same HORs and did not require species-specific tools. The tools for non-human primate HORs were built the same way as was previously described for human HORs [54]. All annotations are publicly available at the following URLs: https://genome.ucsc.edu/s/fedorrik/primatesX (chromosomes X) and https://genome.ucsc.edu/s/fedorrik/primatesY (chromosomes Y).

The SF annotation coverage in siamang is sometimes not continuous, as some monomers are not annotated due to significant divergence of gibbon alpha satellite monomers from their progenitor Ga class monomers. However, most monomers in both centromeric and sub-telomeric arrays are identified as Ga, which indicates SF4. In orangutan centromeres, most monomers are identified as R1 and R2 which indicates SF5. In chimpanzee and human autosomes and X chromosomes, active arrays are formed by J1 and J2 (SF1), D1, FD and D2 (SF2) and W1-W5 (SF3) monomers (the new families). The detailed notes on the coverage and contamination issues in the tracks are provided in Note S7.

More detailed analysis of active HOR arrays in pairs of closely related *Pan* and *Pongo* species was performed as follows. We used our HOR-tracks and StV (structural variant)-tracks of the assemblies to extract full-length HORs from the active array. Sometimes the major StVs with duplications were also extracted, duplications removed and the HORs converted to full-length. Such sets were added to the full-length sets. 100-300 HORs for each species were randomly selected, combined with such a sample for another twin species, aligned and used to build minimum evolution phylogenetic tree (HOR tree). The branches on this tree were treated as HORhaps (HOR haplotypes[54]).

The sequences for each HORhap (a tree branch) were collected, re-aligned and used as HMMs for HMMER HORhap classification tool built as described in [54]). The same MSAs (multiple sequence alignment) were used to derive HORhap consensus sequence (simple majority), and average divergence between HORs in MSA was determined as an estimate of intra-array divergence.

Minimum evolution trees of the HORhap consensus sequences were built and analyzed to determine which HORhap was more derived (younger) or less derived (older), and which was near equi-distant to both species, which was considered to be an approximation to the major HOR in the common ancestor of the 2 species. Active HORhaps were usually more derived in these trees.

The HORhap classification tool was used to build the HORhap annotation of the assemblies and produce the HORhap tracks. These were examined manually to verify that at least some of the resulting HORhaps were "well regionalized" which meant that they formed arrays composed almost exclusively of one HORhap. If two or more HORhaps were interspersed with each other and were closely related (sat in the neighboring branches in the HOR-tree), they would be merged and treated as one HORhap. If they were not closely related, they were treated as different HORhaps which jointly make the same array. Sometimes there were regions made solely by one HORhap and others made by the same HORhap jointly with some other one. This was duly noted and such regions were interpreted as closely related but different.

Using all the above information we picked up the consensus sequences for the active HORhaps (the ones which hosted CDRs) for each chromosome for both species, compared them and determined the number of differences per HOR and per monomer. The differences in the consensus sequences listed were further analyzed to check the percentage of each base at a given position to make sure the differences do not result just from slight variation of nucleotide frequencies at this position.

#### Classifications into PARs, ancestral, ampliconic, and X-transposed regions

**Pseudoautosomal regions (PARs).** In order to find initial candidate PAR regions, we utilized SEDEF (v1.1) [55] to identify homologous segments in genome assemblies in which common repeats were masked using Tandem Repeats Finder (TRF) (v.4.0.9)[56], RepeatMasker (v4.1.2-p1) [57], and Windowmasker (v2.2.22) [58]. Satellite DNA content was annotated using RepeatMasker. pCht repeats, a 32-bp satellite repetitive sequence found in non-human primates including gorillas, bonobos, and chimpanzees, were identified using BLAST [59]. Genomic regions with high sequence identity (>99% matched region and >98% including indels and at least 100 kb) between X and Y chromosomes were annotated as initial candidate PAR intervals. The initial candidate PAR intervals were then manually adjusted by considering the locations of long tandem repeat arrays and ending locations of high-identity alignments. Dotplots of lastz [36] alignment in the candidate regions guided the manual process. Tandem Repeats Finder [56] and Noise Canceling Repeat Finder [60] were used to identify the endpoints of tandem repeats, and intervals of these repeats were excluded. Lastz alignments of the remainder of the candidate interval were manually trimmed to remove lower-identity tails.

**Overview of the workflow for non-PAR sequence class annotations.** We annotated the X and Y chromosomes of the studied species into sequence classes, following the annotations designed by Skaletsky et al.[12], with modifications. First, we created a satellite repeat track containing consecutive stretches of predominantly satellite sequences. This was achieved by merging neighboring RepeatMasker annotations (bedtools `merge -d 1000`). We only kept regions larger than 0.25 Mb that did not overlap PARs. Next, we identified potentially ampliconic regions. This was done by combining palindrome-forming regions as discovered by PALINDROVER version 20230615 (available at https://github.com/makovalab-psu/T2T_primate_XY/tree/main/src) and regions with high intrachromosomal identity. For intrachromosomal similarity, each coordinate of a 5-kb window on the Y chromosome with 2-kb step that mapped to another region with the minimal identity of 50% using blastn version 2.5.0+ was kept, and considered a hit. In order to analyze the matches to ampliconic regions, and not to the repetitive parts of the Y chromosome, we hardmasked the sequences beforehand. All RepeatMasker annotations matching the keywords SAT, GAP, or LINE were hardmasked, as well as subterminal satellites for chimpanzee, bonobo, and gorilla, and HSAT in human. For mapping, we used blastn version BLAST 2.5.0+ with the parameters `-perc_identity 50` and the same hardmasked version of the Y chromosome as a reference. In order to exclude self-alignments (each window mapping back to its origin), we excluded all alignments whose start or end coordinates fell within the coordinates of the window from which they were generated. The coordinates of hits within close distance were kept (`-d 100000`) and filtered similarly as for the repeats, requiring a length of at least 90 kb in order to filter out spurious hits to repeats. The coordinates identified by PALINDROVER and those identified by the intrachromosomal similarity analysis were merged. We shortened or split ampliconic regions when appropriate in order not to overlap with PARs and satellite repeat track (using the bedtools command `subtract`). Finally, for the Y chromosome, regions annotated as none of the above-mentioned classes were annotated as potentially ancestral (X-degenerate), and confirmed as such if they intersected with the coordinates of any ancestral (X-degenerate) genes. The remainder of the non-PAR sequence for the X chromosome was then designed as ‘ancestral’, and, for the Y chromosome as class ‘other’ in the first pass, and, in cases where subregions of this class overlapped with ampliconic genes, as ‘ampliconic’. For both X and Y chromosomes, to improve the continuity of the annotations, in the second pass, each ‘other’ region that was nested within two regions of the same annotation was also annotated as such.

Please see Note 4 to find information about our search of X-transposed regions.

Please also note that many figures include a “Sequence class” track to help contextualize other data with the regions of the chromosomes. In those figures an additional alpha satellite “class” is included, which can be a helpful proxy for the centromeres and surrounding satellites. The location of alpha satellites were extracted from the RepeatMasker output.

#### Palindromes

**Palindromes Detection.** Palindromes were derived from lastz [36] chromosome self-alignments (part of the pairwise alignments). Only alignments to the reverse strand and above the main diagonal were considered; moreover any portion of an alignment extending below the main diagonal was discarded. Following [12], the remaining alignments were considered as candidates if they had sequence identity ≥ 98%, length ≥8 kb, and spacer ≤500 kb. Candidates were then subjected to a blacklist filter to reject those in satellite or certain repeat regions. Specifically, intervals annotated as Satellite, any class beginning with 'Satellite/', 'Low_complexity', or 'Simple_repeat' were collected from the repeat annotations (see Supplementary Methods' section 'Classifications into PARs, ancestral, ampliconic, and X-transposed regions'). Any candidate palindrome with at least 80% of its bases covered by such an interval was rejected. The resulting software to detect palindromes was called PALINDROVER and is available on GitHub.

**Finding pairwise sharing of palindromes between species**. lastz [36] alignment was performed for all possible pairwise palindrome combinations (considering arms only) using the human-chimp.v2 scoring parameters appropriate for primates and `--notransition`, as we describe elsewhere. Alignments with identity <85%, gaps >5%, <500 matched bases, or covering <40% of either arm were discarded. Pairs of palindromes with non-discarded alignments were considered to be shared between species (Table S16).

**Clustering of homologous palindromes.** Palindromes were grouped by overlap—if an arm of a palindrome A overlapped an arm of palindrome B, A and B were considered part of the same group. Alignments (from the previous paragraph) were then used without respect to groups—if any arm of a palindrome in group C aligned to any arm in group D, this was counted as one aligning palindrome. As a result, we obtained clusters of homologous palindromes across species (Table S47).

#### Segmental duplications (SDs)

The analysis of SD content in humans and non-human primates was performed using a previously described method [61]. Briefly, we identified homologous segments using SEDEF (v1.1)[55] in genome assemblies in which common repeats were masked using Tandem Repeats Finder (TRF) (v.4.0.9)[56], RepeatMasker[57], and Windowmasker (v2.2.22)[58]. Satellite DNA content was annotated using RepeatMasker. pCht repeats, a 32-bp satellite repetitive sequence found in non-human primates including gorillas, bonobos, and chimpanzees, were identified using BLAST[59]. Among the SDs identified by SEDEF, only duplicate regions with sequence identity >90% and a minimum length of 1 kb were kept. We also excluded SDs composed of >70% satellite sequences. Lineage-specific SDs of a species A were defined by comparing the putative homologous SD loci of the remaining ape species to the species A assembly. This was repeated by taking each species as the reference and comparing the homologous SDs of the remaining species. The projecting of SD loci was performed by aligning 10-kb SD flanking sequence of ape species to the reference assembly via minimap2.24[35].

#### Structural variants

Ape SVs on the sex chromosomes were identified using CHM13v2.0 as the reference. The pairwise alignment of the human chromosome X and Y was performed on the chromosome X and Y of the six ape species using minimap2.24[35] with options `-m 10 -A 1 -B 2 -O2,12 -n2 -g 100 -cx asm20 -r 200,100000 --eqx -Y -s 1000`. PAV [62] was used to discover structural variants (insertion, deletion and inversion) 50 bp to 300 kb in length. For larger variants, we applied Saffire SV variant calling pipeline (https://github.com/wharvey31/saffire_sv). Rustybam v0.1.29 (https://github.com/mrvollger/rustybam) post-processed alignments (orienting contig alignments, trimming overlaps, filtering out alignments with fewer than 1 Mb), and a series of python scripts retrieved larger variants from alignment files (https://github.com/wharvey31/saffire_sv). Specifically, rustybam was used to break cigar strings which contained insertions or deletions of >30 kb to create a broken alignment file. After creation of the oriented and broken alignment files, these were compared against each other to identify simple insertions and deletions in both contig and reference space. Inversions were called on the negative strand of alignments, and combined with neighboring 'transpositions' to identify nested inversions. Transpositions were classified based on alignments that mapped >30 kb away from their expected alignment coordinates within neighboring syntenic blocks. Gaps were defined as the complement of the oriented alignment file. Gaps that contained flanking alignments originating from the same contig were examined for their length differential between contig and reference space, and differentials of ≥30 kb were classified as either inserted or deleted. Duplications were called based on overlapping alignments. We also recovered smaller PAV variants spanning within the filtered-out deletion calls from PAV (>300 kb). On the other hand, smaller variants overlapping the Saffire SV large deletions were filtered out using bedtools subtract (v2.29.2). For inversion variants, PAV and Saffire SV inversion calls with reciprocal coverage >0.95 were merged. The merged PAV and Saffire SV deletions and inversions defined the final deletion and inversion callset. The insertions from PAV and duplications from the Saffire SV relative to human defined the final insertion callset. We defined human-specific structural variants relative to CHM13v2.0 Y by intersecting the variant loci of six ape species using bedtools (v2.29.2). Overlapping deletions in the six ape species relative to human reference chromosomes were classified as putative human-specific insertions, whereas the overlapping insertions were considered as putative human-specific deletions. The phylogenetic branch of origin was predicted using maximum parsimony. Variants shared by bonobo and chimpanzee and present only in these two species were considered to originate in the *Pan* lineage. As a limitation of this analysis, the SVs for branches including ancestors of the reference species, i.e. human ancestors (i.e. human-chimp-bonobo, human-chimp-bonobo-gorilla, and human-chimp-bonobo-gorilla-orangutan common ancestors) were not computed due to the lack of reference variant call set for human.

#### rDNA array validations

**Chromosome spreads, Fluorescent In-Situ Hybridization (FISH), and immuno-FISH.** For the preparation of chromosome spreads, cells were blocked in mitosis by the addition of Karyomax colcemid solution (0.1 μg/ml, Life Technologies) for 6-7h and collected by trypsinization. Collected cells were incubated in hypotonic 0.4% KCl solution for 12 min and prefixed by addition of methanol:acetic acid (3:1) fixative solution (1% total volume). Pre-fixed cells were collected by centrifugation and then fixed in Methanol:Acetic acid (3:1). Spreads were dropped on a glass slide and incubated at 65°C overnight. Before hybridization, slides were treated with 0.1mg/ml RNAse A (Qiagen) in 2×SSC for 45 minutes at 37°C and dehydrated in a 70%, 80%, and 100% ethanol series for 2 min each. Slides were denatured in 70% formamide/2×SSC solution pre-heated to 72°C for 1.5 min. Denaturation was stopped by immersing slides in 70%, 80%, and 100% ethanol series chilled to -20°C. Labeled DNA probes were denatured separately in a hybridization buffer by heating to 80°C for 10 min before applying to denatured slides. Fluorescently labeled probes for human rDNA (BAC clone RP11- 450E20) and *SRY* (BAC RP11400O10) were obtained from Empire Genomics (Williamsville, NY). Specimens were hybridized to the probe under a glass coverslip or HybriSlip hybridization cover (GRACE Biolabs) sealed with the rubber cement or Cytobond (SciGene) in a humidified chamber at 37°C for 48-72 hrs. After hybridization, slides were washed in 50% formamide/2×SSC 3 times for 5 min per wash at 45°C, then in 1×SSC solution at 45°C for 5 min twice and at room temperature once. For immuno-FISH, slides labeled by FISH were subjected to antigen unmasking in hot (65°C) Citrate buffer, pH 6.0, for 1 hour before processing for immunofluorescence. Slides were blocked with 5% bovine serum albumin (BSA) in PBS/0.5% Triton X-100. Primary antibody (rabbit polyclonal anti-UBF, Novus Biologicals, cat.# NBP1-82545) and secondary antibody (goat anti-rabbit Alexa Fluor 647, Thermo) were diluted in 2.5% (weight/volume) BSA/PBS/0.5% Triton X-100. Specimens were incubated with primary antibody at a minimum overnight, washed 3 times for 5 minutes, incubated with secondary antibody for 2-4 hours and washed again 3 times for 5 min. All washes were performed with PBS/0.5% Triton X-100. Slides were mounted in Vectashield containing DAPI (Vector Laboratories). Z-stack images were acquired on the Nikon TiE microscope equipped with 100× objective NA 1.45, Yokogawa CSU-W1 spinning disk, and Flash 4.0 sCMOS camera. Image processing was performed in FIJI.

**Estimating rDNA copy number and activity from FISH and immuno-FISH images.** Sum intensity projections of Z-planes were generated, and individual rDNA arrays were segmented based on threshold applied to the entire image. The fluorescence intensity of the regions of the same chromosomes that did not contain the rDNA was used to subtract the local background. The background-subtracted integrated intensity was measured for each array. The sum of all integrated intensities of all rDNA loci represented the total amount of rDNA per cell, and the fraction of this total signal was calculated for each array. The total rDNA copy number was estimated from Illumina sequencing data (see section ‘Estimating rDNA copy number from k-mer coverage' below). The fraction of the total rDNA fluorescence intensity was used as a proportion of the total rDNA copy number to determine the number of rDNA copies on the chrY chromosome in S. orangutan and gibbon. Similarly, the fraction of the total UBF fluorescence intensity was used to estimate the transcriptional activity of the chrY rDNA arrays.

**Estimating siamang Y chromosome rDNA copy number.** Two computational methods were used to confirm the rDNA copy number estimate from FISH. The first method was a basic coverage comparison. Using ribotin (https://github.com/maickrau/ribotin), we constructed a representative rDNA unit from the siamang chromosome Y assembly. Aligning all the HiFi reads from this region of the assembly to the representative rDNA sequence yields a median coverage of 410x, while the 100 kb leading up to the rDNA array on chromosome Y has a median coverage of 28x. This gives an estimated copy number of 14.6 units of rDNA, or roughly 14-15 units (410/28=14.6). The second method was based on a comparison to known rDNA copy numbers in the human CHM13 assembly. We explored rDNA coverage for rDNA clusters in the assembly graph from CHM13 chromosomes 14 and 22 and siamang chromosome Y. For each cluster, we found regions shared between all rDNA copies using ribotin. We compared the total coverage of the human rDNA regions (with known rDNA copy numbers of 16 and 21 units for chromosomes 14 and 22, respectively) with coverage for non-rDNA regions of corresponding chromosomes to estimate HiFi coverage depression $c_d$ for rDNA (caused by high frequency of GC repeats in homopolymer compressed space). These estimates were close for both CHM13 chromosomes 14 and 22 at 1.78 and 1.86, respectively, which averages to 1.82. We assumed that coverage bias in siamang rDNA is similar to the CHM13 dataset given a similar GC repeat content (34.1% in CHM13 chromosome 14, 33.9% in CHM13 chromosome 22, and 34.4% in siamang chromosome Y), as assessed by methods in T2T-Polish (https://github.com/arangrhie/T2T-Polish) [31]. Finding shared regions between all rDNA copies in siamang chromosome Y using ribotin, we estimated the total coverage of these regions to be 384.2×. Given that the coverage of non-rDNA chromosome Y regions is 39.5×, we have an estimated copy number of 17.7 copies, or roughly 17-18 rDNA units (384.2/39.5*1.82=17.7).

**Estimating rDNA copy number from k-mer coverage.** We used a k-mer based method to estimate the copy number of the rDNA, based on Nurk et al., 2022[26]. The 18S gene has relatively uniform coverage due to its high degree of conservation and more typical %GC (56%)[26], so 18S was used for copy number determination. The gorilla and Sumatran orangutan have published 45S rDNA references[63]. The bounds of the 18S subunit within these sequences were determined by mapping the first and last 31-mer from the human sequence to the primate one, which was successful with exact single k-mer matches. For the other species there was no reference, so the human 18S sequence was used. A set of 2-kb normalization windows were identified that are within 1% of the total GC content of the 18S reference. An additional step ensured that 31-mers from these windows do not occur elsewhere in the genome. 31-mers from the 18S and normalization set were counted in Illumina sequencing data using jellyfish[64], then counts were divided by their multiplicity. The median 31-mer count of the 18S was divided by that of the matched windows to yield a total copy number. A custom python script was used for mathematical operations. Script available at https://github.com/makovalab-psu/T2T_primate_XY/tree/main/45S_rDNA_CN.

Illumina sequencing data was published in the SRA (SRX21756818,SRX21758603,SRX21765246,SRX21765787,SRX21765788,SRX21765789) and pre-existing Illumina data was used for the bonobo (SRX7685076)[22]. Table S31A summarizes the output from the *k*-mer pipeline. Tables S30B and S30C summarize the estimates of copy number for siamang and S. orangutan, respectively.

#### Methylation

**CpG methylation calling.** In order to generate CpG methylation calls across the X and Y chromosomes, Meryl (v1.3)[19] was used to count k-mers and compute the 0.02% most frequent 15-mers in each ape draft diploid assembly:

```
meryl count k=15 output assembly.k15.meryl assembly.fa.gz

meryl print greater-than distinct=0.9998 assembly.k15.meryl >

repetitive-k15.tsv
```

ONT were re-basecalled because 5mC methylation information was not present for all flowcells. Similarly, PacBio HiFi reads were regenerated because not all HiFi reads had accompanying kinetics information necessary for getting 5mC methylation information. ONT reads were re-basecalled with Guppy v6.3.8 using model `dna_r9.4.1_450bps_modbases_5mc_cg_sup_prom.cfg`. When needed, HiFi reads were generated from the subreads using pbccs v6.4.0, and 5mC methylation information was calculated from the kinetics data for all read sets using Primrose v1.3.0:

```
ccs --hifi-kinetics --minLength 10 --maxLength 50000 --minPasses 3 --minSnr 2.5
--minPredictedAccuracy 0.99 movie.subreads.bam > hifi_reads.bam

primrose --keep-kinetics hifi_reads.bam > 5mC.hifi_reads.bam
```

ONT and PacBio reads were mapped to the corresponding draft diploid assemblies with Winnowmap (v2.03) [29] using the following parameters:

```
winnowmap -W repetitive-k15.tsv -a --cs --eqx --MD -x (one of: map-ont or
map-hifi) -k 15 -y <assembly> <reads.fq.gz> > output.sam
```

Samtools (v1.17) [65] was used to filter secondary alignments and unmapped reads prior to CpG track generation:

```
samtools view -bu -F 260 output.sam | samtools sort --write-index -o
output.bam##idx##output.bam.bai
```

Modbam2bed (v0.6.2) was used to summarize modified basecalls and generate a CpG methylation track viewable in IGV.

**Centromere Dip Region (CDR).** Previous studies have linked a dip in CpG methylation within an alpha satellite HOR array to the region of the array that defines the kinetochore assembly domain[66,67]. Following CpG methylation track generation from ONT and PacBio data, manual inspection of CpG density along HORs were used to define the CDR for each centromere.

**Methylation analysis.** Average methylation levels were calculated by taking all methylation levels within the PAR1, non-PAR X, non-PAR Y, and (when applicable) PAR2 regions. Average methylation across 100kp windows were taken from the CpG sites. Humans had CpGs removed if the difference in methylation levels between ONT and Pacbio methylation were greater than 0.05 or CpGs were in the Yq12 region (high frequency of incongruent methylation between ONT and Pacbio). The means were calculated along with mean standard error from bootstrapping for each region and the significance was tested using a Wilcoxon Rank-Sum test (Extended Data Fig. 3 and Fig. S14A). Similarly, custom R scripts were written (available at https://github.com/makovalab-psu/T2T_primate_XY/tree/main/methylation) to calculate average methylation levels at annotated genes and features including PARs, repeats, and sequence classes (Extended Data Fig. 3 and Fig. S14).

To plot the smoothed average methylation profiles of each species across transcription units, we generated, for every gene, and at every base position within 3-kb upstream, and downstream, of the transcription start site and transcription end site, respectively, the average methylation found in the 500-bp bin centered on each base position. For gene body regions, all CpG methylation coordinates within them were normalized to 12,000 units to be comparable to all other genes. 12,000 units were selected to depict gene body positions because gene length was on average 4 times the 3-kb upstream and downstream regions profiled in the plots. Smoothing was then carried out by calculating the average methylation found in the 500 unit position bins centered on each unit position. Subsequently, methylation was averaged across all genes of a species at every upstream/downstream base position and gene body unit position, yielding a single average methylation profile representative for that species (Fig. S14B-C).

Spearman's correlations between promoter methylation (average CpG methylation in 1000 bps upstream of TSS) and gene expression (counted from Illumina Next-seq 2000 reads using subread featureCounts, v2.0.6) were calculated for genes in the PAR1, non-PAR X, and non-PAR Y (Fig. S14D). Correlation was also tested for statistical significance ($H_0$: correlation = 0) using the Spearman's test. In Fig. S14D, correlations to the right of the solid red line (p<0.01) are considered nominally significant, with non-PAR X genes being highly significant compared to other regions in part due to larger numbers of genes (PAR1, N=15; non-PAR X, N=818; non-PAR Y, N=19).

#### Diversity

Samples collection and data processing. We collected 129 published, high-coverage genomic data sets from four projects [6869 70 71]. The samples consist of: 13 bonobos (*Pan paniscus*), 57 chimpanzee (18 *Pan troglodytes troglodytes*, 19 *Pan troglodytes schweinfurthii*, 9 *Pan troglodytes ellioti*, and 11 *Pan troglodytes verus*); 49 gorillas (1 *Gorilla gorilla diehli*, 9 *Gorilla beringei graueri*, 12 *Gorilla beringei beringei*, and 27 *Gorilla gorilla gorilla*); and 5 Sumatran orangutans (5 *Pongo abelii*) and 5 Bornean orangutans (5 *Pongo pygmaeus*). Detailed information on all samples is provided in Table S42. To analyze the performance of sequencing alignment and variant calling with previous vs. T2T reference genomes, we downloaded the previously published reference genome for each species (GCA_008122165.1, GCA_015021865.1, GCF_000258655.2, GCA_015021855.1, GCF_002880755.1, GCF_000001545.4, GCA_015021835.1). Because these reference genomes of gorillas, bonobos, and orangutans do not include chromosome Y, we integrated Y chromosome scaffolds from Cechova et al. [22] into the non-T2T reference genomes.

**Species mapping and variant calling.** For each subspecies, we executed the independent alignment and variant calling analysis with T2T reference and previous reference genomes following the approaches used for the human T2T analyses [72]. Briefly, the Illumina reads of each sample were aligned to the corresponding reference using BWA-MEM V0.7.17-r1188 [73]. Then we performed variant calling on sex chromosomes with GATK v4.4.0.0 HaplotypeCaller and joint genotyping with the GenotypeGVCFs tool[74]. Low-confident variants were removed using SelectVariants and variantFiltration based on the GATK's hard-filtering parameters (for SNPs: `-filter QD < 2.0 QUAL < 30.0 SOR > 3.0 FS > 60.0 MQ < 40.0`; For indels: `-filter QD < 2.0 QUAL < 30.0 FS > 200.0`). Variants with a genotype quality below 20 were further removed to enhance the accuracy of our variant call set. For all analyses, we assessed the mappability and mismatch rate with samtools v1.6 [65] using the samtools stats tool.

**Impact of masking PAR in variant calling.** We applied the masking strategy to improve the completeness of the variant callset, similar to the one used previously[75]. For each species T2T reference genome, we generated karyotype-specific references using PAR annotations. Specifically, the chrY-PAR was masked in the XY reference, while the whole chrY was masked in the XX reference. Then, reads from XX and XY samples were aligned to the corresponding masked reference genomes. The remaining variant calling and filtering procedures are explained in section 'Species mapping and variant calling'. The masked reference genomes, alignments and variant calls are available within the NHGRI AnVIL[76].

**Genetic diversity analysis.** We estimated the nucleotide diversity within different regions for each subspecies using VCFtools V0.1.16 [77]. The window size was 100 kb. For chromosome X, we calculated nucleotide diversity in the PARs, and the remaining regions which excluded PARs, satellites, and ampliconic regions. For chromosome Y, we assessed the diversity in ancestral regions, and the remaining regions which excluded amplicons, satellites, and PARs.

#### Y chromosome phylogeny and TMRCA calculations

Previously published short-read genome-wide sequencing data for male great ape samples (21 chimpanzees, 2 bonobos, and 11 gorillas; Table S42 [68–70]) were used for the construction and dating of intra-specific Y-chromosomal phylogeny. The variants were called as described in the previous section.

Only regions defined as ancestral in each of the species' Y-chromosomal assembly were used, followed by removal of indels, calls where ≥10 % of high-quality reads supported another allele and sites where >2 chimpanzees and >1 gorilla had missing genotypes, using VCFtools (v.0.1.16) [77]. No missing data were allowed for the bonobo samples. A total of 7,084,961 bp (including 26,587 SNVs) were left after filtering for chimpanzees, 6,929,306 bp (including 6,443 SNVs) for bonobos, and 6,807,422 bp (including 4,792 SNVs) for gorillas. The respective sequences from the *de novo* assembled Y assemblies for each of the species were included in the phylogeny construction and dating.

The Bayesian Markov chain Monte Carlo phylogenetics software BEAST v1.10.4 was used to estimate the time-to-most-recent common ancestor (TMRCA) for the nodes of interest [78]. In the absence of good mutation rate estimates for great ape male-specific regions of the Y chromosome (MSY), two different human MSY mutation rates were used–3.07(95% CI: 2.76-3.40)$\times10^{-8}$ single-nucleotide mutations/nucleotide/generation [79] and 0.76(95% CI: 0.67-0.86)$\times10^{-9}$ single-nucleotide mutations per bp per year[80]. These were scaled according to the male generation time estimates of 31 years for human, 24 years for chimpanzees and bonobos (assumed), and 20 years for gorillas[81], and translated into 1.28(95% CI: 1.15-1.42)$\times10^{-9}$ and 0.98(95% CI: 0.86-1.11)$\times 10^{-9}$ single-nucleotide mutations per bp per year for chimpanzees and bonobos, and 1.53 (95% CI: 1.38 – 1.70)$\times10^{-9}$ and 1.17(95% CI: 1.04-1.33)$\times10^{-9}$ single-nucleotide mutations per bp per year for gorillas, using the mutation rates from Helgason et al. [79] and Fu et al. [80], respectively. Note that the true MSY mutation rate for the great apes is likely to be higher than the estimates for humans, as has been reported for the autosomes [82].

The MCMC runs per species were performed with 200,000,000 generations, logging every 1,000 steps; the first 20,000,000 generations were discarded as burn-in using constant-sized coalescent tree prior and a strict clock. The GTR substitution model with empirical base frequencies was identified as the best fit to the chimpanzee data and HKY with empirical base frequencies to the bonobo and the gorilla data, according to the Bayesian Information Criterion (BIC) as implemented in IQ-TREE v1.6.12 [40]. A prior with a normal distribution based on the 95% CI of the substitution rate was applied. In the runs, only the variant sites were used; the composition of invariant sites was specified in the BEAST xml file. A summary tree was produced using Tree-Annotator (v.1.10.4) and visualized using the FigTree software (v.1.4.4, http://tree.bio.ed.ac.uk/software/figtree/).

### Supplemental Notes

#### Note S1. Confirming species identity of the two orangutan samples

Sumatran and Bornean orangutans are separated into different species based upon cytologic, genomic and other criteria. They are geographically isolated from each other and exhibit significant genetic divergence [83,84]. The interbreeding of captive animals from these species is well-documented[85]. It was therefore important that only cell lines from non-hybrid individuals be included in our T2T assemblies. There are genomic signatures that are characteristic of each species; these include a pericentric inversion on chromosome 2[86,87], and distinct features of the mitochondrial DNA[83,84]. The presence of a Yq nucleolar organizing region (NOR) is unique to the Sumatran orangutan which also has two distinct Y lineages that arose from a naturally occurring pericentric inversion event that did not affect male fertility[88,89]. In our analysis we confirmed species identity of our two orangutan samples with the following approaches:

**mtDNA analysis.** Accession X97707.1 is the Sumatran orangutan mitochondrial sequence from the 1996 publication[83]. It is identical to chrM in the 2018 orangutan reference genome ponAbe3. The 2011 orangutan genome paper[90] asserts that the 2011 version of the orangutan reference genome was derived from "Susie; Studbook no. 1044; ISIS no. 71", a Sumatran orangutan female. This Sumatran orangutan mitochondrial sequence aligns to the same scaffold (in our T2T Sumatran orangutan) that the Bornean orangutan T2T mitochondrial sequence does, but with higher sequence identity (≈98% vs. ≈93%). This suggests that the mother of our T2T Sumatran orangutan was a Sumatran orangutan, and that the mother of our T2T Bornean orangutan probably was not.

**Y chr analysis.** Previous literature[88] asserts that Sumatran orangutans have two distinct Y types. One of these has an insertion of *CDY* gene families on the p arm [89]. Alignment of T2T Sumatran and Bornean Ys, annotated with positions of members of ampliconic gene families, is shown in Fig. N1A below. The order of genes in our T2T assemblies is as listed in the figure legend. However, *RBMY* and *TSPY* have overlapping regions, as do *DAZ* and *CDY*. This order is mostly the same in both Sumatran and Bornean T2T assemblies, in contrast to the figure from Glaser et al. [91] included as an inset in Fig. N1A), and is generally consistent with Glaser's order for Bornean orangutan. In particular, we do not observe the Sumatran orangutan feature in which Glaser et al. show an additional *DAZ* on the "other side" of *RBMY*/*TSPY*. However, our Sumatran T2T Y assembly has a stretch of *CDY* intermixed with *RBMY* and *TSPY*, consistent with it being of Sumatran origin[89], and our Bornean T2T Y assembly does not, consistent with it being of Bornean origin[89].

In the case of our T2T Sumatran orangutan, we observed a non-inverted Y chromosome [88], with a very similar gene order to the one in Bornean orangutan. However, its sequence divergence from the Y chromosome of Bornean orangutan, the presence of *CDY* intermixed between *RBMY* and *TSPY*, and the presence of an NOR[88], suggests that the father of our Sumatran T2T orangutan was indeed a Sumatran orangutan.

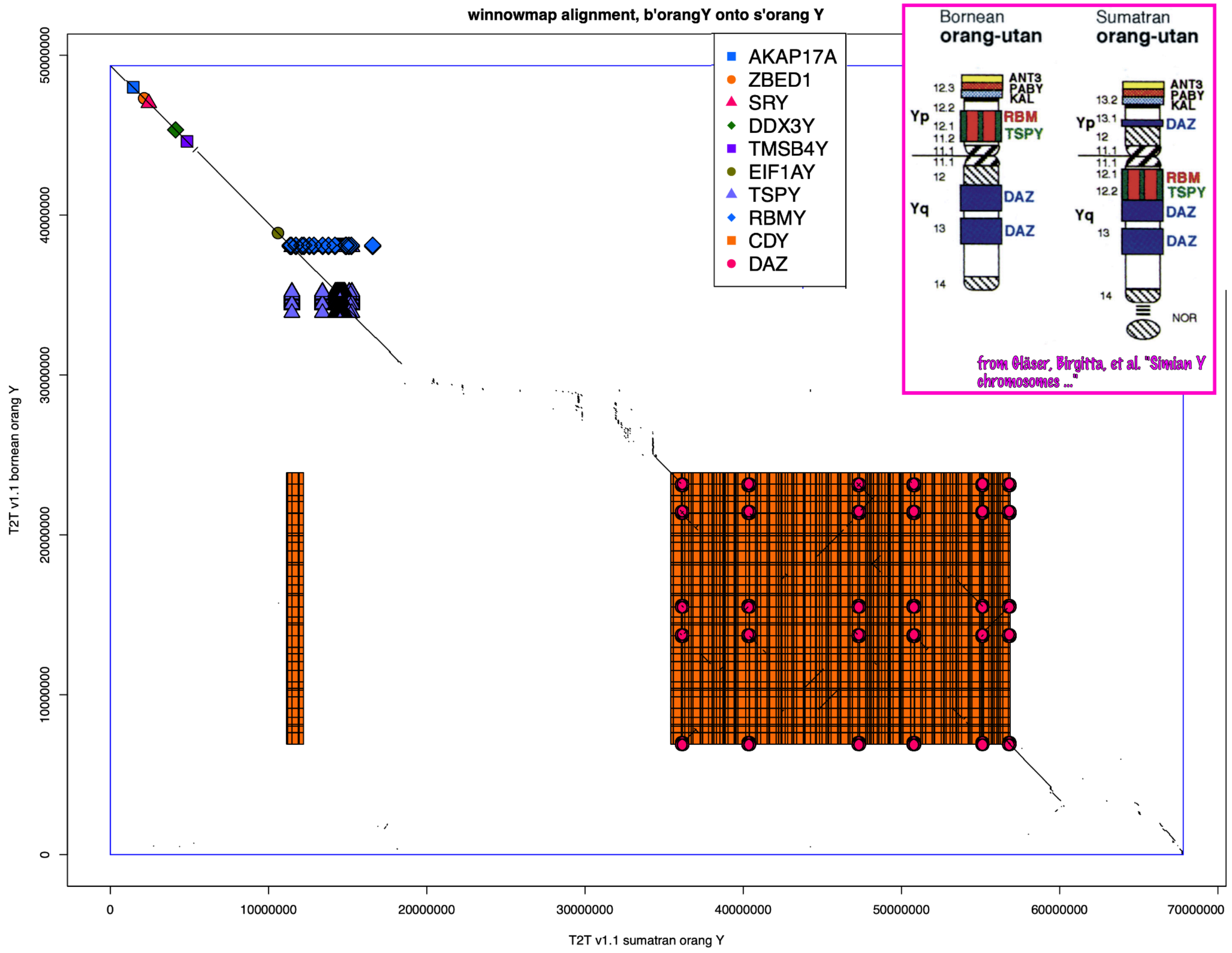

**Figure N1A. An alignment of the Y chromosomes between the T2T Bornean (the Y axis) and Sumatran (the X axis) orangutans.** Gene families are shown where both orangutan Ys aligned to corresponding human Y genes (CHM13). Underlying dotplot shows mappings of Bornean onto Sumatran orangutans.

#### Note S2. Chimpanzee subspecies identification

**Chimpanzee subspecies identification.** The subspecies (*Pan troglodytes verus*) for the chimpanzee (mPanTro3) was determined based on the similarity of mitochondrial sequences, which has been the standard since chimpanzee subspecies discrimination using mitochondrial sequence was determined to be possible[92]. The mitochondria for mPanTro3 were sequenced with the rest of the DNA and thus co-assembled with the sex chromosomes and autosomal sequences. See the Supplementary Methods for Mitochondrial Genome Assembly for details on how the final mitochondrial sequence was generated. Sequences for comparison were identified from the tree in Figure 2 from Vega *et al.* [93] and subsequently obtained from the NCBI (Table N2A). All these sequences are whole mitochondrial genomes except for KJ606391-KJ606393, which contain only a hypervariable region (HVR) of the non-coding displacement loop (D-loop)[93]. Some sequences had ambiguous nucleotides represented according to the International Union of Pure and Applied Chemistry (IUPAC) standard[94,95]. These bases were arbitrarily replaced with an appropriate standard base (A, C, G, or T) using seqtk v1.4 [96]: `seqtk randbase input.fa > output.fa`.

Sequence similarity was determined using the Mash distance[95], which was calculated with Mash (commit 41ddc61) using the following commands: `mash sketch -M -i -k 21 -o input.fa.msh input.fa` and `mash dist mPanTro3.mt.fa.msh vega.whole-mt.fa.msh > output.tsv`. The mitochondrial sequence with the shortest distance (highest number of shared hashes) to mPanTro3's mitochondrial sequence was HM068589.1 (*P. t. verus*), and mPanTro3's mitochondrial sequence was closer to all *P. t. verus* mitochondrial sequences than to mitochondrial sequences from other subspecies (Table N2B).

This determination that mPanTro3 is *P. t. verus* was confirmed by a phylogenetic tree (Fig. N2A). The same set of whole chimpanzee mitochondrial genomes, plus one bonobo mitochondrial genome (HM015213.1), were aligned using Clustal Omega v1.2.4 [97,98]: `clustalo -i input.fa --outfmt=phy -o clustalo.phy`. The tree was built using IQTREE v2.2.2.6 [39–41] with the bonobo sequence specified as the outgroup species: `iqtree -s input.phy -m TEST -B 1000 -o HM015213.1`. The best-fit model according to the Bayesian Information Criterion (BIC) [99] was TN+F+I+G4.

The alignment of mPanTro3's HVR to KJ606391-KJ606393 also confirms the assignment. The HVR for mPanTro3 was identified by finding the forward and reverse primer sequences (respectively D-88 and D-441 from Morin *et al.* [92]) using Bowtie2 v2.5.1 [100]: `bowtie2-build mPanTro3.mt.fa mPanTro3.mt.bt2idx` and `bowtie2 --local -x mPanTro3.mt.bt2idx -f -U primers.fa -S output.sam`. The primer sequences mapped uniquely and without variation (i.e., no clipping, mismatches, or indels). Based on the alignment positions, the sequence between the primers was extracted using Samtools v1.17[65]: `samtools faidx mPanTro3.mt.fa MT:15464-15827 > mPanTro3.mt.hvr.fa`. This 364 bp sequence was then aligned to KJ606391-KJ606393: `bowtie2-build vega.hvr.fa vega.hvr.bt2idx` and `bowtie2 --local -x vega.hvr.bt2idx -f -U mPanTro3.mt.hvr.fa -S output.sam`. The reported alignment was to KJ606392.1, which is *P. t. verus* (alignment starts at the 5$^{th}$ base of KJ606392.1 with the following CIGAR string: 25S222M1I116M).

**Mitochondrial genome assembly.** The mitochondrial DNA was sequenced alongside the nuclear DNA, and the sequences corresponding to the mitochondrial DNA were thus co-assembled with the rest of the genome. The mitochondrial sequence was identified based on the alignment of the human reference mitochondrial sequence (NC_012920.1) to the final assembly using Mashmap v2.0 [24]: `mashmap -r 95 -s 10000 -r NC_012920.1.fa -q assembly.fa`. The resulting alignments were filtered to keep matches with >85% identity and >5 kb alignment length to contigs <100 kb in length. For the chimpanzee (mPanTro3), the mitochondrial sequence was derived from a single node in the assembly graph. The mitochondrial assemblies were circularized (i.e., trimmed to go around the sequence only once), oriented (i.e., switched between strands) to keep the phenylalanine tRNA sequence on the forward strand, and rotated (i.e., changed which base is the start base) to begin with the phenylalanine tRNA sequence. The phenylalanine tRNA sequence was identified with tRNAscan-SE v2.0.11[101] (`-M mammal`). Overlap of the mitochondrial sequence with itself to identify where to trim the sequence was identified by alignment with nucmer v4.0.0rc1[102] (`--nosimplify`

`--maxmatch -f -b 5000`). Trimming, orienting, and rotating the sequence was accomplished with SAMtools faidx v1.16.1[65].

**Table N2A. Mitochondrial sequences for comparison, identified from the tree in Figure 2 of Vega *et al*. [93].** Some accession versions have been updated since 2014, and those updated versions were used here. Abbreviations: bp = base pairs, *P.* = *Pan*, *t.* = *troglodytes*.

| Accession | Taxonomic group | Sequence type | Sequence length (bp) |
|---:|---:|---:|---:|
| GU112741.1 | *P. t. verus* | Whole mitochondria | 16,561 |
| GU112742.1 | *P. t. ellioti* | Whole mitochondria | 16,564 |
| HM015213.1 | *P. paniscus* | Whole mitochondria | 16,569 |
| HM068575.1 | *P. t. ellioti* | Whole mitochondria | 16,562 |
| HM068577.1 | *P. t. schweinfurthii* | Whole mitochondria | 16,557 |
| HM068585.1 | *P. t. ellioti* | Whole mitochondria | 16,567 |
| HM068586.1 | *P. t. ellioti* | Whole mitochondria | 16,564 |
| HM068589.1 | *P. t. verus* | Whole mitochondria | 16,556 |
| HM068592.1 | *P. t. schweinfurthii* | Whole mitochondria | 16,564 |
| HM068593.1 | *P. t. verus* | Whole mitochondria | 16,560 |
| JF727162.2 | *P. t. troglodytes* | Whole mitochondria | 16,563 |
| JF727164.2 | *P. t. troglodytes* | Whole mitochondria | 16,557 |
| JF727168.1 | *P. t. troglodytes* | Whole mitochondria | 16,564 |
| JF727172.2 | *P. t. troglodytes* | Whole mitochondria | 16,558 |
| JF727174.1 | *P. t. troglodytes* | Whole mitochondria | 16,563 |
| JF727177.1 | *P. t. troglodytes* | Whole mitochondria | 16,561 |
| JF727180.2 | *P. t. troglodytes* | Whole mitochondria | 16,562 |
| JF727184.1 | *P. t. schweinfurthii* | Whole mitochondria | 16,558 |
| JF727186.1 | *P. t. schweinfurthii* | Whole mitochondria | 16,557 |
| JF727187.1 | *P. t. schweinfurthii* | Whole mitochondria | 16,555 |
| JF727198.1 | *P. t. schweinfurthii* | Whole mitochondria | 16,556 |
| JF727200.1 | *P. t. schweinfurthii* | Whole mitochondria | 16,558 |
| JF727202.2 | *P. t. ellioti* | Whole mitochondria | 16,560 |
| JF727203.1 | *P. t. ellioti* | Whole mitochondria | 16,563 |
| JF727205.1 | *P. t. ellioti* | Whole mitochondria | 16,564 |
| JF727210.3 | *P. t. verus* | Whole mitochondria | 16,559 |

| | | | |
|---|---|---|---|
| JF727212.2 | *P. t. verus* | Whole mitochondria | 16,557 |
| JF727213.1 | *P. t. verus* | Whole mitochondria | 16,558 |
| JF727215.1 | *P. t. verus* | Whole mitochondria | 16,557 |
| KJ606391.1 | *P. t. schweinfurthii* | Hypervariable region | 367 |
| KJ606392.1 | *P. t. verus* | Hypervariable region | 518 |
| KJ606393.1 | *P. t. troglodytes* | Hypervariable region | 460 |

**Table N2B. Mash distances between mPanTro3's whole mitochondrial sequence and the whole mitochondrial sequences from samples where the subspecies has previously been identified.** The entries are sorted by Mash distance.

| ***Pan troglodytes* subspecies** | **Accession** | **Mash distance** | ***p*-value** | **Shared hashes (out of 1,000)** |
|---:|---:|---:|---:|---:|
| *verus* | HM068589.1 | 0.000680979 | 0 | 972 |
| *verus* | HM068593.1 | 0.00448671 | 0 | 835 |
| *verus* | JF727213.1 | 0.00451781 | 0 | 834 |
| *verus* | JF727212.2 | 0.00464278 | 0 | 830 |
| *verus* | GU112741.1 | 0.00597612 | 0 | 789 |
| *verus* | JF727210.3 | 0.00969064 | 0 | 689 |
| *verus* | JF727215.1 | 0.0103562 | 0 | 673 |
| *ellioti* | JF727202.2 | 0.0108702 | 0 | 661 |
| *ellioti* | HM068585.1 | 0.0112201 | 0 | 653 |
| *ellioti* | JF727205.1 | 0.0116661 | 0 | 643 |
| *ellioti* | GU112742.1 | 0.0118471 | 0 | 639 |
| *ellioti* | JF727203.1 | 0.0118471 | 0 | 639 |
| *ellioti* | HM068575.1 | 0.0121679 | 0 | 632 |
| *ellioti* | HM068586.1 | 0.0122141 | 0 | 631 |
| *troglodytes* | JF727172.2 | 0.0189925 | 0 | 505 |
| *troglodytes* | JF727180.2 | 0.0191181 | 0 | 503 |
| *schweinfurthii* | JF727187.1 | 0.0192445 | 0 | 501 |
| *troglodytes* | JF727174.1 | 0.0193079 | 0 | 500 |
| *schweinfurthii* | JF727184.1 | 0.0194352 | 0 | 498 |
| *schweinfurthii* | JF727186.1 | 0.0195632 | 0 | 496 |
| *troglodytes* | JF727177.1 | 0.0196275 | 0 | 495 |
| *schweinfurthii* | JF727198.1 | 0.0196919 | 0 | 494 |
| *schweinfurthii* | HM068577.1 | 0.0198213 | 0 | 492 |
| *schweinfurthii* | HM068592.1 | 0.0198862 | 0 | 491 |
| *troglodytes* | JF727168.1 | 0.0198862 | 0 | 491 |
| *schweinfurthii* | JF727200.1 | 0.0199514 | 0 | 490 |
| *troglodytes* | JF727162.2 | 0.0200167 | 0 | 489 |
| *troglodytes* | JF727164.2 | 0.0204789 | 0 | 482 |

**Figure N2A.** Cladogram of *Pan* whole mitochondrial sequences, rooted using *P. paniscus* as an outgroup. Nodal support values were determined by bootstrap with 1,000 replicates.

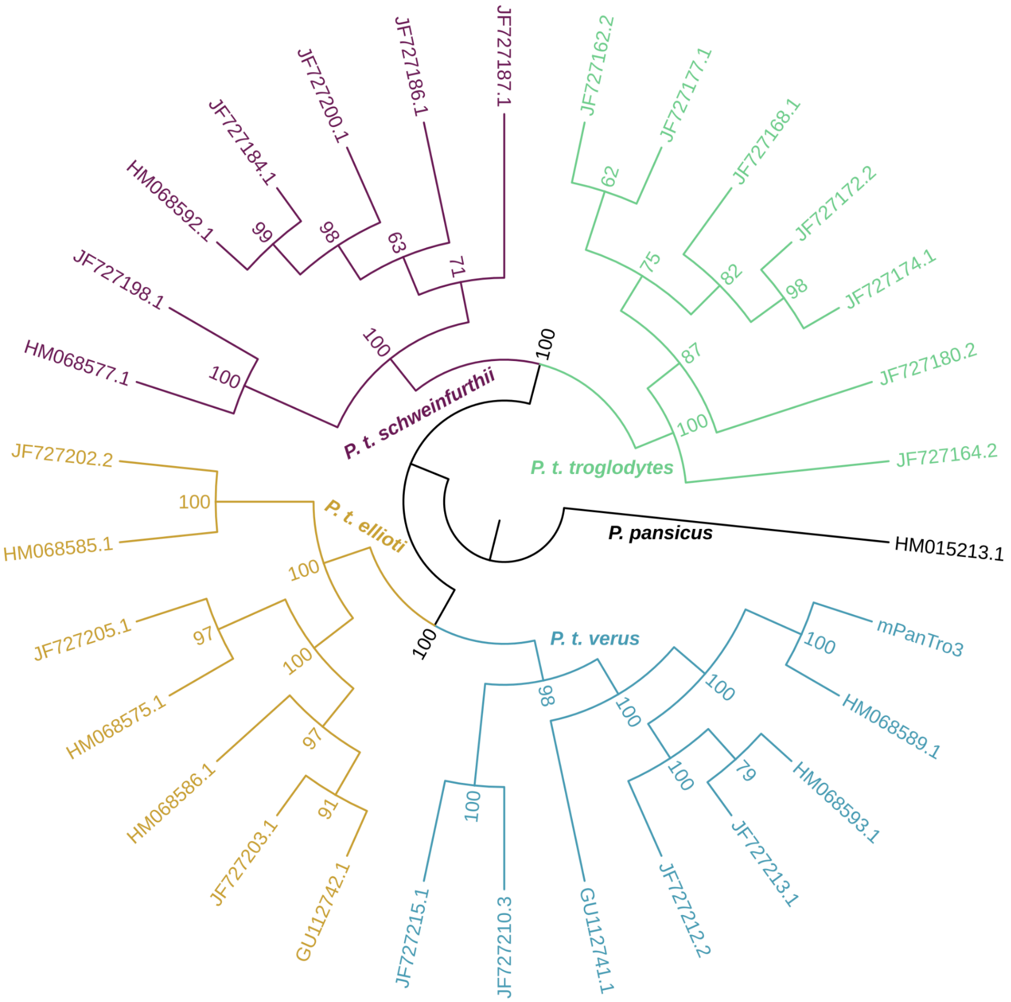

### Note S3. Bonobo PAR2 and Ariel satellites

The newly discovered here bonobo pseudoautosomal region 2 (PAR2) on chrX:156,408,085-156,502,125 and chrY:46,801,687-46,897,825 is immediately followed by a novel bonobo-species-specific satellite Ariel, which has a 63-bp repeat unit (chrX: 156,502,116-156,522,247 and chrY: 46,897,816-46,906,281) and spans for 20 kb at both chrX and chrY, making it distinct from PAR2 in human and from PAR1 shared by great apes. The assembly in this region is reliable (except for a likely 453-bp deletion, in the middle of the Ariel repeat cassette, on chromosome X), and the entire bonobo PAR2 and Ariel satellite are contained within four ultralong ONT reads on chrX (Fig. N3A). In the bonobo assembly, the Ariel satellite is also present on chromosomes 1, 2B, 3-14, 16, and 18 (based on alignments of the canonical Ariel repeat unit with <5% divergence). The repeat cassette is always within 4 Mb of the end of a chromosome, and, in most cases, is within 50 kb. Three of these chromosomes, 2B, 3, and 9, also contain all or part of the PAR2 sequence immediately adjacent to the Ariel satellite, as in chromosomes X and Y (Fig. N3B). The copy on chromosome 9 is inverted relative to X and Y, and there is a second, partial copy on chromosome 3. To a lesser extent than the PAR2 and Ariel satellite sequence, these three chromosomes also share some similarity beyond PAR2 and the Ariel satellite to the ends of their chromosomes with the immediately adjacent portion on chromosome X (Fig. N3C).

**Ariel repeat alignment.** The canonical Ariel repeat unit, 5'-ATAATATCCACACCATGCCCTATCACTGATCTAATCCACACCATCGCTTCCAATACTAATGTA-3', was aligned to the full diploid assembly using BLAST+ v2.14.0 [103,104]:

```
makeblastdb -input_type nucl -in dip-asm.fa -title dip-asm -hash_index -out
dip-asm; blastn -query ariel.fa -db dip-asm -out ariel-x-dip-asm.tsv
-perc_identity 95 -qcov_hsp_perc 95
```

**Bonobo PAR2 alignment.** PAR2 and flanking region were aligned to the autosomes in the “primary” pseudohaplotype assembly (GCF_029289425) using Winnowmap v2.03 [29]:

```
winnowmap -x asm20 -o output.paf asm.fa chrx-par2.fa
```

**Chromosome end alignments.** To use alignments to check for additional similarity after bonobo PAR2 between (a) chromosomes X and Y and (b) the autosomes with bonobo PAR2 sequence, chromosomes ends were extracted with SAMtools v1.18 [65] based on the coordinates of the matches from the previously-performed Winnowmap v2.03 [29] PAR2 alignments. Example command with chromosome X:

```
samtools faidx asm.fa chrX:156,502,125- >> chromosome-ends.fa.
```

The coordinates for chromosome ends are the following:

```
chr2B_haplotype1-0000013:144,801,512-, chr3_haplotype1-0000015:196,275,038-,
chr9_haplotype1-0000014:1-47,980, chrX:156,502,125-, chrY:46,897,825-
```

The alignments were performed with Mashmap v2.0 [24]:

```
mashmap -f none -k 16 --pi 95 -s 500 -r chromosome-ends.fa -q
chromosome-ends.fa -o chromosome-ends.ssv
```

**Figure N3A. ONT read alignments to bonobo chromosome X PAR2 and Ariel satellite.** IGV [18] screenshot showing four ultralong ONT reads (purple) spanning both PAR2 (red) and Ariel repeats (green). The PacBio HiFi alignments (not shown) also align well and support the 453-bp deletion in the middle of the Ariel repeat cassette.

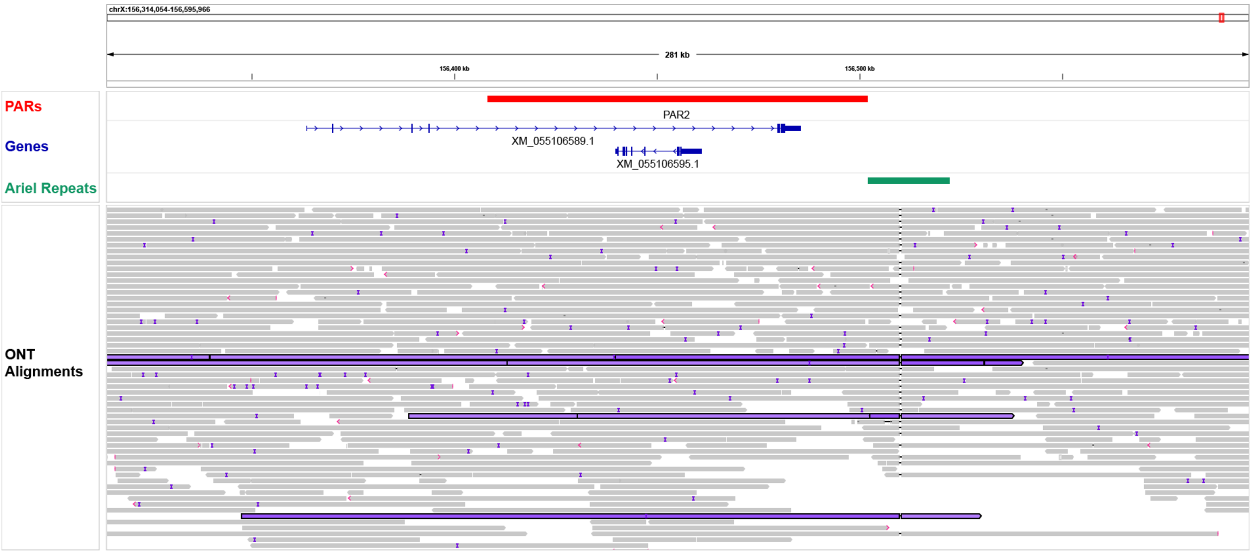

**Figure N3B. Dotplots of bonobo chromosome X PAR2 alignments to bonobo chromosomes 2B, 3, and 9.** Dotplots of Winnowmap v2.03[29] alignments of chromosome X PAR2 and flanking region to chromosomes 2B (left), 3 (center), and 9 (right). The x-axis is chromosome X, and PAR2 is demarcated by red lines. The sizes of the Ariel satellite on chromosomes 2B and 9 are similar to the Ariel satellite on chromosome X (and Y), but noticeably smaller on chromosome 3. While smaller on chromosome 3, the Ariel satellite block is also duplicated with part of PAR2 approximately 125 kb downstream.

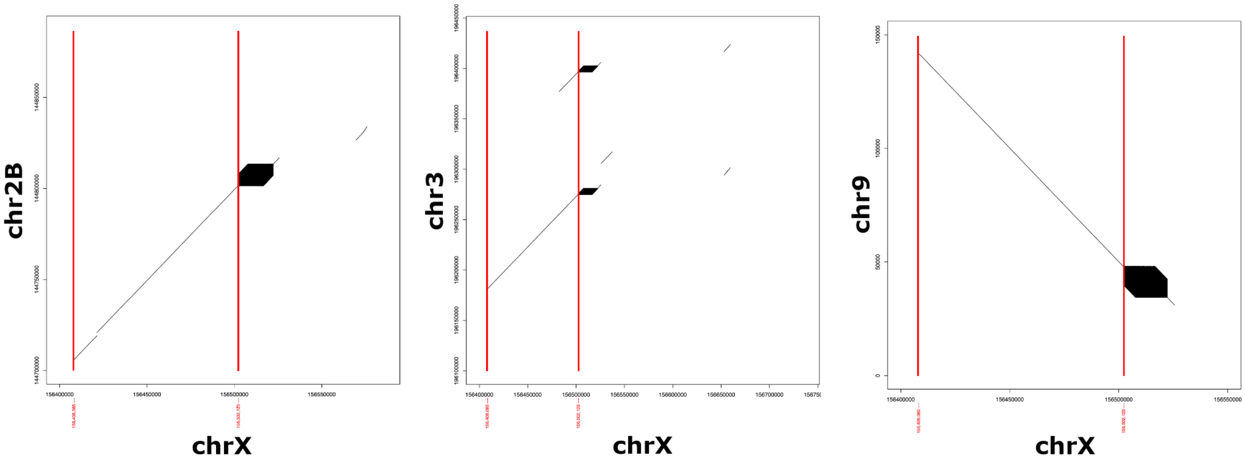

**Figure N3C. Dotplots of post-PAR2 chromosome X alignments to chromosomes 2B, 3, and 9.** Dotplots of MashMap[24] approximate alignments of chromosome X after PAR2 to the ends of chromosomes 2B (left), 3 (center), and 9 (right). With the exception of some gaps in similarity, chromosomes 2B, 3, and 9 are roughly similar to chrX after PAR2 until they end. Chromosome X has approximately 3.5 Mb of sequence after PAR2, whereas chromosomes 2B, 3, and 9 have, respectively, approximately 80 kb, 160 kb, and 45 kb. Chromosome X also has a large expansion relative to chromosome 2B.

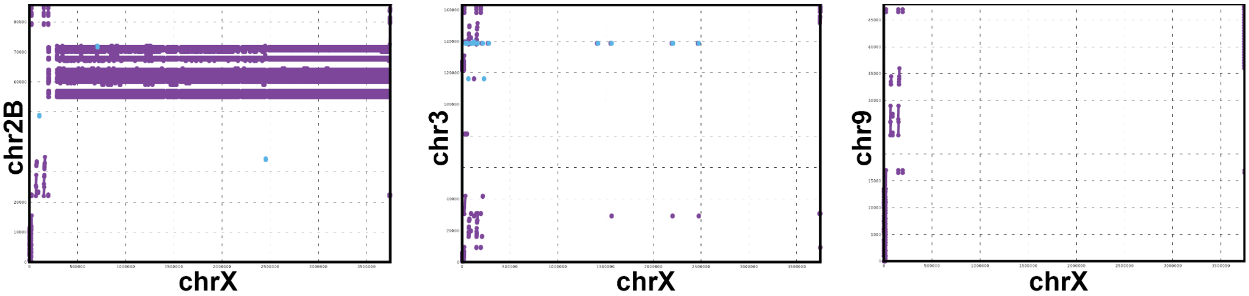

#### Note S4. Searching for X-Transposed Regions

The human Y chromosome has a large (3.4-Mb) transposition (duplication) from the human X chromosome, which occurred ~3-4 MYA, i.e. after the divergence of the human and chimpanzee lineages [12,105]. This X-transposed region (XTR) is characterized by >99% sequence identity with the corresponding region on the X chromosome, and it has since been split into two blocks due to a subsequent inversion event. It has a different evolutionary history relative to the other regions that have some sequence similarity with the X chromosome, such as PARs and other sequence classes in the male-specific region—ampliconic and ancestral. We confirm the finding that the appearance of the human XTR occurred after the divergence of human and chimpanzee lineages because we see no evidence of the human XTR on the Y chromosomes of the other apes (Fig. N4A). We also found no evidence of other XTRs, i.e., of sequence transposed from the X to the Y chromosome in one or more non-human primates (Fig. N4B).

**Searching for human XTR in non-human primate Y chromosomes**. Alignments between human (HG002) chromosome Y from T2T-CHM13 v2.0[75,106] and the non-human primate Y chromosomes (this study) were performed with lastZ[36] (see Supplemental Methods – Pairwise alignments) and plotted using R v4.3.0[107] with ggplot2 v3.4.3[108]. The colors for the human sequence classes match the colors used for sequence classes throughout this manuscript, and they were taken from Rhie et al.[13], along with the positions of the sequence class intervals. The R script can be found at https://github.com/makovalab-psu/T2T_primate_XY/tree/main/xtr_search. The input to the R script requires lastZ's "`--format=rdotplot`" output, as described in the lastZ manual (https://lastz.github.io/lastz/#fmt_rdotplot). Since the alignments were run with a variant of "`--format=general`" (with fields name1, zstart1, end1, name2, strand2, zstart2+, end2+, nmatch, id%, and cigarx), the following GNU AWK v4.2.1 command was used to convert to the other format:

```
awk 'BEGIN{FS=" +"; OFS="\t"}{ts=$2; te=$3; qs=$6; qe=$7; if($5 == "-"){tmp=te;
te=ts; ts=tmp} print qs, ts; print qe, te; print "NA", "NA"}' < file.lz >
file.dots
```

**Searching for non-human XTR in non-human primate Y chromosomes.** Two methods were employed to search for non-human XTRs in the non-human primate Y chromosomes. The first method searched for a unique Y sequence with high similarity to X within the same species; this would only find an XTR if it were unique to a given species (or a closely related one). For each species, alignments (see Supplemental Methods – Pairwise alignments) between Y and X for each species (i.e., bonobo Y vs. bonobo X, gorilla Y vs. gorilla X, etc.) were considered if they had ≥94% identity, had >1 kb of noncontiguous matches, and did not overlap the previously-annotated Y PAR region(s) (see Supplemental Methods – Classifications into PARs, ancestral, ampliconic, and X-transposed regions). These positions were further filtered by removing overlaps with any alignment to another non-human primate species' (outside the same genus) Y chromosome (also at >=94% identity and >1 kb matches). The remaining coordinates were primarily ancestral regions (based on separate annotation) and repeats, suggesting that no species-specific XTRs arose for the non-human primates.

The second method considered similarity between the Y and the X for a given species and relied on the sequence class annotations (see Supplemental Methods – Classifications) into PARs, ancestral, ampliconic, and X-transposed regions; this dropped any assumption or requirement that a transposition event from the X is unique to a given species. Interchromosomal segmental duplications between a given species Y and X chromosomes were used as an initial set, and these were reduced by removing overlap with the annotated sequence classes. All remaining sequences (when there were any for a given species) were short (approximately ≤20 kb) and/or low identity (some ~97%, most <95%). The precise origin of these otherwise unclassified sequences is not yet known, but no large transpositions from the X chromosome were evident for any species other than humans. The unfiltered alignments are plotted as a dotplot with the sequence class annotations colored in the background (Fig. N4B). Plotting was done using R v4.3.0[109] with ggplot2 v3.4.3[110] and cowplot v1.1.1[111], and the full R script is available at https://github.com/makovalab-psu/T2T_primate_XY/tree/main/xtr_search.

**Figure N4A. Similarity between the Ys of human and non-human primates.** Dotplots based on lastZ[36] alignments of the human (HG002) Y chromosome to the respective non-human primate Ys. The background colors show the sequence class of the respective positions on the human Y chromosome. Certain repetitive elements were soft-masked before mapping, so no dots are expected in the plot despite the expected sequence similarity. The XTR (pink) was unmasked, and the human sequence has little-to-no similarity with the non-human primate sequences, indicating that the XTR is uniquely transposed onto human Y. See a similar dotplot between two human Y chromosomes in Extended Data Fig. 3 from [13] for comparison.

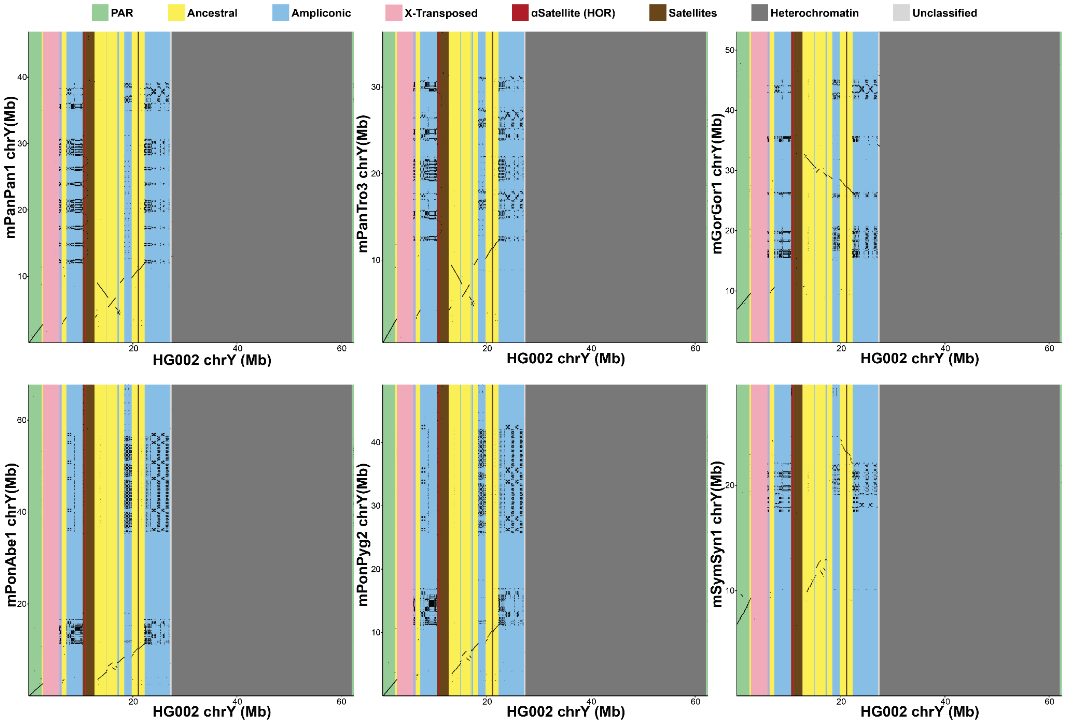

**Figure N4B. Similarity between the Y and X chromosomes of non-human primates.** Dotplots based on lastZ[36] alignments of the non-human primates Y chromosomes to their respective X chromosomes. The background colors show the sequence class annotations on the Ys. Certain repetitive elements were soft-masked before mapping, so no dots are expected in the plot despite the expected sequence similarity. No alignments are present in the Unclassified (light gray) category for the orangutans, and only short (approximately ≤20 kb) and/or low percent identity alignments (mostly <95%, though some ~97%) are present in the same category for the other species.

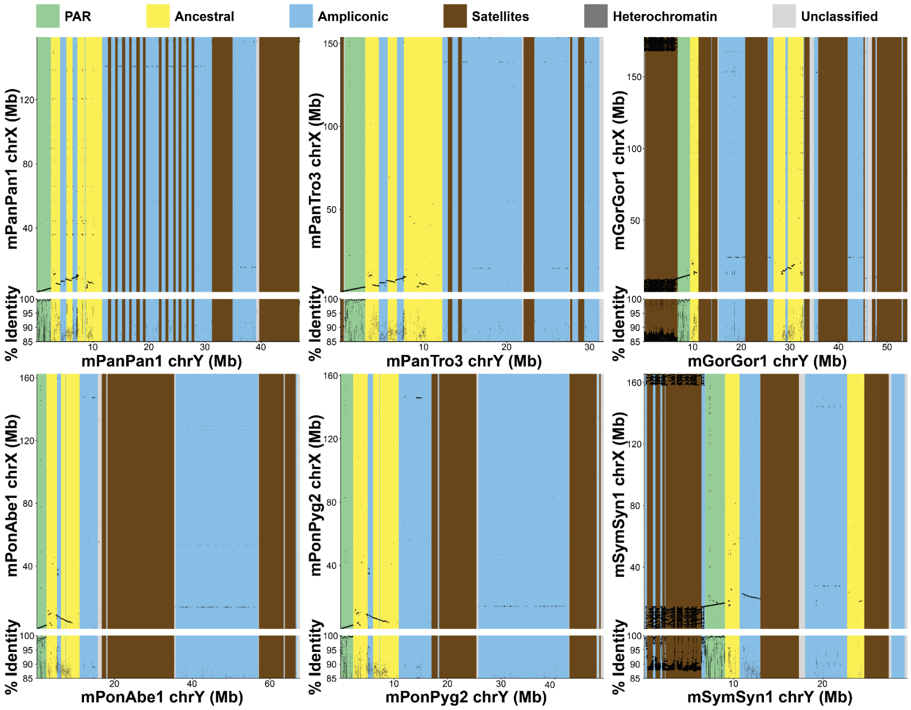

#### Note S5. The comparison of ampliconic regions, palindromes, and segmental duplications

We have studied the ampliconic regions, palindromes, and segmental duplications on the Y chromosomes of great apes. Per Skaletsky et al. 2023[12], ampliconic regions are long multi-copy regions with >50% sequence identity (although in practice higher values are observed). These regions represent parts of the Y chromosome with a unique evolutionary history shaped by rapid evolution and consisting of multi-copy amplicons hosting multi-copy gene families.

Within ampliconic regions, palindromes represent inverted repeats with ≥8-kb arms of at least 98% identity, separated by no more than 500 kb. Conserved palindromes are especially gene-rich, and we show that shorter spacers, as well as longer arms, result in higher sequence identity between chromosome arms (Fig. S7, Table S15), presumably due to gene conversion. Therefore, palindromes represent a distinctive subset of the ampliconic regions.

In contrast, segmental duplications are >1-kb multi-copy regions with >90% identity as originally described by Bailey et al. 2001[112]). The feature is computed genome-wide and is based on softmasking common repeat sequences such as retrotransposons and satellites to establish seeds and then constructing optimal pairwise alignments of the minimum length and % identity [61]. While SDs identify regions of both inverted and direct orientation, these parameters mean that SDs, ampliconic or palindromic regions are handled differently based on alignment parameters and how common repeats are processed, justifying the need for multiple computational approaches to comprehensively characterize these features of the Y chromosome. Thus, ampliconic regions, palindromes, and segmental duplications sometimes capture different regions, albeit with a very large degree of overlap.

In the figures below, ampliconic regions are plotted in **blue**, palindromes in **gray**, and intra-species segmental duplications in **orange**.

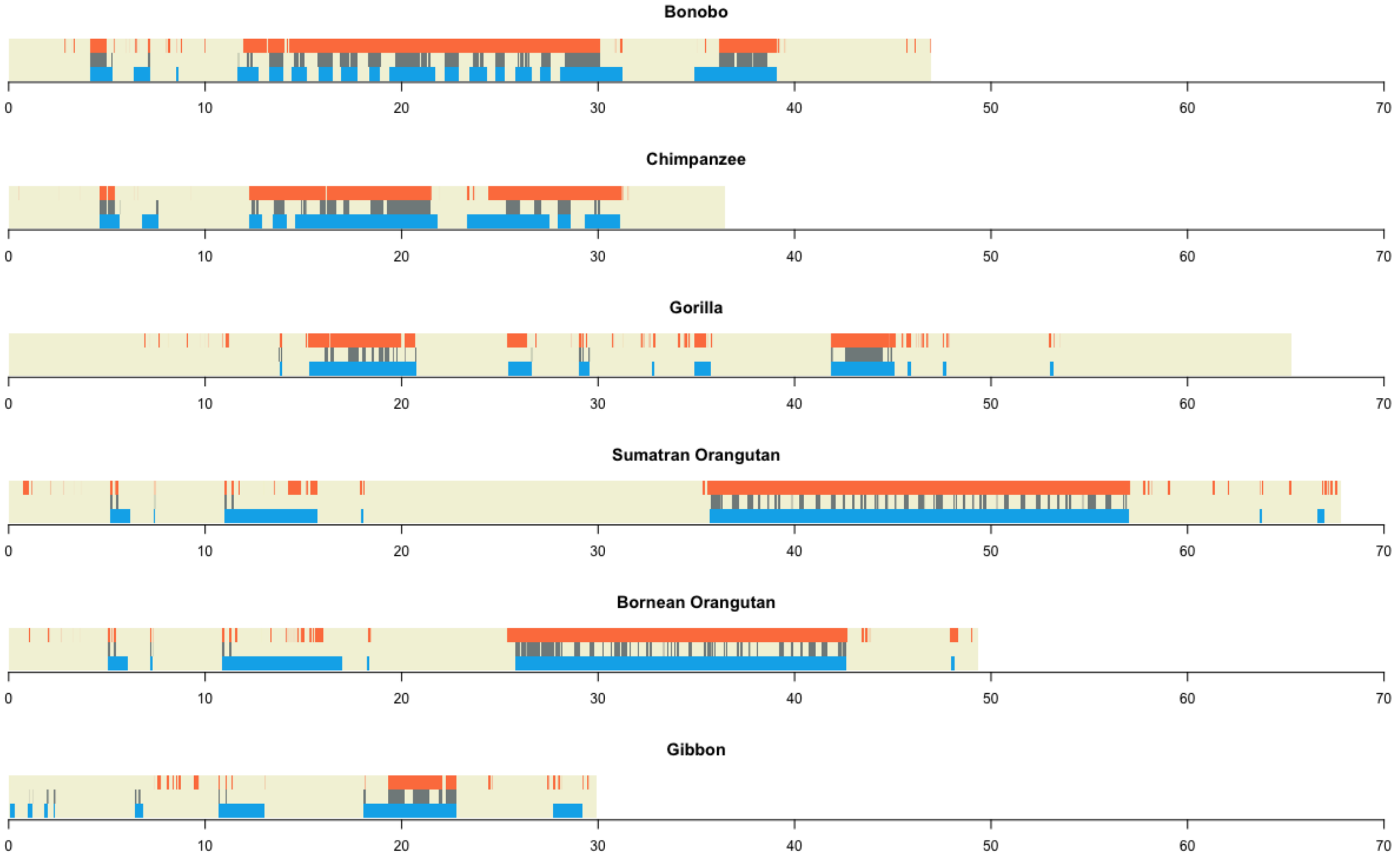

### Note S6. Lineage-specific (LS) repeat expansions

The X and Y LS insertion patterns between closely related species were largely shared, with few exceptions, including an orangutan-specific region of SINE insertions on the Y (Table S25), an additional LS stretch of SINEs ~18-kb-long on the Sumatran orangutan Y, a LS pCHT/StSat expansion at the terminal end of the bonobo Xq, and a LS pCHT/StSat expansion on the terminal end of the chimpanzee Yp. Moreover, the bonobo Yq arm contains a small, but densely populated region of LS DNA element insertions, eight LS loci composed of ACRO composite subunit array expansions, and a LS SINE expansion spanning ~98 kb. While human and gorilla Xs carry evenly distributed TE insertions, the gorilla Y carries a unique TE insertion distribution, with five distinct, densely populated regions of all TE types punctuated by stretches of satellite and simple repeats (Extended Data Fig. 2). In fact, all 40 LS insertions on gorilla sex chromosomes are present on the Y, 18 of which are expansions of satellites.

Analysis of repeat content across all LS insertions revealed variable TE types and satellite arrays contribute to LS insertion patterns among the primates in this study (Fig. 4A, Ext. Data Fig. 2, Table S24, Table S25). For example, while most LS TE insertions in the *Pongo* genus are shared, the Bornean orangutan contains a higher level of LS SINE insertions. Moreover, chimpanzee contains an increase in LS RNA insertions (i.e., tRNA, scRNA, snRNA, srpRNA and rRNA) taking into account both sex chromosomes compared to bonobo, suggesting that the unique repertoire of TE insertions contributes to primate genome heterogeneity.

### Note S7. Centromere satellite analysis

**1. Gorilla.** Here, as an example, we describe the analysis of gorilla alpha satellites (AS) in full detail to illustrate what more subtle information can be extracted from annotation tracks developed for this project. Note that StV annotations, which show altered monomer order in AS HORs, are different from the broadly used genome-wide SV annotations. We pay special attention to dramatic changes in centromere identity further called “interlayers" for brevity, where the previously existing centromere array was split and inactivated by expansion of a different HOR, which may belong to the same or to a different suprachromosomal family (SF). The interlayers we describe in gorilla belong to the latter kind and show how the most dramatic of possible changes happens in a centromere (save the introduction of a completely different satellite).

**1.1. cenX.** Gorilla cenX has an unusually complex structure that features not only the SF01/SF3 interlayer, which is shared by *Pan* and *Homo*, but also two additional interlayers, which are specific to gorilla (SF3/SF2 and SF2/SF1, in that order, Fig. N7A). Also, most of the centromeric AS array in gorilla is inverted relative to *Homo* and *Pan* with both inversion breakpoints in SF3. The SF3 array is based on 12-mer periodicity shared with *Homo* and *Pan* cenX HORs, the SF2 and SF1 cenX HORs are unique to gorilla. Additionally, an SF5 HOR array is present on the right flank, which is unique to gorilla. Below we will consider the details of gorilla cenX organization.

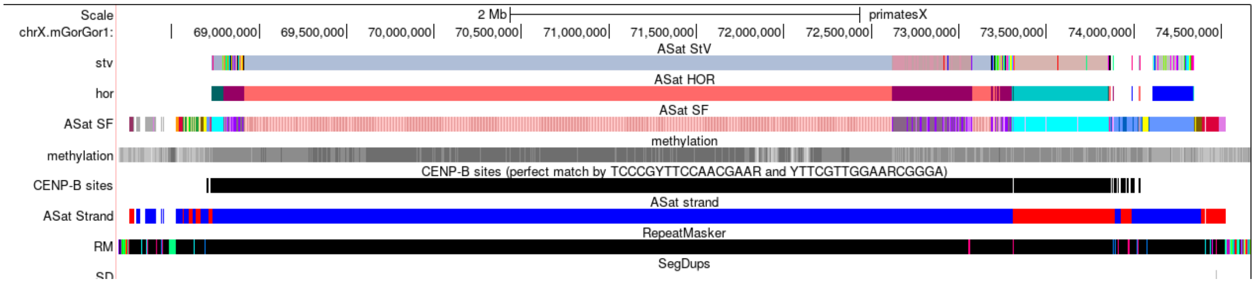

**Figure N7A. Inversion and interlayers in gorilla cenX.** General view of gorilla cenX. Various close-ups of the parts of this centromeric region are shown in Fig. S15E and Fig. N7B-N7L.

**1.1.1. Inversions.** As the SF3 part is shared with *Pan* and *Homo*, only the flanks of the gorilla array go in reverse orientation, same way as in the other two genera, and the proximal parts are inverted (Fig. N7B). Hence gorilla has experienced an inversion of all the central part of the centromere with both breakpoints in SF3. It is not clear whether it happened after the interlayers occurred in SF3, or before them when the SF3 centromere was still undisrupted and active. In either case, we would note that SF2 and SF1 sequences have been inserted in SF3 in the same orientation. In human cen1, which features the SF3/SF1 interlayer, the SFs are in opposite orientations[54].

The left breakpoint can be located quite clearly. The exact site is in this 23bp window: chrX.mGorGor1:68,731,465-68,731,487. The right breakpoint is less precise because of some TE insertions on the border. As shown in Fig. N7B, there is an insertion of L1PA2 on the border and a small piece of irrelevant AS, which may have been transduced here by L1 element from another location (R1, R2 and Ga monomers). So, due to the TE insertion, the right breakpoint window is 7.3 kb at chrX.mGorGor1:73,306,162-73,313,490.

The inversion involved the whole centromere core (likely a few Mb) and occurred after the divergence of the gorilla lineage from the human-chimpanzee clade. However, we cannot discern whether this event preceded or followed the consecutive SF3/SF2 and SF2/SF1 interlayers that occurred in the same branch.

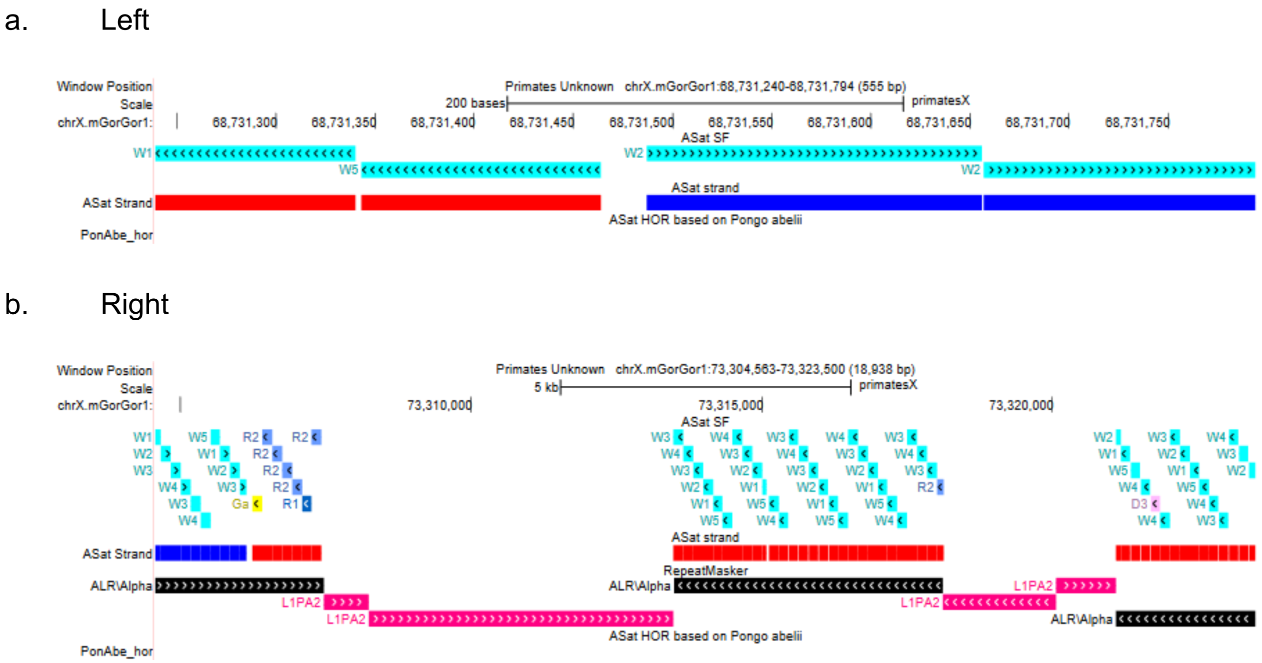

**Figure N7B. CenX inversion breakpoints.**

**1.1.2. Centromere interlayering.** SF01/SF3 interlayer implies that the SF01 centromere that once existed in the common ancestor of African apes has been split and replaced by SF3 centromere in all upstream branches. Unlike in humans, the remnants of the dead SF01 array are clearly seen in gorilla and *Pan* on both flanks. We next consider both flank arrays in detail.

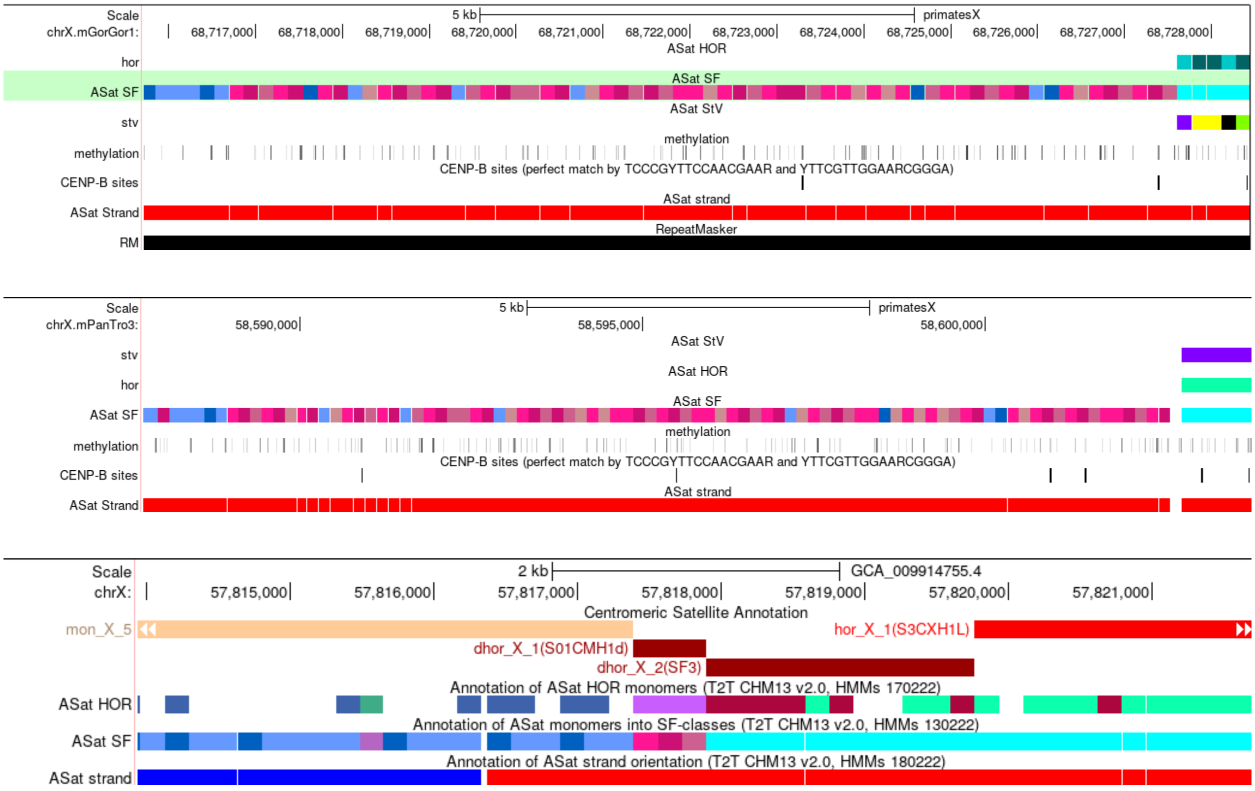

**Figure N7C. The left-flank cenX SF01 array in African apes.**

Fig. N7C shows that in gorilla (upper panel) and chimpanzee (middle panel), the left flank SF01 array (different hues of cherry color sprinkled with blue in SF-track) is relatively large, located between the SF5 array on the left (dark and light blue) and SF5 array on the right (cyan), and it goes in reverse orientation, same as the active SF5 centromere in these species (red in the strand track). In human cenX (bottom panel), only three monomers are left from this array (hues of cherry in SF-track), which are located in the same position between SF5 (blue) and SF3 (cyan). Note the inversion in the blue layer on the left (blue color in the strand track).

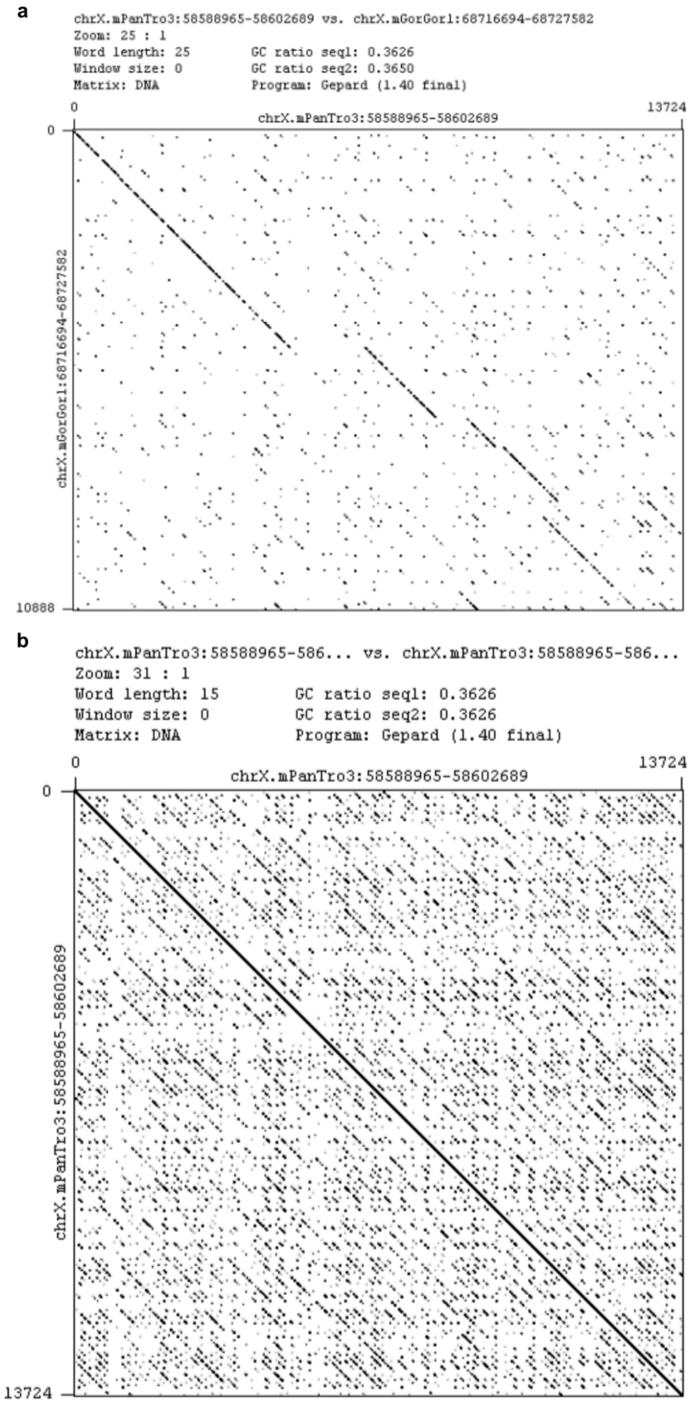

**Figure N7D. Analysis of the left-flank SF01 array in gorilla and chimpanzee. (a)** The dotplot comparison reveals structural variation that differentiates the linear organization of the left flank of the gorilla and chimpanzee X centromere arrays, that is, the broken diagonal indicates a number of deletions in gorilla and one in chimpanzee. **(b)** shows the self-plot for the chimpanzee array.

Fig. N7D shows the dot-plot analysis of the left-flank SF01 array. The diagonal line (a) is 96.9% which is about the expected divergence for the two species. This means that the regions are clearly orthologous. But the self plot (b) shows that the HOR structure, which is expected for the SF01 array, is completely disrupted (no clear additional diagonals parallel to the central one), which means that the array was inactivated long time ago and had enough time to have the HOR structure disrupted by mutations. Thus, it seems that the SF01 centromere had died before the separation of gorilla and chimpanzee, which aligns with both already having the younger SF3 centromeres at that time.

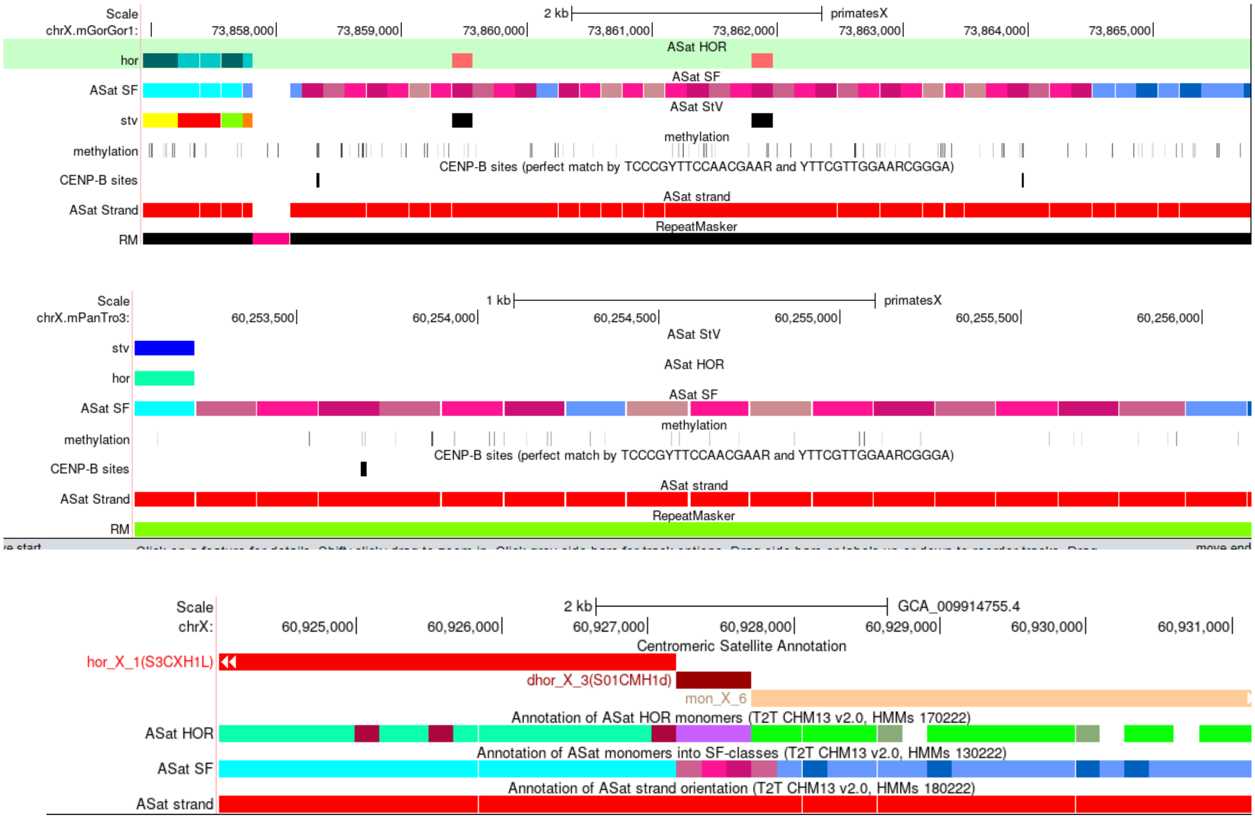

**Figure N7E. The right-flank cenX SF01 array in African apes.**

Figure N7E shows the right flank of the SF01 array in the same way as in Fig. N7C. The vestigial array is relatively large in gorilla, smaller in chimpanzee, and only 4 monomers long in humans. In all three cases, the location is the same, between SF5 (cyan) on the left and another SF5 (blue) on the right.

The best diagonal piece in Fig. N7Fa is 96.35% which is not far from the value for the other flank. The second best diagonal is 94.7%, which is substantially lower. This suggests that the region indicated in (a) is an orthologous piece of ancestral array shared by both species. The other regions represent pieces that have survived only in one of the species. The self-plot shows a typical pattern of disrupted HORs (side diagonals are all broken). Thus, the features in the left- and right-flank SF01 arrays are similar, which is consistent with them once being parts of the same array.

**1.1.3. SF3.** The SF3 arrays are situated symmetrically on the flanks between the more distal SF01 arrays and more proximal SF2 arrays. They are formed by different HORs, a 16mer on the left flank and 19mer on the right. Both HORs are based on SF3 12mer with a monomer structure identical to that of *Homo* and *Pan* cenX 12mer HORs. It is likely that the active centromere of the human-gorilla ancestor (before the interlayering) was also formed by the 12mers, and the current 16 and 19 mers are just recent amplifications and therefore are pseudocentromeres (arrays formed by recently re-amplified old material) as opposed to relic centromeres, which are the intact (not re-amplified) pieces of the old material usually represented by dHORs (divergent HORs) or monomeric layers. Moreover, the small pieces with the 12mer structure can be seen at the flanks of the arrays on both sides (see details in HOR section below). As we reported above, the centromere in gorilla is inverted relative to *Homo* and *Pan*. Both inversion breakpoints are in SF3 AS.

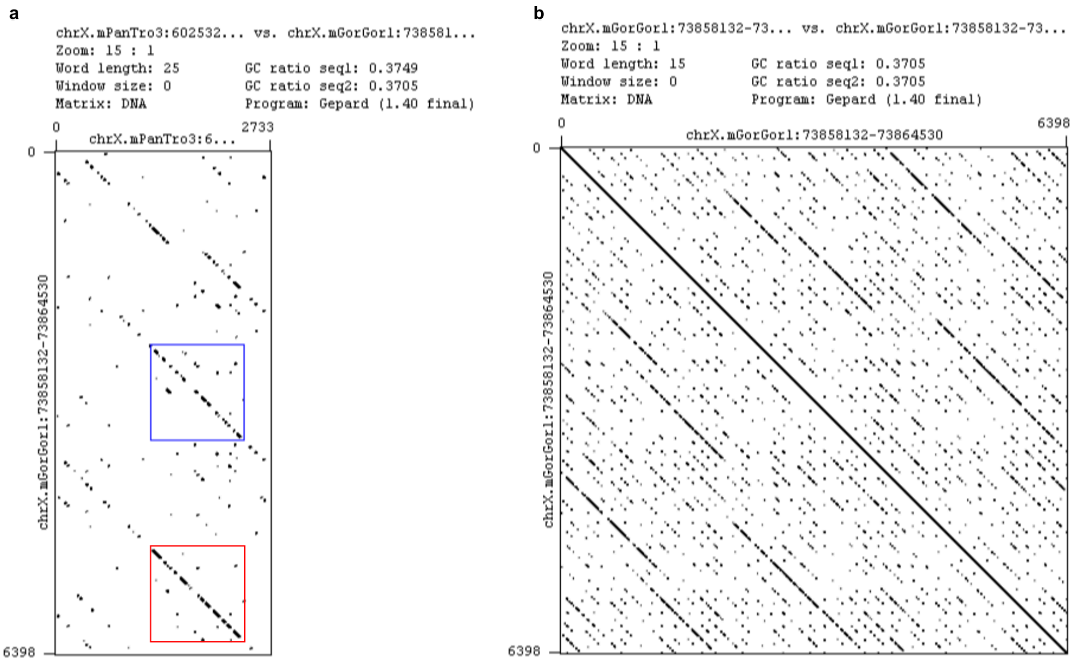

**Figure N7F. Analysis of the right-flank SF01 array in gorilla and chimpanzee.** (a) The dot-plot comparison which seems to have a piece of a better diagonal which would indicate a shared orthologous piece (marked by a red rectangle) and some pieces of not so good diagonals which likely represent non-orthologous pieces. A second best diagonal piece, which we used as control, is marked by a blue rectangle, (b) Shows the self plot for the larger (gorilla) array.

**1.1.4. SF2.** It can be seen in Fig. N7F that SF2 on the right flank is rearranged with a piece of SF1 inserted in it, which is followed by four macro-repeats (see [54] for macro-repeats definition), each of which has a part made by SF2, a part made by SF1, and one insertion of *Alu*Y. The length of the repeat copy is about 14 kb.

a. Bird eye view

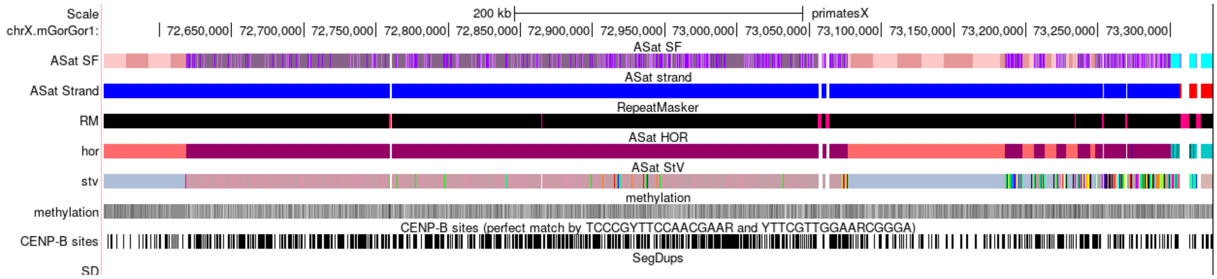

b. Close-up of macro-repeats

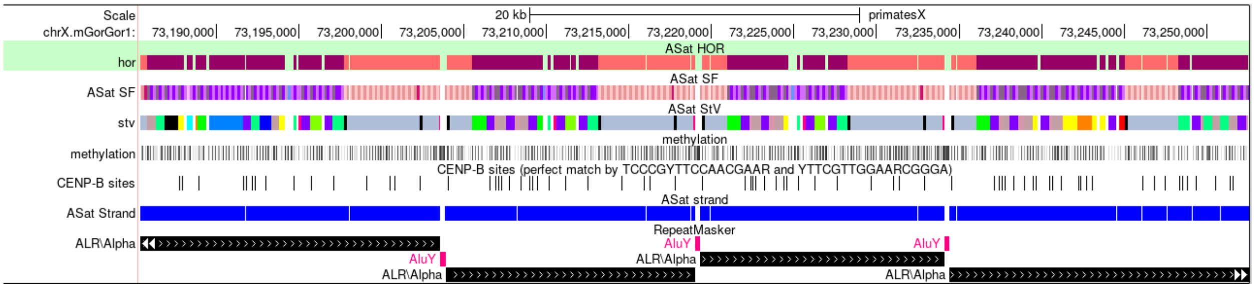

**Figure N7F. The right flank SF2 array in gorilla cenX with SF1 insertion and SF2/SF1/*Alu*Y macro-repeats.** The length of macro-repeats varies likely due to deletions of different parts in individual copies, which occurred after repeat expansion.

**1.1.5. SF1.** SF1 forms the active centromere in gorilla cenX (3.7Mb), which is confirmed by CDR. There seems to be no periodicity in this array other than J1J2 dimers, which is a hallmark of all SF1 sequences. So, the HOR in this case is a dimer, which is only 95% identical to a human dimeric dHOR (S1CMH1d), which is in turn identical to consensus J1J2 dimer derived across all human SF1 HOR consensus sequences. So, these only two known cases of dimeric J1J2 HORs do not appear to be the same.

**1.1.6. HORs and CDRs**

**1.1.6.1. SF5 HORs.** Unidentified minor SF5 HORs include the smaller SF5 HOR arrays, which are not covered in the HOR classification tool (shown in Fig. N7G):

- SF5 11mer, 6 complete copies chrX.mGorGor1:74,034,172-74,048,714
- SF5 16mer, 3 complete copies chrX.mGorGor1:74,094,253-74,103,049
- SF5 13mer, 2 complete copies chrX.mGorGor1:74,324,658-74,339,742

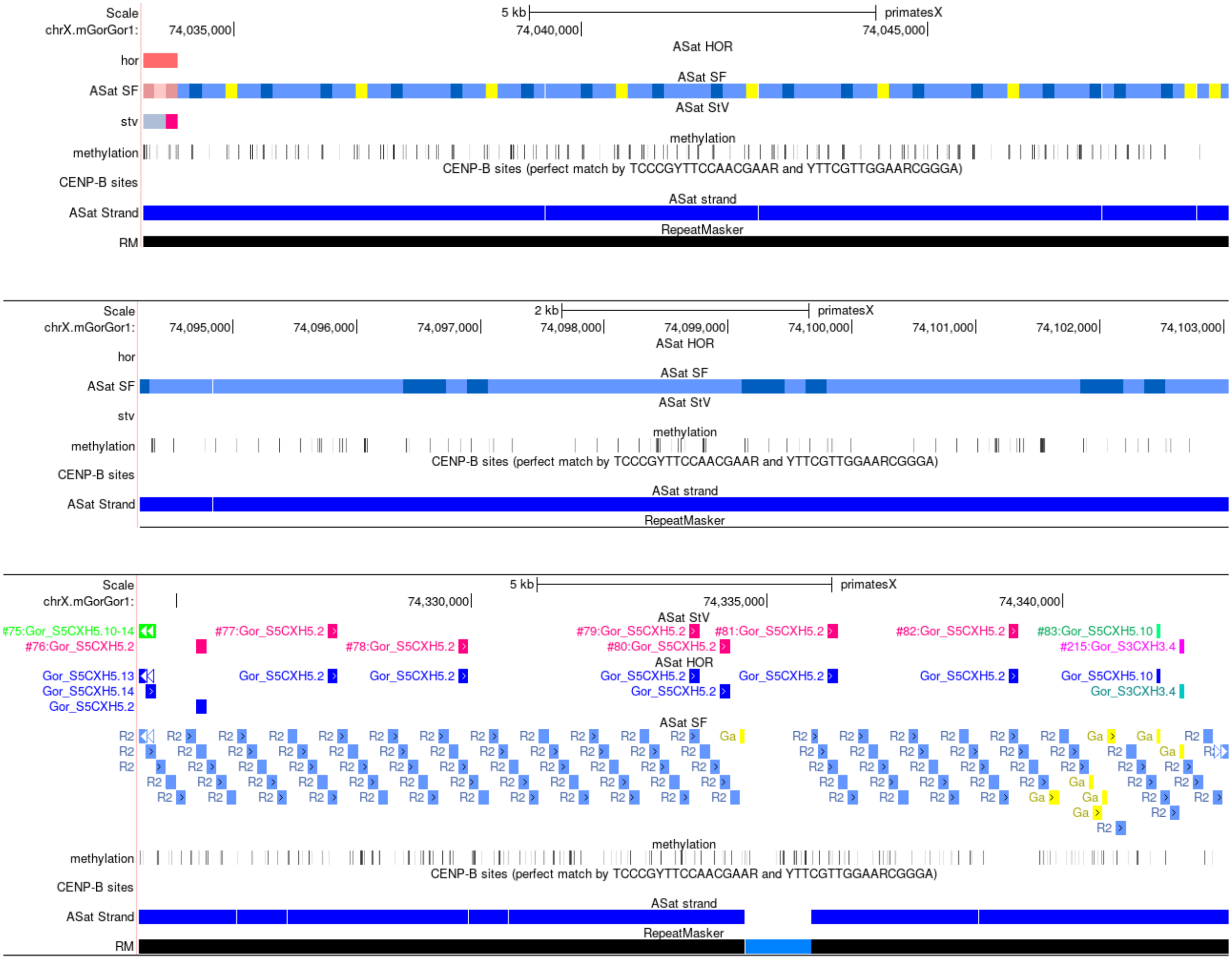

**Figure N7G. Minor HORs not covered by HOR-track are revealed by monomer periodicities in the SF-track or occasional false coverage hits in the HOR-track.** These HOR mini-arrays are located on both sides of the large SF5 HOR array S5CXH5.

**1.1.6.2. SF3 HORs.** In *Homo* and *Pan*, the HOR structures are the same, and gorilla has different structures on Xp and Xq, but they are likely recent amplifications in already dead arrays (pseudocentromeres). The HOR that was active in the gorilla ancestor was likely the same 12-mer as in *Pan* and *Homo*, as it is a common denominator between the two inactive gorilla HORs.

Homo and Pan 12mer W1W2W3W4W3W4W5W1W2W3W4W5 live HORs

Gorilla Xq SF3 19mer W1W2W3W4W3W4W5W1W2W3W4W5W1W2W3W4W3W4W5 dead HOR, q-side
Gorilla cenXp 16mer W1W2W3W4W5W1W2W3W4W3W4W5–W2W3W4W dead HOR, p-side

However, the arrays of such HOR would have their obvious specific signatures if one looks at concatenated arrays. Namely the alternation of 7-mer (green) and 5-mer (gray) components of this ancestral repeat would be 7-5-7-5 in 12-mer array, 7-5-7-7-5-7 in 19-mer array, and 5-7-5-5-7-5 in 16-mer array. So, the presence of doublets 7-7 and 5-5 or their absence would allow us to determine the type of array even in small pieces of ancestral arrays that differed in sequence from the present-day HORs. We have applied this technique to the small SF3 pieces at the flanks of the arrays which had ulcer identity as described below.

P-arm chrX.mGorGor1:68,727,597-68,731,489

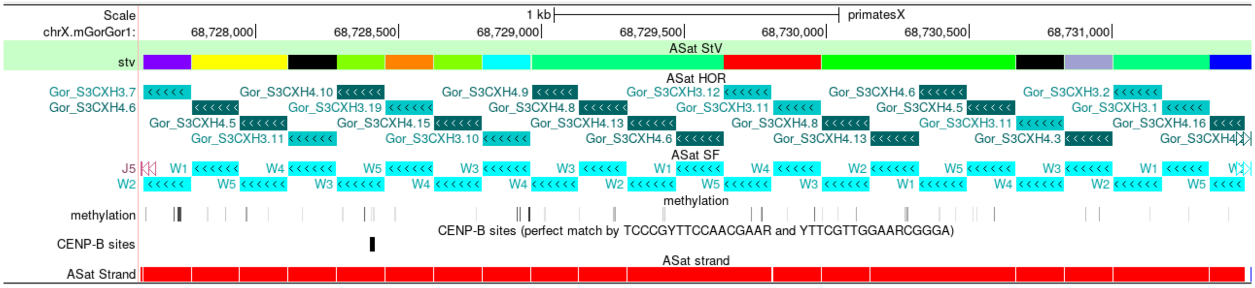

Q-arm chrX.mGorGor1:73,854,486-73,857,805

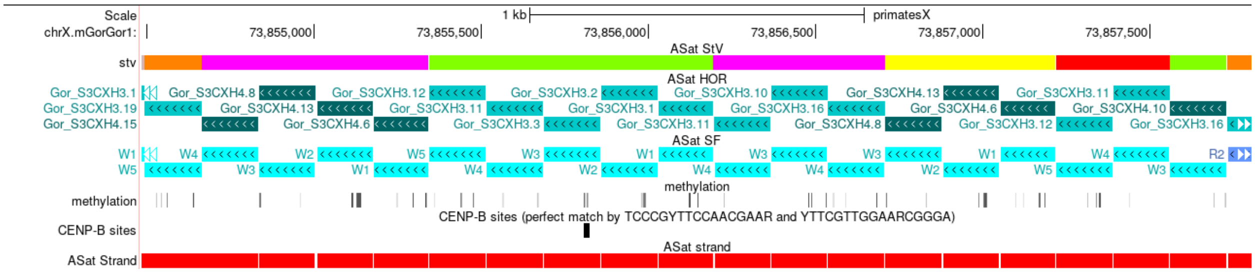

**Figure N7H. The very distal pieces of gorilla cenX SF3 array are similar and composed of S3CXH3 (19mer) and S3CXH4 (16-mer) monomer mix.** Two very small distal SF3 pieces shown in Fig. N7H are in the original reverse orientation and would constitute the edges of the original gorilla SF3 array. Both bear the signature of a 16-mer repeat: 7mer_5mer_5mer (albeit one deleted W5 in the q-arm piece). Note the sparsity of the CENP-B predicted sites. They may be ancestral to the S3CXH4 16-mer pseudocentromere.

The other mixed identity regions match the 12-mer structure where the 7-mer and the 5-mer are alternating and two 5-mers do not go one after another. These regions are as follows:

chrX.mGorGor1:73,300,740-73,306,144

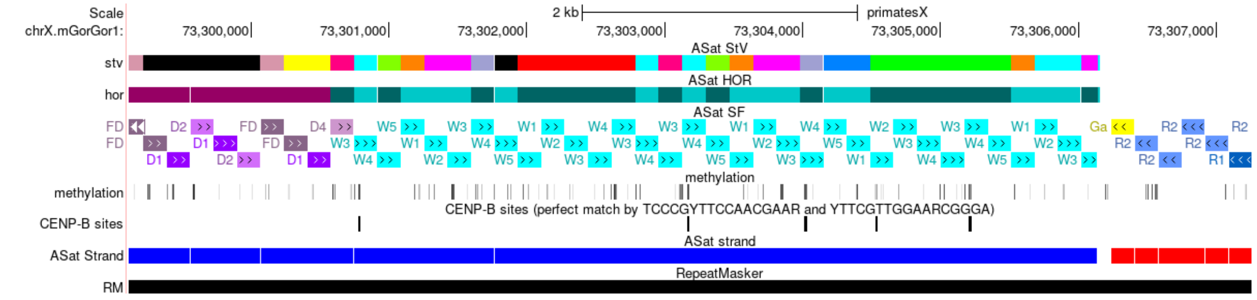

chrX.mGorGor1:73,313,484-73,318,112 (4.6 kb)

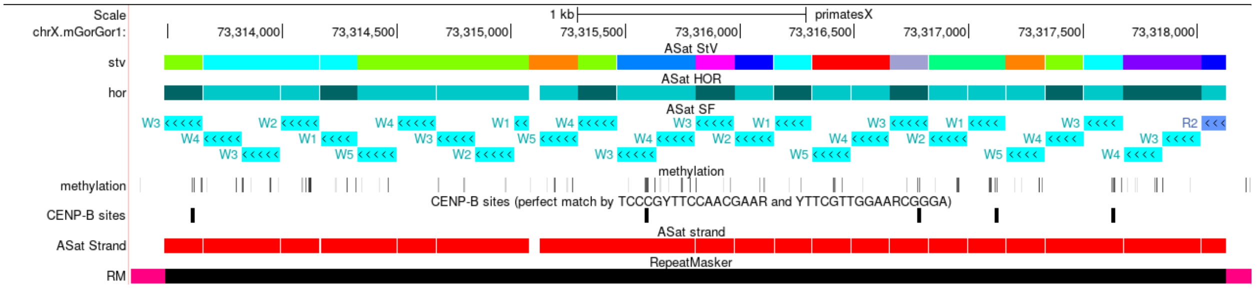

**Figure N7I. Two small regions with unclear identity bear the signature of the 12-mer array.** Two SF3 pieces shown in Fig. N7I are currently adjacent on the q-side and only separated by L1 insertions, but they are in different orientation, so they likely come from different flanks of pre-inversion cenX. The piece shown in  Fig. N7J is on the p-side, but it is in an acquired direct orientation and was brought here from the q-side upon inversion. The piece starts at inversion breakpoint and ends before the W5W2 dimer which is the hallmark of 16-mer HOR.

chrX.mGorGor1:68,731,475-68,737,443 (5.9 kb)

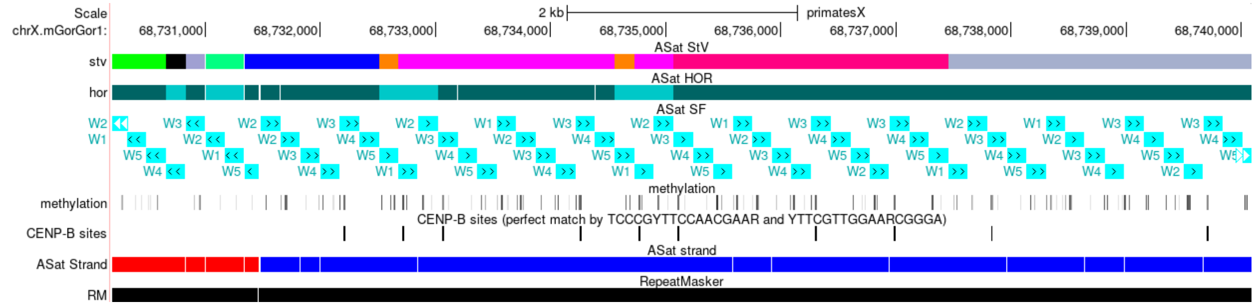

**Figure N7J. One more unclear-identity region bears the signature of the 12-mer array.** Thus, the succession of W monomers in all these regions is compatible with 12-mer (alternating 7-mers and 5-mers) and not compatible with either 16-mer (adjacent 5-mers) or 19-mer (adjacent 7-mers) HORs. We conclude that these are pieces of ancestral 12-mer structure that formed the cenX in human-gorilla common ancestor.

**1.1.6.3. SF2 HORs.** For simplicity we designated the left-flank SF2 array as SF2_1, the larger right-flank SF2 array at  chrX.mGorGor1:72,617,225-73,075,469 (458kb) as SF2_2, and the leftmost small array at chrX.mGorGor1:73,248,237-73,300,236 (52kb) together with the SF2 parts of the macro-repeats as SF2_3. Interestingly, the HOR composition of these arrays separated by SF1 expansions was different. The SF2_1 array consists of a mixture of StVs based on a 7-mer HOR that contains one FD monomer, which is a D1/D2 hybrid and is characteristic of most modern SF2 HORs in humans. The large SF2_2 array consists exclusively of an StV that has a stretch of 2-13 FD monomers (monomer1 of the HOR) in tandem followed by a D1D2 dimer (monomers 2 and 5 of the HOR, respectively). So, the StV formulae in this array is (1)n-2_5 where n=2-13. Finally the SF2_3 array is similar to SF2_1 and consists of various StVs in which all monomers of the HOR are present.

**1.1.6.4. SF1 HOR.** There seems to be no periodicity in this array other than J1J2 dimers, which is a hallmark of all SF1 sequences. So, the HOR in this case is a dimer, which is 95% identical to human S1CMH1d, which is in turn identical to a consensus J1J2 dimer. Note that an even smaller (4.2-kb) isolated piece of SF1 HOR array has been inserted in SF5 on the right flank at chrX.mGorGor1:74,029,936-74,034,171.

**1.1.6.5. HOR homogeneity.** The SF3 HORs look very regular (in terms of repeat periodicity) as opposed to the other HORs in gorilla centromere. This suggests that they are very recent amplifications of the dead SF3 material which featured ruined HOR structure (disrupted HORs), but the regularity was reinstated by a secondary amplification which has brought about a different HOR (e.g. 19mer instead of more ancestral 12mer). This dead material has survived mostly as small islets of diverged 16-mer and 12-mers mentioned above. These pieces are likely relic centromere pieces (dHORs) and the SF3 HORs are likely pseudocentromeres.

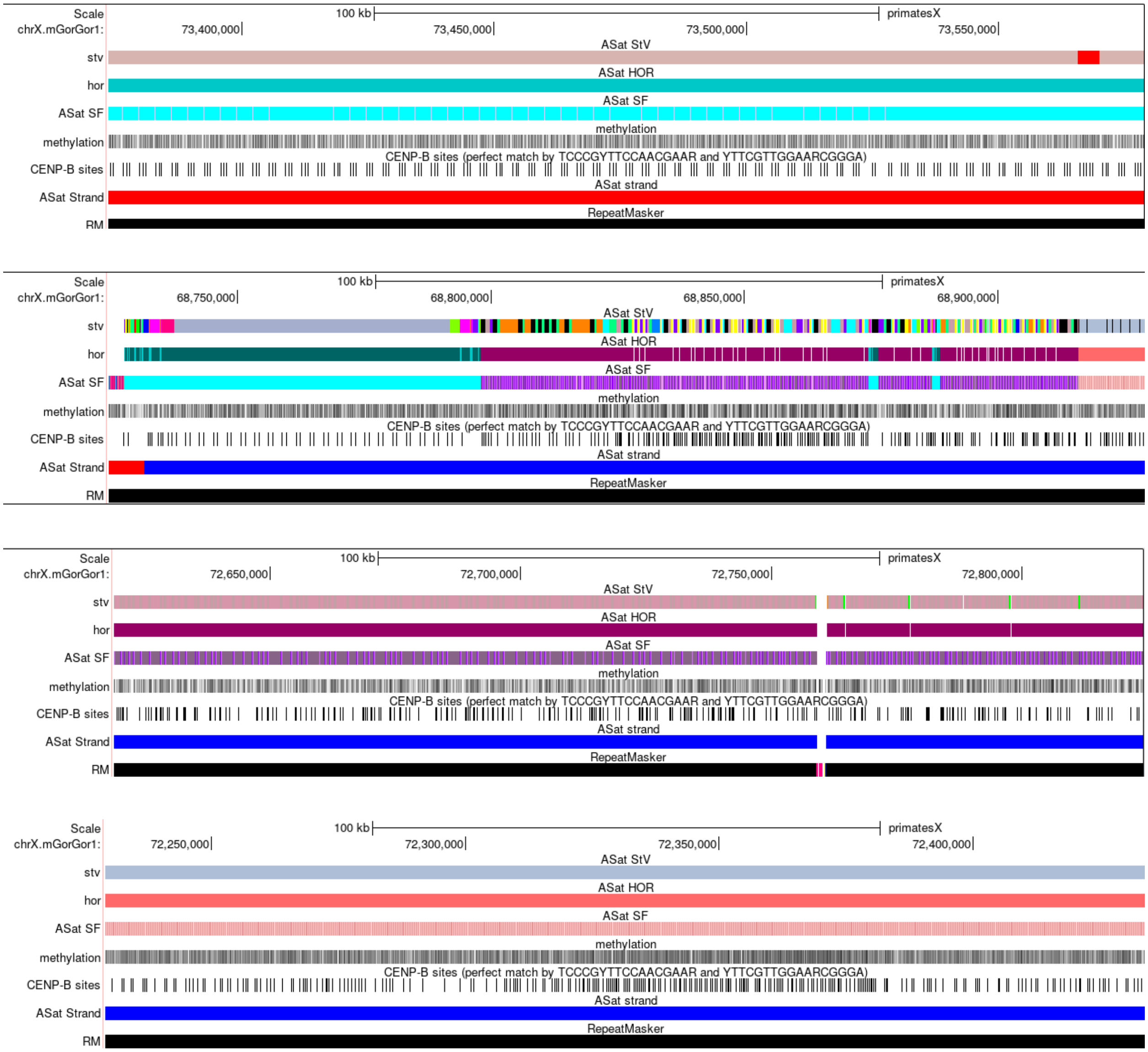

**Figure N7K. A striking regularity of CENP-B predicted sites in presumed recent SF3 pseudocentromere (S3CXH3) outmatches inactive SF2 arrays and an even active SF1 array.** It is likely just the sign of a more recent amplification.

**1.1.3. TE insertions.** Besides numerous and trivial TE insertions of various ages, which are common in monomeric AS regions[54,113], we observed some rarer cases in the inactive SF3, SF2 and SF01 arrays. Such occasions are rare because these are the younger arrays formed by new family repeats, so that they have been exposed to L1 insertions only for a short time since their emergence, and are not supposed to have TEs that are older than the arrays themselves. Thus, for gorilla, only L1PA2 and L1PA3 human-shared L1 elements are expected in new family arrays. The data were consistent with this expectation. The annotated list of insertions in the new arrays is as follows: (1) L1PA2 with the pieces of some other associated repeats at chrX.mGorGor1:72,758,934-72,760,908, perhaps brought into this location together with the L1PA2 element; (2) two closely spaced L1PA2 insertions at chrX.mGorGor1:73,056,190-73,066,124, which look like a duplication of an L1 insertion together with an AS piece; (3) three copies of *Alu*Y repeat in three SF2/SF1 super-HORs (one per repeat); (4) two closely spaced L1PA2 elements at inversion breakpoint, which do not look like a duplication, at chrX.mGorGor1:73,306,154-73,321,068. In the latter site, one L1PA2 element has a small piece of an AS unrelated to the context attached (seven monomers of SF5 in the SF3 array); this piece may have been transduced from the previous L1 location; and (5) finally, a small piece of L1PA2 element (300 bp) is located on the SF3/SF01 border at chrX.mGorGor1:73,857,808-73,858,107. It probably represents the remains of a largely deleted element that is unique to gorilla. Neither human nor *Pan* have an L1 at this location.

Two unusual L1PA2 elements are shown below:

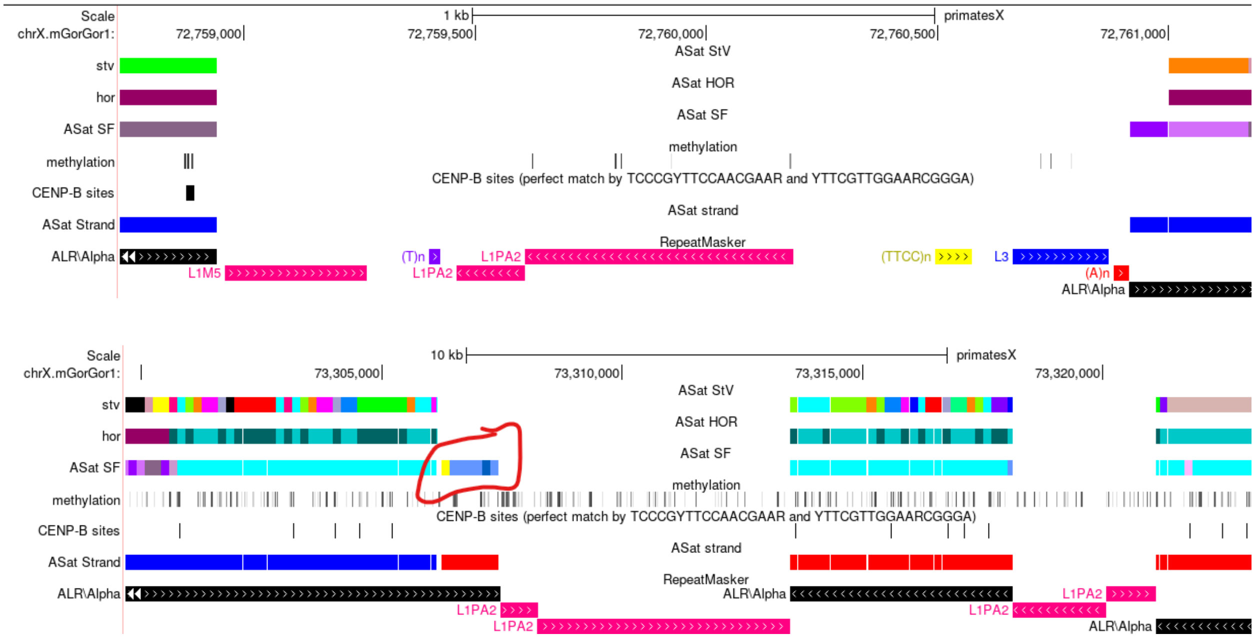

**Figure N7L. Two unusual L1PA2 elements in gorilla cenX AS.** Upper panel, L1PA2 with the pieces of some other associated repeats at chrX.mGorGor1:72,758,934-72,760,908 (described in the text). Lower panel, two closely spaced L1PA2 elements at an inversion breakpoint at chrX.mGorGor1:73,306,154-73,321,068 (described in the text. An unrelated AS piece possibly transduced from the previous TE location is indicated.

##### 1.2. cenY.

The centromeric region of the gorilla, designated as cenY, is constituted by SF1 HORs, which exhibit no relation to the SF4 HORs that typify the cenYs in both *Homo* and *Pan* genera. It is important to underscore that the SF4 HORs in *Homo* and *Pan*, despite sharing the same family designation, do not exhibit a close genetic relationship. Their similarity is comparable to that of any two arbitrarily chosen SF4 monomeric sequences, characterized by an absence of co-linearity and an average divergence of approximately 16% between monomers. Consequently, this significant genetic disparity precludes the possibility of conducting meaningful comparative analyses across these lineages.

The SF1 centromere in gorillas is devoid of older structural layers, which may imply recent centromere repositioning. This inference is supported by evidence that the centromere was inserted and subsequently expanded within a symmetrical segmental duplication (SD), which encompasses fragments of orange (SF10) and red (SF9) AS, as illustrated in Figure N7M. This pattern of segmental duplications, containing interspersed

AS, is a recurrent feature observed across ape genomes, indicating a common evolutionary mechanism in centromere evolution within this clade.

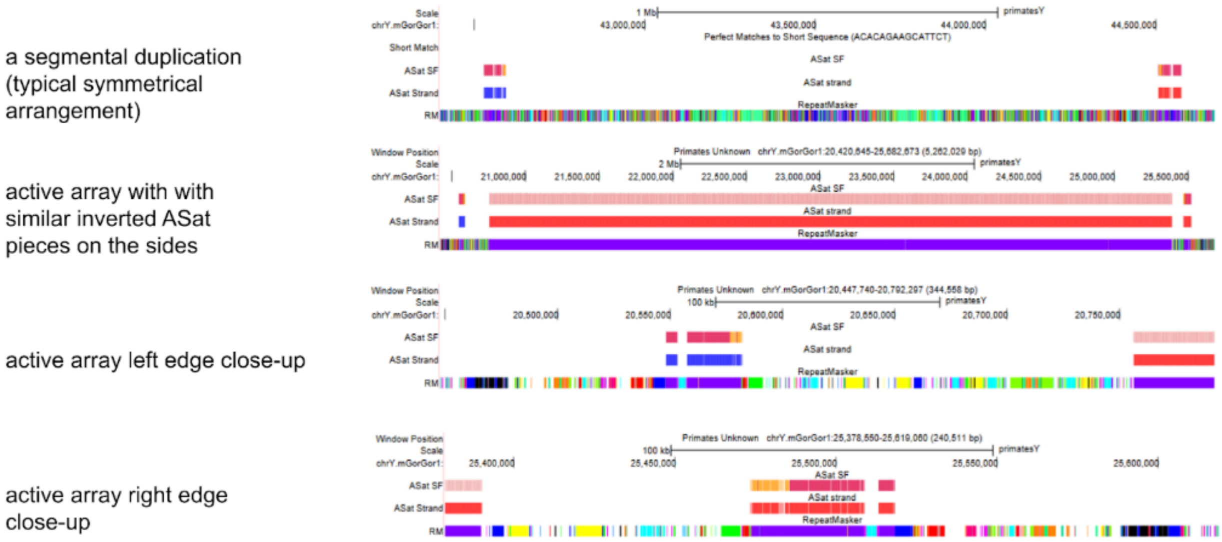

**Fig. N7M. SF1 centromere in gorilla cenY was inserted and expanded in palindromic AS-containing segmental duplication (SD).** In the top panel, an example of intact gorilla palindromic SD is shown. Two pieces of ancient red and orange AS (SF9 and SF10) can be seen in the SF-track on the flanks of the SD. The strand track shows that these pieces are in the opposite orientations which testifies to palindromic organization. The RepeatMasker track shows a symmetrical succession of other variously colored repeated elements around the center of the SD. A similarly organized red/orange AS-containing palindromic SD has been described in the human Y[105], others are present in chimpanzee and bonobo and are massively amplified in both orangutan species (Fig. S15A). The second panel shows the SF1 active array of cenY and two nearly symmetrical pieces of red/orange AS in its flanks. The bottom two panels show close-ups of the two flanks. The imperfectly symmetrical arrangement of RepeatMasker colored elements around the centromere can be observed which demonstrates that the active array has been inserted into a palindromic SD.

##### 1.2.1 HOR arrays, TE insertions, and frequency of CENP-B sites

The characterization of HORs within the gorilla cenY has been conducted with a focus primarily on identifying the major components, while smaller segments possessing more intricate structures at the array boundaries were not included. Furthermore, there was no attempt to refine the initiation site of HORs to simplify the complexity inherent in StV annotation. The beginning of the array at chromosome location chrY.mGorGor1:20,756,444-20,838,163 might hold unidentified sequences, suggesting these segments could be valuable targets for further analyses in future studies. The gorilla cenY comprises a minimum of five distinct HOR arrays as detailed in Table S29, among which the 18-mer S1CYH1L HOR constitutes the primary active array, spanning approximately 3.5 megabases. Additionally, smaller arrays are situated on the centromere's right flank, as depicted in Figure N7N. These arrays exhibit variable densities of functional CENP-B sites, with the active array displaying the highest density—six sites per 18-mer HOR, meaning that 6 out of 9 J2 monomers possess the site. The densities in other arrays decrease progressively: S1CYH5 (cyan) at 2/12, S1CYH2 (green) at 1/20, S1CYH3 (blue) at 2/20 or 3/20, and S1CYH4 (pink) at 1/20. Additionally, a solitary insertion of L1PA2 was identified within gorilla cenY, further detailed in Figure N7N, highlighting the genomic complexity and diversity within this centromeric region.

A

| HOR | CENP-B sites | Basic HOR length (mon) | Array size | Dimeric structure |
|---|---|---|---|---|
| Gor_S1CYH1L | yes | 18 | 3500 kb | 1-18 (J1J2)9 |
| Gor_S1CYH2 | yes | 20 | 520 kb | 1-20 (J1J2)10 |
| Gor_S1CYH3 | yes | 20 | 248 kb | 1-20 (J1J2)10 |
| Gor_S1CYH4 | yes | 20 | 150 kb | 1-20 (J1J2)10 |
| Gor_S1CYH5 | yes | 12 | 33 kb | 1-12 (J1J2)6 |

B

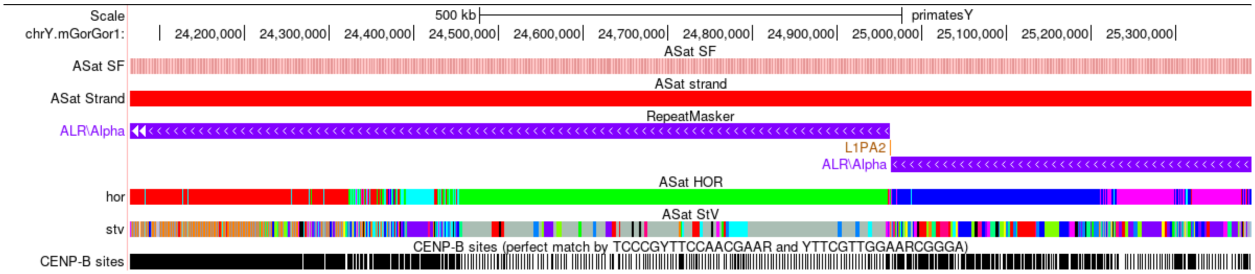

**Figure N7N. Five loosely related HOR arrays in gorilla cenY. (A)** Shows the list of the HORs with "L" in the name indicating the active array, and lengths in monomers and numbers of J1J2 dimers in a HOR shown. **(B)** Browser panels showing multiple HOR arrays in the right flank of the centromere. Shows the region overlapping the right end of the active array (red) and going through the smaller arrays till the end of the AS array. Note L1PA2 insertion in RepeatMasker track, the variously colored arrays in the HOR track (color code is given in (A)), and different density of CENP-B sites shown in the bottom track. Note that Gor_S1CYH3 (blue) array is mostly composed of various StVs, a small island of full-length HORs is located at chrY.mGorGor1:25,114,591-25,135,611.

While certain monomers across different HORs seem to share a common ancestry (e.g. monomers S1CYH1L.17, S1CYH5.5, and S1CYH5.9, all feature an identical 4bp insertion post position 95) the HORs do not exhibit collinearity and cannot be considered sister HORs. Their relationship is not immediately apparent, aside from their composition, which includes J1J2 dimers characteristic of the SF1 signature.

#### 2. Active HOR comparisons in closely related species of *Pan* and *Pongo*

**2.1. Summary.** The centromeres in the twin species are made by the same HORs, but by no means could be seen as "the same" arrays that evolved separately for a given time (0.5 MY for orangutans and over 1-2 MY for *Pan*) by accumulating random mutations. Even in the closest arrays, the HORs do not mix on the trees, but make up either separate branches or separate sectors in the fan-like tree shapes (i.e. each species is represented by its own species-specific HOR haplotypes). Thus, since separation of species, one or both respective arrays have undergone cycles of re-modelling that replaced the bulk of the active array. In other words, almost all HOR copies are different between species, and the differences are the same in all HORs (i.e. species-specific). Moreover, in most cases, the change was achieved not in one but in two (or more) discrete cycles, and these previous-generation dead HORhap arrays sit at the flanks of the current live ones. Finally, only the oldest HORhaps represented by only a few copies, which sit on the periphery of the arrays or often at their very tips, are not species-specific or at least much less so. Their HORhap consensus sequences are near equidistant from both species' active array HORhaps. In one case, where such HORs have survived in both species (*Pan* cenX) they mix in the same cluster of branches in the HOR tree. These HORs are presumably the surviving copies that represent the centromeres of the common ancestors of the twin species.

**2.2. Evolutionary scenarios.** The main feature of AS evolution is the layered expansion process that leads to the "expanding centromere" model, where the centromere constantly grows in the middle and shrinks at the periphery[54,114]. In the simplest situation, the growing core is at any one moment formed by just one AS variant that expands rapidly. Once in a while, the identity of the growing core changes and an expansion of a new variant starts. Typically (but not necessarily), the seed of a new growth is located within the old growing array, so that the latter is split and displaced to the flanks, where it no longer grows, and starts shrinking. This complex scenario of compartmentalized expansion/contraction was proposed instead of the older notion of stochastic expansion/contraction process, to explain the complex layered structure of primate centromeres, which was revealed in the last two decades. As layered expansion is such a well-structured process, it is unlikely to be driven purely by stochastic recombination processes, therefore a “kinetochore selection” hypothesis was proposed, which explains it by egotistic selection drive, where the kinetochore acts as an amplification machine and drives expansion of a satellite variant currently covered by CENP-A, which primarily determines location of the other kinetochore proteins and of various recombination proteins that might be associated with it[54,114].

**2.3. Methods.** Given the set of tracks we have developed for each assembly, we have endeavored to produce the following five elements to perform the detailed analysis of AS in twin species. The StV track was used to collect the StV statistics (1) and determine the number of full-length HORs, or the StVs with duplicated monomers, which could easily be converted to full-length. Such StVs were extracted, aligned, and used to build the HOR tree (2), the branches of which were used to build Multiple Sequence Alignments (MSAs), which then became HMMs and were used in the HMMER-based tool to classify HORs in chromosome assemblies and build the HORhap annotations (3) of complete centromeres. Such annotations were visualized as the UCSC Browser tracks and examined manually to establish the HORhap regionalization. The HORs in the MSAs were also compared to each other and average degree of intra-array divergence was determined (4). Lastly, the MSA were used to derive consensus sequences for each HORhap and build the HORhap consensus tree (5). Below we provide detailed examples of how we used these elements to perform the analysis and arrive at conclusions stated in the main text, Ext. Data Fig. 4 and Fig. S15. Below we provide figures that are more detailed than those in the main text, or present additional information. Note that all HORs are numbered in StV tracks from left to right, and we often use these numbers to show locations of HORs within an array instead of coordinates.

##### 2.4. Chimpanzee/bonobo

###### 2.4.1 cenX

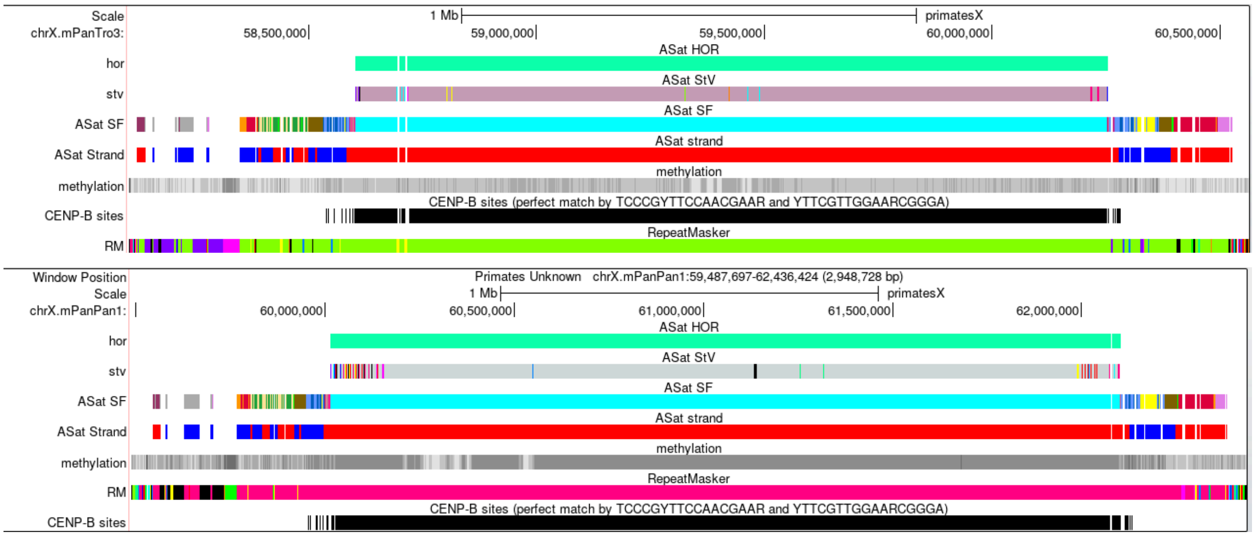

**Figure N7O Centromere landscapes in chimpanzee and bonobo as revealed by the Browser tracks.**

Figure N7O shows the stable flank patterns of monomeric layers in the SF-track and of inversions in the strand-track outside area covered by the active HOR (HOR-track), and predicted CENP-B sites are abundant in HOR arrays of both species. However, there are notable differences in the HOR array: (1) two closely spaced insertions of L1Pt elements at the left flank in chimpanzee, which are absent in bonobo (two yellow bands in RepMask track, and the gaps in all other tracks), note that the RM (RepeatMasker) track uses random colors, so the same repeated elements appear in different colors in different assemblies; (2) CDR (a series of hypomethylated bands that overlaps the HOR array indicating the kinetochore position[54] within it) is located in the middle of an HOR array in chimpanzee and at the left flank in bonobo; (3) both arrays are formed mostly by a full-length 12-mer SF3 (cyan in the SF-track) HOR shown by different shades of gray in the two species. StVs with altered monomer order (other colors) are present mostly at array ends, but their exact patterns are distinctly different.

**Table N7A. StV content in *Pan* twin species cenX active HOR arrays.**

| StV (Chimpanzee) | cnt |
|---|---|
| Pan_S3CXH1L.12-1 | 783 |
| Pan_S3CXH1L.12-5_2-1 | 4 |
| Pan_S3CXH1L.12-9_2-1 | 2 |
| Pan_S3CXH1L.12-7 | 2 |
| Pan_S3CXH1L.12-5 | 2 |
| Pan_S3CXH1L.12-3_11-1 | 2 |
| Pan_S3CXH1L.12-3_1 | 2 |
| Pan_S3CXH1L.9-7_10-9_5-4_5-1 | 1 |
| Pan_S3CXH1L.7-3_9-1 | 1 |
| Pan_S3CXH1L.7_11-7 | 1 |
| Pan_S3CXH1L.7-1 | 1 |
| Pan_S3CXH1L.12-8 | 1 |
| Pan_S3CXH1L.12-7_11-1 | 1 |
| Pan_S3CXH1L.12-11_5-4_10-1 | 1 |
| Pan_S3CXH1L.12-11_2-1 | 1 |
| Pan_S3CXH1L.12-10_2-1 | 1 |

| StV (Bonobo) | cnt |
|---|---|
| Pan_S3CXH1L.12-1 | 966 |
| Pan_S3CXH1L.12-9_7-1 | 8 |
| Pan_S3CXH1L.12-8 | 7 |
| Pan_S3CXH1L.12-6_2-1 | 7 |
| Pan_S3CXH1L.12-4_1 | 6 |
| Pan_S3CXH1L.12-10_6-1 | 6 |
| Pan_S3CXH1L.7-2_8-1 | 5 |
| Pan_S3CXH1L.12-10_2-1 | 4 |
| Pan_S3CXH1L.7-1 | 3 |
| Pan_S3CXH1L.12-9_1 | 2 |
| Pan_S3CXH1L.12-8_4-1 | 2 |
| Pan_S3CXH1L.12-3_11-1 | 2 |
| Pan_S3CXH1L.9-7_10-9_5-4_5-4 | 1 |
| Pan_S3CXH1L.6-1 | 1 |
| Pan_S3CXH1L.4-1 | 1 |
| Pan_S3CXH1L.12-9_5-4_5-1 | 1 |
| Pan_S3CXH1L.12-6 | 1 |
| Pan_S3CXH1L.12-3_4-1 | 1 |
| Pan_S3CXH1L.12-2_8-2_8-7_11-7 | 1 |
| Pan_S3CXH1L.12-2_8-2_8-1 | 1 |
| Pan_S3CXH1L.12-2_8-1 | 1 |

StVs. Both species are similar in having almost exclusively full-length HORs (Table N7A; shown as S3CXH1L12-1 (not 1-12), because AS is in reverse orientation, and the monomer order in the StV-tracks and StV table is also inverted). However, the rare other StVs are different in two species. All StVs with copy numbers more than one are shown in the tables (full data in Table S30).

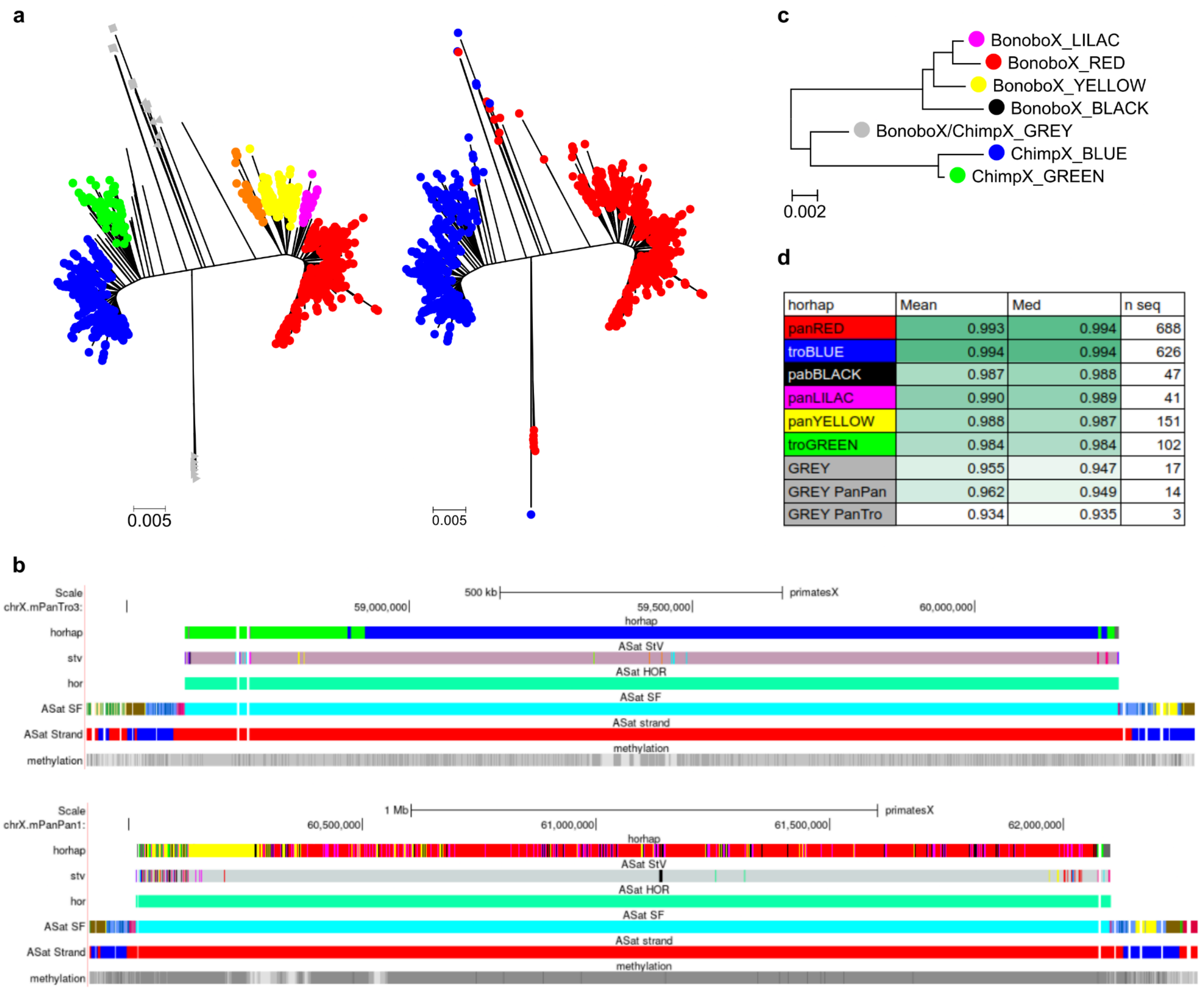

| horhap | Mean | Med | n seq |
|---|---|---|---|
| panRED | 0.993 | 0.994 | 688 |
| troBLUE | 0.994 | 0.994 | 626 |
| pabBLACK | 0.987 | 0.988 | 47 |
| panLILAC | 0.990 | 0.989 | 41 |
| panYELLOW | 0.988 | 0.987 | 151 |
| troGREEN | 0.984 | 0.984 | 102 |
| GREY | 0.955 | 0.947 | 17 |
| GREY PanPan | 0.962 | 0.949 | 14 |
| GREY PanTro | 0.934 | 0.935 | 3 |

**Figure N7P. HOR tree (A), HORhap Browser tracks (B), HORhap consensus tree (C) and intra-array divergence (D) for S3CXH1L HOR in two *Pan* species.** See comments in the text.

HORhaps. Panel A in Figure N7P shows a minimum evolution HOR-tree of 300 random full-length HORs (Pan_S3CXH1L.12-1) from each of the two species, colored in two different ways. On the right tree, robust branches (marked by different colors) were used to generate MSAs further used as HMMs (Hidden Markov Models) in the HORhap annotation tool and consensus HORhap sequences (each colored branch treated as a different HORhap). The average length of the twigs in the HOR tree likely indicates the so-called “amplification age” of the HORhaps. More numerous red and blue HORs have on average shorter twigs, which likely indicates less intra-array divergence and more recent amplification. Other branches (except the grey one) have longer twigs, which suggests an older amplification. The grey, loose cluster of branches features the longest twigs, which indicates that these HORs resulted from yet older amplification events. The right tree is colored by species; the chimpanzee HORs are blue and the bonobo HORs are red. The grey cluster is the only zone in which the sequences of both species mix. Thus, the GREY HORhap likely represents the remnants of the centromere of the common ancestor of both species. There are 14 full-length HOR copies in bonobo and three in chimpanzee, all located at the very right tips of respective arrays.

Fig. N7P panel B shows the regionalization of the HORhaps identified. In chimpanzee, the blue HORhap is active as it overlaps with CDR. It is characteristically large and occupies a more central position. The green HORhap is located on the flank and is much smaller. The layered expansion model[54] predicts that it represents an older HOR generation, which was likely active before the expansion of the blue HORhap and is unlikely to be active now. In bonobo, the large red and the small yellow seem to play similar roles. However, it is not fully confirmed by the CDR position (because it partially overlaps the yellow HORhap). Note that the black and lilac

HORhaps are not well regionalized, these HORs are always interspersed with the red HORhap, so they are harder to interpret.

Bonobo does have few full-length copies of green HORs characteristic of the older array in chimpanzee. Two copies at the very right tip, interspersed with grey, and few copies at the left tip (Fig. N7Q). However all but one copies at the latter location are parts of the macro-repeat which also includes some full-length yellow HORs and incomplete lilac and grey HORs (which therefore may be misclassified), so it is likely there are only two independent copies. That gives just four  independent copies altogether, two at each side. It is not clear what is the significance of these. If taken at face value, they may indicate the possibility that bonobo once also had green arrays in the centromeres which were distal to grey arrays.

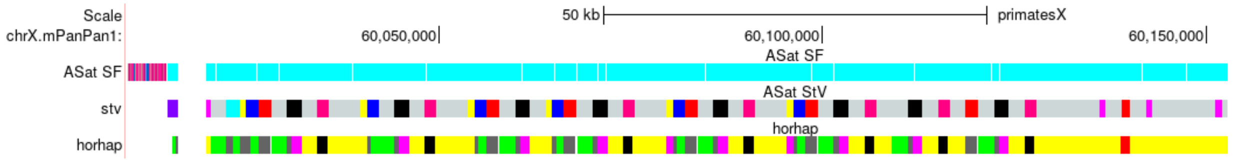

**Fig. N7Q. Green HORs and macro-repeats at the very left tip of the bonobo cenX array.** The HORhap assignments are shown in the HORhap track. Eight copies of characteristic macro-repeat can be seen (some partially deleted). Therefore, eight full-length green HORs in the region actually represent one original copy which was recently amplified in a macro-repeat. In StV track, the grey segments are formed by the full-length HORs, and align with some geen HORhap segments indicating full-length green HORs. Note that HORhap assignments, which are not full-length, especially the shorter ones may result from misclassification, as the HORhap HMMs are full-length.

Fig. N7P panel C shows the consensus HORhap tree which usually displays phylogenetic relationships better than the HOR tree, as it indicates the so-called "phylogenetic age" of the HORhaps that shows how derived the sequences are (i.e. what are their relative distances from the root of the tree). The grey branch is likely to represent a common ancestor (of the two species as well as of the HORs involved), as it is close to the root. Red and blue HORhaps are the most derived and phylogenetically youngest of all, as their branching points are the farthest from the root in respective branches. Yellow and green HORhaps likely represent the older generations of HORs. Additionally, in panel D, the mean and median values for intra-array identity are calculated from MSAs used for HMMs that quantify the "amplification age". The layered expansion model calls for the highest identity in the active arrays (red and blue), somewhat lower identity in the medium-age HORhaps (green and yellow), and the lowest in grey. Consistent with these predictions, green is indeed markedly more divergent than blue, and yellow is slightly more divergent than red. Divergence in grey is indeed by far the highest. As grey MSA contained the HORs from two different species (and assemblies), we also calculated the average identity separately for each species. The grey HORhap is still by far the most divergent.

**2.4.2. Patterns of AS-containing SDs in the arms of chromosome Xs in chimpanzee and bonobo.** The patterns of AS-containing SDs are very different between the two *Pan* species (Fig. S15A). Bonobo has a simple pattern, where only the SDs, which have Ga (yellow) and Ha (brown) monomers are present on the p-side, while only the SDs, which have the Ca (red) and Ba (orange) monomers are present on the q-side. Judging by the AS patterns all yellow/brown SDs are likely to be the offspring of the same single SD that was first multiplied in several steps, which included inversions to form a group of six SDs, and this block was duplicated in direct orientation to yield a pattern of 12 yellow/brown SDs on the short arm. The same is true for the red/orange SDs that contain the same AS piece in both direct and reverse orientations. The AS part of this SD is the same as the one we have previously described in humans[105].

**2.4.3 cenY.**

CenY in chimpanzee assembly is inverted relative to bonobo and humans (Fig. S15A). Whether this should be considered an inversion or the whole chromosome is flipped because the short arm has become longer than the long arm due to repeat expansions is hard to ascertain. In any case, inversion breakpoints are outside the AS. Flanking AS regions are small and contain yellow (SF4) and brown (SF6) monomeric layers and are mostly SDs (Fig. 5B, main text).

StVs. The active HOR in both species is an SF4 29mer S4CYH1L. Chimpanzee cenY active array is small (1.2Mb) and has a lot of full-length HORs. The bulk of the array is mostly formed by irregular alternation of 1-29 and 1-17_17-29 (a duplication of mon17) StVs (Table N6B). The CDR overlaps a cluster of 1-3_10-29 StVs all of which are located in this region.

**Table N7B. StV content in *Pan* twin species cenY active HOR arrays.** Full data are shown in Table S30.

| StV (Chimpanzee) | cnt |
|---|---|
| Pan_S4CYH1L.1-29 | 130 |
| Pan_S4CYH1L.1-17_17-29 | 62 |
| Pan_S4CYH1L.1-17_19-29 | 24 |
| Pan_S4CYH1L.1-3_10-29 | 11 |
| Pan_S4CYH1L.1-18_17-29 | 10 |
| Pan_S4CYH1L.1-9_12-18_17-29 | 4 |
| Pan_S4CYH1L.1-18_17-18_21-29 | 3 |
| Pan_S4CYH1L.17-29 | 2 |
| Pan_S4CYH1L.1-5_4-29 | 2 |
| Pan_S4CYH1L.1-4_12-29 | 2 |
| Pan_S4CYH1L.1-18_10-18_10-17_17-29 | 2 |
| Pan_S4CYH1L.1-12_27-29 | 2 |
| Pan_S4CYH1L.26-29 | 1 |
| Pan_S4CYH1L.2-6_14-20 | 1 |
| Pan_S4CYH1L.26 | 1 |
| Pan_S4CYH1L.23 | 1 |
| Pan_S4CYH1L.2-29 | 1 |
| Pan_S4CYH1L.1-9_12-29 | 1 |
| Pan_S4CYH1L.17 | 1 |
| Pan_S4CYH1L.1-3_27-29 | 1 |
| Pan_S4CYH1L.1-20 | 1 |
| Pan_S4CYH1L.1_17-29 | 1 |

| StV (Bonobo) | cnt |
|---|---|
| Pan_S4CYH1L.29-14_6-2 | 133 |
| Pan_S4CYH1L.29-18_3-2 | 100 |
| Pan_S4CYH1L.29-1 | 92 |
| Pan_S4CYH1L.29-18 | 83 |
| Pan_S4CYH1L.29-11_3-1 | 79 |
| Pan_S4CYH1L.29-21_19-18_3-2 | 62 |
| Pan_S4CYH1L.29-17 | 49 |
| Pan_S4CYH1L.29-12_12-1 | 33 |
| Pan_S4CYH1L.29-12_22-1 | 25 |
| Pan_S4CYH1L.29-19_17-11_3-1 | 19 |
| Pan_S4CYH1L.29-21_3-1 | 15 |
| Pan_S4CYH1L.29-22_3-1 | 13 |
| Pan_S4CYH1L.29-22 | 7 |
| Pan_S4CYH1L.29-27_22-14_6-2 | 6 |
| Pan_S4CYH1L.29-21 | 5 |
| Pan_S4CYH1L.29-14_12-1 | 5 |
| Pan_S4CYH1L.29-12_22-12_22-1 | 5 |
| Pan_S4CYH1L.29-18_14-1 | 4 |
| Pan_S4CYH1L.29-16 | 4 |
| Pan_S4CYH1L.29-14_6-2_16-14_6-2_16-14_6-2 | 4 |
| Pan_S4CYH1L.25-18_3-2 | 4 |
| Pan_S4CYH1L.22-1 | 4 |

Bonobo active array is 3.7 Mb long, has relatively few full-length HORs and no StVs with duplication of mon17 (Table N7B). The 0.8Mb array at 32,603,841-33,438,639 is formed by two long concatemeric StVs (with an islet in between) made exclusively of monomers 12-22 (the segment duplicated in green StV in adjacent region on the right; see below). There are just three full-length HORs in the left 2Mb of the array (#466, 467, 472, right before and between the 2 large blocks of 12mer StV). In the right half,  there is a significant number of 1-29 and 1-12_12-29 StVs.

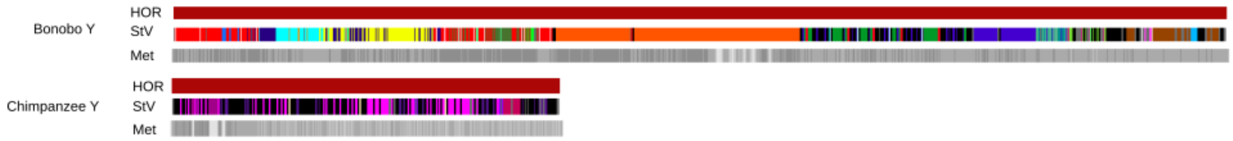

**Fig. N7R. StV patterns and kinetochore positions in cenY are dramatically different between chimpanzee and bonobo.** The HOR, StV and methylation cenY tracks are shown for the two Pan species. The same 29mer HOR shared by both is shown by the brown color in the HOR-track, but the CDRs show dramatically different kinetochore positions (Met-track, middle in one and left flank in the other). The StV track shows a huge expansion of a novel shortened 12mer HOR (monomers 12-22 of the 29mer; marked by orange color) in bonobo, which is absent in chimpanzee and might have caused the shift in kinetochore position. Full-length HORs are shown in black in StV-tracks. One can see that it constitutes about a half of the HORs in chimpanzee and appears only in the right half of the array in bonobo. Some structure suggesting black/lilac macro-repeat is seen in the chimpanzee array.

The detailed list below shows different distinct bonobo cenY regions (from left to right) shown in the StV track in Fig. N7R. It features the color(s) in the StV track, coordinates of each region, size, major StVs, and sometimes comments on obvious other features. The list shows a complex history of local amplifications not shared with chimpanzee cenY.

1. Red, dark blue, cyan, dark blue, yellow, red mixed with dark green, chrY.mPanPan1:31,298,693-32,597,917 (1.2Mb), complex patchwork of arrays each dominated by a particular StV (hence the colors) all of which lack monomer 1 and have various other deletions. In general, the coloring in this region shows the HOR end StVs like 18-29, 17-29 and 16-29 in dark colors and longer StVs in bright colors. The following detailed example shows the monomeric structure of dominating HOR, region and color in parentheses (numbers of HORs according to the StV track) and dominating color: 2-6_14-29 (#1-80, red), 18-29 (#80-105, dark blue), 2-3_18-19_21-29 (#106-169, cyan), mix of the above (#170-214), mix of 2-3_18-29 (yellow) and 18-29(dark blue) (#215-341), 2-6_14-29 (red) with patches of 17-29 (dark green) (#342-465). These arrays are sprinkled with some other, less frequent StVs, often in patterns indicative of local low-copy super-HOR repeats.
2. Orange, chrY.mPanPan1:32,603,841-33,438,639 (0.8Mb), two long concatemeric StVs (#468 and #473 with an islet in between) made exclusively of StV 12-22 (the segment which is also duplicated in green StV in the adjacent region to the right).

   The following regions are all dominated by StVs which have monomer 1.

3. Black, green, chrY.mPanPan1:33,475,354-34,034,571: 1-29 (black, full-length) and 1-22_12-29 (green, dup12-22 same as the major 12mer concatemer StV to the right). This region (#474-585) is composed of 5 green/black macro-repeats.
4. Blue, greenish-blue, chrY.mPanPan1:34,034,978-34,373,351 (340kb) 1-3_11-29 (blue, #588-644) and 1-3_11-17_19-29 (greenish-blue, it is a blue StV with additional deletion of monomer 18).
5. Black, brown: chrY.mPanPan1:34,357,827-34,892,286 (0.5Mb), full-length HOR 1-29 (black) and 1-12_12-29 (brown, dup12).

Note that the CDR is located in region #2 on 12-22 concatemers and that 1-3_10-29 StVs which overlapped the CDR in chimpanzee are absent in bonobo.

Summary on StVs: The chimpanzee cenY looks more ancestral by structure, and in bonobo the right part has perhaps two islands (regions 3 and 5) which are apparently more reminiscent of the ancestral structure (have some full-length HORs) and the left part is presumably younger and is formed by structurally more derived expansions of variously deleted StVs.

<u>HORhaps.</u> For HORhap analysis we used all full-length HORs and StVs with monomer 12 or monomer 17 duplications from which one duplicated monomer was removed to bring their length to a full-length HOR. The chimpanzee centromere is well represented by this analysis but in bonobo 1-29 plus 1-12_12-29 StVs only compose ~20% of the whole array. However, the latter presumably would represent the older parts of the array which would be closer to chimpanzee, so our analysis would estimate the differences between the two species in a conservative manner.

Results of the cenY HORhap analysis in *Pan* are shown Ext. Data Fig. 4 and their interpretation and terminology are illustrated in section 2.4.1 above. The technical details are given in Fig. N7S. Panel B shows that the HORhaps for the full-length chimpanzee HORs (dark green) and the StV with monomer 17 duplication (bright green) are somewhat different (they do not mix in the tree). However, as all HORhaps were similar in amplification age and all were specific to chimpanzee, we treated them as one HORhap and designated green (Fig. N7SB). In bonobo, there were three well differentiated branches (HORhaps), two of which were represented only by full-length HORs (blue and grey HORhaps) and one had both full-length and duplicated StVs (cyan HORhap). The results of HORhap annotation show that various StVs in the bulk of the array mostly correspond to the younger (short twigs) blue HORhap, and the older (long twigs) cyan and grey HORhaps are located peripherally. The average intra-array divergence values shown in Fig. N7SA confirm that blue HORhap is younger in amplification age (more homogeneous) than black and grey ones (more divergent). Finally, the consensus tree shown in Fig. N7SC allows to ascertain the phylogenetic age and shows that the grey HORhap is almost equidistant to blue and green active HORhaps and hence is close to the HORs of the common ancestor of *Pan* species, while cyan is likely to represent the intermediate generation of bonobo cenY HORs. Fig. N7SD aligns the regions in the StV map described above to the HORhap arrays (see the legend).

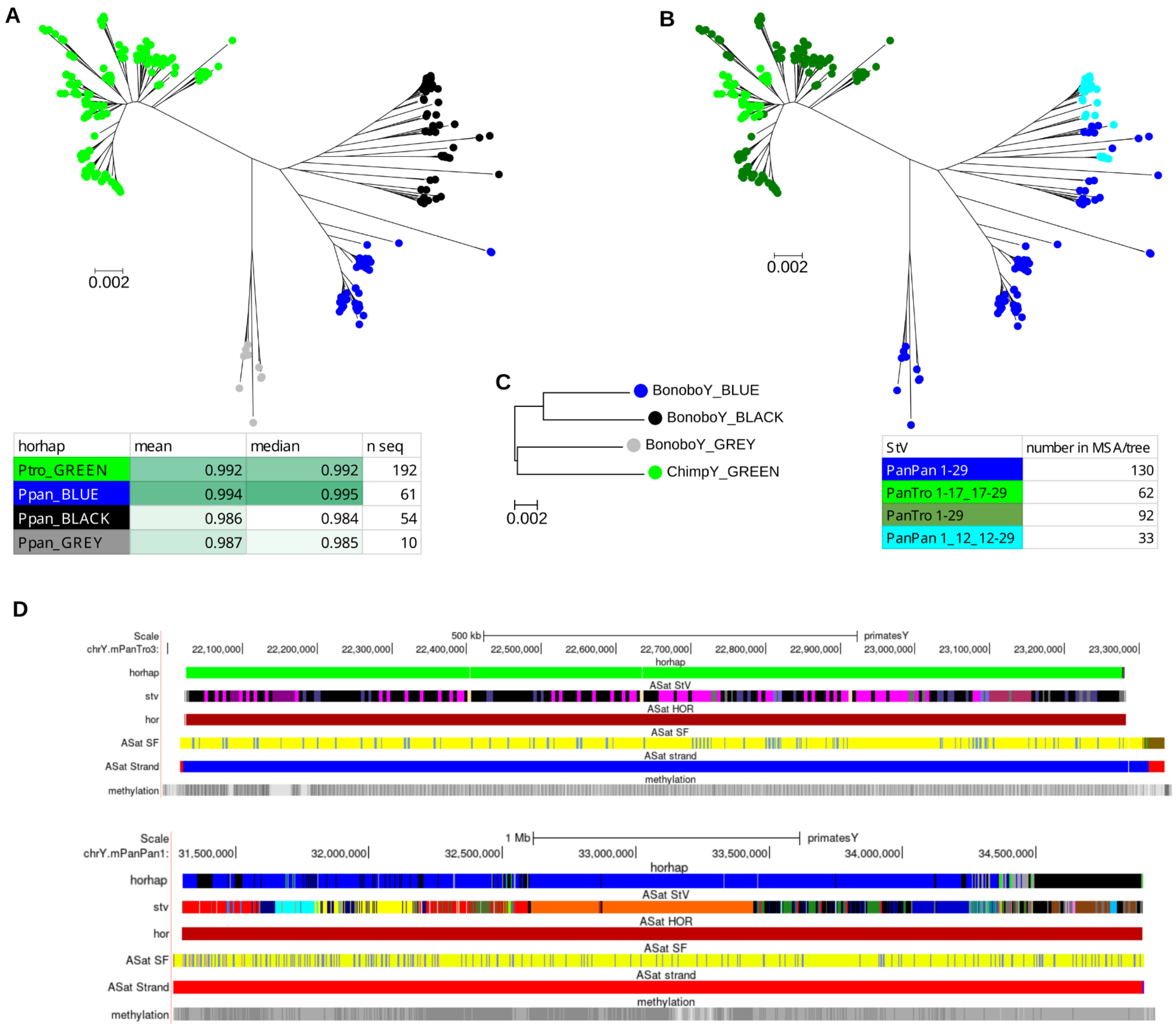

**Figure N7S. (A)** Tree of all full length HORs from both species (S4CYH1L.29-1), chimpanzee S4CYH1L.1-17_17-29 and bonobo S4CYH1L.29-12_12-1. Duplicated monomers were excised to allow alignment with full-length HORs. Robust branches were used to define Horhaps (marked by colors), create HMMs and consensus HORhaps. In the color legend, the intra-array divergence and a number of HORs used are shown for each HORhap (this tree is shown in Ext. Data Fig. 4). Note that the blue and green active HORhaps are more homogeneous than the older black and grey ones. **(B)** Same tree marked by StV shows distribution of full-length HORs and HORs with duplicated monomers into the HORhaps. **(C)** HORhap consensus tree shows the phylogenetic age of HORhaps. **(D)** Genome Browser tracks show correspondence of HORhaps to regions with specific StV structure listed in StV section.. One can appreciate that the older grey and black HORhaps form region 5, and regions 1-5 are all covered by active blue HORhap despite their structural diversity.

Note that the bonobo BLACK HORhap array, in its central part, contains StVs with a truncated (~128 bp long) monomer 1. This polymorphism can be visualized by a short match with the kmer TGCAGATTCCCCAAAGGAAGGTATCAAAAC (Fig. N7T). Chimpanzee has no such short monomers.

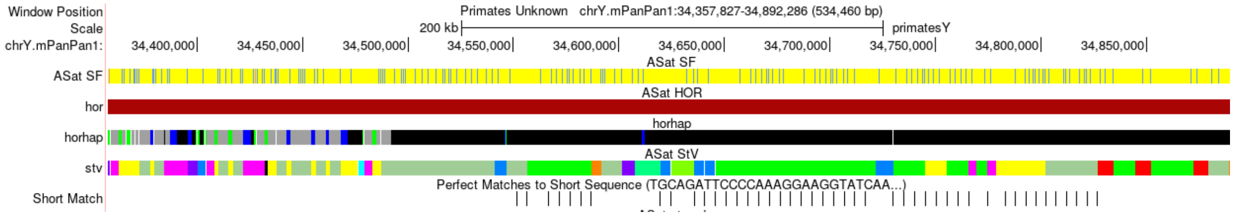

**Fig. N7T. HORs with truncated monomer 1 in the BLACK HORhap array.**

Summary on HORhaps. Clearly the HORhap structure is different between species. Chimpanzee does not have any full-length HORs older than the green HORhap and does not have any significant overlap with bonobo. The latter has retained at least 2 previous generations of HORhaps, of which the grey is approximately equi-distant to active HORhaps of both species and is likely to be close to active centromeres of the common ancestor of chimpanzee and bonobo.

##### 2.4.4. The patterns of AS-containing SDs in the arms of chrY in chimpanzee and bonobo.

The patterns of AS-containing SDs are very different between the twin *Pan* species. Bonobo has a simple pattern where only the SDs which have Ga (SF4, yellow) and Ha (SF6, brown) monomers are present on the p-side, while only the SDs which have the Ca (SF9, red) and Ba (SF10, orange) monomers are present on the q-side (Fig. S15A). Judging by the AS patterns all yellow/brown SDs are likely to be the offspring of the same single SD which was first multiplied in several steps that included inversions to form a group of 6 SDs and this block was duplicated in direct orientation to yield a pattern of 12 yellow/brown SDs on the short arm. The same is true of the red/orange SDs which contain apparently the same AS piece in both direct and reverse orientations (Fig. N7U). The AS part of this SD is the same which we described previously in human chromosome Y[105].

#### Human copy (direct)

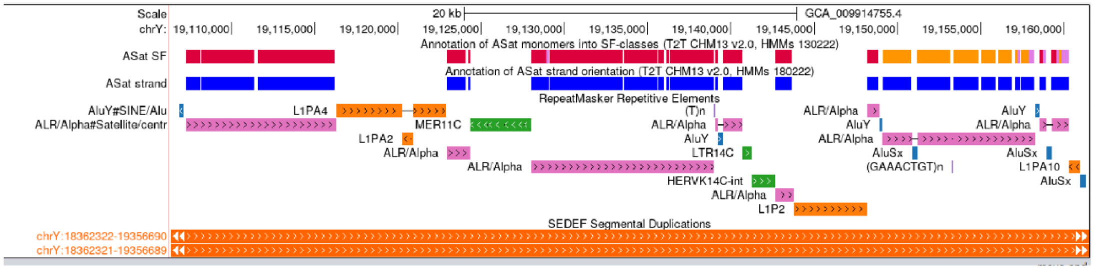

#### Human copy (reverse)

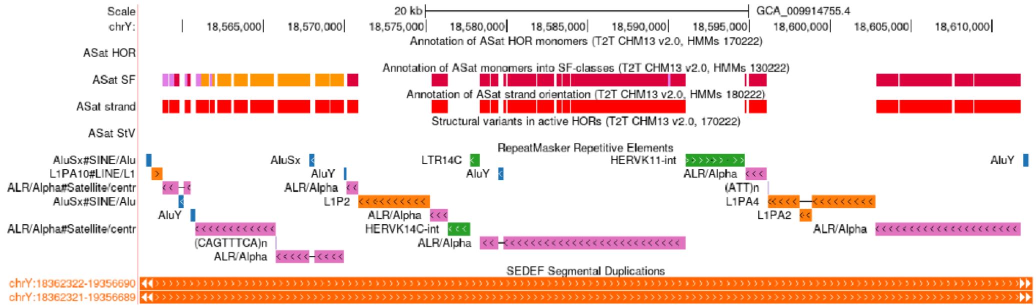

#### Bonobo copy (reverse)

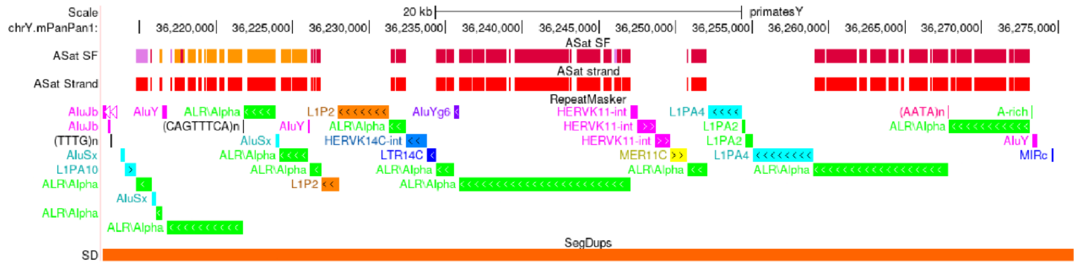

#### Chimpanzee copy (reverse)

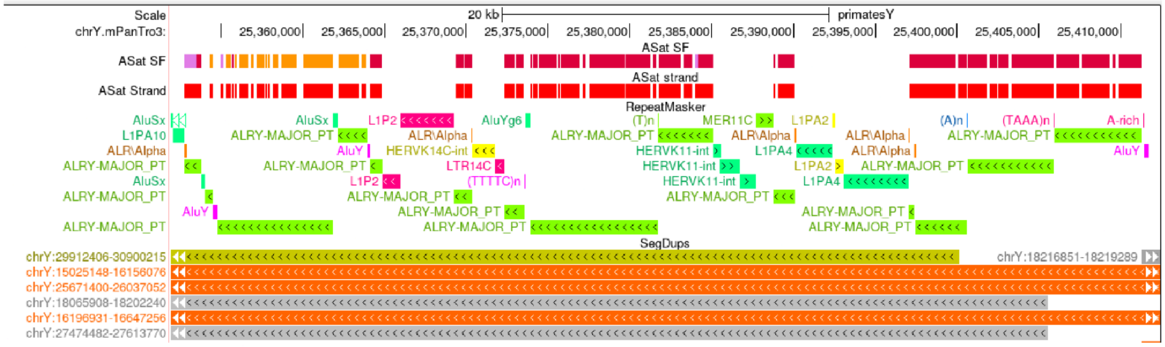

**Fig. N7U. Comparison of the copies of the red/orange cenY AS-containing SD in humans and *Pan* species.** The copies of the red/orange SD AS array are pretty much the same in all 3 species. Sometimes the parts of HERVK11-int at chrY.mPanPan1:38842982-38843970 are classed as MER11C and sometimes the whole repeat, but it seems it is always the same element, perhaps variously deleted.

##### 2.5. S. orangutan/B. Orangutan

Analysis of centromeres in orangutans is presented in Ext. Data Fig. 4 and Fig. S15A-D. Here we will just provide a few additional comments and details to fortify the conclusions stated in the main text.

**2.5.1. cenX.** Despite the apparent collinearity of the AS arrays and similar CDR position in the two species (Fig. N7V), the composition of the active cenX array (orange) is different, as also shown in Ext. Data Fig. 4 and Fig. S15B-D. This may be additionally illustrated by distribution of perfect matches to a kmer caactctgtgagttcaacacacacatcacaaa (32mer) partially specific to B. orangutan and centered on a single-nucleotide difference between the HOR consensus sequences of the two species (Fig. N7V). It indicates the presence of a small number of HORs characteristic of B. orangutan at the right flank of S. orangutan array. This suggests that B. orangutan structure is likely similar to that of the common ancestor of both species, and the S. orangutan active array have experienced the expansion of a new HOR variant (not marked by the kmer) and lost most of the ancestral array after the separation of the two species. Some remnants of the ancestral array have been preserved at the right flank. That aligns with the HORhap patterns shown in Ext. Data Fig. 4C.

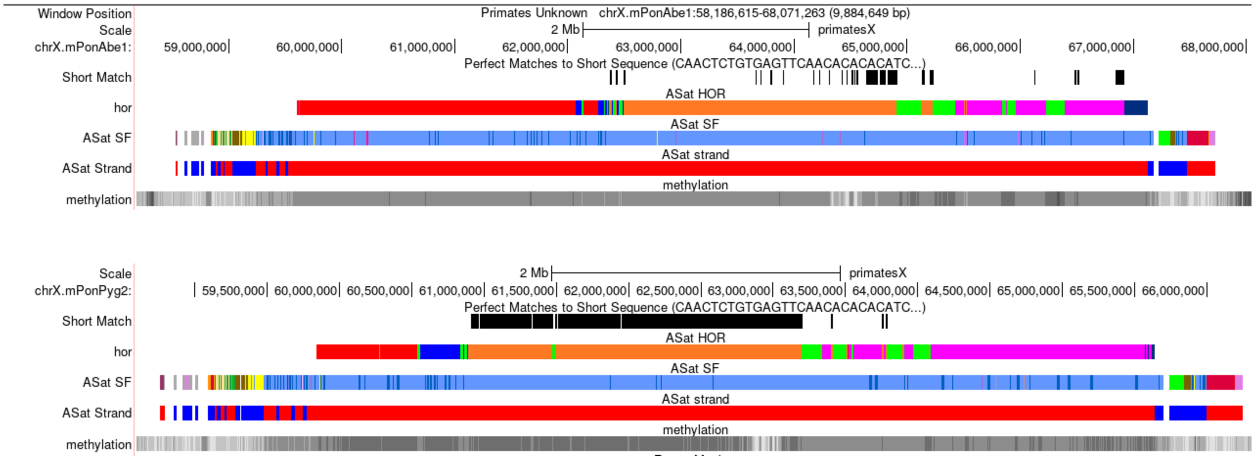

**Fig. N7V. B. orangutan-specific kmer highlights the differences in cenX active arrays in the two species.**

**2.5.2. cenY.** The values of the intra-array identity for orangutan cenY HORhaps are shown in Table N6C to additionally document the point of Ext. Data Fig. 4. The more derived and more recently amplified red and grey active HORhaps are more homogeneous than other HORhaps with identity values matching apparent "amplification age" noted from the HOR trees in Ext. Data Fig. 4.

**Table N7C. Intra-array identity in orangutan cenY HORhaps.**

| horhap | mean similarity |
|---|---|
| Ppyg_GREY | 0.9995 |
| Ppyg_RED | 0.9990 |
| Pabe_GREEN | 0.9987 |
| Pabe_BLUE | 0.9983 |
| Ppyg_LILAC | 0.9982 |
| Pabe_BLACK | 0.9939 |

Note: Grey HORhap has only 2 full-length copies which are in duplication, so there is only one independent copy, and the divergence value should be disregarded.

From our analysis of 100 randomly picked HORs in Ext. Data Fig. 4, we have concluded that each species has its own HORhaps which are not present in the other one. However, close-ups of HORhap annotation tracks in Fig. N7W allow to view the entire lengths of the arrays and show some very limited apparent presence of B. orangutan HORhaps (blue and cyan) in S. orangutan and of S. orangutan HORhap (lilac) in B. orangutan. We have examined this cross-contamination more closely by manually reviewing respective regions in our HORhap and StV annotations and found no cross-presence of full-length HORs. Apparent cross contamination is due to complex StV structure in the regions which fragments the HORs into small pieces and often precludes reliable HORhap identification. Note that grey color in StV tracks identifies the regions formed by full-length HORs, and the areas of the HORhap tracks overlapping grey color in StV track are highly reliable.

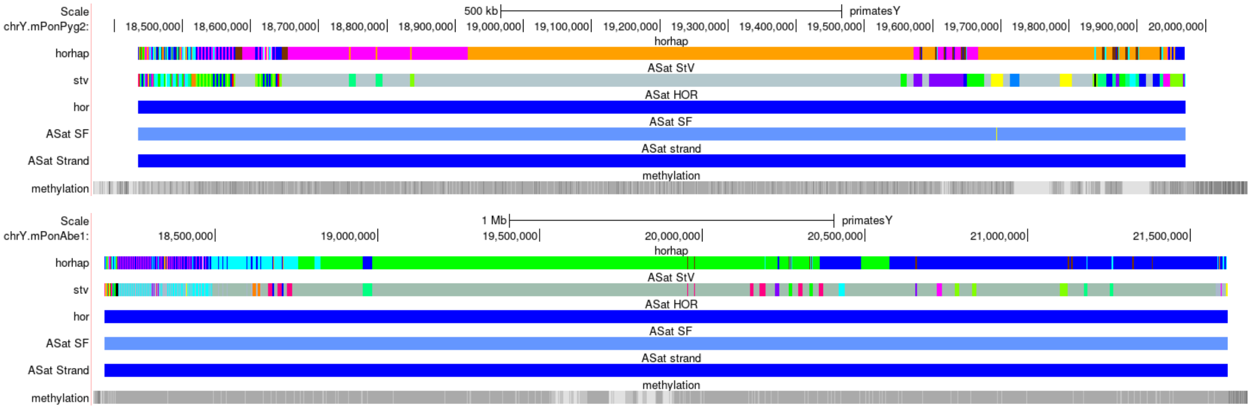

**Fig. N7W. Close-ups of orangutan cenY HOR arrays.**

#### Note S8. Additional FISH validations of rDNA and PARs

Using samples from individuals and species without assembled genomes, we cytogenetically characterized rDNA regions, in order to assess their localization in great and lesser apes, performing fluorescent in situ hybridization (FISH) experiments using rDNA-carrying BAC clone RP11-450E20 [115] as a probe. To confirm the sex of the cell lines tested, other probes were used. In particular, the great apes were tested using the pseudoautosomal region(PAR)-specific clone (RP11-990G10), while the gibbon lines were characterized using an X-specific clone (CH271-132L14) obtained from the *Nomascus leucogenys* (NLE) BAC library.

The experiments were performed on metaphase spreads of different hominid species: *Homo sapiens* (HSA; human), *Pan troglodytes* (PTR; chimpanzee), *Gorilla gorilla* (GGO; gorilla), *Pongo abelii* (PAB; Sumatran orangutan), and *Pongo pygmaeus* (PPY; Bornean orangutan), as well as three lesser apes—*Nomascus leucogenys* (NLE; white-cheeked gibbon), *Nomascus concolor* (NCO; black crested gibbon), and *Hylobates lar* (HLA; lar gibbon).

**Methods.** To prepare metaphase spreads, a skin cell line of GGO and lymphoblastoid cell lines of PTR, PPY, PAB, and three different gibbons (NCO, NLE, HLA) were used. In particular, we used two Chimpanzee (PTR 1 and PTR 8), one Gorilla gorilla (GGO 9), one Sumatran (PAB 16) and one Bornean (PPY 19) orangutan cell lines.  Human lymphocytes derived from a normal donor peripheral blood sample were stimulated with phytohemagglutinin (PHA) and used for human spreads.

DNA extraction from selected BACs was done with Biorad Quantum Prep Plasmid Miniprep Kit. FISH experiments were performed essentially as previously described[116]): two hundred nanograms of the DNA probe labeled by nick-translation with Cy3-dUTP, Cy5-dUTP, or Fluorescein-dUTP were precipitated by ion exchange alcohol precipitation with human Cot DNA and finally denatured for 2 minutes at 70°C and hybridized at 37°C overnight. Post-hybridization wash was performed at 60°C in 0.1×SSC (three times, high stringency). Gibbon cell lines were washed with 2×SSC at 42°C, having used probes of different derivations (human and gibbon) in a co-hybridization. At the end, the slide was stained with DAPI, producing a Q-banding pattern. The fluorescence signals coming from Cy3, Cy5, FITC and DAPI were detected separately with specific filters using a Leica DMRXA epifluorescence microscope equipped with a cooled CCD camera (Princeton Instruments) and recorded as grayscale images.

**rDNA region.** As a positive control, FISH experiments on human metaphases revealed the presence of rDNA on all the acrocentric chromosomes (13, 14, 15, 21, 22) at the pter or pcen (Fig. N8A), in agreement with those shown by [115].

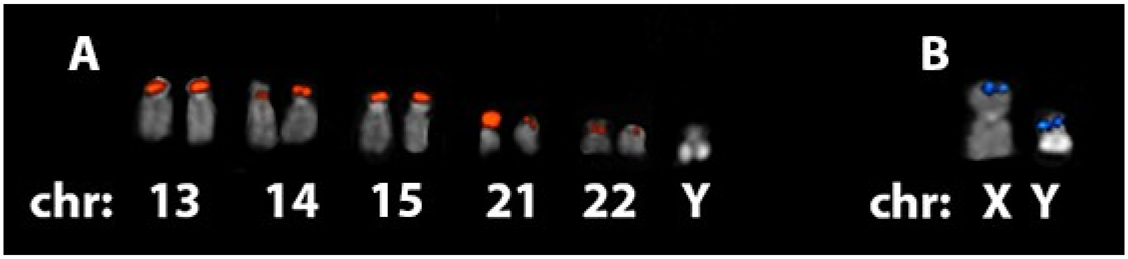

**Figure N8A. FISH experiment on human metaphase.** (A) The rDNA-carrying BAC RP11-450E20 (red signals) showed the presence of rDNA on the acrocentric 13, 14, 15, 21, 22 human chromosomes. (B) The RP11-990G10 probe (blue signals) confirms the location on the p arm of both sex chromosomes.

In both PTR cell lines, rDNA probe signals were located on a few acrocentric chromosomes (chr 14, 15, 17, 22 and 23 - homologous to human XIII, XIV, XVIII, XXI and XXII, respectively). No signals were detected on acrocentric chromosomes (12, 13, 16; Fig. N8B).  We detected a difference in signal intensity between PTR1 and PTR8 (Table N7A).

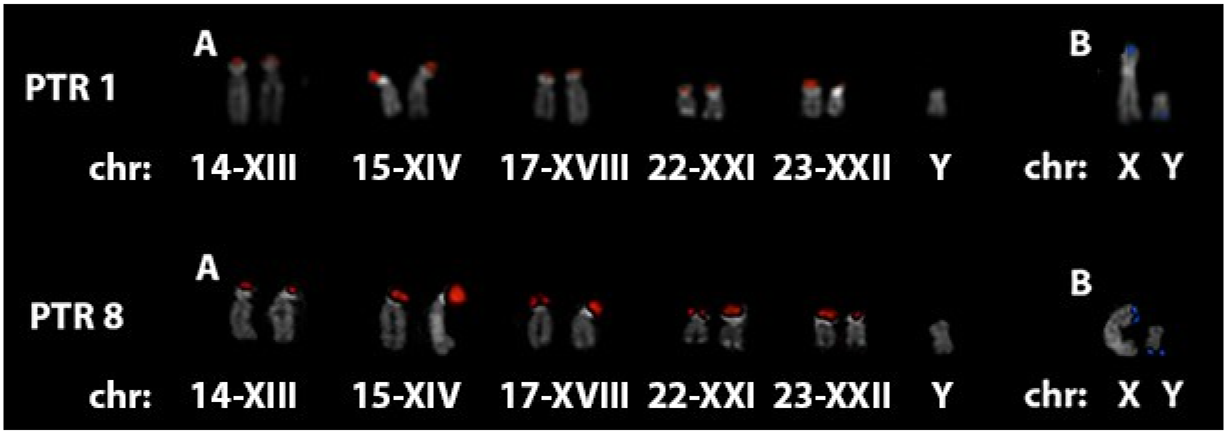

**Figure N8B. FISH experiment on PTR1 and PTR8 (*Pan troglodytes*) metaphases.** (A) RP11-450E20 probe (red signals) showed the presence of rDNA on the acrocentric 14, 15, 17, 22, and 23 while (B) The RP11-990G10 probe (blue signals) had unusual localization on the q arm of the Y chromosome. The Arabic numbers indicate the chromosomes with the nomenclature of the PTR karyotype, while the Roman numbers refer to the nomenclature of the corresponding human homologous chromosomes.

The male gorilla showed the presence of rDNA on the two small acrocentrics 22 and 23 (homologous to the human XXI and XXII) and on the pter of chromosome 1 (in heterozygous state) (Fig. N8C). None of the large acrocentrics (11, 12, 14, 15, 16) showed the presence of rDNA.

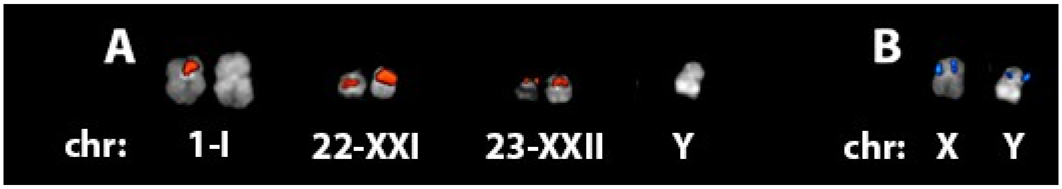

**Figure N8C. FISH experiment on GGO9 (*Gorilla gorilla*) metaphase.** (A) The rDNA-carrying BAC (red signals) showed the presence of rDNA on the acrocentric chromosomes homologous to XXI and XXII human chromosomes and in heterozygosis on chr I, while (B) The RP11-990G10 probe (blue signals) confirms the location at the p arm of both sex chromosomes, but on the X chromosome it is duplicated. The Arabic numbers indicate the chromosomes with the nomenclature of the GGO karyotype, while the Roman numbers refer to the nomenclature of the corresponding human homologous chromosomes.

In PAB (Fig. N8D) and PPY (Fig. N8E), rDNA was on chromosomes IIq (PPY in heterozygosis), IIp, IX, XIII, XV (PAB in heterozygosis), XXI, XXII. However, PAB showed the presence of rDNA in heterozygosis also on chromosome XIV, while in PPY no signal of the rDNA probe was detected on this chromosome. In both cell lines, rDNA mapped at the qter of chr Y.

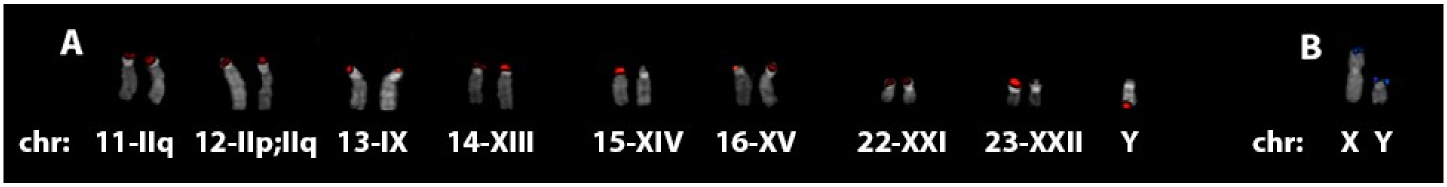

**Figure N8D. FISH experiment on PAB16 (*Pongo abelii*) metaphase.** (A) The RP11-450E20 probe (red signals) showed the presence of rDNA on the acrocentric IIq, IIp, IX, XIII, XIV, XV, XXI, XXII and Y chromosomes, while (B) The RP11-990G10 probe (blue signals) confirmed the location on the p arm of both sex chromosomes. The Arabic numbers indicate the chromosomes with the nomenclature of the PAB karyotype, while the Roman numbers refer to the nomenclature of the corresponding human homologous chromosomes.

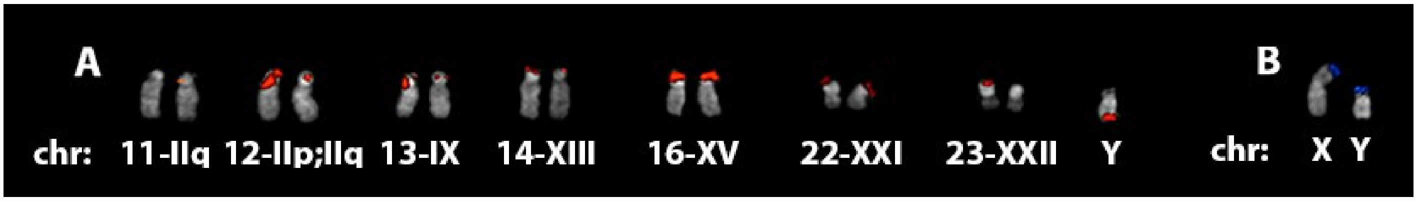

**Figure N8E. FISH experiment on PPY19 (*Pongo pygmaeus*) metaphase.** (A) The rDNA-carrying probe RP11-450E20 (red signals) showed the presence of rDNA, (B) The RP11-990G10 probe (blue signals) confirmed the location on the p arm of both sex chromosomes. The Arabic numbers indicate the chromosomes with the nomenclature of the PPY karyotype, while the Roman numbers refer to the nomenclature of the corresponding human homologous chromosomes.

In both *Nomascus* gibbons (*N. concolor* - Fig. N8F and *N. leucogenys* - Fig. N8G), we highlighted the same pattern of rDNA probe signals, localized at the pter of chromosomes 24, 25 and Y. However, *Hylobates lar* (Fig. N8H) carries rDNA only on chromosome 12 at a pericentromeric location, highlighting potentially genus-specific rDNA content and locations.

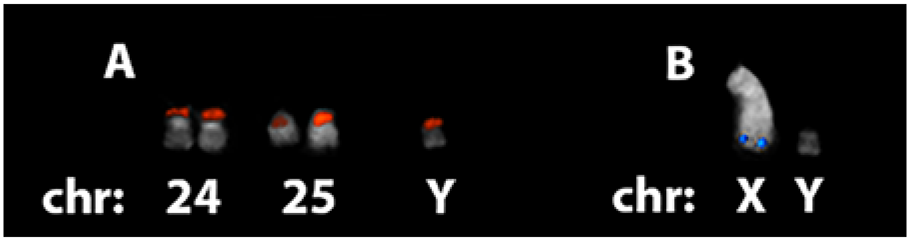

**Figure N8F. FISH experiment on NCO (*Nomascus concolor*) metaphase.** (A) The BAC RP11-450E20 (red signals) showed the presence of rDNA on chr 24, 25 and Y. (B) The CH271-132L14 probe (blue signals) hybridized only at the X qter.

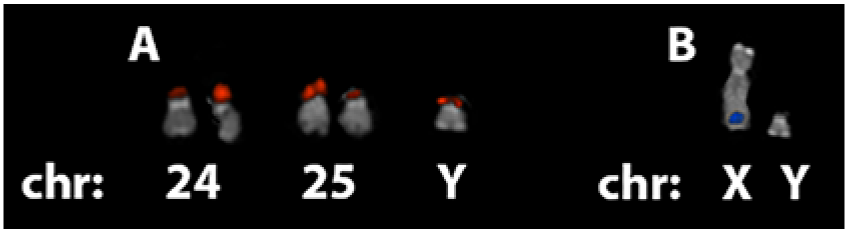

**Figure N8G. FISH experiment on NLE (*Nomascus leucogenys*) metaphase.** (A) The rDNA-carrying BAC (red signals) showed the presence of rDNA at chr 24, 25 and Y. (B) The CH271-132L14 probe (blue signals) hybridized only at the X qter

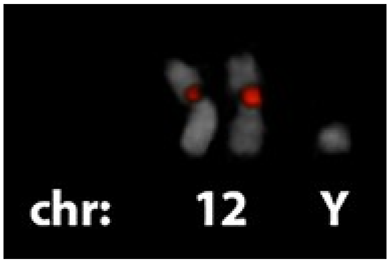

**Figure N8H. FISH experiment on HLA (*Hylobates lar*) metaphase.** The BAC rDNA RP11-450E20 (red signals) showed the presence of rDNA.

**PAR region.** All great apes X chromosomes had the PAR region at the pter and in particular a duplication of the PAR region on the gorilla X chromosome has been reported (Fig. N8C).

On the Y chromosome the PAR region is always found at the pter, with the exception of *Pan troglodytes*, which is the only species on which the PAR region probe was localized at the qter, rather than the classic pter.

The PAR region was not tested for gibbons due to higher divergence between gibbons and human. The probe (CH271-132L14) used on gibbon cell lines showed the expected localization at the qter of chrX only on *Nomascus* cell lines. Instead no signals of this probe were detected on *Hylobates*.

**Sample information**:

- PTR1 is a male *Pan troglodytes* (called Tank) lymphoblastoid cell line
- PTR8 is a male *Pan troglodytes* (called Carl) lymphoblastoid cell line
- GGO9 is a male *Gorilla gorilla* skin-cell line

- PAB16 is a male *Pongo abelii* (called Sinjo - 1833) lymphoblastoid cell line
- PPY19 is a male (named Sumbo - L1847) orangutan hybrid (between a Sumatran and a Bornean) LB cell line
- NCO is a male *Nomascus concolor* lymphoblastoid cell line
- NLE is a male *Nomascus leucogenys siki* lymphoblastoid cell line with karyotype 53, XY+14
- HLA is a *Hylobates lar* lymphoblastoid cell line of a gibbon called Eddie

**Conclusions.** With these experiments we validated the localization of rDNA and PAR in great and lesser apes, showing how, from an evolutionary point of view, these regions have progressively modified both their localization and their copy number not only on the acrocentric chromosomes but also on the Y chromosomes of the various species analyzed (Fig. N8I).

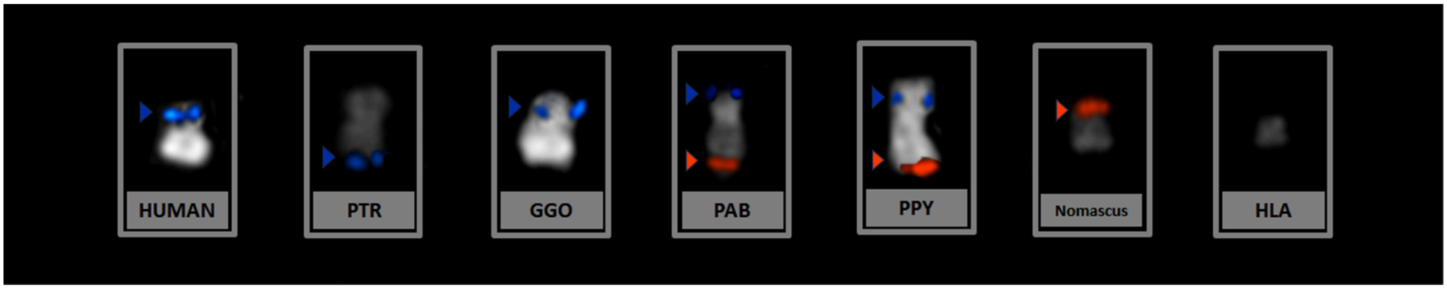

**Figure N8I. Summary of Y chromosome hybridizations of all the species analyzed (human, PTR, GGO, PAB, PPY, *Nomascus* and HLA) with the signals of rDNA-carrying BAC (red) and PAR region probe RP11-990G10 (blue).**

**Table N8A. Overview of all identifying signals with detailed localization in the cell lines tested using the rDNA-carrying BAC clone RP11-450E20 and a sex-chromosomes-specific clone (RP11-990G10 for HSA, PTR, GGO and PPY lines; CH271-132L14 for NCO and NLE).**

| Cell line | chr | RP11-450E20 localization | RP11-990G10 localization |
|---|---|---|---|
| HSA | 13 | +pter/++pter | -/- |
| | 14 | +pcen/++pter | -/- |
| | 15 | ++pter/++pter | -/- |
| | 21 | ++pter/+pter | -/- |
| | 22 | +pcen/+pcen | -/- |
| | X | - | +pter |
| | Y | - | +pcen |
| PTR1 | 14-XIII | +pter/+pter | -/- |
| | 15-XIV | ++pter,+pcen/+pter,+pcen | -/- |
| | 17-XVIII | +pter/+pter | -/- |
| | 21-XXI | +pter/+pter | -/- |
| | 23-XXII | ++pter/+pter | -/- |
| | X | - | +pter |
| | Y | - | +qter |
| PTR8 | 14-XIII | +pter/+pter | -/- |
| | 15-XIV | +pter/++pter | -/- |
| | 17-XVIII | +pter/+pter | -/- |
| | 22-XXI | +pter/++pter | -/- |
| | 23-XXII | ++pter/+pter | -/- |
| | X | - | +pter |
| | Y | - | +qter |

| | | | |
|---|---|---|---|
| **GGO9** | 1-I | +pter/- | -/- |
| | 22-XXI | pcen+,qcen+/pcen+,qcen+ | -/- |
| | 23-XXII | +pter/+pter | -/- |
| | X | - | +pcen,+pter |
| | Y | - | +pter |
| **PAB16** | 11-IIq | +pter/+pter | -/- |
| | 12-IIp;IIq(2A) | +pter/+pter | -/- |
| | 13-IX | +pter/+pter | -/- |
| | 14-XIII | +pter/++pter | -/- |
| | 15-XIV | ++pter/- | -/- |
| | 16-XV | +pter/- | -/- |
| | 22-XXII | +pter/+pter | -/- |
| | 23-XXII | ++pter/+pter | -/- |
| | X | -/- | pter+ |
| | Y | ++qter | pter+ |
| **PPY19 cell line** | 11-IIq | -/+pter | -/- |
| | 12-IIp;IIq(2A) | ++pter/+pter | -/- |
| | 13-IX | ++pter/+pter | -/- |
| | 14-XIII | +pter/+pter | -/- |
| | 16-XV | ++pter/++pter | -/- |
| | 22-XXII | +pter/+pter | -/- |
| | 23-XXII | +pter/- | -/- |
| | X | -/- | pter+ |
| | Y | +qter | pter+ |
| **Cell line** | **chr** | **RP11-450E20 localization** | **CH271-132L14 localization** |
| **NCO cell line** | 24 | +pter/+pter | -/- |
| | 25 | +pcen/+pcen | -/- |
| | X | - | +qter |
| | Y | +pter | - |
| **NLE cell line** | 24 | +pter/++pter | -/- |
| | 25 | ++pcen/+pcen | -/- |
| | X | - | +qter |
| | Y | +pter | - |
| **HLA cell line** | 12 | +pcen/+pcen,+qcen | -/- |
| | Y | - | - |

### Note S9. Ancestral (X-degenerate) genes on the Y chromosome: Comparisons with other studies and gene conversion analysis.

**Resolution of ancestral genes.** Previous evaluations of ancestral gene content on the Y chromosome of African apes suggest lower conservation of ancestral genes[117–119] than identified in this work (Fig. 6). This appears to be due to two factors (1) an increase in Y chromosome sequence quality and (2) increased resolution of gene annotations. Although mostly concordant, some genes were identified as present in these T2T assemblies that had been previously identified as missing in certain taxa (e.g. *TMSB4Y* in chimpanzee[117,118]). However, most discrepancies were due to changes in annotations. For example, previous annotation versions have suggested that some genes are functional, possibly based on their expression profiles, but are now annotated as pseudogenes in human (*TLXNGY*, previously known as *CYorf15a* and *CYorf15b,* and *PRKY*). Further, by aligning multispecies gametologs, we call additional pseudogenes based on truncations that differ from previous studies[117–119]. In sum, the increased quality of the T2T sex chromosome sequences, combined with curated annotation sets for these genes, has further elucidated their evolutionary history within the ape species studied here. Note that, at least in some cases, we have sequenced different individuals than those that were analyzed before, and some intraindividual variability is expected and may also contribute to the observed differences.

**Gene conversion analysis.** Curiously, we found little evidence of X-Y gene conversion in coding regions of Y-linked ancestral genes and their X-chromosomal homologs. We used GeneConv[120] to identify putative gene conversion within the T2T XY assemblies; we identified only two potential regions of gene conversion. The first is gene conversion between *NLGN4X* and *NLGN4Y* (Table N9A), a gametologous gene pair that previously had gene conversion reported for its intron[22]. The second is an interesting case of potentially ancestral gene conversion between *KDM6A* on the X chromosome and *UTY* on the Y chromosome, where these gametologs have higher sequence identity (measured with *p*-distance) to each other in African apes (86%) than in the other three studied taxa (83%; Table N9B). Our results differ from the ones previously reported for gene conversion in these genes[22] because we only analyzed their protein-coding regions. Nevertheless, the lack of evidence for gene conversion increases our confidence in the alignments and topologies of phylogenetic trees used for the selection analysis.

**Table N9A. GeneConv output for all ancestral genes with measured likelihood of X-Y gene conversion within protein-coding regions.** Column headers (as per GeneConv[120]) are defined as: "Type" Global or Pairwise inner fragment, global being more conservative than pairwise; "Species" Common name of a focal taxon from this study; "Genes" which X/Y gene pair was analyzed; "Sim P-value" p-value calculate using 10,000 permutations; "BC KA P-value" Bonferroni-corrected Karlin-Altschul P-values of each fragment; "Alignment Begin" beginning of fragment; "End" end of fragment; "Length" total length of the analyzed fragment; "Num Poly" number of polymorphic sites within the analyzed fragment; "Total Difs" total number of differences between the two sequences.

| Type | Species | Genes | Sim P-value | BC KA P-value | Alignment Begin | End | Length | Num Poly | Total Difs |
|---|---|---|---|---|---|---|---|---|---|
| Global | Human | *NLGN4X/Y* | 0.0094 | 0.02861 | 1081 | 1400 | 320 | 21 | 71 |
| Global | S. orang | *NLGN4X/Y* | 0.0492 | 0.13437 | 1357 | 1694 | 338 | 27 | 48 |
| Pairwise | Human | *NLGN4X/Y* | 0.0022 | 0.00409 | 1081 | 1400 | 320 | 21 | 71 |
| Pairwise | S. orang | *NLGN4X/Y* | 0.0099 | 0.0192 | 1357 | 1694 | 338 | 27 | 48 |

**Table N9B.** Pairwise percentage of sequence identity between X-Y gametologs (*KDM6A*/*UTY*) across studied primates. Notably, X-Y sequence identity is higher within the African apes (green) than in other apes (yellow).

| | Bonobo *KDM6A* | Bonobo *UTY* | Chimp *KDM6A* | Chimp *UTY* | Human *KDM6A* | Human *UTY* | Gorilla *KDM6A* | Gorilla *UTY* | Borang *KDM6A* | Borang *UTY* | Sorang *KDM6A* | Sorang *UTY* | Siamang *KDM6A* | Siamang *UTY* |
|---|---|---|---|---|---|---|---|---|---|---|---|---|---|---|
| Bonobo *KDM6A* | | | | | | | | | | | | | | |
| Bonobo *UTY* | 86 | | | | | | | | | | | | | |
| Chimp *KDM6A* | 100 | 86 | | | | | | | | | | | | |
| Chimp *UTY* | 86 | 100 | 86 | | | | | | | | | | | |
| Human *KDM6A* | 100 | 86 | 100 | 86 | | | | | | | | | | |
| Human *UTY* | 86 | 99 | 85 | 99 | 85 | | | | | | | | | |
| Gorilla *KDM6A* | 100 | 86 | 100 | 86 | 100 | 85 | | | | | | | | |
| Gorilla *UTY* | 86 | 99 | 86 | 99 | 86 | 99 | 86 | | | | | | | |
| Borang *KDM6A* | 100 | 86 | 100 | 86 | 100 | 86 | 100 | 86 | | | | | | |
| Borang *UTY* | 83 | 95 | 83 | 95 | 83 | 95 | 83 | 95 | 83 | | | | | |
| Sorang *KDM6A* | 100 | 86 | 100 | 86 | 100 | 86 | 100 | 86 | 100 | 83 | | | | |
| Sorang *UTY* | 83 | 95 | 83 | 95 | 83 | 95 | 83 | 95 | 83 | 100 | 83 | | | |
| Siamang *KDM6A* | 100 | 85 | 100 | 86 | 100 | 85 | 100 | 86 | 100 | 83 | 100 | 83 | | |
| Siamang *UTY* | 83 | 95 | 83 | 95 | 83 | 95 | 83 | 95 | 83 | 98 | 83 | 98 | 83 | |

### Note S10. Significant shifts in ampliconic gene copy number

To estimate evolution of the copy number of ampliconic genes on the Y chromosome, we used CAFE (v5.1.1)[121]. CAFE fits a birth-death process model across the phylogeny for estimating the expansion or contraction of gene family copy-size across a phylogeny, isolating branches where large-scale changes have occurred that violate a single-rate birth-death model. We used a phylogeny inferred from all single-copy orthologous sites on chrY across species. To rescale branch-lengths into years we used the estimate of the human chrY mutation rate of $8.88 \times 10^{-10}$ mutations per year as a proxy for all great apes and rescaled branch lengths to millions of years[79]. The resulting rescaled tree, with branch-lengths in millions of years, used for estimation of gene-copy evolution was:

```
(Symphalangus_syndactylus.mSymSyn1_Y:48.6983411036036,(Pongo_abelii.nPonAbe1_Y:
0.698506081081081,Pongo_pygmaeus.mPonPyg2_Y:0.7118148648648649):32.946111486486
49,(Gorilla_gorilla.mGorGor1_Y:14.684040427927926,(chm13.chm13_Y:10.69708,(Pan_
troglodytes.mPanTro3_Y:2.4742399774774775,Pan_paniscus.mPanPan1_Y:2.43342804054
05404):7.452579391891892):5.674023986486486):30.526842004504502);
```

For Y ampliconic genes, we used the estimated number of copies based on the assembly and manual curation, restricting counts to the inferred non-pseudogenized copies in each species (Fig. 6). For the human HG002 reference genome we used the number of non-pseudogenized copies[75]. Since CAFE does not support zero gene copies at the root, we excluded such gene families from downstream inferences. All subsequent analyses considered only the following seven gene families: *CDY*, *DAZ*, *HSFY*, *RBMY*, *TSPY*, *VCY*, and *FRG1*.

Under a single-rate parameter model across gene families, we estimate the overall birth-death rate (**λ**) to be 0.021 events per million years. This is approximately half of the inferred rate of 0.05 events per million years from[122] but still ~10x as high as reported for non-Y gene families across primates[123], although the latter reference is based on non-T2T assemblies.

We find evidence of deviation from a single birth-date rate accelerated evolution for three of the seven tested gene families (*CDY* $p$<0.001; *RBMY* $p$<0.001; *TSPY* $p$<0.001; Bonferroni $p$-value threshold 0.05 / 7 = 0.007; Fig. N10A, Table N10A). For three genes, we could detect individual lineages with significant gene family contractions or expansions. For *CDY*, the bursts are largely driven by a significant increase and a significant decrease in copy number in the Sumatran and Bornean orangutan lineages, respectively ($p$ =2.00e-07 and p=7.47e-04, respectively). The significant inferred events for the *RBMY* family are a significant increase in copy number in the bonobo lineage ($p$=4.46e-14) and significant decreases in the chimpanzee (p=1.70e-04) and Sumatran orangutan ($p$=1.53e-03) lineages. *TSPY* experienced several significant copy number increases and decreases, with a pronounced expansion in the human lineage ($p$=2.64e-06).

Overall these results are broadly consistent with previous findings for Y-chromosome ampliconic gene family evolution in primates[122]. A limitation of our analysis is that we are only able to use a single point estimate of the ampliconic gene copy from the reference genomes of each species. Therefore, these results should be confirmed with the analysis of intraspecific variation in copy number.

**Figure N10A.** Evidence of chrY ampliconic copy number change across primate genomes for *CDY*, *RBMY*, and *TSPY* gene families. Tips are annotated with observed numbers of gene-family copies (internal nodes reflect predicted copy-number from the single rate model in CAFE). Branch colors reflect phylogenetically significant increase (orange) or decrease (blue) of copy-number.

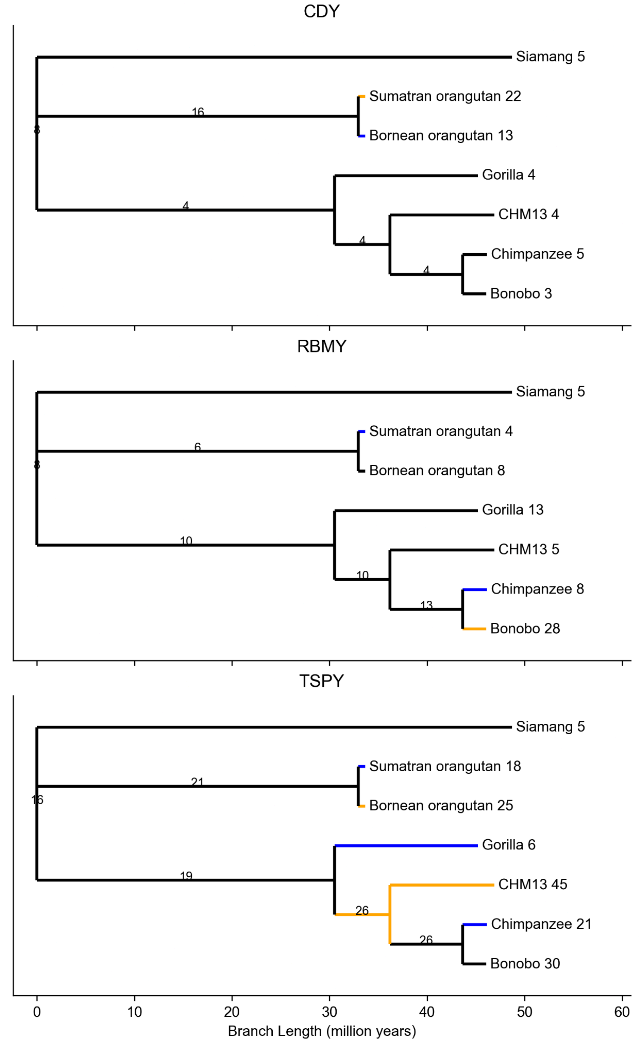

**Table N10A. Branch-specific *p*-values for deviation in copy-number of from a single-rate model from CAFE5 along the primate phylogeny.** Significant *p*-values (using Bonferroni-corrected threshold 0.05/13 = 0.0038, as there are 13 nodes in the tree) are shown in bold. *P*-values indicating significant expansions are shown in orange, and the ones indicating significant contractions in blue.

| **Branch** | ***CDY*** | ***RBMY*** | ***TSPY*** |
|---|---|---|---|
| Siamang | 0.643997 | 0.643997 | 0.0339888 |
| Sumatran orangutan | **2.00E-07** | **0.00152769** | **0.00155311** |
| Bornean orangutan | **0.000746571** | 0.00479986 | **0.000312361** |
| Gorilla | 0.0406925 | 0.179081 | **1.64E-05** |
| Human | 0.834446 | 0.0147669 | **2.64E-06** |
| Chimpanzee | 0.0955534 | **0.000169962** | **0.00298104** |
| Bonobo | 0.23104 | **4.46E-14** | 0.0254488 |
| Siamang --- (Bornean Orangutan, Sumatran Orangutan) | 0.024002 | 0.692508 | 0.209156 |
| Siamang---Gorilla, (Human, (Chimpanzee, Bonobo))) | 0.289665 | 0.367422 | 0.393576 |
| Human, (Chimpanzee, Bonobo)---Gorilla, (Human, (Chimpanzee, Bonobo))) | 0.758542 | 0.859516 | **0.00235146** |
| (Chimpanzee, Bonobo)---Human, (Chimpanzee, Bonobo) | 0.793386 | 0.117838 | 0.928161 |

### Note S11. Candidate *de novo* gene analysis

**Methods.** Genes on the Y chromosomes predicted to be novel according to the NCBI[124,125] and CAT[46] annotation pipelines were manually filtered for putative *de novo* gene candidates as follows. First, all pseudogenes and novel genes with a BLAT hit in the human genome CHM13 T2T (Jan 2022)[106] were discarded. The remaining novel gene candidates were then blasted against the NCBI non-redundant protein sequences (nr) database[126] using default parameters. All novel genes with homology to annotated proteins were discarded. Candidate *de novo* genes within PAR were discarded, as they are not specific to the Y chromosome. A total of two genes were the final *de novo* gene candidates on the non-recombining part of the Y.

Upstream and downstream syntenic genes of each of the two *de novo* gene candidates were retrieved from the corresponding gff3 files. Only conserved annotated protein-coding genes were used to define synteny; lncRNAs and pseudogenes were not used. For verification of *de novo* origin, we searched for sequences homologous to the *de novo* genes within all genomes available from this study, as well as outgroup species covering Old World monkeys, New World monkeys, and mouse (Table N11A). For this, we used BLASTn[104] locally with the transcript sequences of the corresponding *de novo* genes with the following command:

```
> blastn -query putative_denovo_all_spliced.fasta -db species.dna.toplevel.fa
-outfmt "6 qseqid qstart qend qseq sseqid sstart send sstrand sseq evalue
length pident" -out species_denovo_blast.csv
```

Homologous hits for each *de novo* gene candidate were manually verified, assigned to corresponding exons, and checked for synteny. In case of multiple hits for one exon with similar E-value (<10e-5) and query cover, only the hit in the syntenic region and close to other exons of the same *de novo* gene was kept as the best hit. Exons that were found on the Y chromosome, but not on the X chromosome, of a species were searched for in the X-Y chromosome alignment file. Aligned regions including 100 bp up- and downstream of the target exon were extracted from MAF files using the get_all_dn_XY.py script (available on GitHub https://github.com/makovalab-psu/T2T_primate_XY/tree/main/denovo_genes) and were manually examined for corresponding exon regions. All best hits were then compared to TE and repeat regions as annotated above using the script TE_denovo.py (available on GitHub in https://github.com/makovalab-psu/T2T_primate_XY/tree/main/denovo_genes). The age of non-coding origins was inferred using the pairwise divergence time estimated by timetree.org[127] between the species containing the coding *de novo* gene candidate and the furthest outgroup species with a BLASTn hit. The transcription factor binding site (TFBS) motifs were predicted as described in[128] taking 1 kb upstream of the first position of the best hit and 100 bp downstream.

Different properties of the proteins encoded by the *de novo* genes were predicted. NetSurfP-3.0[129] was used to predict secondary structure elements, disordered regions were predicted using flDPnn[130], determining the percentage of residues in disordered regions using the binary predictions (threshold for disorder = 0.3) and the mean disorder propensity over the whole sequence. Aggregation propensity as the normalized a4v sequence sum for 100 residues (Na4vSS) was predicted by AGGRESCAN[131]. Solubility was predicted using Protein-Sol[132] and categorized into 'low solubility' if below the experimental average, 'high solubility' if above the experimental average, or 'average solubility' if within 0.1 of the experimentally determined average. 3D protein structures were predicted using ESMFold [133].

**Table N11A. Outgroup genomes used for blast**

| Scientific name | Common name | Ensembl genome version |
|---|---|---|
| *Homo sapiens* | Human | Xchr: T2T-CHM13; Ychr: T2T-hg002 |
| *Macaca mulatta* | Macaque | Mmul_10 |
| *Saimiri boliviensis* | Bolivian squirrel monkey | SaiBol1.0 |

| | | |
|---|---|---|
| *Callithrix jacchus* | White-tufted-ear marmoset | mCalJac1.pat.X |
| *Carlito syrichta* | Tarsier | Tarsius_syrichta-2.0.1 |
| *Mus musculus* | Mouse | GRCm38 |

**Results.** *De novo* genes are defined as orphan genes without or with a limited number of homologs in closely related species. *De novo* genes thus emerged from ancestral non-coding regions of the genome. Multiple lines of evidence exist that *de novo* genes may, albeit very rarely, become fixed and assume essential functions[134,135]. Several *de novo* genes play a role in fertility in different species[136–138], and many have testis-specific expression[137,139,140]. This makes the Y chromosome a promising candidate for *de novo* gene emergence. However, excellent assemblies of closely related species, which are required to unambiguously identify non-genic homologous sequences in synteny in genomes of closely related species, have been scarce[141]. Here, we investigated our new high-quality T2T assemblies of the Y chromosome for six ape species for *de novo* gene emergence.

We were able to trace the origins for two *de novo* gene candidates on the non-PAR Y chromosome, one specific to siamang, and the other specific to bonobo. Both *de novo* gene candidates belong to the ampliconic sequence class genes and, therefore, are not well conserved between the X and Y chromosomes. Both candidate *de novo* genes have annotated genes in close proximity that aided in finding the syntenic regions across species (Table N11B). Homologous non-coding sequences within syntenic regions were detected for the two *de novo* genes in close outgroup species (Old and New World monkeys). None of the *de novo* genes have detectable homology in the X and Y chromosome or other chromosomes with the syntenic genes in mouse.

**Table N11B. *De novo* gene properties**

| Species | Name | GC | exons (coding) | upstream gene | downstream gene | non-coding origin | sequence class |
|---|---|---|---|---|---|---|---|
| Bonobo | LOC129395657 | 0.66 | 4 (4) | *RNA5-8SN5* | *TSPY* | 30 MYA | AMPLICONIC |
| Siamang | LOC129476750 | 0.53 | 6 (3) | *VCY* | *NLGN4Y* | 90 MYA | AMPLICONIC |

The *de novo* gene candidate *LOC129395657* in bonobo comprises four exons with exons 2 and 3 completely within an interspersed repeat (SSU-rRNA_Hsa, or a small subunit of the rRNA). The gene is located between conserved genes *RNA5-8SN5* and *TSPY*. Twelve copies of the *LOC129395657* gene exist on the bonobo Y chromosome, all of which have the potential for being protein-coding as they contain complete open reading frames (ORFs) and are in proximity (but varying orientation) to *TSPY* copies. However, only one copy of this *de novo* gene is a predicted gene according to the gene annotation pipelines in bonobo. Three homologous copies of *LOC129395657* were identified on the chimpanzee Y chromosome, two of which can result in an intact ORF when transcribed, and one with a premature stop codon. None of the homologous copies detected by BLASTn are predicted genes according to our gene annotation. One homologous sequence containing a premature stop codon is present in the human Y chromosome close to one of the *TSPY* gene copies, but not within the *TSPY* composite repeats[107]. Another homologous sequence spanning all exons of *LOC129395657* was identified on chr1 in gorilla, but lacked the start codon. Partial homologous sequences can be detected in Y chromosomes of orangutans and on chr23 of siamang. No sequence homology to X chromosome regions in bonobo or other primate species was detected by either BLASTn or by examining the X-Y chromosome alignments.

For further verification of the *de novo* gene candidate in bonobo, all regions with homology to the complete transcript sequence were screened for TFBS motifs. The total number of TFBS motifs in upstream regions is highest in the originally detected bonobo *de novo* gene candidate and five of the homologous copies on the bonobo Y chromosome, but also in the upstream region of the non-coding chimpanzee homolog with a premature stop codon. Other homologous potentially protein-coding sequences in bonobo and chimpanzee have a lower number of motifs comparable to the non-coding homologs in gorilla and human. This underlines the specificity of *LOC129395657* to bonobo, as the sequences with protein-coding abilities in chimpanzee have lower chances of being transcribed(due to the lack of TFBS motifs)  than the bonobo *de novo* gene.

The protein encoded by the Y-chromosome specific *de novo* candidate gene *LOC129395657* is predicted to be highly soluble and non-aggregating, which are beneficial properties for novel proteins and have been observed previously as a characteristic of *de novo* proteins[142]. In line with the solubility predictions, the protein is highly disordered according to the secondary structure prediction, the 3D structure prediction and disorder prediction (Table N11C).

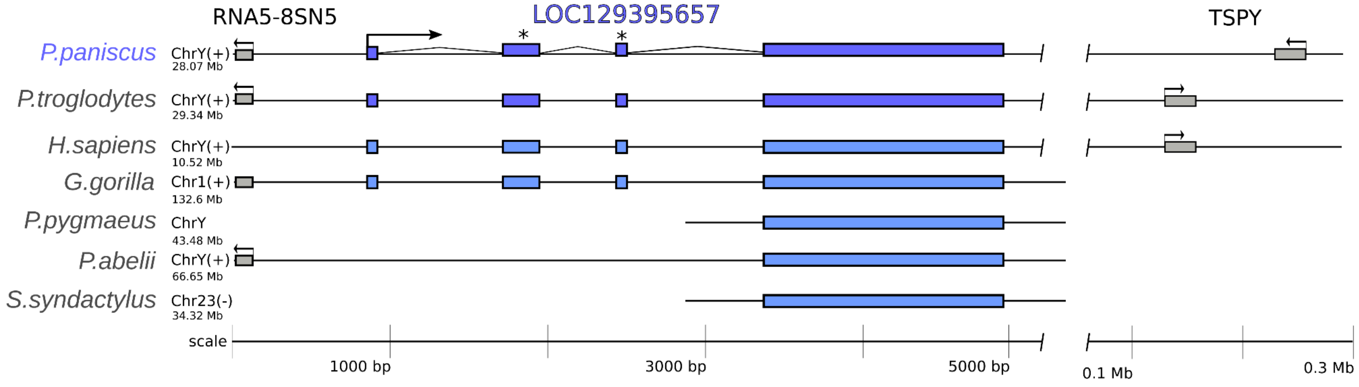

**Figure N11A. Emergence of the *de novo* gene candidate *LOC129395657* on the bonobo Y chromosome.** The *de novo gene* candidate (blue) emerged between conserved genes (grey) *RNA5-8SN5* and *TSPY* with homologous sequences in other primates. Most homologous sequences lack the ORF because of a missing start codon or premature stop codons (light purple). Stars indicate the exons located completely within the interspersed repeat region. Starting positions of the regions shown are taken from the respective chromosome assembly and are indicated below chromosome name and strand.

The *de novo* gene candidate *LOC129476750* in siamang comprises six exons with the ORF located in the first three exons (Fig. SN11B). The non-coding exon 6 is partially located in a simple repeat region. Exon 3 is partially located on a MER34B TE, that includes the stop codon completing the ORF in siamang. The *de novo* gene candidate is located between annotated genes *VCY* and *NLGN4Y*, but the *de novo* gene candidate itself is present in only one copy on the siamang Y. A homologous sequence is present on the human Y chromosome covering all exons, but lacking the ORF. Partial homologous sequences were detected on the Y chromosomes of chimpanzee and bonobo. Homologous sequences of exons 1 and 4-6 are present on X chromosomes of all great ape genomes assembled here, in syntenic regions between the conserved genes *VCX* and *NLGN4X.* The upstream regions of all homologous sequences for exon 1 were examined for TFBS motifs. The *de novo* gene candidate on the siamang Y chromosome contained the highest number of motifs compared to all other homologous hits, underlining the unique gene-like properties of *LOC129476750* in siamang.

In contrast to the bonobo *de novo* protein described above, the siamang *de novo* gene candidate *LOC129476750* encodes a relatively structured protein, not prone to aggregation but of low solubility, according to predictions (Table N11C). The secondary structure and 3D structure predictions suggest a helical protein with low amounts of disorder, which is expected to be rare in *de novo* proteins as structure is difficult to obtain from scratch[135].

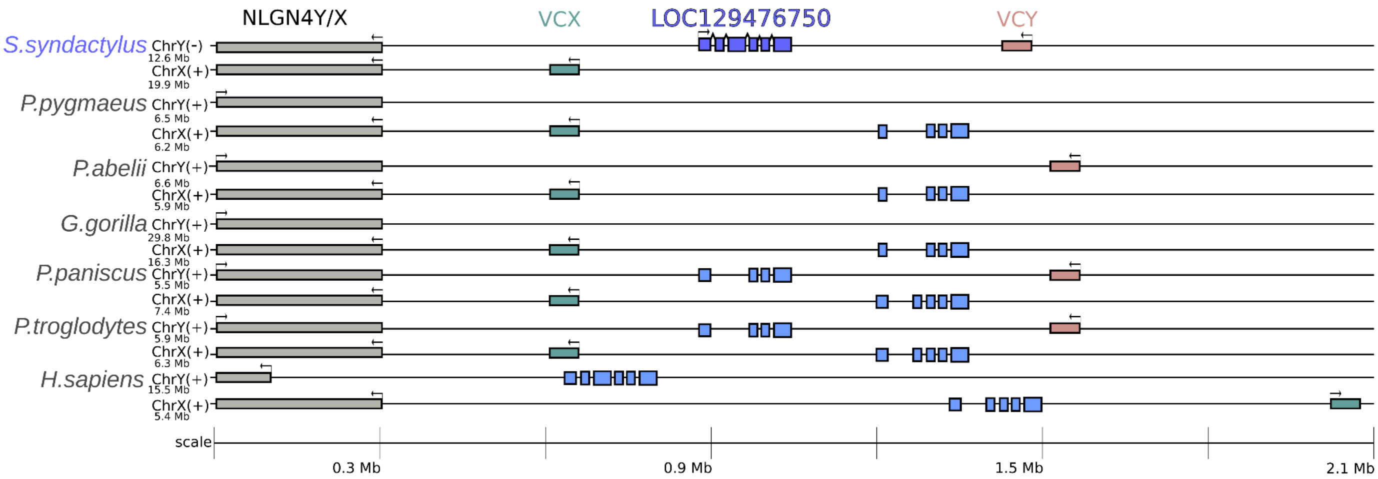

**Figure N11B. Emergence of the *de novo* gene candidate LOC129476750 on the siamang Y chromosome.** The *de novo gene* candidate (blue) emerged between conserved genes *NLGN4Y* (grey) and *VCY* (red), *NLGN4X* (grey) and *VCX* (green) in the corresponding X chromosome regions. Starting positions of the regions shown are taken from the respective chromosome assemblies and are indicated below chromosome name and strand.

**Table N11C. *De novo* protein properties**

| name | length | helix | sheet | coil | disorder (%) | disorder (mean) | solubility | aggregation |
|---|---|---|---|---|---|---|---|---|
| *LOC129395657* | 359 | 0.04 | 0.01 | 0.95 | 0.52 | 0.34 | high | negative |
| *LOC129476750* | 212 | 0.39 | 0.09 | 0.52 | 0.08 | 0.12 | low | negative |

**Discussion.** The predicted novel genes contained two *de novo* emerged genes specific to the ampliconic regions of the respective Y chromosomes. The *de novo* genes analyzed here were predicted as protein-coding by standard gene annotation pipelines (see Methods), making them high-confidence *de novo* proteins. Because of the imperfect gene annotation pipelines, we manually curated genes of interest. However, many low-confidence *de novo* gene candidates and newly emerging *de novo* genes, as well as the multiple copies of the bonobo *de novo* gene *LOC129395657,* might have been overlooked by our analyses because of our strict filters. While both candidate *de novo* genes are located within ampliconic regions, only the bonobo-specific *LOC129395657* is present in twelve copies on the bonobo Y. The siamang *de novo* candidate gene *LOC129476750* is only present once on the siamang Y chromosome. Both *de novo* gene candidates had exons located in TEs, possibly influencing their emergence. A previous study of *de novo* genes in Drosophila identified a connection between *de novo* gene emergence and TEs hypothesizing that highly mutable genomic regions around TEs may enable *de novo* gene birth[143].

#### Note S12. Chromosome-wide selection analysis

We performed selection analyses on chromosomes X and Y in chimpanzees and gorillas only, due to the limited sample size in the other great apes analyzed. Furthermore, we pooled subspecies for chimpanzees and gorillas resulting in datasets of 58 chimpanzees (36 females and 22 males) and 50 gorillas (39 females and 11 males). For chromosome X, we analyzed only females.

**Filtering.** We extracted chromosome X and Y datasets and filtered them separately with BCFtools[65]. We then filtered out ampliconic and repeated regions for the chromosome X datasets, and satellite, PAR, ancestral, ampliconic, and repeated regions for the chromosome Y datasets. We retained only biallelic SNPs and required a missing genotype rate of <50%. This resulted in 249,348 biallelic loci for the gorilla chrX dataset, 537,925 biallelic loci for the chimpanzee chrX dataset, 719 biallelic loci for the gorilla chrY dataset, and 1,095 biallelic loci for the chimpanzee chrY dataset.

Next, we calculated the proportion of missing genotypes per individual and removed outliers from our analysis. We removed one individual (Serufuli with 99.7% missing data) from the gorilla chrX dataset, four individuals (Noemie with 52.68%, Banyo with 43.35%, Annie with 42.48% , Kopongo with 31.13% missing data) from the chimpanzee chrX dataset, and three individuals (Alfred with 62.4%, Yogui with 62.31% , Brigitta with 61.86% missing data) from the chimpanzee chrY dataset. No samples were filtered out from the gorilla chrY dataset.

**Methods.** We calculated both nucleotide diversity (π)[144] and Tajima's D[145] on chromosome X in sliding windows of 100 kb with a step size of 20 kb and on the entire chromosome Y, for each species separately. Nucleotide diversity was computed from the site frequency spectrum as $\pi = \binom{n}{2}^{-1} L^{-1} \sum_{i=1}^{n-1} i(n-1)\xi_i$, where *L* is the length of the region in base pairs, $\xi_i$ is the count of sites with *i* copies of the alternate allele, and *n* is the sample size. To handle missing data for both nucleotide diversity and Tajima's D, we subsampled the site frequency spectrum down to the smallest sample size in each window using $\xi_j^h = \sum_{i=j}^{n-1} \xi_i^n \frac{\binom{i}{j}\binom{n-i}{h-j}}{\binom{n}{h}}$, where $\xi^k = \{\xi_1^k, \xi_2^k, \cdots, \xi_{k-1}^k\}$ is the site frequency spectrum for a sample of size $k$, $\xi_l^k$ is the count of sites with $l$ copies of the alternate allele, $n$ is the number of haplotypes in the full sample size, and $h < n$ is the largest number of haplotypes with non-missing data in the window (the target SFS size).

**Results.** The negative value for Tajima's D could suggest that the population size was recently reduced and/or that there was a recent positive or negative selection. Likewise, a positive value suggests a recent population expansion or balancing selection. The chromosome-X-wide distribution of Tajima's D values are plotted in Figure N12A for gorillas and Figure N12B for chimpanzees. Mean Tajima's D across windows was $-1.142881$ and $0.1796825$ chimpanzees and gorillas, respectively.

We identified genes in regions with extreme negative values by calculating the 1% quantile from the empirical distribution of D values, for each species separately, as indicated by the red horizontal lines plotted in Figures N12A and N12B. To identify genes in these extreme regions, consecutive windows meeting or exceeding this threshold were merged and then intersected with NCBI RefSeq gene annotations. These are given in Table N12A for gorilla and Table N12B for chimpanzee.

Because the non-recombining portion of chrY evolves as a single linkage group, we only computed the statistic for the entire chromosome, which resulted in $D = 0.110911$ and $D = -0.0230875$ for gorillas and chimpanzees, respectively.

The nucleotide diversity for the chimpanzee chromosome X was $\pi_X^C = 1.273\times10^{-3}$ and for the chimpanzee chromosome Y was $\pi_Y^C = 7.660 \times 10^{-4}$ resulting in a Y/X ratio of $\pi_Y^C/\pi_X^C = 0.6017$. The nucleotide diversity for the gorilla chromosome X was $\pi_X^G = 1.023 \times 10^{-3}$ and for the gorilla chromosome Y was $\pi_Y^G = 1.950 \times 10^{-4}$

resulting in a Y/X ratio of $\pi_Y^G/\pi_X^G = 0.1906$.

Figures N12C and N12D plot the window-based nucleotide diversity results for gorilla and chimpanzee X chromosomes, respectively.

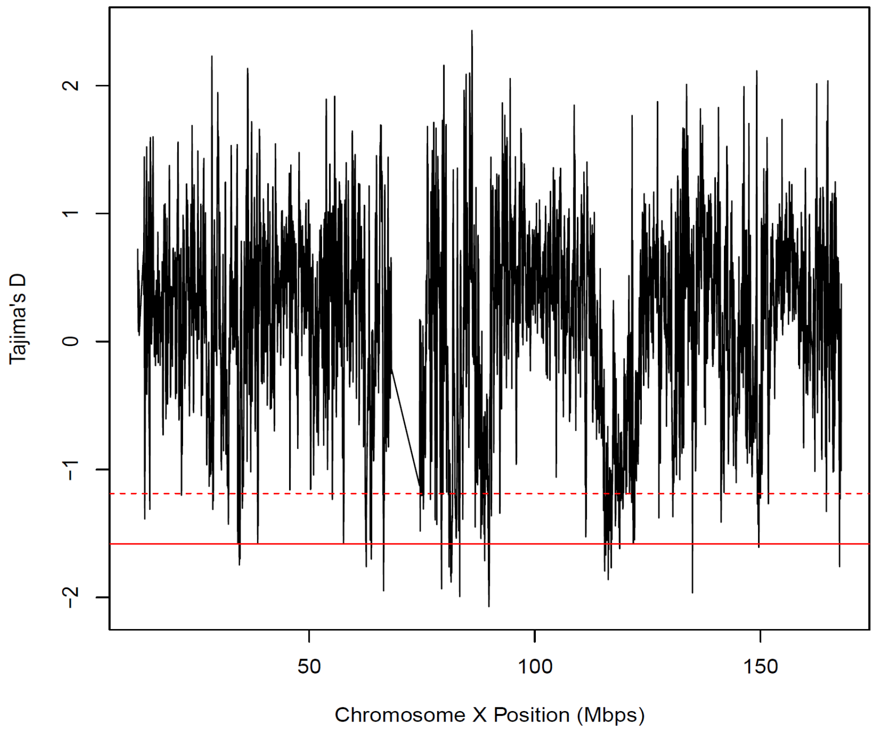

**Figure N12A. Tajima's D in 100-kb sliding windows with 20-kb steps for gorilla chrX.** The dashed red line indicates the most extreme 5% quantile and the solid red line indicates the most extreme 1% quantile.

**Table N12A. Top 1% Tajima's D regions on gorilla X and their associated genes**

| Start | End | Associated genes |
|---|---|---|
| 34460000 | 34619999 | *KLHL15,EIF2S3* |
| 34620000 | 34759999 | *LOC115932527,ZFX,LOC129530076* |
| 38560000 | 38659999 | -- |
| 57560000 | 57659999 | *RBM10,UBA1,CDK16,USP11* |
| 62540000 | 62699999 | *XAGE2,LOC115932194,LOC101148088* |
| 63700000 | 63839999 | *HUWE1* |
| 66440000 | 66599999 | -- |
| 79240000 | 79399999 | *AR* |
| 80940000 | 81059999 | *LOC101151098* |
| 81320000 | 81679999 | *NALF2,EDA,LOC115932441* |
| 83300000 | 83419999 | *OGT,GCNA* |
| 88840000 | 88939999 | -- |
| 89660000 | 89979999 | *ATP7A,PGK1,TAF9B,CYSLTR1* |
| 115400000 | 115539999 | *DHRSX,TCEAL1,MORF4L2,LOC129530127,GLRA4,PLP1,LOC129530130* |
| 115560000 | 115759999 | *DHRSX,PLP1,LOC129530129,RAB9B,LOC101142541,LOC101145971* |
| 116220000 | 116319999 | *DHRSX* |
| 116340000 | 116439999 | *DHRSX,IL1RAPL2,LOC101146664* |
| 116860000 | 116979999 | *DHRSX,IL1RAPL2* |
| 118720000 | 118859999 | *LOC101124366,MORC4,RBM41* |
| 121820000 | 121919999 | *ARSD,TMEM164* |
| 134860000 | 134999999 | *GRIA3* |
| 149580000 | 149679999 | -- |
| 167460000 | 167579999 | *VAMP7,LOC115932124* |

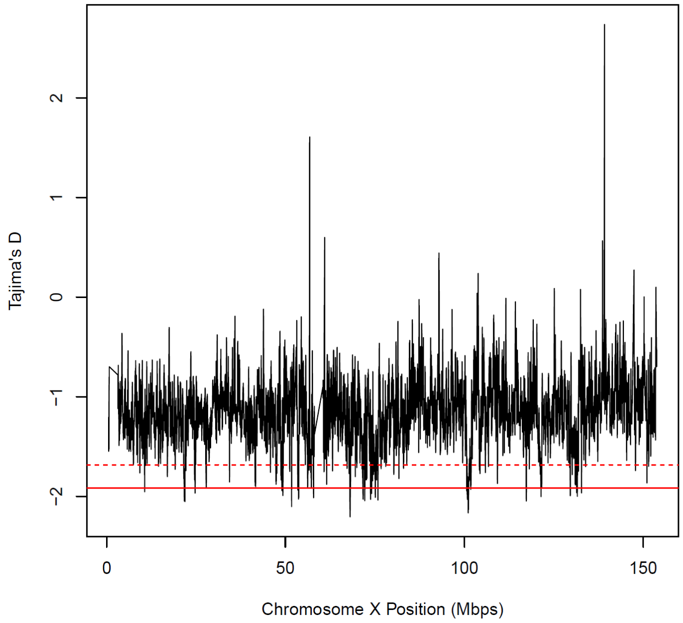

**Figure N12B. Tajima's D in 100-kb sliding windows with 20-kb steps for the chimpanzee chrX.** The dashed red line indicates the most extreme 5% quantile and the solid red line indicates the most extreme 1% quantile.

**Table N12B. Top 1% Tajima's D regions for chimpanzee X and their associated genes**

| Start | End | Associated genes |
|---|---|---|
| 10580000 | 10679999 | *WWC3,COL4A6,COL4A5,IRS4,LOC129138600,LOC112207135* |
| 21660000 | 21819999 | *LOC129138882,LOC107971195,LOC112207136* |
| 21820000 | 21959999 | *LOC129138882* |
| 24680000 | 24799999 | *LOC112207723,LOC129138882,LOC112207193,ZFX* |
| 49000000 | 49099999 | *LOC112207723,LOC112207154,GLOD5,LOC112207116,GATA1,HDAC6* |
| 49260000 | 49379999 | *LOC112207723,KCND1,GRIPAP1,TFE3,CCDC120,PRAF2,WDR45,LOC112207276* |
| 51680000 | 51799999 | *LOC112207723,LOC473612,LOC107970979,GSPT2* |
| 53440000 | 53539999 | *LOC112207723,SMC1A,LOC112206922,RIBC1,HSD17B10* |
| 53620000 | 53799999 | *LOC112207723,HUWE1* |
| 56180000 | 56279999 | *LOC112207723,KLF8* |
| 57820000 | 57939999 | *LOC112207723* |
| 67980000 | 68219999 | *LOC104004304,DGAT2L6,AWAT1,P2RY4,ARR3,RAB41,PDZD11,KIF4A,LOC107971127,GDPD2,LOC101059407* |
| 71640000 | 71839999 | *PUDP,LOC100615774,LOC107971096,LOC107971130,LOC129138865* |
| 72160000 | 72279999 | *PUDP,SLC16A2* |
| 73620000 | 73759999 | *LOC104004678,PUDP* |
| 73900000 | 74059999 | *LOC104004678,PUDP,PBDC1* |
| 74960000 | 75099999 | *LOC104004678,PUDP* |
| 75720000 | 75819999 | *LOC107971128,PUDP,ATP7A,PGAM4* |
| 100560000 | 100659999 | *LOC100612348* |
| 100700000 | 100859999 | *LOC100612348,LOC465775,BEX4* |
| 100960000 | 101159999 | *TCEAL9,BEX3,LOC100614699,LOC129138744,RAB40A* |
| 101680000 | 101799999 | *TBL1X,FAM199X,ESX1* |
| 117180000 | 117359999 | *ARHGAP6,SEPTIN6,SOWAHD,LOC737451,LOC112207195,LOC129138610* |
| 121340000 | 121439999 | *ARHGAP6,XIAP,LOC112207171* |
| 129540000 | 129659999 | *FRMPD4,FRMD7* |
| 131320000 | 131419999 | *FRMPD4,GPC3* |
| 131500000 | 131599999 | *FRMPD4* |
| 132660000 | 132779999 | *FRMPD4,LOC104004010,LOC129138847,ZNF75D,LOC112206982* |

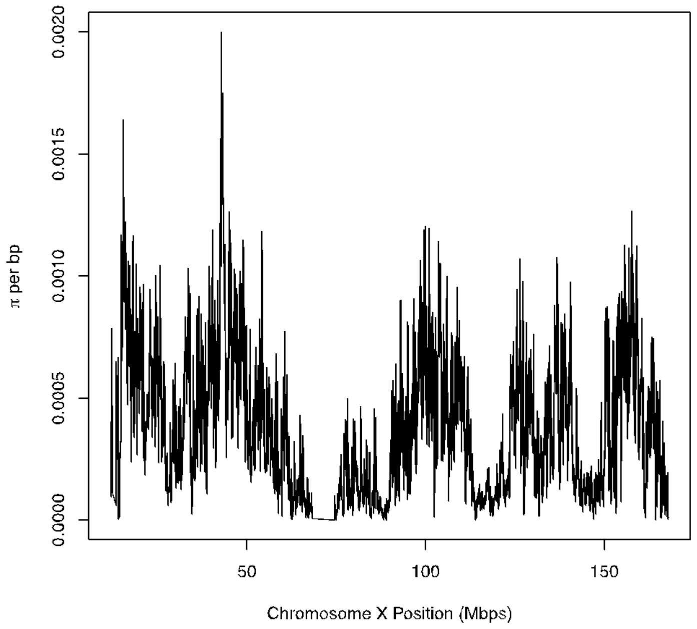

**Figure N12C. Nucleotide diversity in 100-kb sliding windows with 20-kb steps for the gorilla chrX.**

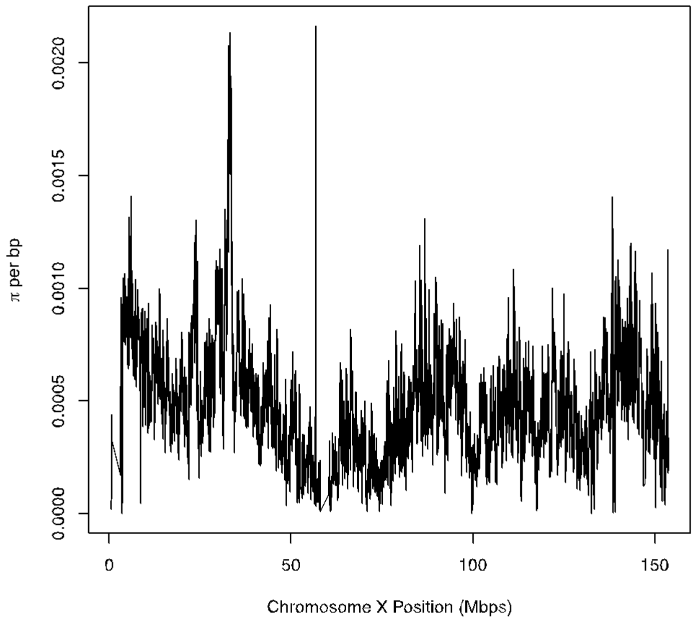

**Figure N12D. Nucleotide diversity in 100-kb sliding windows with 20-kb steps for the chimpanzee chrX.**

We next examined the Y chromosome of chimpanzee and gorilla for evidence of natural selection in more detail. Since there is no recombination for most parts of the Y (outside of PARs), there are no 'local' effects of selection on genealogy and only chromosome-wide effects can be observed. Both directional positive and negative selection will reduce genetic diversity, with positive selection further shifting the site frequency spectrum (SFS) towards an inflation of low-frequency variants.

We first compared the nucleotide diversity of the Y chromosome to that of the X chromosome. The expected ratio under neutral conditions hinges on the male-to-female effective population size ratio ($N_m/N_f$), the mutation rate difference between the X and the Y, and the specific demographic model. For populations of constant size, coalescence theory provides analytical expectations. The Y chromosome's effective population size is half that of the male's ($N_Y = N_m/2$), while that of the X chromosome depends on both male and female effective population sizes, $N_X = 9N_mN_f/(4N_m + 2N_f)$ [146]. Using these relations, the $N_Y/N_X$ ratio becomes $(2N_m+N_f)/9N_f$. The lowest possible $N_Y/N_X$ value, in mating systems with very low male effective population size, is thus 1/9. For systems with equal male and female effective population size, the ratio is 1/3 (Fig. N12E).

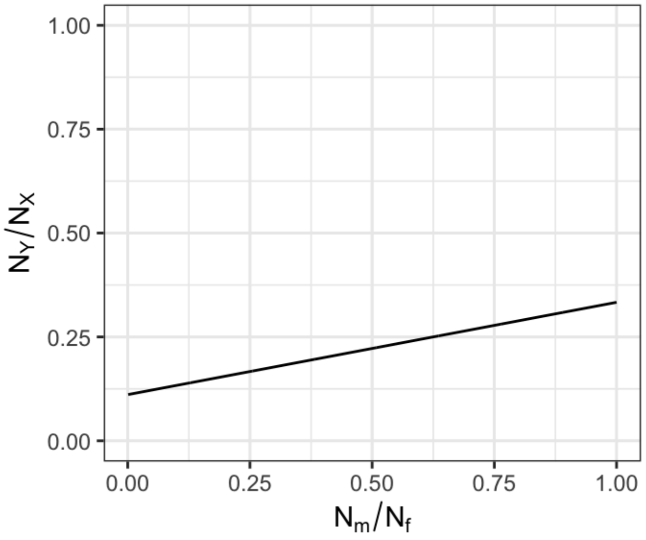

**Figure N12E. Relationship between the Y to X effective population size as a function of the male-to-female effective population size in a neutral model with constant population size.**

However, chromosome diversity can also be impacted by mutation rate differences. In both chimpanzee and gorilla, the Y chromosome substitution rate was found to be 1.8-2 times higher than that for the X chromosome (Fig. 1D). Thus, assuming neutrality, constant population size, and a doubled Y chromosome mutation rate, the lowest expected Y/X diversity ratio is 0.22 in case of very low male-to-female effective population size. For equal male and female effective population sizes, the expected Y/X diversity ratio is 0.67.

In our chimpanzee data, the Y/X diversity ratio stands at 0.602, while for gorillas it is 0.191. This means that, under the assumption of constant population size, gorillas have a Y/X diversity ratio consistent with a significantly low male effective population size ($N_m < 0.1\ N_f$). In contrast, chimpanzees seem to have a male-to-female effective population size that is more balanced. These results are broadly consistent with a smaller male effective population size in gorillas than in chimpanzees due to polyandrous mating in the former[147].

However, non-equilibrium demographics and population structure might also skew the Y-to-X genetic diversity ratio. To assess this effect, we simulated genetic variation data based on previously inferred demographic models for chimpanzees[69] and gorillas[71]. We adjusted population sizes in the model according to Wilson Sayres et al.'s approach[146], simulating separate X and Y chromosome population sizes to match a specific male-to-female ratio. We adjusted the X chromosome's recombination rate to 2/3 of the autosomal value to reflect the lack of recombination in males and set the Y chromosome's recombination rate to zero. An autosomal mutation rate was assumed for the X chromosome and a doubled rate for the Y.

Our simulations indicate that the expected Y/X diversity ratio, under realistic demographic models, exceeds that of constant-sized populations (Fig. N12F). Even for low male-to-female effective population size, the simulated Y/X diversity ratio is on average 0.8. This suggests that our observed Y/X diversity ratio of 0.191 in gorilla is unexpectedly low, aligning with the hypothesis of selection reducing diversity on the non-recombining Y chromosome, as was suggested for the human Y[146]. In chimpanzee, the observed Y/X diversity ratio of 0.602 seems consistent with a neutral model only for a very low male effective population size ($N_m < 0.25\ N_f$).

However, if male effective population size is higher, which is likely the case for chimpanzees [147], the observed pattern would also be consistent with selection reducing diversity on the Y.

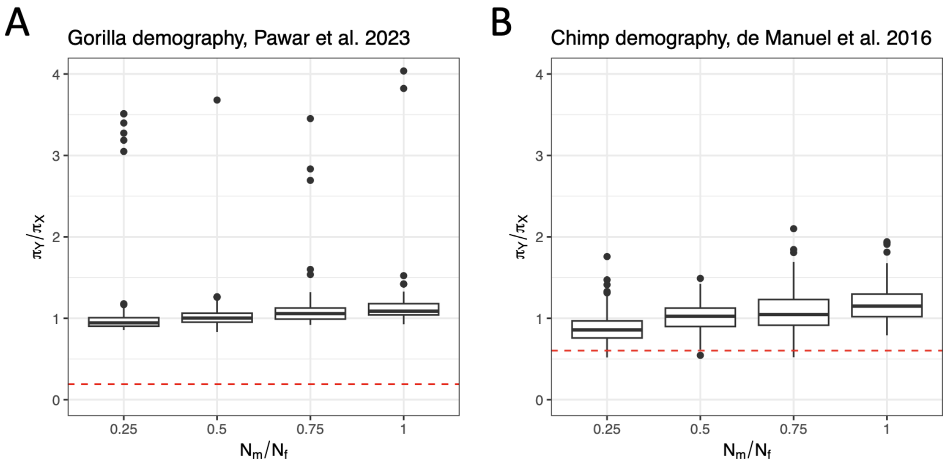

**Figure N12F. Relationship between the Y-t- X nucleotide diversity ratio as a function of the male-to-female effective population size in demographic models previously estimated for gorilla (A) and chimpanzee (B).** We assume a twice as large mutation rate on the Y than on the X. Boxplots show the distribution of 80 replicate whole-chromosome simulations.

The strongly diminished diversity on the Y chromosome in gorilla could result from background selection due to harmful mutations, consistent with observed purifying selection on the Y chromosome ($d_N/d_S$ < 1 for ancestral genes in both species). However, recent positive selection might also lead to reduced linked neutral diversity on the Y. Moreover, a selective sweep would increase the abundance of rare mutations, thereby decreasing Tajima's D. The measured Tajima's D values for chimpanzee and gorilla on the Y are -0.0230875 and 0.110911, respectively. Simulations using demographic models indicate these values align well with those simulated under neutrality for a wide range of male-to-female population size ratios (Fig. N12F). Only for gorilla, at very low levels of male-to-female population size ratio, the observed Tajima's D value appears at the lower end of the distribution consistent with the importance of recent positive selection. Note however that purifying selection at non-recombining sequences can also lead to a decrease in Tajima's D at neutral sites[148]. Further, sex-biased migration could lead to higher levels of population structure experienced by the Y chromosome than predicted from a demographic model that was estimated using autosomal data. Both effects on the expected distribution of Tajima's D are currently not modeled. In sum, purifying selection is the more parsimonious explanation for patterns of diversity on the Y in gorillas and chimpanzees.

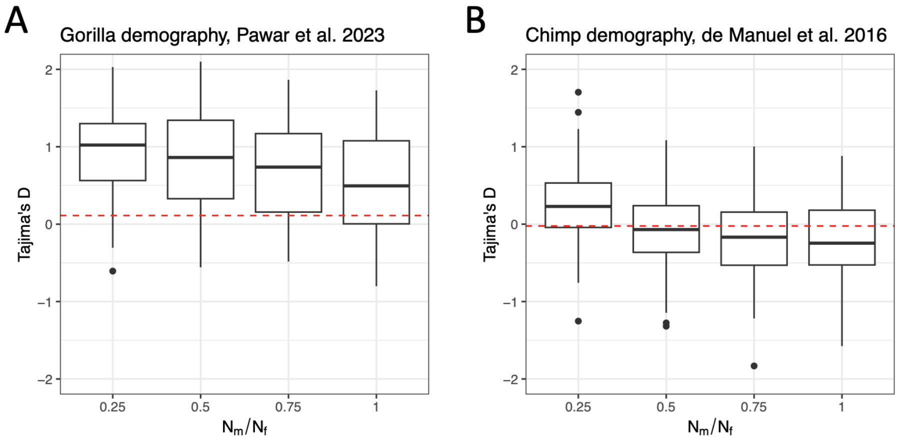

**Figure N12G. Relationship between simulated Tajima's D for the Y chromosome as a function of the male-to-female effective population size in demographic models previously estimated for gorilla (A) and chimpanzee (B).** Boxplots show the distribution of 80 replicate whole-chromosome simulations.

### Note S13. Selection analysis of ancestral (X-degenerate) Y genes using diversity data

**Phylogenetic inference.** We examined the phylogenetic history of the X and Y chromosomes using individuals with XY karyotypes (N = 80). The results are presented in Figures N13A and N13B.

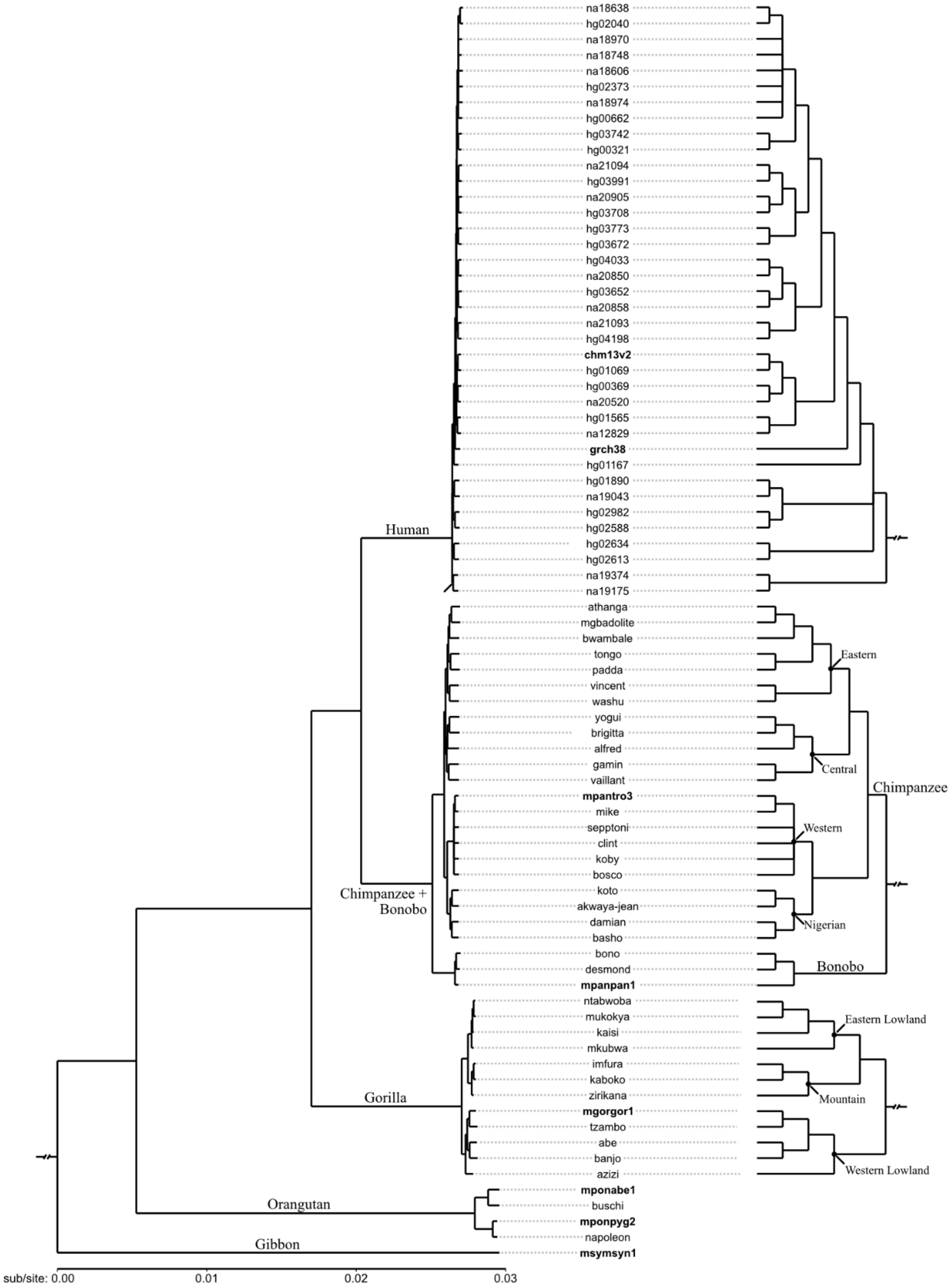

**Figure N13A.** Evolutionary history of X chromosomes among primate populations. Maximum-likelihood phylogeny (left) is presented as cladograms (right) to highlight relationships among short branches. All nodes with ultrafast bootstrap support ≤95% are collapsed as polytomies and bolded names reflect tips associated with reference assemblies.

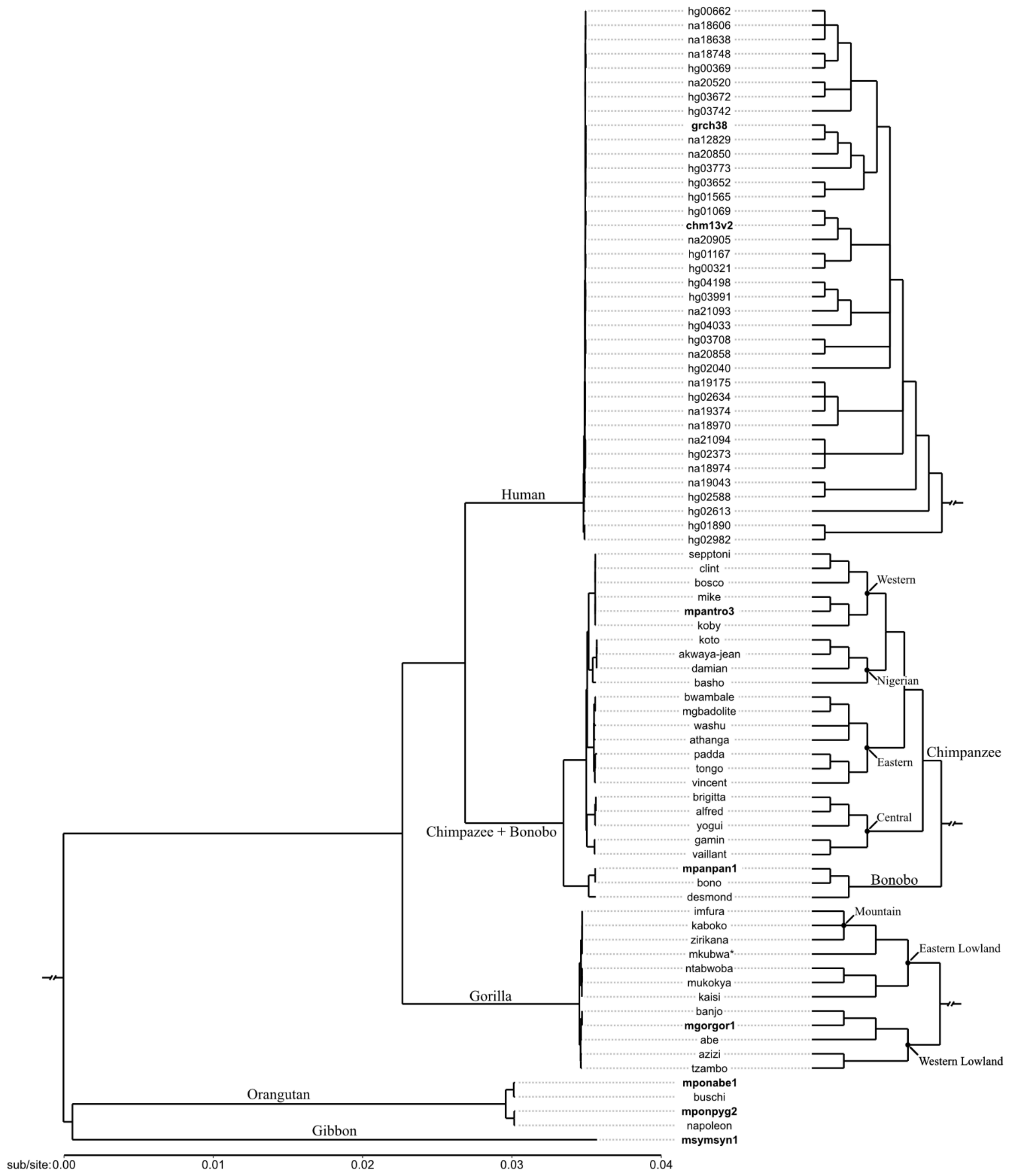

**Figure N13B.** Evolutionary history of Y chromosomes among primate populations. Maximum-likelihood phylogeny (left) is presented as cladograms (right) to highlight relationships among short branches. All nodes with ultrafast bootstrap support ≤95% are collapsed as polytomies and bolded names reflect tips associated with reference assemblies.

Though the species-level topologies (Fig. N13C) of both the X and Y chromosome phylogenies are concordant and agree with previously published data[22], we observed discordance within species lineages (normalized topological distance[149] = 0.61, where 0 reflects complete concordance and 1 reflects complete discordance). In addition to differences in branching order within populations, topological disagreement was present throughout the human subclade, where the evolutionary history of chromosome X broadly disagrees with previously published chromosome Y haplogroup relationships[150]. The relationships between the populations of mountain and eastern lowland gorillas reflect two unique evolutionary histories. Whereas these two populations appear reciprocally monophyletic for chromosome X, the evolutionary history of chromosome Y suggests a paraphyletic relationship, with the mountain population's chromosome Y being derived from the eastern lowland population. By computing the mean evolutionary distance between all tips, we also observed elevated rates of sequence evolution for chromosome Y (1.88e-2 ± 1.22e-2, mean substitutions/site ± s.d.) when compared with chromosome X (1.30e-2 ± 8.81e-3).

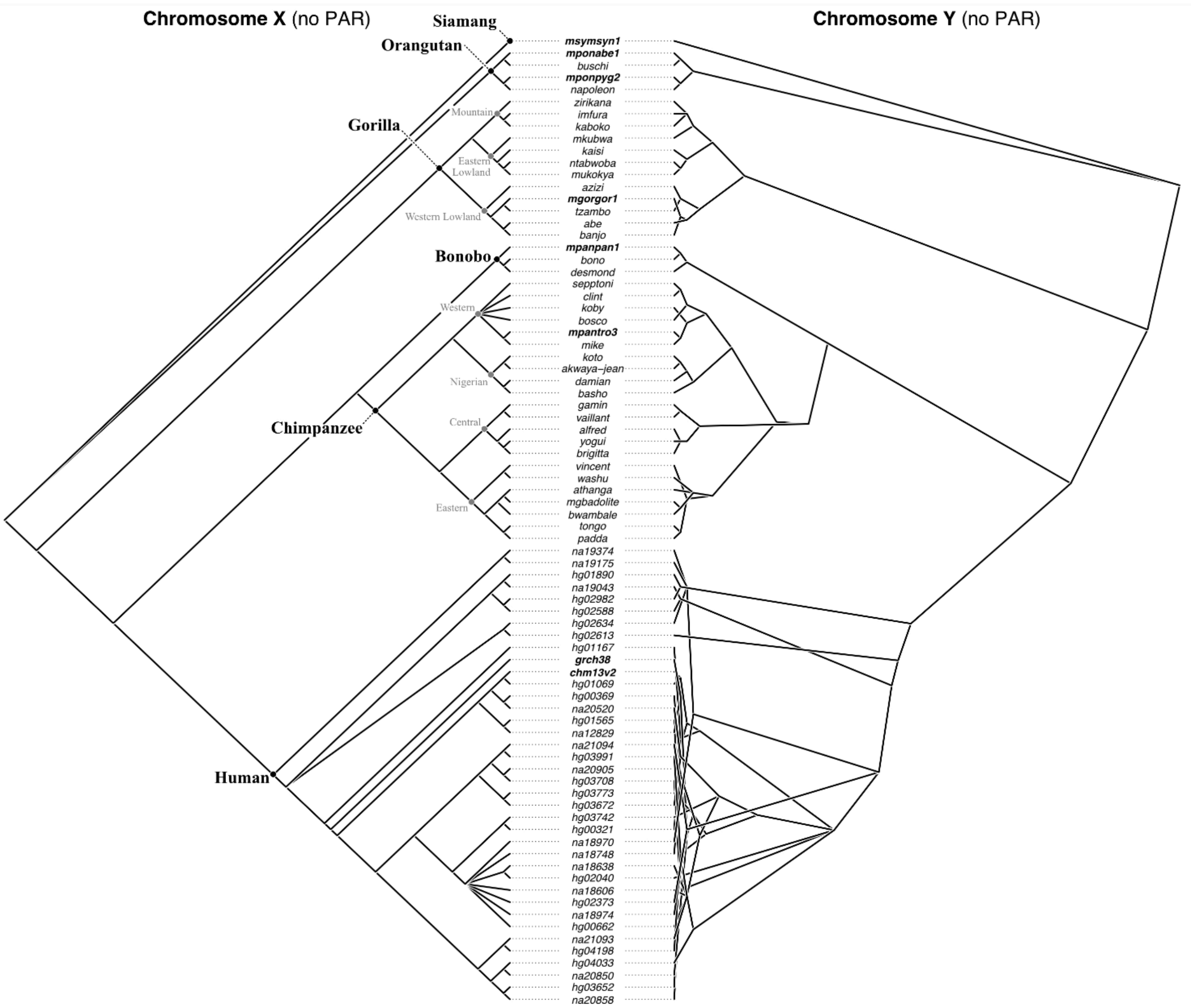

**Figure N13C.** Cotangle plot highlighting discordance between X and Y chromosome phylogenies. Topology of the X chromosome cladogram (left) is fixed, whereas the topology of the Y chromosome cladogram is permitted to rotate and overlap. Both topologies' branches do not possess units of length.

**Branch-site tests of positive selection.** Following phylogenetic reconstruction, we performed branch-site tests of positive selection using ancestral genes present as single copies across all sampled taxa. Although *UTY* showed evidence of episodic positive selection (*q*-value = 3.48e-3; FDR cutoff = 0.05) in the ancestral lineage to all modern chimpanzees, closer inspection of the underlying codon alignment revealed this signal was driven by a 5' indel exclusive to the chimpanzee lineage. Thus, while unique polymorphism exists within the chimpanzee lineage (relative to all other sampled primates), non-synonymous substitution bias alone does not explain this observation.

**McDonald-Kreitman selection tests**. To identify potential signatures of selection on the Y chromosome, we conducted the McDonald-Kreitman (MK) test[151] on the ancestral genes. The alignments we used for the MK test were the same alignments used for the branch-site tests of positive selection above. We focused our analysis on male chimpanzees and gorillas, for which there was sufficient population sequencing data available for our analysis (21 non-reference individuals for chimpanzee and 11 non-reference individuals for gorilla).

In chimpanzees, there were only three ancestral genes on the Y that had a sufficient number of segregating sites to allow the MK test α value to be calculated. These genes were *KDM5D*, *UTY*, and *ZFY*. *KDM5D* had an α of -1.25 (*p*=0.658), *UTY* had an α of 0.267 (*p*=1) and finally, for *ZFY*, α was 0.524 (*p*=0.638). An analysis concatenating all ancestral genes on the Y showed an α value of 0.444 across all genes with a *p*-value of 0.114.

In gorillas, only one gene, *UTY,* had a sufficient number of segregating sites to conduct an MK test. Its α value was 1 and the *p*-value was 0.237. We then concatenated all the genes, as we did with the chimpanzees, and ran a whole-chromosome MK test on all ancestral genes on the Y. The results of this analysis were also not significant with an α of -0.846 (*p*=1).

#### Methods

**Phylogenetic history of primate populations.** To gain insight into the evolutionary history of X and Y chromosomes among great ape populations, XY-karyotyped samples were used to reconstruct phylogenetic trees where each phylogeny was composed of the same individuals–allowing for direct comparisons between topologies. One chimpanzee sample, Taweh, was removed due to an uncertain karyotype. Regarding human sample selection for representation within the phylogenetic dataset, the 1KGP sample with the highest mean sequencing depth (genome-wide) was selected for each chromosome Y haplogroup[150]. In addition to population samples, each primate T2T assembly and GRCh38 were included.

Prior to alignment, pseudoautosomal regions were removed from each species' variant call file, as regions of recombination between X and Y may interfere with X- and Y-specific phylogenetic inference. For each XY karyotyped individual selected for phylogenetic inference, `bcftools consensus` (version 1.9) [65] was used to emit sample-specific short variants onto each species' respective T2T X/Y assembly (`--haplotype A` was used to project alternative alleles). X and Y chromosome alignments were generated using CACTUS (version 3.1.20211107152837)[37]. Intraspecies guide-tree relationships were estimated using the neighbor-joining topology computed from genotype Manhattan distances using the `vcfR` (version 1.14.0) [152] and ape (version 5.7)[153] packages in R (version 4.3.0). The interspecies guide-tree topology was obtained from[154].

Each cactus alignment graph was converted to Multiple Alignment Format (MAF) using `hal2maf` (version 2.2)[38] with T2T-CHM13 set as the reference assembly, pseudoautosomal regions removed (Table S9), ancestral sequences removed (`--noAncestors`), and parology edges removed (`--onlyOrthologs`). A custom BioPython script was used to extract 1-to-1 orthology blocks and convert the alignment format to FASTA, where each extracted alignment block contained a single sequence per species. X and Y maximum-likelihood phylogenies were inferred using IQTree (version 2.0.3)[155] with the best-fit substitution model estimated by ModelFinder[40] and node support estimated using 10,000 ultrafast bootstrap replicates[41]. Nodes with <95% ultrafast bootstrap support were collapsed as polytomies.

**Branch-site tests of positive selection.** Coding sequences for ancestral genes on chromosome Y were extracted for each XY-karyotyped individual using vcf2fasta (https://github.com/santiagosnchez/vcf2fasta), where a manually curated annotation was applied for chrY to identify the coordinates for each gene's CDS. Only genes present as single copies across all species were used in these analyses. These include the following: *AMELY, DDX3Y, KDM5D, RPS4Y1, RPS4Y2, SRY, TMSB4Y, UTY*, and *ZFY* for chrY. As each annotation dataset is composed of multiple isoforms for each gene, only the longest isoform for each gene was used in subsequent analyses of positive selection (isolated using AGAT; version 1.2.0)[156]. For each gene, a codon alignment was generated using MACSE (version 2.07)[157].

The adaptive branch-site random effects likelihood (aBSREL) model implemented in HyPhy (version 2.5.50)[158] was used to test for signatures of episodic positive selection. Input gene alignments were cleaned using Gblocks (version 0.91b)[159] with default parameters prior to processing with HyPhy. Foreground branches were specified separately as each population, species, and multi-species clade (e.g., western chimpanzees, *Pan troglodytes*, and the monophyletic assemblage of Pan+Homo+Gorilla, respectively). To accommodate multiple testing, aBSREL *p*-values were adjusted using the qvalue package (version 2.23.0)[160] where the false discovery rate was set to 0.05 (*q*-value≤0.05).

**McDonald-Kreitman tests.** We started by examining each ancestral gene individually. To do so, we used a multi-fasta alignment of the T2T reference for the species, the non-reference individuals making up the population data, and the human T2T reference (used as the outgroup to measure divergence). To determine synonymous and non-synonymous polymorphisms within our population samples for each species, we used iMKT[161] . iMKT also calculated the divergence from the aligned human T2T outgroup sequence. The multi-fasta option in iMKT considers four-fold degenerate sites and zero-fold degenerated selected sites when determining synonymous and non-synonymous polymorphisms. From the counts of these polymorphisms, we estimated the proportion of substitutions under positive selection (α) and conducted a Fisher's exact test to determine statistical significance.

Due to small sample size, gene lengths, and limited diversity between individuals in the population, we did not have enough statistical power to detect selection on individual genes. Therefore, we sought to increase the power of the test by concatenating the divergence and polymorphism output files generated by iMKT for each gene into a larger file containing the values from all Y chromosome ancestral genes. Using this concatenated file, we ran another MK test on all ancestral genes combined from the Y chromosome.

### Note S14. The analysis of chromosome Y phylogenies and TMRCA using new references and variant calls

**Methods.** Previously published male chimpanzee, bonobo, and gorilla resequencing datasets[68–70] were used for the construction and dating of the Y-chromosomal phylogeny. All confident sites were called using GATK[74], only ancestral regions were used, followed by removal of indels, calls where ≥10 % of high-quality reads supported another allele and sites with >9% of missing genotypes. The Bayesian Markov chain Monte Carlo phylogenetics software BEAST v1.10.4[78] was used to estimate the time-to-most-recent common ancestor (TMRCA) for the nodes of interest (see Supplementary Methods for details).

**Results.** Using the species-level variant set to reconstruct and date the intraspecific Y chromosome phylogenies (Fig. S17; Table S42), we found the overall topology to be nearly identical to the one built previously[162] for the samples that overlapped between the current and previous[162] analyses. The estimated TMRCAs were also generally in line with previous estimates[162].

**Figure N14A. MSY phylogenies for chimpanzees (top), bonobos (middle), and gorillas (bottom) with times to the most recent common ancestor (TMRCA).** Branch lengths are drawn proportional to the estimated times between successive splits according to BEAST analysis. Point estimates of TMRCA are given adjacent to the nodes with the 95% highest posterior density (HPD) intervals shown in squared brackets. TMRCA estimates using two mutation rates are shown – the Helgason et al. 2015 mutation rate estimate (top) and the Fu et al. 2014 mutation rate (bottom, in brackets; see Supplemental Methods - Y chromosome phylogeny and TMRCA calculations) [79,80]. Species/subspecies are indicated, and names of individuals are given at the tips of branches followed by their species/subspecies designation as follows: PTT - *Pan troglodytes troglodytes*; PTS - *P. t. schweinfurthii*; PTE - *P. t. ellioti*; PTV - *P. t. verus*; PPA - *Pan paniscus*; GGG - *Gorilla gorilla gorilla*; GBB - *G. beringei beringei*; GBG - *G. b. graueri*. kya – thousand years ago.

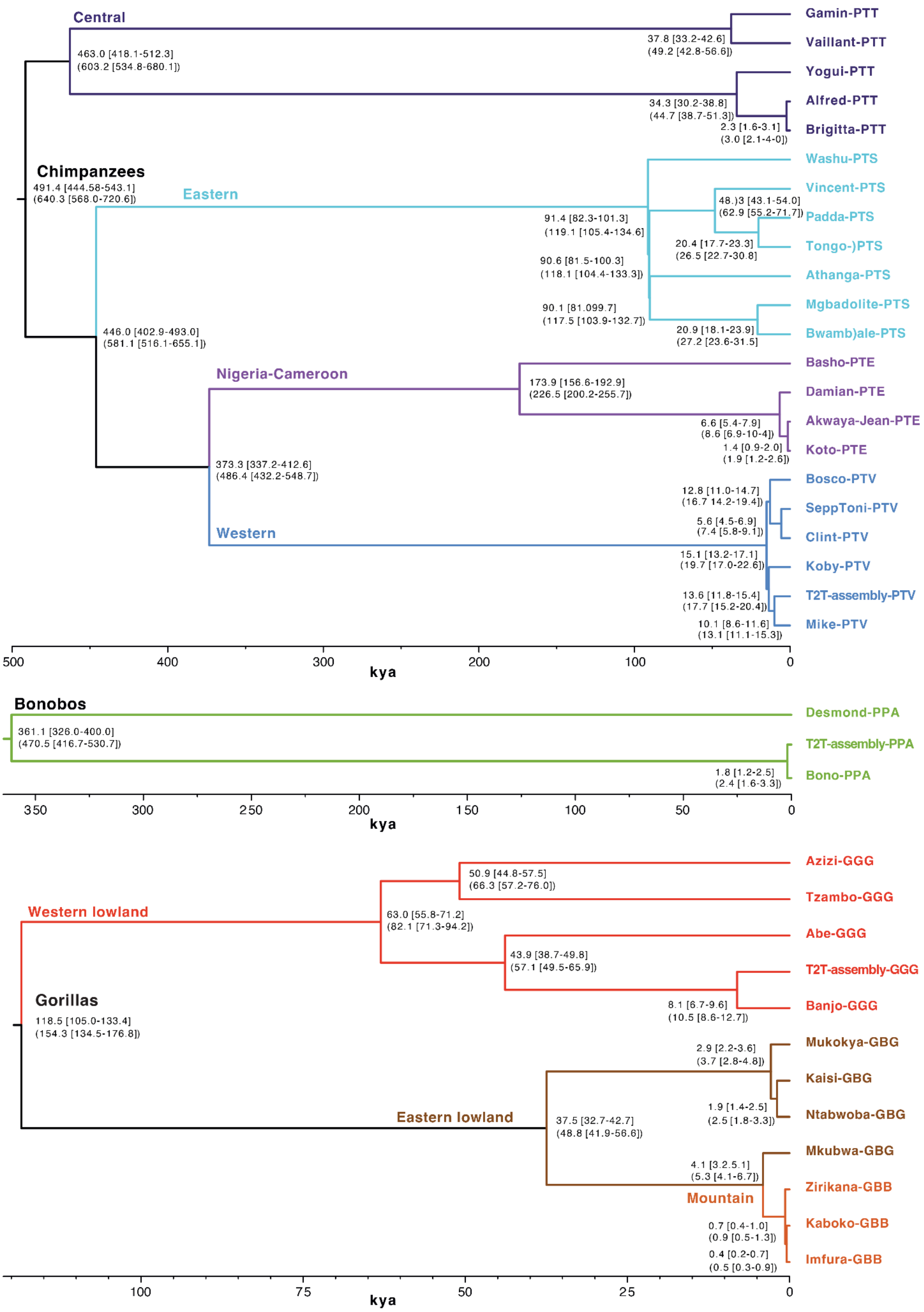
Central
Gamin-PTT
Vaillant-PTT
Yogui-PTT
Alfred-PTT
Brigitta-PTT
463.0 [418.1-512.3]
(603.2 [534.8-680.1])
37.8 [33.2-42.6]
(49.2 [42.8-56.6])
34.3 [30.2-38.8]
(44.7 [38.7-51.3])
2.3 [1.6-3.1]
(3.0 [2.1-4-0])
Chimpanzees
491.4 [444.58-543.1]
(640.3 [568.0-720.6])
Eastern
Washu-PTS
Vincent-PTS
Padda-PTS
Tongo-)PTS
Athanga-PTS
Mgbadolite-PTS
Bwamb)ale-PTS
48.)3 [43.1-54.0]
(62.9 [55.2-71.7])
91.4 [82.3-101.3]
(119.1 [105.4-134.6]
20.4 [17.7-23.3]
(26.5 [22.7-30.8]
90.6 [81.5-100.3]
(118.1 [104.4-133.3])
90.1 [81.099.7]
(117.5 [103.9-132.7])
20.9 [18.1-23.9]
(27.2 [23.6-31.5]
446.0 [402.9-493.0]
(581.1 [516.1-655.1])
Nigeria-Cameroon
Basho-PTE
Damian-PTE
Akwaya-Jean-PTE
Koto-PTE
173.9 [156.6-192.9]
(226.5 [200.2-255.7])
6.6 [5.4-7.9]
(8.6 [6.9-10-4])
1.4 [0.9-2.0]
(1.9 [1.2-2.6])
373.3 [337.2-412.6]
(486.4 [432.2-548.7])
Western
Bosco-PTV
SeppToni-PTV
Clint-PTV
Koby-PTV
T2T-assembly-PTV
Mike-PTV
12.8 [11.0-14.7]
(16.7 14.2-19.4])
5.6 [4.5-6.9]
(7.4 [5.8-9.1])
15.1 [13.2-17.1]
(19.7 [17.0-22.6])
13.6 [11.8-15.4]
(17.7 [15.2-20.4])
10.1 [8.6-11.6]
(13.1 [11.1-15.3])
500
400
300
200
100
0
kya
Bonobos
Desmond-PPA
T2T-assembly-PPA
Bono-PPA
361.1 [326.0-400.0]
(470.5 [416.7-530.7])
1.8 [1.2-2.5]
(2.4 [1.6-3.3])
350
300
250
200
150
100
50
0
kya
Western lowland
Azizi-GGG
Tzambo-GGG
Abe-GGG
T2T-assembly-GGG
Banjo-GGG
50.9 [44.8-57.5]
(66.3 [57.2-76.0])
63.0 [55.8-71.2]
(82.1 [71.3-94.2])
43.9 [38.7-49.8]
(57.1 [49.5-65.9])
8.1 [6.7-9.6]
(10.5 [8.6-12.7])
Gorillas
118.5 [105.0-133.4]
(154.3 [134.5-176.8])
Eastern lowland
Mukokya-GBG
Kaisi-GBG
Ntabwoba-GBG
Mkubwa-GBG
2.9 [2.2-3.6]
(3.7 [2.8-4.8])
1.9 [1.4-2.5]
(2.5 [1.8-3.3])
37.5 [32.7-42.7]
(48.8 [41.9-56.6])
4.1 [3.2.5.1]
(5.3 [4.1-6.7])
Mountain
Zirikana-GBB
Kaboko-GBB
Imfura-GBB
0.7 [0.4-1.0]
(0.9 [0.5-1.3])
0.4 [0.2-0.7]
(0.5 [0.3-0.9])
100
75
50
25
0
kya

doi:10.1038/s41586-023-06457-y.

14. Kearse, M. *et al.* Geneious Basic: an integrated and extendable desktop software platform for the organization and analysis of sequence data. *Bioinformatics* **28**, 1647–1649 (2012).
15. Dutra, A. S., Mignot, E. & Puck, J. M. Gene localization and syntenic mapping by FISH in the dog. *Cytogenet. Cell Genet.* **74**, 113–117 (1996).
16. Mao, Y. *et al.* Structurally divergent and recurrently mutated regions of primate genomes. *bioRxiv* 2023.03.07.531415 (2023) doi:10.1101/2023.03.07.531415.
17. Baid, G. *et al.* DeepConsensus improves the accuracy of sequences with a gap-aware sequence transformer. *Nat. Biotechnol.* **41**, 232–238 (2023).
18. Robinson, J. T., Thorvaldsdóttir, H., Wenger, A. M., Zehir, A. & Mesirov, J. P. Variant Review with the Integrative Genomics Viewer. *Cancer Res.* **77**, e31–e34 (2017).
19. Rhie, A., Walenz, B. P., Koren, S. & Phillippy, A. M. Merqury: reference-free quality, completeness, and phasing assessment for genome assemblies. *Genome Biol.* **21**, 245 (2020).
20. Challis, R. J., Kumar, S., Stevens, L. & Blaxter, M. GenomeHubs: simple containerized setup of a custom Ensembl database and web server for any species. *Database* **2017**, (2017).
21. Dolezel, J., Bartos, J., Voglmayr, H. & Greilhuber, J. Nuclear DNA content and genome size of trout and human. *Cytometry. Part A: the journal of the International Society for Analytical Cytology* vol. 51 127–8; author reply 129 (2003).
22. Cechova, M. *et al.* Dynamic evolution of great ape Y chromosomes. *Proc. Natl. Acad. Sci. U. S. A.* **117**, 26273–26280 (2020).
23. Cheng, H. *et al.* Haplotype-resolved assembly of diploid genomes without parental data. *Nat. Biotechnol.* **40**, 1332–1335 (2022).
24. Jain, C., Koren, S., Dilthey, A., Phillippy, A. M. & Aluru, S. A fast adaptive algorithm for computing whole-genome homology maps. *Bioinformatics* **34**, i748–i756 (2018).
25. Rautiainen, M. & Marschall, T. GraphAligner: rapid and versatile sequence-to-graph alignment. *Genome Biol.* **21**, 253 (2020).
26. Nurk, S. *et al.* The complete sequence of a human genome. *Science* **376**, 44–53 (2022).
27. Kolmogorov, M., Yuan, J., Lin, Y. & Pevzner, P. A. Assembly of long, error-prone reads using repeat

graphs. *Nat. Biotechnol.* **37**, 540–546 (2019).

28. Vasimuddin, M., Misra, S., Li, H. & Aluru, S. Efficient architecture-aware acceleration of BWA-MEM for multicore systems. in *2019 IEEE International Parallel and Distributed Processing Symposium (IPDPS)* (IEEE, 2019). doi:10.1109/ipdps.2019.00041.
29. Jain, C., Rhie, A., Hansen, N. F., Koren, S. & Phillippy, A. M. Long-read mapping to repetitive reference sequences using Winnowmap2. *Nat. Methods* **19**, 705–710 (2022).
30. Koren, S. *et al.* Canu: scalable and accurate long-read assembly via adaptive -mer weighting and repeat separation. *Genome Res.* **27**, 722–736 (2017).
31. Mc Cartney, A. M. *et al.* Chasing perfection: validation and polishing strategies for telomere-to-telomere genome assemblies. *Nat. Methods* **19**, 687–695 (2022).
32. Jain, C. *et al.* Weighted minimizer sampling improves long read mapping. *Bioinformatics* **36**, i111–i118 (2020).
33. Sahakyan, A. B. *et al.* Machine learning model for sequence-driven DNA G-quadruplex formation. *Sci. Rep.* **7**, 14535 (2017).
34. bedtools: a powerful toolset for genome arithmetic. https://bedtools.readthedocs.io/en/latest/index.html#.
35. Li, H. Minimap2: pairwise alignment for nucleotide sequences. *Bioinformatics* **34**, 3094–3100 (2018).
36. Harris, R. S. *Improved Pairwise Alignment of Genomic DNA*. (2007).
37. Armstrong, J. *et al.* Progressive Cactus is a multiple-genome aligner for the thousand-genome era. *Nature* **587**, 246–251 (2020).
38. Hickey, G., Paten, B., Earl, D., Zerbino, D. & Haussler, D. HAL: a hierarchical format for storing and analyzing multiple genome alignments. *Bioinformatics* **29**, 1341–1342 (2013).
39. Nguyen, L.-T., Schmidt, H. A., von Haeseler, A. & Minh, B. Q. IQ-TREE: a fast and effective stochastic algorithm for estimating maximum-likelihood phylogenies. *Mol. Biol. Evol.* **32**, 268–274 (2015).
40. Kalyaanamoorthy, S., Minh, B. Q., Wong, T. K. F., von Haeseler, A. & Jermiin, L. S. ModelFinder: fast model selection for accurate phylogenetic estimates. *Nat. Methods* **14**, 587–589 (2017).
41. Hoang, D. T., Chernomor, O., von Haeseler, A., Minh, B. Q. & Vinh, L. S. UFBoot2: Improving the Ultrafast Bootstrap Approximation. *Mol. Biol. Evol.* **35**, 518–522 (2018).
42. Earl, D. *et al.* Alignathon: a competitive assessment of whole-genome alignment methods. *Genome Res.*

**24**, 2077–2089 (2014).

96. Li, H. *seqtk: toolkit for processing sequences in FASTA/Q formats*. (2023).

102. Marçais, G. *et al.* MUMmer4: A fast and versatile genome alignment system. *PLoS Comput. Biol.* **14**,

e1005944 (2018).

108. Wickham, H. *ggplot2: Elegant Graphics for Data Analysis*. (Springer, 2016).

109. R Core Team. *R: A Language and Environment for Statistical Computing*. (R Foundation for Statistical Computing, 2023).

110. Wickham, H. *ggplot2: Elegant Graphics for Data Analysis*. (Springer, 2016).

132. Hebditch, M., Carballo-Amador, M. A., Charonis, S., Curtis, R. & Warwicker, J. Protein–Sol: a web tool for

predicting protein solubility from sequence. *Bioinformatics* **33**, 3098–3100 (2017).

146. Wilson Sayres, M. A., Lohmueller, K. E. & Nielsen, R. Natural selection reduced diversity on human y

chromosomes. *PLoS Genet.* **10**, e1004064 (2014).

156. Dainat, J., Hereñú, D. & Pucholt, P. AGAT: Another Gff Analysis Toolkit to handle annotations in any GTF. *GFF format. Zenodo*.

157. Ranwez, V., Douzery, E. J. P., Cambon, C., Chantret, N. & Delsuc, F. MACSE v2: Toolkit for the Alignment of Coding Sequences Accounting for Frameshifts and Stop Codons. *Mol. Biol. Evol.* **35**, 2582–2584 (2018).

158. Kosakovsky Pond, S. L. *et al.* HyPhy 2.5—A Customizable Platform for Evolutionary Hypothesis Testing Using Phylogenies. *Mol. Biol. Evol.* **37**, 295–299 (2019).

159. Castresana, J. Selection of conserved blocks from multiple alignments for their use in phylogenetic analysis. *Mol. Biol. Evol.* **17**, 540–552 (2000).

160. Storey, J. D. & Tibshirani, R. Statistical significance for genomewide studies. *Proc. Natl. Acad. Sci. U. S. A.*

**100**, 9440–9445 (2003).

161. Murga-Moreno, J., Coronado-Zamora, M., Hervas, S., Casillas, S. & Barbadilla, A. iMKT: the integrative McDonald and Kreitman test. *Nucleic Acids Res.* **47**, W283–W288 (2019).

162. Hallast, P. *et al.* Great ape Y Chromosome and mitochondrial DNA phylogenies reflect subspecies structure and patterns of mating and dispersal. *Genome Res.* **26**, 427–439 (2016).
